# The genomic formation of First American ancestors in East and Northeast Asia

**DOI:** 10.1101/2020.10.12.336628

**Authors:** Chao Ning, Daniel Fernandes, Piya Changmai, Olga Flegontova, Eren Yüncü, Robert Maier, N. Ezgi Altınışık, Alexei S. Kassian, Johannes Krause, Carles Lalueza-Fox, Andrea Manica, Ben A. Potter, Martine Robbeets, Kendra Sirak, Veronika Siska, Edward J. Vajda, Leonid A. Vyazov, Ke Wang, Lixin Wang, Xiyan Wu, Xiaoming Xiao, Fan Zhang, David Reich, Stephan Schiffels, Ron Pinhasi, Yinqiu Cui, Pavel Flegontov

## Abstract

Upward Sun River 1, an individual from a unique burial of the Denali tradition in Alaska (11500 calBP), is considered a type representative of Ancient Beringians who split from other First Americans 22000–18000 calBP in Beringia. Using a new admixture graph model-comparison approach resistant to overfitting, we show that Ancient Beringians do not form the deepest American lineage, but instead harbor ancestry from a lineage more closely related to northern North Americans than to southern North Americans. Ancient Beringians also harbor substantial admixture from a lineage that did not contribute to other Native Americans: Amur River Basin populations represented by a newly reported site in northeastern China. Relying on these results, we propose a new model for the genomic formation of First American ancestors in Asia.

**One Sentence Summary:** Ancient Beringians do not form the deepest American lineage, but harbor admixture from Amur River Basin populations.

## Main Text

The peopling of America remains an area of active debate in archaeogenetics (*1, 2*): the time and place of the “Beringian standstill” (*3-6*), dates and routes of the first colonization (*5, 7-9*), and the Asian source for the later migration about 5000 cal yr BP (*4, 10, 11*) are all contentious. Two recent discoveries are critical for constraining models of American settlement (*2*). First, the Upward Sun River 1 individual (USR1) from a unique double infant burial of the Denali tradition dated to ca. 11500 cal yr BP (*12*) was subjected to deep (17×) whole-genome sequencing and was shown to represent a previously unknown genetic lineage (Ancient Beringians) that split from the American stem between 22000 and 18000 cal yr BP (*4*), much earlier than the two major clades known before, the northern Native Americans (NNA) and southern Native Americans (SNA) that split between 17500 and 14600 cal yr BP (*13-19*). The region where the divergence of Ancient Beringians took place remains unknown, but Moreno-Mayar *et al*. (*4*) favored a model where the divergence occurred in Alaska, with the northern and southern American clades diverging south of the ice sheets. Other genomes relevant for reconstructing the earliest history of proto-Americans in Asia belong to an individual from the Duvanny Yar site (9800 cal yr BP), Kolyma River, northeastern Siberia (*10*) and an older individual from the Ust#x2019;-Kyakhta-3 site (14000 cal yr BP) in the Baikal region (*20*). Both the Kolyma1 and Ust#x2019;-Kyakhta individuals were modeled as proto-American lineages diverging deeper than USR1 and having Siberian admixture of 21-27% (*20*). According to Sikora *et al*. (10), this Paleo-Siberian population represented by the Kolyma1 individual was later largely replaced by the Neo-Siberians, with individuals from a 7,600-year-old site at Devil#x2019;s Gate Cave in the Russian Far East being their type representatives (*10, 21*). The Kolyma1-related genetic component now is prominent only among Chukotkan, Kamchatkan and some West Siberian indigenous groups (Kets, Selkups, Khanty, Mansi), and in ancient Paleo-Inuit (*10*).

To date, our knowledge about prehistoric populations from mainland East Asia is relatively limited. Six Early Neolithic individuals (∼7600 cal yr BP) from Devil#x2019;s Gate Cave in the Amur River Basin (ARB) were sequenced at coverage ranging from 0.1 to 6.6× and were shown to be genetically continuous with present-day populations in the ARB and with a majority of present-day Siberians (*10, 21*). Other genetically similar individuals from the ARB were reported by Ning *et al*. (*22*) and Wang *et al*. (*23*): 28 individuals dated between ca. 7400 cal yr BP and 11^th^-13^th^ centuries CE. To investigate the prehistory of the ARB deeper in time, we generated whole-genome shotgun data from 12 individuals from the Houtaomuga site in Northeast China, who lived between 12000 and 2300 cal yr BP. The oldest individual dated to ∼12000 cal yr BP was sequenced to a high coverage (45×) and is the earliest sequenced representative of the ARB genetic cluster.

We developed a new method for admixture inference based on exhaustive “mapping” of a target group on a “skeleton” admixture graph as a mixture of an increasing number of sources and another method that searches the graph space automatically. To allow comparison of less graph models of different complexity, we undertook a study on simulated genetic data (Supplementary Texts 1 and 2) and developed a new model-comparison approach resistant to overfitting. We utilized the new data from the Houtaomuga site for constraining admixture proportions in a graph connecting key Siberian, East Asian, and American lineages. Leveraging the power of that admixture graph and the novel methods, we demonstrate that the USR1 lineage descends from admixture between NNA and ARB populations represented by the Houtaomuga group, questioning the monocladality of USR1 with Native Americans suggested previously (*4*).

## Results

### An archaeogenetic transect at the Houtaomuga site

An initial sample screening step was performed on 22 ancient individuals from the Houtaomuga site, in the Jilin province of China (Fig. 1, Supplementary Text 3). Based on endogenous ancient DNA yields and authentication (see Supplementary Text 4), the libraries of 12 individuals (two Early Neolithic (EN), four Middle Neolithic (MN) and six Late Bronze Age/Early Iron Age) were sequenced to an adequate coverage (0.1-45x) on the Illumina HiSeq X10 platform (Data S3). The genomic coverages obtained ranged between 0.51–1.40-fold for the Early Neolithic-2, Middle Neolithic, and Iron Age specimens, while a ∼12000-year-old Early Neolithic-1 individual was sequenced to a coverage of 45-fold from a UDG-treated library. Deamination frequencies on the 5#x2019; and 3#x2019; ends of the sequences of these 12 individuals ranged from 6.8 to 40.7%, values typical of aDNA (Data S3).

**Fig. 1.**
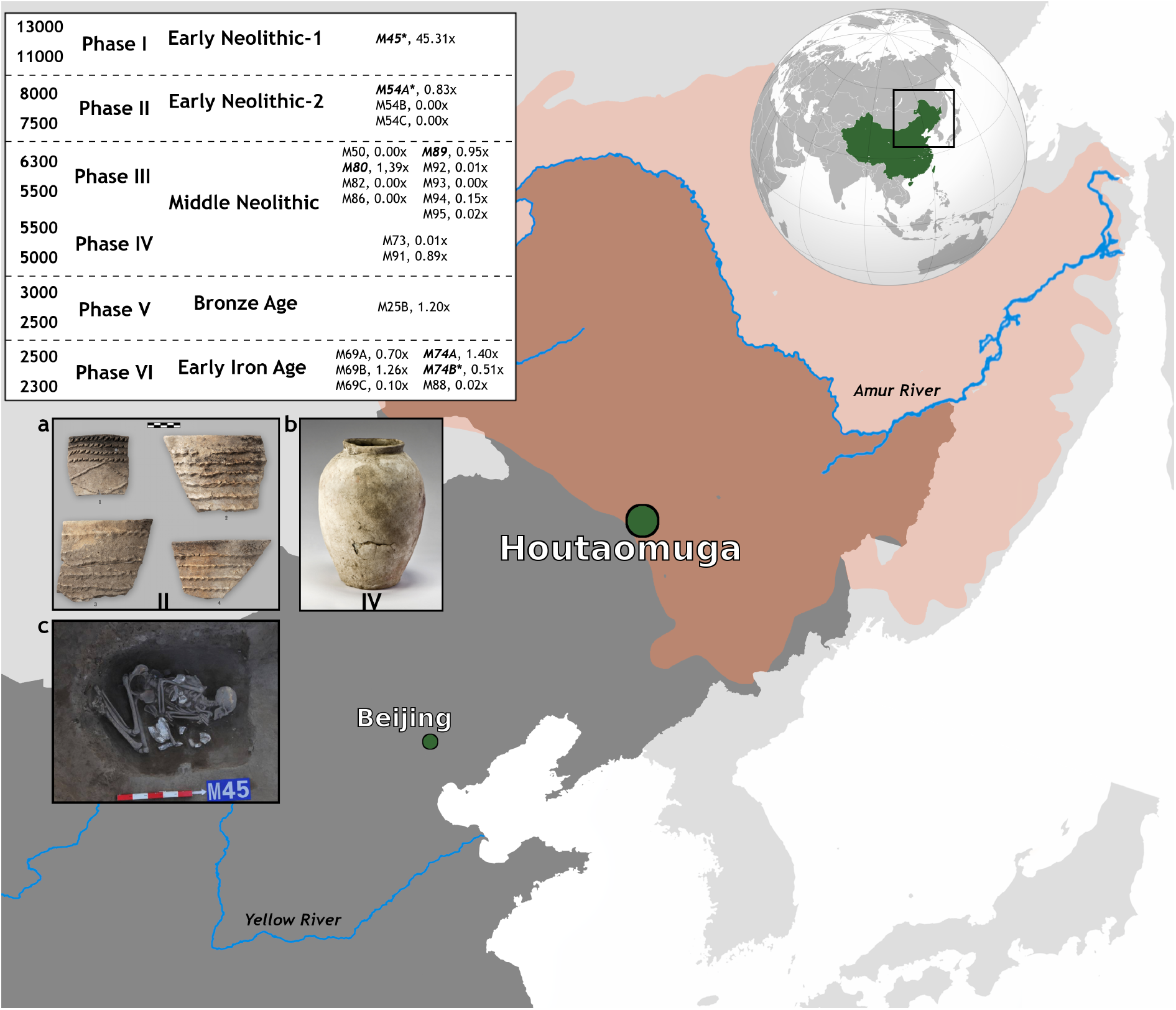
Geographic location of the Houtaomuga site analyzed in this study. The table in the inset includes chronological information and genomic coverage for all screened individuals. Bold and italicized individual codes denote individuals used for population genetic analysis; asterisks denote individuals used for *qpGraph* modeling. Dates are based on confidence intervals from radiocarbon dating results (Data S3). (**A, B**) Ceramic shards and vessels from Phases II and IV, respectively. (**C**) The burial of individual M45. The Amur river basin is marked by color.

Based on contamination estimates, some samples were not included into the reference Houtaomuga group used for admixture graph analyses, leaving a final set of six (Fig. 1, Data S3). Among these six Houtaomuga individuals, three (M45, M54A, M74B) had no detectable West Eurasian admixture according to an analysis based on a simple admixture graph (Supplementary Text 5). We leveraged this feature of Houtaomuga individuals for constructing an admixture graph uniting diverse Siberian lineages admixed with various West Eurasian groups (Supplementary Text 5). For all the downstream graph-based analyses, we created two alternative Houtaomuga groups with pseudohaploid genotype calls: 1/ M45 and M54A; 2/ those two EN individuals and an Iron Age individual M74B.

According to a principal component analysis (PCA), all nine Houtaomuga individuals passing a missing rate cut-off clustered with other Early Neolithic ARB individuals from the Devil#x2019;s Gate Cave and Boisman sites (*10, 23*) and were close to a cluster composed of present-day Tungusic and Nivkh speakers in the ARB (Fig. S41). These results suggest long-term genetic continuity in the ARB, in line with previous studies (*10, 22, 23*).

### A new admixture cline between Asians and Americans

We assembled a dataset based on a panel of 1.24 million SNPs across the human genome (*24, 25*) and generated two versions: a version without transition polymorphisms including 208,649 variable autosomal sites and a full version including 1,062,979 sites (Supplementary Text 5). In the PC1/PC2 space both USR1 and Kolyma1 individuals lie between the American and East Asian clusters, with Kolyma1 located much closer to the Asian cluster (Fig. 2). This result remained unchanged when different dataset versions and PCA protocols were used (Supplementary Text 6, Fig. S40, Fig. S51), and the same results were reported before (*4, 10, 20*). As shown for the simulated USR1 individual, her position in the PC space cannot be interpreted as evidence for or against Siberian admixture since a similar position was reproduced in the absence of Siberian-American gene flow (Fig. 2, Supplementary Text 2). Whole diploid human genomes without missing data were simulated using *msprime* (*26*) according to the admixture graph shown in Fig. S43c (see also Fig. S21, Table S8, Supplementary Text 2).

**Fig. 2.**
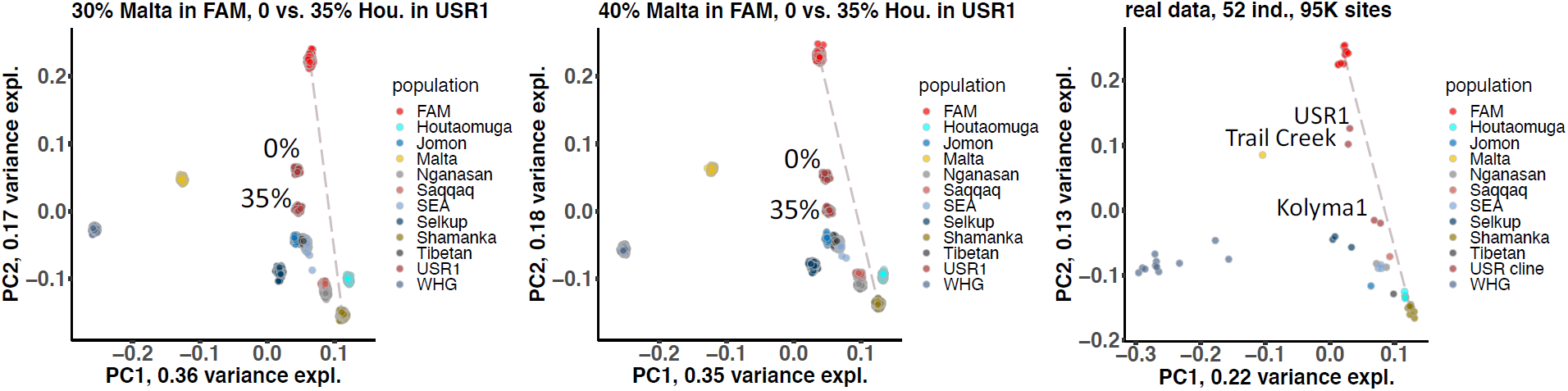
PC1/PC2 plots constructed using PLINK on simulated (**A**,**B**) and real data (**C**). PCs were calculated using 50 simulated ind. and the same number of real counterparts. In the case of the real data, 3 USR1-related individuals were added: Trail Creek and two versions of the Kolyma1 genome (labelled in panel **c**). Dataset sizes in terms of sites were also nearly identical, ca. 95,000. Centroids of the FAM and Shamanka EN clusters are connected with a dashed line. Plots for 100 simulation iterations were overlapped. Panels **A** and **B** show results for different ratios of Mal#x2019;ta ancestry in the simulated FAM and USR1 groups: 30/30% and 40/30%. Clusters of USR1 points are labelled according to the simulated proportion of Houtaomuga ancestry.

Both USR1 and Kolyma1 individuals were modelled to date as mixtures of a proto-American lineage (abbreviated as FAM_EA_ below) and a West Eurasian lineage (*4, 10*). The Kolyma1 individual was alternatively modelled as a mixture of a proto-American, a Siberian, and a West Eurasian lineage, and the same model was proposed for the Ust#x2019;-Kyakhta individual (*20*). To check if these simple models fit the data, we applied the newly developed admixture inference protocol based on admixture graphs to these key individuals. We constructed a skeleton admixture graph for Siberians, East Asians, and Americans, as described in Supplementary Text 5. Building a graph for Siberians is difficult because most groups have a substantial amount of West Eurasian ancestry coming from different sources and probably in multiple waves (*27*). We decided to simplify the graph and used distal sources to account for almost any type of West Eurasian ancestry: Mal#x2019;ta-like, West European hunter-gatherers (WHG), and the basal Eurasian ghost source (*28*). We added these three gene flows by default to each Siberian branch on the skeleton graph. Two similar versions of the skeleton graph were constructed, including 14 groups on the dataset without transitions (Fig. 3a,b, Fig. S37) or 16 groups on the complete dataset (Fig. S39).

**Fig. 3.**
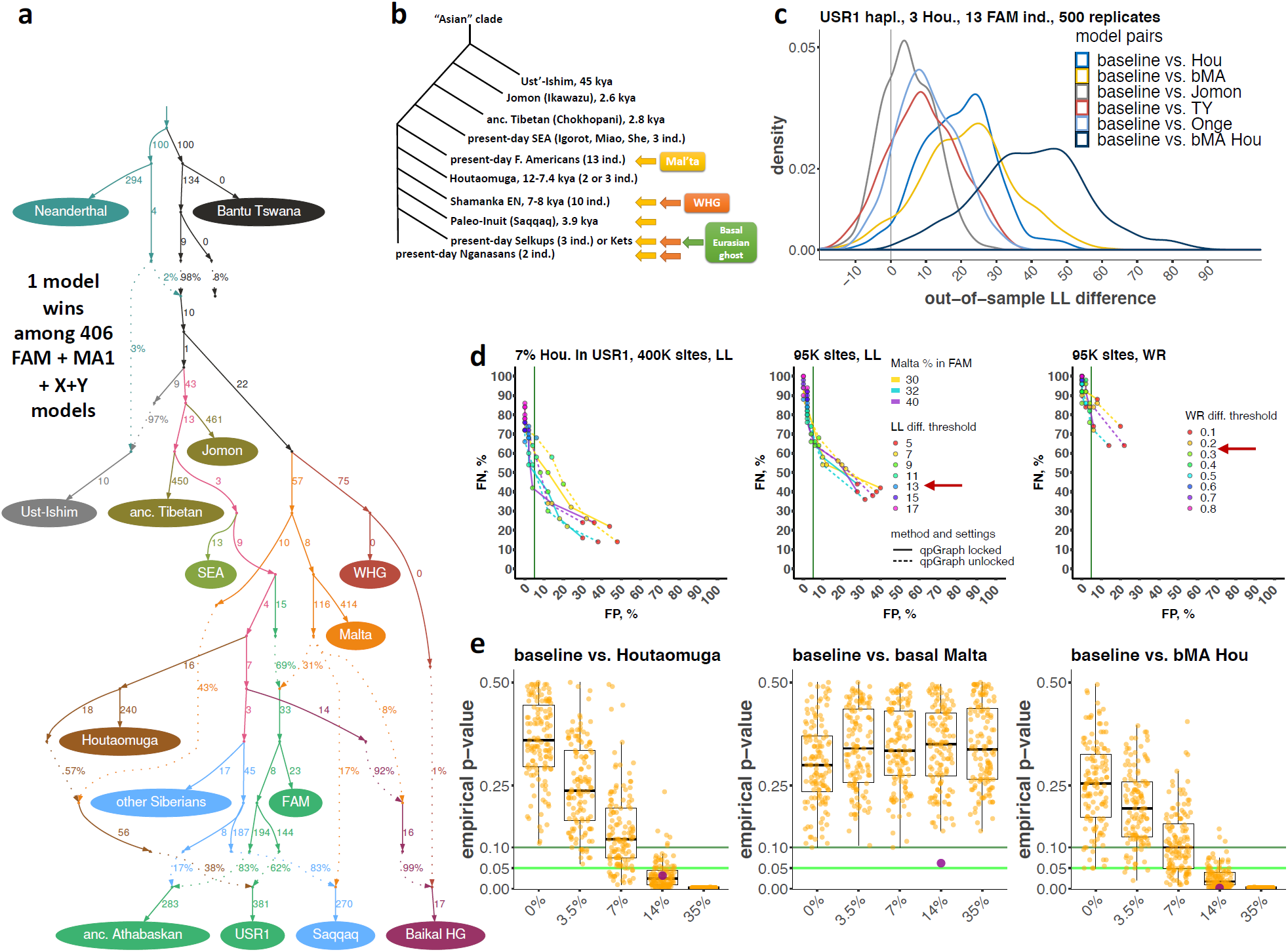
Modelling the genetic history of the USR1 individual. The graph in (**A**) presents a model in the 4-way admixed class that is significantly different from the other 405 models of that class. Panel (**B**) displays the branching order in the Asian clade and the West Eurasian gene flows incorporated into the skeleton model. In panel (**D**) false positive and false negative rates of the graph mapping protocol on simulated genetic data are shown (baseline and “FAM_EA_ + Mal#x2019;ta + X” models compared). LL and WR difference thresholds are coded by point colour, and the optimized thresholds are marked by arrows. Panel (**E**) shows results of out-of-sample testing of admixture graph models on resampled replicates of simulated data. Each dot shows a one-tailed empirical *p*-value. The analysis was repeated for 130 replicates of the history and for different proportions of <u>Houtaomuga</u> ancestry in <u>USR1</u>. The bright green line marks the significance threshold of 0.05. Model comparison *p*-values on real data (ca. 407,000 sites across all groups) are shown as magenta dots in panel **E**, and the distributions of out-of-sample LL differences across 500 bootstrap replicates are shown in panel **C** for six model pairs labelled according to the legend.

Although the skeleton graph presented here may be a simplification of the real situation, it fits the data (Supplementary Text 7d), and we believe it is reasonably powerful for inferring admixture in Asians and Americans as demonstrated on genetic data simulated using *msprime* (Supplementary Text 2) and on positive controls (ancient and present-day Athabaskan and Eskimo-Aleut speakers, and 10,000 years-old individuals from Brazil) described in Supplementary Text 8. Moreover, the same branching order of Asian lineages (Onge, Tianyuan, Jomon, Tibetans, Southeast Asians, Americans and Siberians, see Table S16) was inferred using a different approach (*TreeMix*) (*29*).

We mapped USR1 (Supplementary Text 7), Kolyma1, and Ust#x2019;-Kyakhta individuals (Supplementary Text 9) on both skeleton graph versions. A sister group of FAM_EA_ and a Mal#x2019;ta-related source were assigned as default ancestry sources for the USR1 individual (*4*), and a Mal#x2019;ta-related source was added by default to Kolyma1 and Ust#x2019;-Kyakhta following previous results (*10, 20*). We added one or two additional ancestry sources to each target, testing all possible source combinations within the clade of anatomically modern humans. In the case of USR1 (Data S4a, S4b), only one model in each class was significantly different from others according to the likelihood cut-off of 3 ln-units (*11*), and each additional gene flow resulted in a log-likelihood (LL) difference and/or difference in the worst model residuals (WR) that was associated with a false positive (FP) rate below 5% on data simulated according to the 14-population skeleton graph (Fig. 3d, Table S18, Supplementary Text 2). The mapping protocol was repeated under eight or four setting combinations as detailed in Supplementary Text 7, showing consistent results.

A final model for USR1 was as follows: FAM_EA_ + a Mal#x2019;ta source that for simplicity contributes to all American and Siberian groups (Figs. S37, S39) + a distinct Mal#x2019;ta-related lineage (termed “basal Mal#x2019;ta” for convenience) + a Houtaomuga-related lineage. Exploration of the likelihood space of admixture proportions has put a lower bound on the Houtaomuga-related ancestry in USR1 at ∼6% and the upper bound at 35% (Data S4e, S5). We note that an *f4*-statistic (First Americans, USR1; Southeast Asians, Houtaomuga) deviating from 0 by 3 SE points to attraction between USR1 and Houtaomuga (Table S17). Similar *D*-statistics (a First American group, USR1; a Siberian group, Han) showed the same signal in the original report, but since absolute Z-scores remained below 3.3 the signal was interpreted as a “slight residual affinity” (*4*).

The best model obtained for the Kolyma1 individual was Houtaomuga + FAM_EA_ + Mal#x2019;ta (Data S7a, S7b). The Houtaomuga-like ancestry proportion in Kolyma1 ranged from 37% to 55% (across four setting combinations and both diploid and pseudohaploid genome versions), and the FAM_EA_ ancestry proportion ranged from 19% to 38% (Supplementary Text 9). This result broadly agrees with the previous report (*20*) where 21% ARB ancestry was inferred in Kolyma1. We support the main result by Yu *et al*. (*20*), namely that the Ust#x2019;-Kyakhta individual represents a divergent proto-American lineage, but we are able to model this individual in a simpler way, without the Siberian gene flow (Supplementary Text 9). A low-level (∼6%) gene flow from Kolyma1 or Ust#x2019;-Kyakhta to the Saqqaq lineage has received statistical support according to our criteria, although the direction of the gene flow was impossible to resolve (Supplementary Text 9). In contrast, the direction of the Houtaomuga-USR1 gene flow was well-resolved (Supplementary Text 7). The Kolyma1-Saqqaq relationship was reported before (*10*) and is entirely plausible geographically and chronologically.

Next, we performed an exhaustive exploration of *qpAdm* models for USR1, FAM, and Houtaomuga on the full 1240K-derived dataset, in the context of 12 other groups included into the simpler skeleton graph version (Supplementary Text 10). Using *qpWave*, we tested the cladality of USR1 and various FAM groups: a group of present-day SNA individuals (*11*) used for most admixture graph analyses in this study, 11 alternative present-day SNA groups, and 45 alternative groups composed of ancient SNA and NNA individuals (*13, 14, 18*). The cladality of First Americans and USR1 was rejected for 29 to 33 of 57 groups, depending on settings (*p*-values ranged from 6×10^-7^ to 0.01, Data S8a). The FP rate of this test on data simulated under the 14-population skeleton graph was 7% in the presence of ascertainment bias (*p*-value cut-off = 0.01, 100 datasets simulated using *msprime*, 400,000 randomly sampled sites), and <2% in the absence of ascertainment bias. To determine if the FAM or USR1 group is responsible for the cladality violation, we tested all possible 2- and 3-way admixed models for USR1, the composite FAM group, and Houtaomuga as targets. Among 273 2-way models tested for these targets using *qpAdm* with the “rotating” outgroups setup, no model satisfied the following conditions (*30*): *p*-value > 0.01 and inferred admixture proportions ± 2 SE lie between 0 and 1 for at least one setup. Among 1,092 3-way models tested for the three targets, just one model satisfied these conditions: USR1 = 45±20% FAM + 35±13% Houtaomuga + 20±7% Mal#x2019;ta (Data S8b-d). The FP rate of this *qpWave/qpAdm* procedure on data simulated using *msprime* was 5% in the presence of ascertainment bias and <2% in the absence of ascertainment bias. In other words, for 5 out of simulated 100 datasets i/ the cladality of FAM and USR1 was rejected and ii/ only one of three alternative targets was successfully modeled as a 2- or 3-way mixture according to the criteria outlined above.

Finally, we applied a new model-comparison approach resistant to overfitting (see Materials and Methods) to the best graph models selected above for USR1. The baseline model for USR1 (*4*) was compared to 3-way models “FAM_EA_ + Mal#x2019;ta + basal Mal#x2019;ta” and “FAM_EA_ + Mal#x2019;ta + Houtaomuga", and those were compared to the 4-way model “FAM_EA_ + Mal#x2019;ta + basal Mal#x2019;ta + Houtaomuga” (Supplementary Text 11). Here we focus on two model comparisons where the Houtaomuga-related ancestry source was added. When comparing distributions of out-of-sample LL differences across 500 bootstrap replicates (Fig. 3c, Fig. S49), the “FAM_EA_ + Mal#x2019;ta + Houtaomuga” model fitted the data significantly better (*p*=0.032) than the baseline model for the pseudohaploid USR1 version (Fig. 3c,e). The magnitude of the signal observed on real data in the context of signals on simulated data (Fig. 3e, Table S20) is consistent with the constraints on the ARB ancestry proportions in USR1 (6 to 35%) found by exploring the admixture graph likelihood space (Data S4e, S5).

The gene flow that we revealed in the USR1 lineage challenges previous results which show that the First American source of USR1 forms the deepest lineage in the American clade that bifurcated ca. 20000 years ago in Beringia (*4*). We tested all possible topologies for USR1, an ancient Athabaskan individual (I5321) admixed with Paleo-Inuit (*11*) and representing the NNA clade, a Clovis individual (Anzick) (*16*), and an ancient Peruvian individual (IL7) demonstrating no signal of extra Asian admixture (Data S6). The latter two individuals represented the SNA clade. This American clade with all the necessary Asian and West Eurasian gene flows was grafted on the skeleton graph, and 15 possible topologies were tested under 48 setting/dataset/skeleton graph combinations (Supplementary Text 7g, Data S4f, S4g). This model competition has demonstrated that the basal position of the USR1 source was either significantly rejected under all setting combinations or, if it was not rejected, it invariably resulted in a trifurcation in the American clade. The most supported topology that also almost never produced trifurcations was ((I5321, USR1), (Clovis, IL7)). Thus, the American source of USR1 probably belongs to the NNA clade, in agreement with the geographic location in Alaska. A similar topology exploration incorporating the Kolyma1 and Ust#x2019;-Kyakhta individuals revealed the following topology as a “winner” having no trifurcations in the proto-American clade: ((Ust#x2019;-Kyakhta, Kolyma1), ((I5321, USR1), IL7)) (Supplementary Text 9).

When constructing the skeleton graphs (Supplementary Text 5) and conducting mapping and branching order exploration (Supplementary Texts 7, 8, 9), we used certain conventions that helped to standardize the modelling process, but those conventions make the resulting models unrealistic in certain ways. West Eurasian gene flows added to Siberian and American lineages were always right below the leaf nodes, and they were added separately to each Siberian and American branch. For instance, there was no Mal#x2019;ta-related gene flow into the common ancestor of First Americans and USR1. To check if the results generated using these skeleton graphs hold for more realistic topologies, we performed another model comparison. We showed that the ARB=>USR1 gene flow is also supported in the context of a graph incorporating separate NNA and SNA lineages and a Mal#x2019;ta-related gene flow into the common ancestor of Americans (Supplementary Text 12). When all the necessary extra-American gene flows were incorporated, all three alternative branching orders in the American clade displayed nearly identical model likelihoods, and the branching order (proto-USR1, (proto-Athabaskan, SNA)) again invariably resulted in a trifurcation, in contrast to the other branching orders (Supplementary Text 12). The resulting best model (SNA, (proto-Athabaskan, USR1)) is shown in Fig. 3a.

To check if our main result is robust to more profound changes in the analytical setup, we constructed a dataset that is expected to be free of most types of bias, such as: 1/ uneven deamination damage across ancient DNA samples that leads to ancient DNA attraction; 2/ reference bias (Günther and Nettelblad 2019), 3/ ascertainment bias caused by the paucity of rare variants (Supplementary Texts 2 and 13) that is typical for SNP panels (*31*); 4/ background selection and biased gene conversion which possibly affect 97% of variable sites in the human genome (*32*); 5/ various levels of missing data across groups. Using the new set of 226,472 G ⇔ C and A ⇔ T polymorphic sites, an entirely different protocol for constructing a skeleton graph and a new set of reference populations (Supplementary Text 13), the ARB=>USR1 gene flow was supported (*p*-value = 0.05) using the bootstrap model-comparison method. The direction of the gene flow was also supported by a significant log-likelihood difference (Supplementary Text 13).

### A complex genetic landscape in the Americas

Strong signals (Supplementary Text 8) of Paleo-Inuit admixture were detected along the Chukotko-Kamchatkan/Eskimo-Aleut/Athabaskan cline (Fig. S42). This finding received attention in previous studies (*4, 10, 11, 15, 17, 33*), but here the signal was detected in an unsupervised way. These targets served as positive controls, as well as Native Americans having well-known colonial European admixture (Figs. S40, S42, Supplementary Text 8).

A controversial “population Y” signal in Native Americans (*13-15, 34, 35*) manifests itself as a low-level gene flow from Tianyuan or Onge lineages. We detected the Tianyuan component in one out of five ancient Brazilians from Lapa do Santo (9000-10000 cal yr BP) analyzed here (Fig. S47, Data S6). These individuals were previously analyzed by Posth *et al*. (*14*), and no signal was found. The Tianyuan admixture proportion inferred by us was low, 6%. Three of four ancient Brazilians from Sumidouro (∼10000 cal yr BP) (*13*) analyzed here showed evidence of gene flows from deeply diverging sources, but several sources were almost equally supported (Data S6). In contrast, for the highest-coverage individual Sumidouro5 only the ancestry sources on the Onge and Tianyuan branches and on the adjacent stem edges were among the winning models (Data S6). The Onge- or Tianyuan-related ancestry proportions in Sumidouro5 were 7% or 8% of the genome (Data S6). A previously unreported Onge/Tianyuan signal (Fig. S47) was found at a low level (2% of the genome) in an ancient individual from Belize (individual ID I3443 from Mayahak Cab Pek) who was dated at 9300 cal yr BP (*14*). In five out of 12 ancient individuals from Lapa do Santo, Sumidouro and Belize, the Onge/Tianyuan gene flows were supported by the bootstrap model-comparison approach: in CP18, CP22, Sumidouro7, I3443, and I5456 (*p*-values ranged from 0.026 to 0.046). In the case of the Sumidouro5 individual, the “population Y” signal was also significant on the set of neutrally evolving sites derived from shotgun sequencing data (Supplementary Text 13).

## Discussion

We question a major result on American pre-history, namely that Ancient Beringians represented by the late Pleistocene/early Holocene USR1 Alaskan individual form the deepest branch in the American clade (*4*). Here we introduce an alternative interpretation of Ancient Beringian ancestry that is significantly more likely than the one suggested by Moreno-Mayar *et al*. (*4*). We revealed a new admixture cline that can be represented as a mixture between NNA and a late Pleistocene Siberian population related to contemporary and later groups in the Amur River Basin (ARB). Only two individuals were confidently placed on this cline: USR1 (11500 cal yr BP) in Alaska and Kolyma1 in Chukotka (9800 cal yr BP). These individuals are roughly contemporaneous with the earliest ARB individual reported here and dated to ∼12000 cal yr BP (Supplementary Text 3). Ancient Beringians were shown to be genetically unique among Americans (*4, 13*), and we offer a different explanation for this uniqueness: a unique admixture event in the history of this group instead of their unique phylogenetic position. We summarize various lines of evidence supporting this admixture event in Table S25.

Lexical remnants of a substrate language presumably spoken on the Denali territory before the Athabaskan radiation ca. 2000 – 3000 cal yr BP show similarity with the Chukotko-Kamchatkan language family (Supplementary Text 14a). This suggests that this substrate pre-Athabaskan language could be a member of the hypothetical Chukotko-Kamchatkan–Nivkh linguistic clade (*36*). There is also evidence for a distant relationship between the latter clade on the Asian side and Salishan and Algic language families in North America (reviewed in Supplementary Text 14b). Since the Nivkh speakers are prominent present-day representatives of the ARB gene pool (Data S10), it is possible that these linguistic traces reflect the Pleistocene gene flow revealed in this study between the ARB cluster and Ancient Beringians.

We believe that the admixture signals in the USR1 and Kolyma1 individuals remained on the verge of statistical significance in previous studies (*4, 10*) because the dates of divergence of the First American and ARB lineages from the Asian stem were separated by a relatively short period likely coinciding with the Beringian standstill (*6, 13*), and the ARB lineage(s) contributing to Kolyma1 and USR1 split from the ARB stem shortly after the emergence of the ARB population. In other words, the phylogeny is nearly star-like and thus hardly amenable to standard analytical methods (*37*). Although without further ancient DNA sampling these split times are hard to estimate, this hypothesis seems probable considering the known divergence dates for First Americans and Siberians (*4, 15, 38*) and the radiocarbon dates of the individuals mentioned above. Notably, another analysis by Moreno-Mayar *et al*. (*4*) revealed a statistically significant signal of Asian admixture in USR1 (Table S25). Four simple demographic coalescent models were tested for population pairs USR1/X using *diCal* (*39*), and a significance threshold for LL difference was optimized on simulated data (*4*). A clean split model was rejected for all partner populations tested: present-day Aymara, Karitiana, Athabaskans, Han, Koryaks and Nivkhs. Remarkably, only one model was significantly better than the other three models for the Han/USR1 and Nivkh/USR1 pairs, and that was a second contact model (*4*). And Nivkhs are probably the closest present-day relatives of the ARB cluster according to our graph mapping results, with 81% of their ancestry derived from it (Data S10). We expect that cryptic gene flows similar to the one detected here are common in published admixture graphs, and these undetected gene flows may be important for deciding between competing archaeological interpretations.

The most parsimonious archaeological interpretation of our results, given the geographic anchor points for early NNA, SNA, Ancient Beringians, and ARB individuals between 13000 and 8000 cal yr BP, is that the divergence of the SNA and NNA clades occurred in Asia, immediately after the end of the Beringian standstill (*6*) that, in fact, could have taken place outside of Beringia (Fig. 4). This model explains the ARB signal in Ancient Beringians without invoking numerous independent crossings of the Bering strait, for which there is no clear archaeological evidence. We hypothesize that SNA ancestors were the first to move to Beringia towards the end of the isolation period, NNA ancestors were the second, and ancestors of Ancient Beringians (an NNA sub-group) were the last to migrate into North America, and thus interacted more with the ARB group ancestors spreading from other refugia at the same time. The hypothesis that the major First American clades diverged in East Siberia, and not in Beringia, also makes the “population Y” signal in present-day and ancient Amazonians (*13, 34*) less surprising since it provides more opportunities for contacts between recently diverged American groups and various Asian groups.

**Fig. 4.**
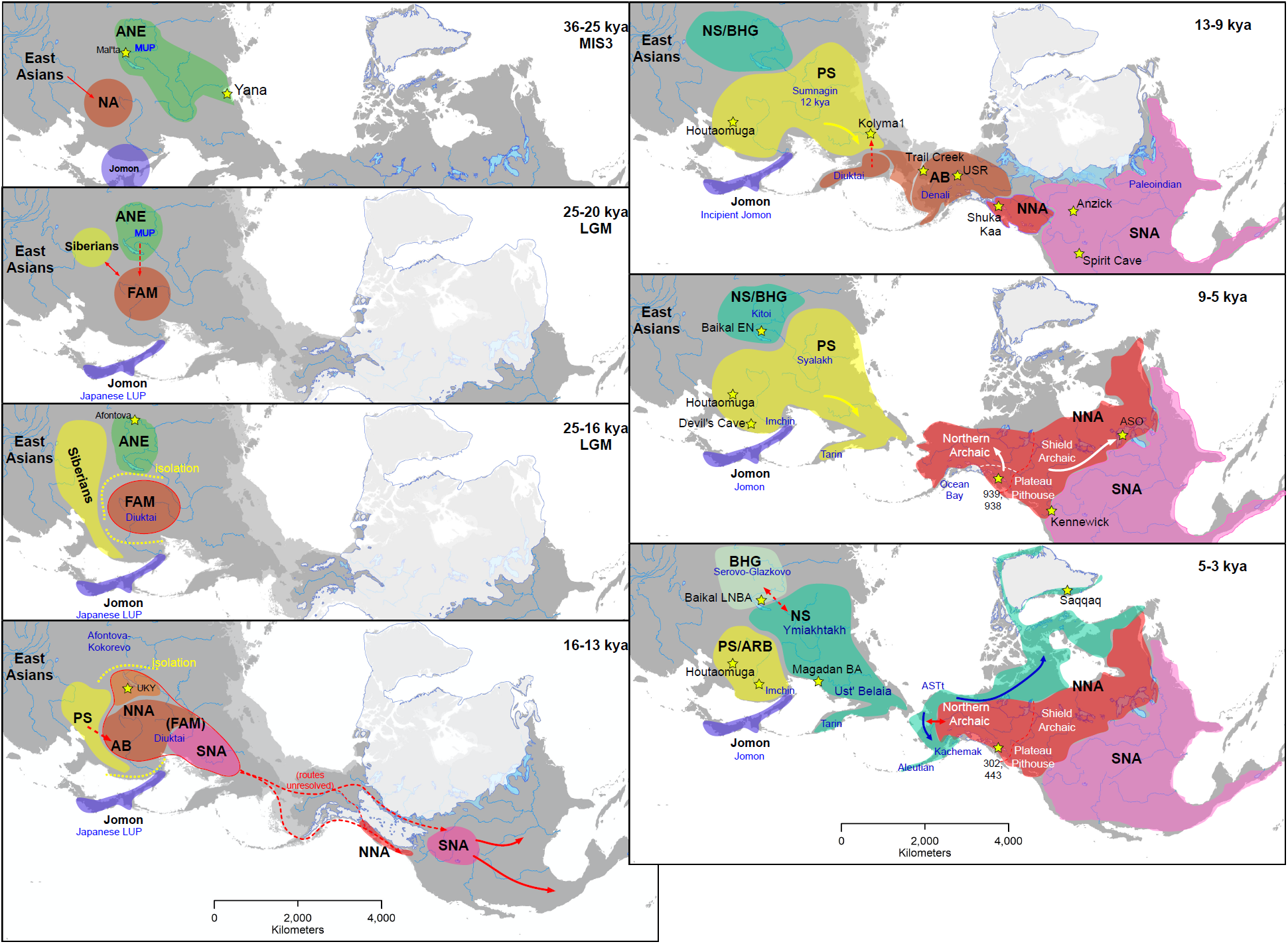
Maps of archaeological cultures and migrations in Northeast Asia, North and Central America. Seven “time slices” are shown. Key archaeological sites and ancient genomes are marked with yellow stars, archaeological cultures with blue text. Past shorelines and glaciers are also shown. The following abbreviations are used: AB, Ancient Beringians *sensu* Moreno-Mayar *et al*. (*4*), ANE, ancient North Eurasians *sensu* Raghavan *et al*. (*40*); BHG, Baikal hunter-gatherers *sensu* Damgaard *et al*. (*41*), FAM, the First American lineage in Asia and America; LGM, last glacial maximum, LUP, Late Upper Paleolithic; MUP, Middle Upper Paleolithic; NNA and SNA, northern and southern Native American lineages *sensu* Raghavan *et al*. (*15*); NS and PS, Neo- and Paleo-Siberians (the meaning of the terms is different from that used by Sikora *et al*. (*10*)); UKY, the Ust#x2019;-Kyakhta-3 site.

## Author contributions

P.F., R.P., and Y.C. supervised the study. L.W., X.X., R.P., and Y.C. assembled the collection of archaeological samples. L.W. was responsible for radiocarbon dating and calibration. C.N., D.F., K.S., R.P., and Y.C. performed laboratory work and supervised ancient DNA sequencing. P.F., C.N., P.C., D.F., O.F., E.Y., N.E.A., C.L.-F., K.W., and S.S., analyzed genetic data, R.M. developed novel software, and B.A.P. prepared archaeological maps. A.S.K. and E.J.V. wrote the supplemental sections on linguistics. P.F., C.N., and R.P. wrote the manuscript with additional input from all other co-authors.

## Data and materials availability

Raw sequence data (bam files) from the 12 newly reported ancient individuals have been deposited at the Genome Sequence Archive at the China National Genomics Data Center under accession number HRA000268. Custom code used in this manuscript is available at a dedicated github repository: https://github.com/uqrmaie1/admixtools.

## Supplementary Materials

Materials and Methods

Supplementary Text, sections 1-14

Figures S1-S55

Tables S1-S25

References

Data S1-S10

