## Supplementary text, figures, and tables for "The genomic formation of First American ancestors in East and Northeast Asia"

**This PDF file includes:**

Materials and Methods
Supplementary Text, sections 1-14
Figs. S1 to S55
Tables S1 to S25
Captions for Data S1 to S10

**Other Supplementary Materials for this manuscript include the following:**

Data S1 to S10

### Contents

**Materials and Methods** 3

**Suppl. Text 1.** Testing *D*-statistics, *qpAdm*, and *qpGraph* on simple simulated histories 8

**Suppl. Text 2.** Testing PCA, *D*-statistics, *qpAdm*, and *qpGraph* on a complex simulated history 20

**Suppl. Text 3.** Archaeological context of the Houtaomuga cemetery 28

**Suppl. Text 4.** DNA sequencing, read mapping, and contamination estimation 32

**Suppl. Text 5.** Building skeleton admixture graphs 34

**Suppl. Text 6.** Principal component analysis 48

**Suppl. Text 7.** Applying the admixture graph mapping approach to the USR1 individual 50

**Suppl. Text 8.** Asian gene flow signals in Native Americans 60

**Suppl. Text 9.** Applying the admixture graph mapping approach to the Kolyma1 and Ust'-Kyakhta individuals 63

**Suppl. Text 10.** *qpAdm* analysis of the USR1 individual 67

**Suppl. Text 11.** A model comparison method for *qpGraph* resistant to overfitting 69

**Suppl. Text 12.** A more realistic graph model of the USR1 history 71

**Suppl. Text 13.** Automated graph space search and a bias-free dataset 72

**Suppl. Text 14.** Linguistic implications of the gene flow between Siberians and Ancient Beringians 76

**Suppl. Figures** 90

**Suppl. Tables** 167

**Suppl. data file captions** 185

#### Materials and Methods

Exploring the performance of *f*-statistic-based methods on a complex simulated history

Although a broad range of methods exists for reconstructing genetic history from genome-wide autosomal data, just few methods became a cornerstone of archaeogenetic studies: principal component analysis or PCA (*42*), *D*-statistics (*37, 43*) and *f*-statistics (*44-46*), *ADMIXTURE* (*47*), *qpWave* (*17*) and *qpAdm* (*48, 49*). *D*- and $f_4$-statistics (which are identical except for the denominator) test if an unrooted four-population tree fits the data, and they are not affected by genetic drift. A deviation of the statistic from 0 means that either the tree topology is wrong, or a gene flow happened between a pair of groups in the tree (in the absence of recurrent mutations or SNP ascertainment bias). However, interpretation of the statistics is ambiguous since prioritizing only one pair of groups that experienced a gene flow is impossible (Lipson 2020 preprint). A more sophisticated admixture inference approach is now used widely, that is a pair of closely related tools named *qpWave* and *qpAdm*. The approach is based on $f_4$-statistics and does not require a detailed knowledge of the population phylogeny beyond few simple assumptions (*28, 48, 49*). This method allows testing combinations of proxy ancestral groups ("sources") for a target, given a diverse set of "outgroups".

The popularity of the methods listed above is explained by their modest computational requirements and versatility: the ability to analyze unphased biallelic genotype data of various types (pseudohaploid or diploid) and generated using targeted enrichment on a panel of sites or shotgun sequencing technologies, and the ability to analyze low-coverage ancient genomes with a high proportion of missing data (*49*). Even though the methods relying on *f*-statistics are central to archaeogenetics (*28, 30, 48, 50-66*), just a handful of studies was devoted to testing their performance in fully controlled conditions, i.e. on simulated genetic data (*13, 45, 49, 51*). The only study reporting detailed testing of *qpAdm* on simulated data is a recent pre-print by Harney *et al.* (*49*). In that study the method was explored under polar conditions (either extremely favourable or extremely unfavourable), and assumption violations and the false positive (FP) rate were not explored.

Since all analytical methods based on *f*-statistics remain untested on realistically complex simulated histories (*13, 45, 49, 51*), we undertook a method development effort guided by a large-scale simulation study. First, we explored if the *D*-statistics, *qpWave/qpAdm*, and *qpGraph* (*45*, Maier *et al.* in preparation) methods behave as expected on two types of simple simulated demographic histories (trees with one admixture event in the history of the target group, Supplementary Text 1). We simulated various demographic histories for 10 or 11 groups (5 individuals per group) ranging from those with branch splits well-separated in time to those with more star-like branch splits, performed 50 independent simulation runs for each history, and subsampled 95,000 or 400,000 diploid variable sites from whole-genome simulations generating 19,500 datasets in total (the number of sites matching typical ancient DNA datasets). In order to reproduce the depletion of rare variants common to SNP panels and associated ascertainment biases (*31*), we applied a global filter on a minor allele frequency (MAF) before randomly sampling variable sites. Sites having MAF <5% were dropped. We demonstrated that under these conditions the methods behave largely as expected (*45, 49*), except for minor biases in certain $f_4$-statistics caused by MAF filtering, that knowing the true phylogeny allows to achieve higher sensitivity in detection of gene flows, and that ranking alternative admixture graph models by log-likelihood, LL, is preferable to ranking by the worst model residual, WR (Supplementary Text 1).

Next, we simulated a complex demographic history with multi-layered admixture (for 15 groups and 54 individuals) that matched a skeleton admixture graph used later in this study for exploring the history of Siberians and Native Americans (Fig. 3a). In total, 3,000 diploid datasets without missing data were generated (95,000 or 400,000 sites, with or without MAF filtering, Supplementary Text 2), and the distribution of individuals in the space of the 1st and 2nd principal components observed on real data was closely reproduced (Fig. 2). Taking advantage of these complex simulated datasets, we first explored the effect of assumption violation on *qpAdm* performance.

We suggest that a conservative interpretation of *qpAdm/qpWave* results should be favoured over the interpretation that became dominant in the literature (*28, 30, 48, 51-66*) since the latter depends on an assumption about the phylogeny that is difficult to control. The conservative interpretation is a strict interpretation of *qpWave* *p*-values (*17*): a *p*-value for an *n*-way model below a threshold means that more than *n* gene flows in any direction connect the outgroups and the tested group of populations. A *p*-value above the threshold cannot be interpreted unambiguously since it may reflect lack of power to reject a particular *n*-way model. Thus, *qpWave* is a tool for rejecting models, but if *qpAdm* is used as a tool for **accepting** models for a particular target group, additional assumptions about the topology of the graph come into play: the proxy source lineages should receive no further gene flow from outgroups after the split with true ancestry sources, i.e. the true and proxy sources should be cladal, and there should be no gene flows from the test groups to outgroups (*28, 48*).

We applied the *qpAdm* admixture inference method as suggested by Harney *et al.* (*49*), using a "rotating" set of outgroups and testing initially the least complex and subsequently more complex models for the target. On the complex simulated histories, we encountered FP rates between 20 and 30% when using *p*-value thresholds commonly employed in published *qpAdm* analyses (0.01 and 0.05). The FP rates mentioned here refer to the whole admixture inference protocol (Supplementary Text 2), and not to a stand-alone *qpWave* or *qpAdm* test. These high FP rates were observed only when the cladality assumption was violated (Supplementary Text 2). A seemingly small-scale deviation from cladality of real and proxy sources (2% of the proxy source genome derived from the "violating" gene flow) was shown to be more serious than a large-scale deviation (10% of the proxy source genome derived from the "violating" gene flow). This behaviour is a combination of two effects. Rejection of the simplest 1-way model for the target can be interpreted as evidence of a gene flow into either the target or the proxy source (and the latter is correct in this case). However, we did not consider the latter possibility, i.e. only one target group was singled out. At the next stage, when two- and three-way models are considered for the target, some unnecessarily complex model do not get rejected, but that might happen because they are either true, or because the method lacks power. Opposite trends in FP rates were encountered when decreasing the *p*-value cut-off in the cases of small-scale vs. large-scale assumption violation, and certain outgroup sets also showed opposite effects (Supplementary Text 2). This behaviour makes method optimization difficult. While the second *qpAdm* assumption (no "backflow") can be satisfied by using outgroups that are much deeper in time than source proxies and targets, the validity of the first assumption ("cladality") is difficult to ensure, especially in high-throughput tests involving dozens of alternative sources and the "rotating outgroup" protocol (*30, 54*, and other recent studies).

The arguments above are valid if a particular group is singled out as a target for modelling (e.g. because of its younger date), and potential ancestry sources are not tested as targets. For circumventing the problem above we propose the following protocol:

i/ take a set composed of a group of interest, potential ancestry sources for this group and "outgroups";

ii/ test cladality of all population pairs using *qpWave* with the "rotating" outgroup setup (*49*); if cladality of the group of interest with any other group is rejected at $p$=0.01, move to the next step;

iii/ test all possible 2-, 3-, and $n$-way models for the group of interest using *qpAdm* with the "rotating" outgroup setup, identify a set of potential source proxies; consider as "fitting" only models demonstrating a $p$-value above 0.01 and inferred admixture proportions ± 2 SE between 0 and 1 (*30*);

iv/ test all possible 2-, 3-, and $n$-way models for the group of interest and its potential sources as targets (with the "rotating" outgroup setup); increase the number of ancestry sources only if no fitting models are found in the current class;

v/ if fitting models are found at the previous step only for the group of interest and not for its potential sources, declare a positive result.

We demonstrated that the FP rate of this procedure on the complex simulated history in the presence of ascertainment bias was 5% ($p$-value cut-off = 0.01, 30% Malta ancestry in FAM and USR1, 400,000 randomly sampled sites with MAF >5%). In other words, for 5 out of 100 datasets simulated under the null hypothesis i/ the cladality of the FAM and USR1 groups was rejected and ii/ only one of three alternative targets (FAM, USR1, Houtaomuga) was successfully modeled as a 2- or 3-way mixture of all possible sources according to the criteria outlined above. The FP rate of this procedure on the same simulated data without the ascertainment bias (50 datasets) was <2% since the cladality of FAM and USR1 was not rejected according to *qpWave* tests. We caution that this procedure was not tested on simulated data mimicking the most challenging "proximal" *qpAdm* setup (*30*) when groups close in time and space are used as targets, sources, and outgroups, and where a network of gene flows between "ingroups" and "outgroups" might be especially complex.

Developing an admixture inference method resistant to overfitting

The popular methods listed above either have implicit assumptions about the population phylogeny that are hard to control (*qpAdm*, see above), or their results are open to many interpretations ($D$/$f_4$-statistics, PCA). Hence answering questions about history often requires explicit modelling of the phylogeny and gene flows, i.e. requires fitting admixture graphs to the data. Diverse graph-fitting methods exist, and *qpGraph* (*45*) is a tool relying on $f$-statistics. It tests if a pre-defined graph fits the matrix of all possible $f_3$-statistics for the incorporated groups, optimizes admixture proportions and edge lengths, and is very fast as compared to similar methods relying on site frequency spectra: *momi v.2* (*67*), *rarecoal* (*11, 68*), and *fastsimcoal* (*69*). Thus, *qpGraph*, especially its recent fast implementation (Maier *et al.* in preparation), allows exhaustive topology exploration for crucial sub-graphs. Unlike most Bayesian methods, *qpGraph* can process any type of biallelic genotype data, which allows key low-coverage ancient individuals to be incorporated. A disadvantage that precluded widespread adoption of the *qpGraph* method is the fact that there is no statistically solid way of comparing models differing in the number of parameters, e.g., gene flows (Lipson 2020 preprint). The best-performing model-ranking metric, log-likelihood (LL) difference (Supplementary Text 1), is susceptible to overfitting.

We used two approaches to deal with overfitting. First, we set the model-comparison thresholds (LL difference and the worst residual, WR, difference) so that the FP rate on simulated data drops below 5% for any dataset/setting combination tested (Supplementary Text 2). If the simulated graph and the graph fitted to real data are identical, if the same classes of models are compared (*n* and *n+1* gene flows) on datasets of similar size, and if the $F_{ST}$ and PCA results on simulated data approach those on real data, we expect the optimized thresholds to yield an FP rate below 5% on real data (Fig. 3d, Table S9). However, this approach is laborious and lacks statistical rigour. Second, we attempted to eliminate the overfitting problem by comparing out-of-sample likelihoods between models of different complexity (Supplementary Text 11). For each bootstrap iteration *i*, we draw bootstrap samples of 5 cM SNP blocks (this block size is a standard in the AdmixTools package) to compute fitted *f*-statistics on this training subset of blocks. We denote the set of SNP blocks which were selected by *I* (~2/3 of blocks) and the set of SNP blocks which were not selected by *O* (~1/3 of blocks), and these sets change with each iteration *i*. The out-of-sample score (LL) of the model is then computed for each bootstrap iteration by comparing *f*-statistics fitted on *I* and those observed on *O*. The difference in out-of-sample LLs for two graphs is calculated for each *i*, and since the distribution of LL differences tends to deviate from normality, an empirical one-tailed *p*-value was computed. We used one-tailed tests since we did not expect that a simpler model would ever demonstrate a significantly better fit than a more complex model. We have shown on the complex simulated datasets that the new method eliminates the overfitting problem, at least if one or two unnecessary gene flows are added (Fig. 3e, Fig. S12-3, Supplementary Text 11). We also optimized the empirical *p*-value threshold based on the data simulated under the baseline model for the USR1 individual and in the presence of ascertainment bias (Supplementary Text 11). The model-comparison method is a part of a new implementation of AdmixTools (AdmixTools 2, Maier *et al.* in preparation). We ensured that model rankings generated by *qpGraph* v. 6412 used throughout this study for consistency, by the latest *qpGraph* version 7365, and by the new AdmixTools 2 implementation are identical, and that LL and WR reported by all the implementations are nearly identical.

Relying on the simulations outlined above, we developed a graph-based method for exhaustive non-greedy exploration of ancestry sources in admixed groups (Supplementary Text 7). A target group is mapped onto all branches of a skeleton admixture graph or its subclade: first as unadmixed, then as a 2-way and 3-way mixture of sources on all possible edges. Other gene flows into the target may be added by default, e.g. the Mal'ta-related admixture in First Americans (*40*). For models having the same number of parameters (alternative branching orders or alternative gene flow sources) a group of best models is defined according to an LL difference threshold of 3 units (*11, 70*). An additional admixture event is considered supported if an LL or a WR improvement exceeds a threshold corresponding to a 5% FP rate on data simulated according to the same skeleton graph/target group combination (Table S9). Alternatively, an additional admixture event is considered supported if a model-comparison empirical *p*-value is significant (below 0.05). A group of best models found in this way can be narrowed down even further by counting the number of 0-length edges corresponding to trifurcations in a clade of interest. Trifurcations are typically an indicator of a mismatch between model and data. To take care of systematic errors, we propose a set of rules for selecting best models across multiple setting combinations tested (Supplementary Text 7).

*qpAdm* analyses and admixture graph mapping on real data

The *qpAdm/qpWave* model selection procedure outlined above was applied to real data in this study (Data S8b-g). Each test was repeated for pseudo-haploid and diploid USR1 versions (4), and for two alternative compositions of the Houtaomuga group (two or three individuals lacking West Eurasian ancestry). Thus, four alternative setups were tested in parallel.

In addition to key individuals USR1 (*4*), Kolyma1 (*10*), and Ust'-Kyakhta (*20*), we mapped on both skeleton graph versions 84 ancient American individuals (*13, 14, 18*) passing a missing rate cut-off of 70% and 13 present-day groups on the 14- and 16-population skeleton graphs using the same setting combinations, significance thresholds and model ranking rules. For details of the mapping procedure see Supplementary Text 8 and for a list of best models see Data S6. We also mapped on the 14-population skeleton graph 230 ancient Eurasian individuals (*10, 11, 30, 41, 53, 71*) and 48 present-day groups (published mostly by Mallick *et al.* (*72*) and Pagani *et al.* (*73*)) passing a missing rate cut-off of 70% and shifted to the Asian cluster in the PC1/PC2 space (Fig. S40). West Eurasian gene flows from the Mal'ta, WHG, and basal Eurasian lineages were added by default to each target to account for diverse West Eurasian sources.

#### Supplementary Text

1. Testing *D*-statistics, *qpAdm*, and *qpGraph* on simple simulated histories

We started from the simple admixture graphs simulated by Moreno-Mayar *et al.* (*13*) and attempted to extend these results. We used *qpGraph* v. 6412, *qpAdm* v. 810 and qpDstat v. 751. The following settings were used for *qpGraph*: outpop, NULL; blgsize, 0.05; lsqmode, NO; diag, 0.0001; hires, YES; initmix, 1000; precision, 0.0001; zthresh, 0; terse, NO; useallsnps, NO. Default settings were used for *qpAdm* and *qpDstat*.

*a. Simulating genetic data*

We simulated two classes of simple demographic histories without multi-layered admixture: those containing many outgroups but just one lineage differentially related to the sources of the gene flow in question, and those containing six lineages differentially related to the sources of the gene flow in question. We call them "information-poor" (Fig. S1) and "information-rich" (Fig. S2) graphs, respectively.

All chromosomes of the human genome were simulated using *msprime* (*26*), with the following settings: recombination rate per bp per generation, $2\times10^{-8}$; mutation rate per bp per generation, $1.25\times10^{-8}$ (*74*); generation time, 29 years (*75*). Fifty independent simulation runs were performed for each setting combination. To mimic the depletion of rare variants typical for widely used SNP panels and associated ascertainment bias, sites having minor allele frequency <5% were removed (corresponding to a minimum allele count of 5 or 6 for the information-poor and information-rich datasets, respectively). Then the datasets were randomly subsampled to 95,000 variable sites to mimic the transition-free dataset used in this study. Alternatively, 400,000 variable sites were sampled to mimic the larger dataset including transitions. This procedure was performed for both graph classes described above.

In both "information-poor" and "information-rich" graphs, only one admixture event was explored in detail: H = I + G (Figs. S1, S2). To explore a range of graphs from nearly star-like to much better resolved ones (*37*), we tested the following combinations of population divergence dates within the clade of interest on the "information-poor" graph, i.e. within the clade uniting F, G, H, and I (Fig. S1): 10/9/8.5 ka; 13/11/10.5 ka; 13/11/8.5 ka; 15/11/8.5 ka. For each combination of divergence dates, we tested three alternative dates for the G→H gene flow: 2.5, 5, and 7.5 ka. Twelve demographic histories were tested in total, and 13 alternative intensities of the gene flow were simulated for each history: 0%, 5%, 10%, 20%, 30%, 40%, 50%, 60%, 70%, 80%, 90%, 95%, 100%. Thus, for the "information-poor" graph we simulated 156 scenarios, 50 iterations each.

The "information-rich" graph (Fig. S2) included six groups differentially related to the two ancestry sources for H: B, C, D, E, F, and J. We tested two alternative divergence dates, 15 and 20 ka, for the clade of interest, i.e. in the clade uniting all groups except for A and Outgroup. For each divergence date, we tested three alternative dates for the G→H gene flow (2.5, 5, and 7.5 ka) and 13 alternative intensities. Thus, for the "information-rich" graph we simulated 78 scenarios, 50 iterations each.

*b. qpGraph tested on the information-poor phylogeny*

The simulated demographic histories were used as input data for calculating *D*-statistics, and for testing admixture models using *qpAdm* and admixture graphs using *qpGraph*. The following five types of graphs were tested: 1) "completely correct", i.e. those matching the topology shown in Fig. S1; 2) "partially correct 1", where directions of the gene flows beyond the clade of interest were reversed (20% B→A and 10% E→D instead of the correct directions 20% A→B and 10% D→E); 3) "partially correct 2", where A was modelled as getting 80% ancestry from B; 4) "incorrect 1", where directions of all the gene flows were reversed (20% B→A, 10% E→D, variable H→G); 5) "incorrect 2", where A was modelled as getting 80% ancestry from B and the G→H gene flow direction was reversed. Since the "information-rich" graph included just one admixture event the "partially correct" topology class was not applicable (Fig. S2). The "partially correct" topologies were introduced to mimic a situation often encountered in practice when a skeleton admixture graph is mildly or grossly incorrect, but a clade of interest is modelled correctly. We also wanted to see under what conditions it is possible to distinguish if the group of interest is a source or a recipient of gene flow, that is why we wanted to introduce the "incorrect" topology classes.

The target group (G in the "incorrect" setup, H in the other setups) was mapped on all possible edges in the skeleton graph as unadmixed or as a mixture of all possible source pairs. Models were ranked according to various metrics (the worst residual, sum of all *f*-statistic residuals, sum of all squared *f*-statistic residuals, log-likelihood).

*c. Finding the best model-ranking metric and threshold for qpGraph*

First, we considered the correct "information-poor" graphs. We tested four alternative model-ranking metrics for distinguishing demographic histories with and without a gene flow event: −ln(likelihood) (LL), sum of all model residuals (absolute Z-scores corresponding to all *f*-statistics) or SR, sum of all squared model residuals or SSR, the worst residual or WR (the Z-score of the worst-fitting *f*-statistic). For each metric, we tested four or five alternative threshold values and captured a trade-off between the false positive and false negative rates using the Matthews correlation coefficient or MCC (Matthews 1975) expressed as: $MCC = \frac{TP\times TN - FP\times FN}{\sqrt{(TP+FP)(TP+FN)(TN+FP)(TN+FN)}}$, where TP is the number of true positive iterations under a given setting, TN is the number of true negative iterations, FP is the number of false positive iterations, and FN the number of false negative iterations. We tested the following thresholds: log-likelihood difference of 3, 4, 5, or 10 units; SR difference of 5%, 10%, 15%, or 20% (calculated as $(SR_{unadm} - SR_{adm})/SR_{adm}$); SSR difference of 5%, 15%, 30%, or 40% (calculated as $(SSR_{unadm} - SSR_{adm})/SSR_{adm}$), and WR difference of 0.1, 0.2, 0.3, 0.5, or 0.8 (expressed in units of standard error intervals). The WR difference threshold of 0.3 SE units was used in a previous publication by Posth *et al.* (*14*). The most stringent thresholds tested were those that under any setting yielded a false positive result for no more than three iterations out of 50 (6% false positive rate). A combination of a model-ranking metric or another admixture inference method and a threshold is here termed "binary classifier" or simply "classifier".

While "mapping" a target group on a skeleton graph, a highest-scoring unadmixed model was compared to a highest-scoring 2-way admixed model. We took the 0% G flow as a true negative baseline and compared the performance of various classifiers (Fig. S3a,c,e,g). As expected (*13, 37*), the performance of the graph mapping protocol was poor on star-like phylogenies, with 10/8.5/7.5 ka graphs being the worst-resolved, and 15/8.5/2.5 ka graphs the best-resolved ones (Fig. S3). Only the most stringent thresholds tested demonstrated nearly ideal performance, with MCC approaching 1 for most gene flow intensities: an LL difference of >10 units, an SR difference of >20%, an SSR difference of >40%, and a WR difference of >0.8 SE. Under a great majority of low-resolution settings (low levels of gene flow) the LL10 classifier surpassed all SR, SSR, and WR classifiers in its performance as measured by MCC. See, for instance, results for the 10/8.5/7.5 ka (Fig. S3a), 13/10.5/5 ka (Fig. S3c), and 15/8.5/2.5 ka graphs (Fig. S3g). We also noticed an asymmetry in performance for low levels of G and I ancestry: low-level G ancestry was easier to detect than low-level I ancestry because on the graph there is only one group (F) differentially related to G and I, and it splits on the G branch. It was generally problematic to detect 5% and 10% G ancestry in H, and 5% to 20% or often even 30% I ancestry in H (Fig. S3, see also Table S1).

In Fig. S4a,c we show an alternative way of presenting these results: the fraction of negative iterations across all intensities of gene flow tested, for the LL10, SR20%, SSR40%, and WR0.8 classifiers. In Table S1 we summarized "confidently detectable admixture proportions" yielding no false negative results among 50 iterations according to the four selected classifiers. Again, we conclude that LL10 outperforms other classifiers in sensitivity to gene flow in low-resolution contexts, while demonstrating false positive rates on a par with the other classifiers (Table S6).

Next, we took a closer look at model ranking. For this purpose, we visualized the top end of the model distribution in the SSR vs. WR and SSR vs. LL coordinates, for each combination of divergence and admixture dates and gene flow intensities. We wanted to assess the properties of most widely used model-ranking metrics, WR and LL. For making a classifier, one needs a metric and a threshold value. But any threshold is useless if the distribution in the SSR vs. WR or SSR vs. LL space is constant or inverted, i.e. if the correct model does not demonstrate the lowest value of a given metric. Thus, we wanted to see how often constant and inverted distributions occur for the WR and LL metrics.

For each iteration, we compared the performance of the metrics in the following way. First, we recorded what types of source pairs fell below chosen LL and WR thresholds (LL3 and WR0.1): "wrong" models where both sources are incorrect, "partially correct" models where only one source is correct, and "correct" models where both sources are correct. The task of ranking models having the same number of parameters (drift and gene flow edges) is different from admixture event detection (Lipson and Reich 2017), thus we chose an LL threshold of 3 ln-units previously used as a statistical significance threshold for comparing models having the same number of parameters (*11*). We also selected a WR threshold that demonstrated similar MCC values and thus provided a similar level of sensitivity (Fig. S3). If, for instance, both correct and partially correct models fell under the chosen WR difference threshold, but only the correct model fell under the chosen LL difference threshold, we concluded that the LL metric "won". Similarly, if correct, partially correct, and incorrect models fell under the chosen WR difference threshold, but only correct and partially correct models fell under the chosen LL difference threshold, we also concluded that the LL metric "won". If only partially correct and/or incorrect models fell under the chosen WR difference threshold, but only the correct model fell under the chosen LL difference threshold, we concluded that the LL metric "won strongly". Second, we recorded the shape of the curve: either "monotonic", or "constant" if at least some points including that corresponding to the correct model formed horizontal line(s), or "inverted" if the correct model did not show the lowest value of the metric. These results are presented in a tabular form in Data S1. See example biplots for high- and low-resolution scenarios in Fig. S5.

The setting 15/8.5/2.5 ka demonstrates the performance of both metrics in nearly ideal conditions (Data S1, Fig. S5a,b). We considered gene flow proportions between 5% and 95%, 550 iterations in total (Data S1). In 500 cases (91%) both metrics performed in the same way given the thresholds of 0.1 SE for WR and 3 ln-units for LL; the WR metric "won" in 6 cases (~1%), the LL metric won in 42 cases (~8%) and won strongly in 2 cases. The curves for the WR metric were constant in 11% of cases, and "inverted" in 0.7%; the curves for the LL metric were never constant, and "inverted" in just 2 cases (0.4%). Thus, even under conditions yielding good resolution and with no reliance on a particular threshold, the WR metric is problematic because of curve flatness. In other words, a wide range of ancestry source combinations, both correct and partially correct, often demonstrate nearly identical worst residuals. As expected, this problem occurred only for low G or I ancestry proportions of 5% or 10%, and demographic histories with 20-80% gene flow intensities were problem-free for both metrics (Data S1). In Fig. S5a-e we visualized a small fraction of results summarized in Data S1, namely those for the 15/8.5/2.5 ka graphs and 5% I ancestry and 95% G ancestry in H. The flatness problem is noticeable in iterations 2, 3, 5, 6, 9, 17, 20, 21, 23, 24, 27, 29, 30, 32, 43, 45-47, 49, and 50. The inversion problem is noticeable in iterations 12, 13, and 42.

Now let us inspect the performance of the metrics under a nearly star-like demography 10/8.5/7.5 ka. In 73% of cases both metrics performed in the same way given the thresholds; the WR metric "won" in 3% of cases, the LL metric won in 24% of cases, and won strongly in 3 cases (~0.5%) (Data S1). The curves for the WR metric were constant in 47% of cases, and "inverted" in 12%; the curves for the LL metric were constant in 4% of cases, and "inverted" in 18% (Data S1). In Fig. S5f-j we visualized results for the 10/8.5/7.5 ka graphs and 40% I ancestry and 60% G ancestry in H. The flatness problem is noticeable in iterations 2, 12, 16, 20, 28, 30, 36, 37, 38, 40, 42, 48, and 49. It is remarkable that even under nearly equal admixture proportions (40% to 60% G ancestry in H) the WR metric demonstrated poor performance because of the flatness problem (Data S1): both metrics performed in the same way in 79% of cases, the LL metric won in 19% of cases, and won strongly in 2 cases (~1.3%). The curves for the WR metric were constant in 22% of cases cf. just one such case for the LL metric (~0.7%). Inverted curves were not observed for these approximately equal admixture proportions. In conclusion, both metrics performed poorly on a star-like phylogeny, but the WR metric was much worse than the LL metric. This effect can be partially attributed to the ascertainment bias (*31*) which shifts expectations of some *f*-statistics and thus increases the background noise. For instance, a certain $f_4$-statistic may be shifted from 0 due to bias, and that results in a WR not associated with the target group and not depending on the combination of ancestry sources for the target.

*d. Detecting the direction of gene flow*

Next, we tested if the graph mapping method is capable of resolving the direction of gene flow. We focused on sets of 95,000 simulated variable sites and first compared graphs of the "incorrect 2" and "partially correct 2" classes under the "mapping" setup. These graphs differ in the direction of the G-H gene flow (G→H is the correct direction) and both have group A modelled incorrectly as getting 80% ancestry from group B. We wanted to test if finding out the direction of gene flow in the clade of interest is possible given these severely non-fitting skeleton graphs (WR >9 SE). We compared models according to the WR and LL metrics. If a gene flow in either G→H or H→G direction approaches 0%, then the topologies H = I + G ("partially correct" graphs) and G = F + H ("incorrect" graphs) approach the same topology ((F, G), (H, I)). If a gene flows in either direction approaches 100%, then the task is formulated as distinguishing between the (outgroups, (F, (I, (G, H)))), and (outgroups, (I, (F, (G, H)))) branching orders.

As anticipated, the WR metric was largely useless for finding out the direction of gene flow in this case (Fig. S6) since the strong signal reflecting the incorrect model for A masked the H-G signal under most settings, except for the setting 15/8.5 ka (Fig. S6d). Even under the most favorable conditions, namely the combination of 15 ka and 8.5 ka divergence dates and three alternative admixture dates (2.5, 5, 7.5 ka), the LL metric allowed to resolve the correct and incorrect directions in all iterations if the simulated gene flow was 20% or higher, given a significance threshold of 3 ln-units applicable for models having the same number of parameters (*11*) (Fig. S6d, Table S2). Under the least favorable conditions tested (10 ka and 8.5 ka divergence dates and three alternative admixture dates of 2.5, 5, and 7.5 ka), the LL metric allowed to resolve the correct and incorrect directions in >95% of iterations only if the simulated gene flow was 90-100% (the gene flow at 7.5) or 80-100% (the gene flow at 2.5 or 5 ka) (Fig. S6a, Table S2).

Next, we compared graphs of the "partially correct 1" and "incorrect 1" classes that have much lower WRs (see the description of the graphs above). Here the resolution was slightly better, and the performance of the LL metric was also better, especially under the 10/8.5 ka setting (Table S2). Under the most favorable conditions (15 ka and 8.5 ka divergence dates), the LL metric allowed to resolve the correct and incorrect directions in >95% of iterations, if the simulated gene flow was 20% or higher (Table S2). Under the least favorable conditions tested (10 ka and 8.5 ka divergence dates), the LL metric provided this level of resolution only if the simulated gene flow was 80-100% (the gene flow at 7.5 ka) or 50-100% (the gene flow at 2.5 or 5 ka). The performance of the method improved on the larger sets of 400,000 sites (Table S2).

Thus, given an "information-poor" demographic history where just one group (F) is differentially related to the true sources of ancestry in the target (the I and G lineages), the main problem for the mapping approach is distinguishing directions of gene flow since even the incorrect direction yields result patterns similar to the correct ones (results not shown), and as illustrated above the directions remain largely indistinguishable for topologies with closely spaced population divergence events (10 and 8.5 ka, 13 and 10.5 ka) even if admixture proportions are ~40%.

*e. Admixture proportions and absolute model fit*

We explored if simulated admixture proportions were inferred correctly under the "mapping" setup (95,000 sites, correct graphs), where they were not locked. The estimated and simulated proportions were linearly dependent, and interquartile ranges of estimated values overlapped the simulated values with no exception (Fig. S7a). The precision depended more on divergence dates rather than on admixture dates (2.5, 5, and 7.5 ka), and was substantially different for the low-resolution (10/8.5 ka) and high-resolution settings (15/8.5 ka): standard deviations (averaged across all admixture dates and simulated gene flow values of 5% to 95%) were 14.4% vs. 2.8%, respectively (Table S3). The other settings (13/10.5 ka, 13/8.5 ka) demonstrated intermediate levels of precision: average standard deviations were 9.7% and 9.2%, respectively.

We also assessed model fit to the data expressed as WR (95,000 sites, correct graphs, "mapping" setup, the correct model H = I + G). A threshold commonly used for declaring an admixture graph model as fitting the data is 3 SE intervals (*45*). Fits remained uniform across all settings, with median fits for a particular setting never exceeding 3 SE intervals (Fig. S8a). However, 26% of all iterations (2,030 of 7,800) demonstrated WRs above 3 SE. Incorrect admixture proportion estimations by *qpGraph* did not influence this result since when admixture proportions were locked at the correct values, 2066 of 7800 iterations demonstrated WRs above 3 SE. We explore the problem of assessing absolute model fits in greater detail in section j.

In the case of the larger dataset (400,000 sites), 27% of all iterations (2,076 of 7,800) demonstrated WRs above 3 SE (Fig. S8b). Precision of admixture proportion estimation has also improved with increasing number of sites (Fig. S7b, Table S3).

*f. Comparing the performance of D-statistics, qpAdm, and qpGraph on the information-poor phylogeny*

Finally, we explored the performance of popular admixture inference methods relying on allele frequencies, namely *qpAdm* (*48*) and *D*-statistics (*37, 44, 45*), and compared them to our graph mapping method.

*qpAdm* tests admixture models for a target group given a set of groups that act as proxies for ancestry sources (often termed "left" populations) and a wider set of ~10 outgroups that are differentially related to the ancestry sources (often termed "right" populations), although a phylogeny is not specified. In the "information-poor" class of graphs, there is only one group differentially related to the sources, so this demographic history is expected to be non-optimal for the *qpAdm* approach. Nevertheless, we started by applying *qpAdm* to this limiting case. The following set of outgroups was used: Outgroup, A, B, C, D, E, F (the only differentially related group). We tested the HI and HG population pairs using *qpWave* (*17*), and then tested the H = I + G model using *qpAdm*. If both HI and HG pairs were consistent with being derived from two distinct ancestry streams (i.e. *p*-values were <0.05 or alternatively <0.01), if the H = I + G model was not rejected (*p*-value for two ancestry streams >0.05 or alternatively >0.01), and if the ancestry proportions inferred by *qpAdm* remained within 0% and 100%, a positive admixture signal was recorded; otherwise a negative signal was recorded. We also used an alternative more sensitive setup with "rotating" outgroups (*49*): when testing the HI and HG population pairs using *qpWave*, groups G or I, respectively, were added to the outgroup set. Thus, we tested two *qpAdm*/*qpWave* setups and two *p*-value thresholds for each.

Individual *D*-statistics are often used for detecting admixture (see, for instance, *4, 76*), in addition to more sophisticated methods like *qpAdm*. For all the simulated datasets ("information-poor" graphs) we calculated all *D*-statistics including the groups in the clade of interest and one outgroup: *D*(I, H; G, X), *D*(I, G; H, X), and *D*(H, G; I, X), where X is any of Outgroup, A, B, C, D, E, F. Among those statistics, we chose complementary statistics: 1) *D*(I, H; G, X), which approaches 0 when the G→H flow approaches 0%; and 2) *D*(H, G; I, X), which approaches 0 when the G→H flow approaches 100%. Among those sets of statistics applied to all the simulated datasets we selected individual statistics yielding fewer non-significant deviations from 0 (absolute Z-score <3), and those were *D*(I, H; G, Outgroup) and *D*(H, G; I, F). These two statistics were used as a single classifier. The result was considered positive if *D*(I, H; G, Outgroup) deviated significantly from 0 in the negative direction (Z-score <–3 or alternatively <–2) and if *D*(H, G; I, F) deviated significantly from 0 in the positive direction (Z-score >3 or alternatively >2). To take care of the multiple testing problem, in nearly all studies an absolute Z-score of >3 is interpreted as significantly deviating from 0, however we also tested a non-standard threshold of 2 SE intervals.

We calculated MCCs for these six alternative classifiers based on *D*-statistics and *qpAdm*/*qpWave* and compared them with the most stringent classifiers based on the graph mapping approach for 95,000 or 400,000 sites: LL10 and WR0.8 (Table S4). Here we ranked the classifiers by performance (in the order of decreasing MCC averaged across all 12 setting combinations) in the case of 5% G ancestry in H and 95,000 sites: 1) *D*-statistics Z2; 2) *qpGraph* LL10; 3) *qpAdm*/*qpWave* (G or I in outgroups) p0.05; 4) *D*-statistics Z3, *qpGraph* WR08, and *qpAdm*/*qpWave* (G or I in outgroups) p0.01; 5) *qpAdm*/*qpWave* 0.05; 6) *qpAdm*/*qpWave* 0.01 (Table S4). It is clear that *D*-statistics with the Z-score threshold of 2 SE was the best-performing classifier, and *qpGraph* LL10 occupied the second place. Remarkably, even the "rotating" *qpAdm*/*qpWave* setup lagged behind in performance. The sensitivity gains conferred by the best-performing classifiers (LL10 and *D*-statistics Z2) were marginal under many high-resolution settings (Fig. S9), but substantial under low-resolution settings like 13/10.5/5 ka (10% gene flow), 10/8.5/2.5 ka (10% gene flow), and 10/8.5/7.5 ka (20% and 30% gene flow).

False positive rates were very similar for all the classifiers compared here. We also note that under low-resolution settings (e.g., 10/8.5/7.5 ka) the *p*-value threshold of 0.01 yielded substantially lower sensitivity as compared to the 0.05 threshold. It is worth noting that under the same low-resolution settings *qpGraph* demonstrated moderate sensitivity to gene flow (Fig. S9), but poor sensitivity to the direction of gene flow (Fig. S6a). However, even under the most optimal settings tested here (15/8.5/2.5 ka and gene flow intensities between 10% and 90%), for which the LL10 and WR0.8 classifiers demonstrated perfect sensitivity and specificity (Fig. S3g), *qpAdm* with a *p*-value threshold of 0.05 demonstrated false negative rates exceeding or equal to 6% for certain gene flow intensities (Fig. S9). This reflects the design of the *qpAdm* classifier since a *p*-value cutoff of 0.05 by definition rejects 5% of correct models (*49*). Indeed, the 0.05 cutoff applied to 50%/50% simulated admixture proportions yielded a false negative rate of 5.2% on average across all settings. The other classifiers do not have this disadvantage since the false negative rate always drops to 0 for an optimal demographic history. Essentially the same results were obtained for sets of 400,000 sites (Fig. S9b, Table S4), with sensitivity uniformly increasing across all classifiers.

We then looked at ancestry proportions and their standard errors estimated by *qpAdm* on sets of 95,000 sites (Fig. S10a), obtaining a result similar to that generated by *qpGraph* (Fig. S7a), but noisier (Table S3). Standard deviations of estimated ancestry proportions across iterations were substantially different for the low-resolution (10/8.5 ka) and high-resolution settings (15/8.5 ka): 28.5% vs. 3%, respectively (Table S3, standard deviations were averaged across all admixture dates and simulated gene flow values from 5% to 95%). Cf. standard deviations of 14.4% and 2.8%, respectively, for *qpGraph* estimates. This additional noise that is most noticeable under the low-resolution setting 10/8.5 ka can be probably attributed to the fact that ancestry proportions estimated by *qpAdm* are not restricted to values between 0 and 1, in contrast to *qpGraph*. With increasing number of sites, the precision of both *qpAdm* (Fig. S10b) and *qpGraph* (Fig. S7b) estimates improved (Table S3).

*g. The information-rich phylogeny*

We used the simulation procedure described in section a and generated a different topology: a simple tree composed of 11 groups (5 individuals per group) with one admixture event in the history of the target group H. The source clades diverged 15 or 20 ka and included 5 and 3 groups (Fig. S2). This graph we call "information-rich" since it incorporates 6 groups differentially related to G and I, the proxies for the sources of gene flow in H. The previous "information-poor" graph contained just one such group. For each divergence date, we tested three alternative dates for the G→H gene flow: 2.5, 5, and 7.5 ka. "Incorrect" graphs were not fitted to the information-rich datasets, just correct ones.

As expected, the performance of all classifiers on this dataset improved as compared to the information-poor dataset, and *qpGraph* LL10 and *D*-statistics with a Z-score threshold of 2 outperformed the other classifiers under challenging settings, for instance the 15 ka divergence date combined with an admixture date at 7.5 ka and a 5% admixture proportion (Fig. S11, Tables S1, S4). We note that the *D*-statistic classifier was constructed in the same complex way as above, and statistics *D*(I, H; G, C) and *D*(H, G; I, F) were used as a composite classifier. The *qpAdm* classifiers were also constructed in the same way as the "rotating"*qpAdm* classifiers above. The average *qpAdm* false-negative rate for all combinations of admixture/divergence dates and for well-resolved admixture proportions between 10% and 90% was expected: 5.1% for the *p*-value threshold of 0.05, and 1% for the *p*-value threshold of 0.01. These false negative rates are much higher than 0% false negative rates attained by the other methods.

The fraction of correct admixture graphs that demonstrated a worst residual with an absolute Z-score > 3 increased as compared to the information-poor dataset, 38% vs. 25-26%.

*h. Resolving alternative ancestry sources on the information-rich phylogeny*

The main purpose of simulating the information-rich demographic history was testing an aspect that remained untested on the information-poor phylogeny: the power to resolve closely related alternative ancestry sources. That is an important question often arising in studies of genetic history. While *qpGraph* provides a natural approach for this via likelihood ranking of models with ancestry sources on various edges of the graph, alternative non-nested *qpAdm* models cannot be easily ranked. They can be just subdivided into two classes: rejected and not rejected.

In Fig. S12 we illustrated the fraction of iterations that showed a *positive* admixture signal for the correct pair G+I and for 12 source pairs including just one correct source (G or I). The results are shown for various admixture proportions and divergence/admixture dates, and for two *qpWave*/*qpAdm*-based classifiers ("rotating outgroup" classifiers with the 0.01 and 0.05 *p*-value thresholds). In Fig. S13 we illustrated the fraction of *qpGraph* iterations where a partially correct source pair was *not* rejected according to an LL difference threshold of 3 units or a WR difference threshold of 0.1 SE. *qpAdm* failed to resolve the F- and G-related ancestry sources (that diverged 8.5 ka ago) across a wide range of admixture proportions, especially if the *p*-value threshold of 0.01 was used. Resolving the I- and C-related sources that diverged 11.5 ka ago was also compromised for admixture proportions up to 20-30%. In contrast, *qpGraph* allowed to resolve the alternative sources across a much wider range of admixture proportions, especially if LL and not WR was used as a ranking metric.

Here we also show admixture proportions for the G+I, F+I, and G+C source pairs estimated by *qpAdm* and *qpGraph* (Fig. S14). Those estimated by *qpGraph* for incorrect sources F or C just slightly deviated from the simulated ones, but those estimated by *qpAdm* deviated considerably for simulated admixture proportions >40%. We also showed *qpAdm* *p*-values for the G+I, F+I, and G+C source pairs (Fig. S15) and worst residuals reported by *qpGraph* for these source pairs (Fig. S16).

The *qpWave*/*qpAdm* classifier with a *p*-value threshold of 0.01 demonstrated poor sensitivity in the case of low admixture proportions (Fig. S11), and it turned out to be completely unsuitable for resolving alternative ancestry sources that diverged 8.5 ka ago (Fig. S12). The classifier with a *p*-value threshold of 0.05 also demonstrated a rather poor resolution of alternative sources. These results suggest that *qpAdm*-based classifiers perform poorly even in nearly perfect conditions: when many outgroups differentially related to the sources are available, there is no missing data, no pseudohaploid data, the true ancestry sources are deeply diverging, and there is no multilayered admixture in the target. *qpAdm* is nowadays often used as a model-ranking tool, especially using the "rotating" setup tested here when unused alternative sources are moved into the outgroup set (*30, 49, 51*). We caution that the *qpWave*/*qpAdm* method should be used for the purpose of model rejection, but not for model ranking, and suggest using *qpGraph* instead if possible, i.e. if the number of admixture "layers" in the history is amenable to *qpGraph* model testing.

*i. Phylogenetic accuracy of mapping when a gene flow is not accounted for*
Another problem unexplored so far is what happens when an admixed group is mapped on a skeleton graph as unadmixed, namely what position on the graph "wins" in these cases. We considered one-way models having the lowest LL and classified them according to ancestry sources: on the I branch, on the G branch, or on any intermediate branch (Table S5). The emerging pattern is rather complex and is influenced by tree topology and divergence and admixture dates. The interval of admixture proportions where intermediate positions for the target group win became wider with increasing divergence between the two ancestry sources and with increasing gene flow date. It was also wider for the information-rich vs. the information poor graphs. In nearly all cases this interval lay within an interval of proportions confidently detectable by the LL10 classifier (Table S1). Thus, for the graphs tested here a phylogenetically incorrect position can always be rejected when two-way models are tested and interpreted using the LL10 or similar classifier.

*j. On measuring absolute model fit to the data*
Above we attempted to find optimal approaches for ranking alternative models. But ranking alternative models and assessing if a given model fits the data well are different questions. Traditionally the worst *f*-statistic residual was used as a measure of absolute fit to the data (*45*), and 3 SE was used as a threshold value. We have shown above that this threshold value is overly stringent (Fig. S8) in the presence of ascertainment bias that was introduced to mimic a typical SNP panel (*31*) and that is expected to affect expectations of some *f*-statistics. While residuals of a single *f*-statistic are expected to be distributed normally across simulation iterations, that is not the case for worst residuals. Below we compare three alternative methods for assessing the absolute model fit: the traditional worst *f*-statistic residuals and two methods that test how close is the distribution of $f_2$-statistic residuals or all *f*-statistic residuals to a normal distribution with a mean of 0 and variance of 1, i.e. the standard normal distribution.

The following alternative methods were applied to quantile-quantile plots: 1) Lin's concordance correlation coefficient (Lin 1989, 2000) implemented in the epiR v.1.0-14 R package and applied to quantile-quantile plots generated by the qqnorm function (the stats v.3.6.2 base R package) under the default settings; 2) percentage of outliers beyond the 95% confidence interval around the standard normal distribution. For the latter approach the qqbounds function from the distr v.2.8.0 R package was used. Symmetric simultaneous confidence intervals were determined relying on the exact Kolmogorov distribution. The following options were passed to the epi.ccc function: ci = "z-transform", conf.level = 0.95. The following options were passed to the qqbounds function: D = Norm(0,1), alpha = 0.95, n = [number of *f*-statistics], withConf.pw = FALSE, withConf.sim = TRUE, exact.sCI = TRUE, exact.pCI = FALSE, nosym.pCI = FALSE.

First, we mapped the target as an unadmixed lineage on the information-rich graphs. Alternative branch points for the target group H were divided into three classes: correct (on the I and/or G branches, depending on the simulated admixture proportions); "correct clade" (those within the C+I clade including its stem and/or within the G+F clade including its stem, depending on the simulated admixture proportions); and incorrect (all the other branch points) (see Fig. S2). The C and I groups diverged 11.5 kya, and the G and F groups diverged 8.5 kya. In the case of the information-poor graphs, all sources within the F+G+I clade and its stem were included into the "correct clade" group, except for the G and/or I branches (Fig. S1). The F/G and I groups diverged 10, 13, or 15 kya. Although the classification of sources was not consistent across different topologies (information-rich and information-poor), we compared the performance of alternative approaches for each topology independently.

In the case of the information-rich graph and 95,000 sites, 99% of correct graphs yielded worst *f*-statistic residuals below 4.13 SE (Fig. S17a). Closely related ancestry sources ("correct clade") and distantly related "incorrect" sources yielded WRs above 4.13 SE under all settings tested (Fig. S17a). The same level of resolution was attained for the information-rich graph and 400,000 sites (results not shown). 99% of correct graphs yielded worst $f_4$-statistic residuals below 4.07 SE. Thus, for this simple demographic history it was possible to find a WR threshold that distinguished correct sources and all the other sources, and thus can be used as a threshold for absolute model fit. Notably, this threshold was higher than 3 SE. Both methods that compared the distribution of $f_2$-statistic residuals to the standard normal distribution allowed to distinguish the "correct" and "incorrect" source classes, but not the "correct" and "correct clade" classes (Fig. S17b,c).

We checked if these results hold for the information-poor graphs. Setting the WR cut-off at ~4 SE (the 99% quantile for the correct unadmixed models, 95,000 sites) or at ~4.2 SE (the 99% quantile for the correct unadmixed models, 400,000 sites) allowed to reject all "correct clade" and incorrect models for 0% and 100% simulated admixture proportions. The methods that check if the distribution of $f_2$-statistic residuals conforms to the standard normal distribution allowed to distinguish correct from incorrect sources, but under many settings they did not allow to distinguish correct and "correct clade" sources (results not shown). $f_2$-statistic residuals of incorrect information-poor models conformed to the general normal distribution even more often than those of the correct or "correct clade" models, which makes this approach unacceptable.

Second, we considered graphs with the target mapped as a 2-way mixture on the information-rich graphs. Alternative source pairs for the target group H were divided into five classes: correct (both sources on the I and/or G branches, depending on the simulated admixture proportions); "correct partially" (a correct + incorrect pair); "correct clade" (a correct + "correct clade" pair or "correct clade" + "correct clade" pair); "correct clade partially" (an incorrect + "correct clade" pair); and incorrect (all the other source pairs) (see Fig. S2). Setting the WR cut-off at ~4.1 SE (the 99% quantile for the correct admixed models, 95,000 sites, Fig. S18a) allowed to reject all incorrect source pairs and nearly all pairs classified as "correct clade partially", across the whole range of simulated admixture proportions. However, it was impossible to distinguish the correct and "correct clade" classes under many settings and admixture proportions (Fig. S18a). The same level of resolution was attained for the information-rich graph and 400,000 sites (results not shown). 99% of correct graphs yielded worst *f*-statistic residuals below 4.14 SE. The methods that check if the distribution of $f_2$-statistic residuals conforms to the standard normal distribution allowed to distinguish most extreme cases, i.e. correct and incorrect source pairs, but under many settings they did not allow to distinguish the other three classes from the correct one (Fig. S18b,c).

We again checked if these results hold for the information-poor graphs. Setting the WR cut-off at ~4 SE (the 99% quantile for the correct admixed models, 95,000 sites) or at 4.05 SE (the 99% quantile for the correct admixed models, 400,000 sites) allowed to reject all incorrect source pairs and all pairs classified as "correct clade partially" (except for the least favorable combination of divergence and admixture dates, 10/8.5/7.5 ka), across the whole range of simulated admixture proportions. However, it was impossible to distinguish the correct and "correct clade" classes as well as correct and "correct partially" classes under many settings and admixture proportions (results not shown).

Finally, we repeated these analyses for the information-rich graph (95,000 sites) and unadmixed models, but now considering all $f_2$-, $f_3$-, and $f_4$-statistic residuals instead of $f_2$-statistic residuals. A method (concordance correlation coefficient) that checks if the distribution of residuals conforms to the standard normal distribution now performed much better and allowed to distinguish correct and "correct clade" sources under all settings except for 15/7.5 ka and 20/7.5 ka (Fig. S19). In that case it was impossible to distinguish only the G and F sources that diverged 8.5 ka. We point out that WR allowed to distinguish correct and "correct clade" sources even under these settings (Fig. S17a). Due to high computational requirements, for the information-rich graph (95,000 sites) and 2-way admixed models we calculated only concordance correlation coefficient for all residuals and for the most challenging setting only: 15/7.5 ka (Fig. S20). Again, WR outperformed concordance correlation coefficient for all *f*-statistic residuals (Fig. S19).

*k. Conclusions*

We attempted further fine-tuning of the *qpGraph*-based classifiers, testing LL difference values between 3 and 12 units in increments of 1 and WR difference values between 0.1 and 1 in increments of 0.1 (Table S6). For each graph/dataset combination we calculated false positive rates across all settings and G ancestry proportions (0% or 100%). The LL10 and WR0.8 classifiers demonstrated nearly identical false positive rates of 1.6% and 1.5%, respectively (averaged across all datasets), thus comparing their performance was justified. The following threshold values resulted in false positive rates below 5% for all four graph/dataset combinations tested in this section: 8 ln-units for LL and 0.7 SE intervals for WR.

Here we focused on one question: demonstrating the existence of a gene flow. But what if the question is different: demonstrating the absence of a gene flow. That question is relevant when constructing skeleton graphs for subsequent mapping. In the case of the information-rich graph, the LL threshold of 8 units can be used for tackling both problems: 95% LL difference percentiles for datasets without simulated gene flow were below 8, and 5% LL difference percentiles for datasets with simulated admixture proportions ranging from 5% to 95% were invariably above 8 (Table S7). However, in the case of the information-poor graph, there is a wide gray zone where 5% percentiles for datasets with simulated admixture were below 8 (Table S7). This gray zone includes 5, 10, and 80-95% admixture proportions. Information-poor sub-graphs often occur within larger graphs: for instance, a small apical clade with many outgroups. Demonstrating the absence of admixture between members of such apical clades that diverged ca. 10 ka is inherently difficult, and that should be kept in mind when analyzing admixture graphs.

We conclude that in the case of simple simulated demographic histories without multi-layered admixture, among other methods based on *f*-statistics, *D*-statistics with a significance threshold of 2 SE intervals is the most sensitive tool for detection of subtle gene flow signals, and *qpGraph* occupies the second place, especially if LL and not WR is used for model ranking. Simpler methods are more sensitive partly because they test more focused hypotheses. Specificity of all methods is very similar and high if appropriate thresholds are used, however finding optimal thresholds is not straightforward.

We caution that not all outgroups ensure equal performance of *D*-statistics, and thus on real data various statistics should be analyzed in parallel. A method based on *qpWave*/*qpAdm* (the sensitive "rotating" setup when unused alternative ancestry sources are moved into the outgroup set) lagged behind in sensitivity to gene flow and demonstrated non-zero false negative rates in ideal conditions that are inevitable by design. Moreover, *qpAdm* failed to resolve alternative ancestry sources that separated 8.5 ka ago, while ranking admixture graphs by log-likelihood allowed to resolve them. Since *qpGraph*, unlike *qpAdm*, provides a natural way of ranking alternative ancestry sources and a way to avoid the overfitting problem (Supplementary Text 11), we believe that *qpGraph* should be preferred for inferring admixture if it is possible to construct a graph without an overly complex network of gene flows.

2. Testing PCA, *D*-statistics, *qpAdm*, and *qpGraph* on a complex simulated history

Even though PCA, *f*-statistics and derived methods are central to archaeogenetic studies (see, for example, *28, 30, 48, 50-66*), just few studies were devoted to testing their performance in fully controlled conditions, i.e. on simulated genetic data (*13, 37, 45, 49, 51, 77*). For instance, Moreno-Mayar *et al.* (*13*) devoted a small section of their paper to simulations that demonstrated that *D*-statistics are insensitive even to 40% admixture proportions under certain demographic scenarios. Other studies were devoted to exploring PCA and related methods on simulated data. Novembre and Stephens (*81*) explored on simulated data patterns observed when PC values are plotted on geographic maps, as well as $PC_n$ vs. $PC_m$ biplots. The authors concluded that "interpreting gradient- and wave-like patterns in PC maps as signatures of historical migration events is problematic because such patterns arise generally under a simple condition: that genetic similarity decays with distance." PC1 vs. PC2 biplots were also shown to display a "horseshoe" artefact under the homogeneous migration scenario. The same artefact was later explored by Frichot *et al.* (*78*) and observed by Harney *et al.* (*49*) under the continuous migration scenario.

McVean (*79*) concluded that "PCA predominantly reflects structure in the expected coalescent time. Consequently, any two demographic models that give the same structure of expected coalescence times will also give the same projections." The author also demonstrated that imbalanced sampling across groups has a major effect on low-dimensional embeddings. Reich *et al.* (*80*) agreed that in the light of new results by Novembre & Stephens (*81*) "if the aim is to study history and document migrations, it is important to carry out additional research to correlate the PCA results with other lines of evidence".

Formal tests for genetic admixture ("admixture" $f_3$-statistics, $D/f_4$-statistics, and *qpAdm*) are often used exactly for this purpose: to prove that a certain cline spotted in a PC space is a result of a migration and admixture of previously isolated groups, and does not reflect isolation by distance or recurrent bottlenecks. Recently *qpAdm* is increasingly used as a main test for admixture hypotheses put forward after inspecting PCA plots (see the long list of studies cited above). The only study reporting detailed testing of *qpAdm* on simulated data is a recent pre-print by Harney *et al.* (*49*). The method was explored by the authors under extremely favourable conditions: the simulated data was free of ascertainment bias since sites were sampled in a random way, the simulated graph included just two non-nested admixture events, the sources for the principal target group diverged about 1200 generation ago (almost 35000 years ago); the source proxies were strictly cladal with the real ancestral groups; several groups differentially related to the ancestry sources were available, 1 million sites was used for most analyses; and only 50/50% simulated admixture proportions were tested for some analyses. The study confirmed that the method behaves as expected under these "laboratory" conditions and offered some important guidelines on the choice of outgroups and model comparison strategies, and also showed that the results are robust to the presence of missing data, imbalanced group sizes, ancient DNA damage, and a particular type of SNP ascertainment: selecting sites heterozygous in one individual from a certain population (*45*). Crucially, the false positive rate of the method remained virtually unexplored. Among challenging scenarios, only multifurcation with subsequent continuing gene flow among several groups was explored, and it was concluded that *qpAdm* is not applicable in this case. Thus, the method was proven to work and fail under polar conditions. But what happens in intermediate cases where arguably most of our species history fits: a history that cannot be represented as a nearly perfect tree, but on the other hand is not characterized by gene flows homogeneous in space and constant in time?

*a. Simulated demographic history*

We decided to test the core suite of archaeogenetic methods (PCA, *D*-statistics, *qpAdm*, *qpGraph*) on a relatively complex simulated demographic history with multi-layered pulse-like admixture events. In the previous section we simulated two classes of simple demographic histories without multi-layered admixture. We showed that all methods based on *f*-statistics behave as expected under these favourable conditions, and if the phylogeny is not approaching a star-like one, they achieve satisfactory false negative (FN) and false positive (FP) rates for detection of a gene flow. Thus, we demonstrated that the software versions tested below are free of major mistakes.

For preparing the complex simulations we followed the protocol described in Supplementary Text 1, section a. The complex history was designed to resemble the skeleton admixture graph used in this study on real data (Fig. S3-1). To assess the effect of the ascertainment bias introduced by the global MAF filter, we prepared two versions of each dataset: generated using a 5% MAF filter followed by random subsampling to 95,000 or 400,000 sites, or generated using random subsampling only. Most results below are based on the larger sets of 400,000 sites with the ascertainment bias, and some crucial analyses were repeated for sets of 400,000 sites without the ascertainment bias. In total, 750 whole-genome datasets were generated, and the four alternative site sampling schemes produced 3,000 final datasets.

The number of groups and the numbers of individuals per group were identical to those included in the skeleton graph on real data plus USR1 (15 groups and 54 individuals). The dates for leaf nodes, divergence and gene flow dates (Fig. S21a, Table S8) were set according to the radiocarbon dates and to our limited knowledge of the history of relevant groups. The same was true for effective population sizes. For the sake of simplicity, data for all simulated individuals was diploid, and no missing data was introduced. We recognize that this is a limitation which makes the simulated and real datasets different. Below all simulated group names are underlined, and real group names are not. We simulated the <u>USR1</u> individual as a mixture composed of 30% of a <u>Mal'ta</u>-related source and 70% of source X. X was simulated as a mixture of the Asian source of First Americans ($FAM_{EA}$) and a <u>Siberian</u> (<u>Houtaomuga</u>-related) group. The simulated source groups diverged 7 kya prior to the admixture event. We tested the following proportions of <u>Houtaomuga</u>-related ancestry in <u>USR1</u>: 3.5%, 7%, 14%, and 35%. This range of simulated proportions completely covers the proportions fitting the real genetic data (see Supplementary Text 7g). We tested simultaneously different levels of the <u>Houtaomuga</u>=><u>USR1</u> gene flow and different ratios of <u>Mal'ta</u> ancestry in <u>FAM</u> and <u>USR1:</u> 30/30%, 32/30%, and 40/30%. The <u>Mal'ta</u>-related gene flows into the <u>FAM</u> and <u>USR1</u> branches followed the <u>$FAM_{EA}$/Houtaomuga</u> admixture event (Fig. S21a, Table S8).

*b. Principal component analysis*

We tested various setups on both real data and their simulated counterparts: 10 PCs based on both ancient and present-day individuals were calculated using *PLINK* v.1.90b6.10 or *smartPCA* v.16000, or alternatively certain individuals were projected on those axes using *smartPCA*. Unless noted otherwise, for inferring the PCs we took only real individuals that had simulated counterparts. The pairwise Wright's $F_{ST}$ distances between simulated groups lay within the range typical for humans (results not shown).

The PC1/PC2 biplot constructed on simulated diploid data with ascertainment bias resembles the plot for Eurasians and Americans on real diploid and pseudohaploid data from the

1240K SNP panel (Fig. S22a-c). This result suggests that the simulation has reproduced key aspects of the actual history. Importantly, the simulated USR1 individual occupies the position between FAM and Siberians even in the absence of any simulated Siberian gene flow. USR1 is also shifted from the FAM cluster towards Mal'ta along PC4 on both simulated and real data (Data S2). Here all individuals plotted were used for inferring the PCs, but the same result was observed when USR1 was projected onto the PCs (results not shown). Indeed, according to McVean (*79*), a third population that diverged from S1 since the split with population S2 is expected to be projected between S1 and S2. Another approach used e.g. by Reich *et al.* (*80*) is projecting both the main ancestry source proxy and the target onto PCs formed by reference groups. That "cancels out" any drift shared by the main source and the target. Even when 35% gene flow was simulated, the shift of USR1 from the FAM cluster was small, and it did not follow the direction of the minor source (Fig. S22d,e). The same result was observed on real data (Fig. S22f).

*c. D-statistics*
For generating subsequent results, we used *qpDstat* v. 751, *qpAdm* v. 810 and *qpGraph* v. 6412. Default settings were used for *qpDstat* and *qpAdm*. Moreno-Mayar *et al.* (*4*) used *D*-statistics *D*(a FAM group, USR1; a Siberian group, another Siberian group) and *D*(a FAM group, USR1; a Siberian group, Han) for investigating a potential gene flow between Siberians and USR1. We investigated similar statistics on simulated data. A statistic *D*(FAM, USR1; X, Siberian) is expected to deviate significantly from 0 (in the positive direction) in the presence of either the Siberian-USR1 or the X-FAM gene flow.

We tested simultaneously different levels of the Houtaomuga=>USR1 gene flow and different ratios of Mal'ta ancestry in FAM and USR1 (Fig. S21). If group X diverged outside of the Asian clade, i.e. when the Mal'ta gene flow was derived from the X branch, a small imbalance in the proportions of Mal'ta ancestry between USR1 and FAM (30 vs. 32%) drove the statistics in the positive direction in the absence of any gene flow from the Houtaomuga branch (Fig. S23b). A larger difference in Mal'ta ancestry (30 vs. 40%) made nearly all statistics *D*(FAM, USR1; X, Houtaomuga) significantly positive across all iterations (Fig. S23c).

These results are not unexpected, and it is known that *D*- or $f_4$-statistics cannot be interpreted unambiguously. Statistics of the type *D*(Reference, Target; Source, Outgroup) are often used to test for gene flows, and very often distant outgroups (e.g., Africans) are used. However, our results show that for the test to be interpretable unambiguously the ancestry components need to be perfectly balanced in the reference and target, which is hard to control in high-throughput analyses. Moreover, there is a higher chance of encountering gene flows into the reference group if genetically distant outgroups are used since many lineages that could contribute gene flows could diverge on the outgroup branch.

*d. qpAdm*
Harney *et al.* (*49*), Narasimhan *et al.* (*30*), and Lazaridis *et al.* (*51*) recommended that "rotating" outgroup/source sets should be used for *qpAdm* analyses. A "rotating" set means that when performing tests all unused alternative ancestry sources for the target are placed into the outgroup set. This approach dramatically increases the sensitivity of the method and improves the resolution among alternative sources, as demonstrated by Harney *et al.* (*49*) and by our analyses on simple graphs (Supplementary Text 1). Informed by the *D*-statistic results above, we assembled three alternative outgroup sets (Fig. S21b): 1) a rotating set including all possible

groups on the graph (11-13 groups); 2) a rotating set including only Asian groups (7-8 groups); 3) a fixed set of 11 groups including all groups except for USR1, FAM, Houtaomuga, & Mal'ta (fixed sets were used in earlier studies, e.g. *28, 48, 54*). Following advice by Harney *et al.* (*49*), we not only tested the expected admixture model, but also explored if simpler models are rejected. The FP and FN rates discussed below refer to the outcomes of this procedure, and not to standalone *qpAdm* tests. The procedure we used singled out only one target group (USR1) and is outlined below:

- test models T=S1, T=S2, T=S3, S1=S2, S1=S3, & S2=S3 using *qpWave*;
- if all are rejected (i.e. *p*-value < threshold), test a model T=S1+S2 using *qpAdm*;
- if it is not rejected (i.e. *p*-value > threshold) AND the inferred admixture proportions ± 2SE lie between 0 and 1 => a positive result;
- test models T=S2+S3 and T=S1+S3 too;
- if all those 2-way models are rejected, test T=S1+S2+S3. If it is not rejected and the inferred admixture proportions ± 2SE lie between 0 and 1 => a positive result;
- FP = ($FP_{\text{2-way models}}$ + $FP_{\text{3-way models}}$)/number of simulation iterations under the null hypothesis.

The condition on simulated admixture proportions (inferred admixture proportions ± 2SE should lie between 0 and 1) followed the protocol by Narasimhan *et al.* (*30*). Other studies (e.g., *28, 54*) used a less stringent condition requiring that the inferred proportions should lie between 0 and 1.

The *qpAdm* protocol focused on a single target has an assumption (*28, 30, 48, 57*) that the source proxies should receive no further gene flow after the split with true ancestry sources, in other words, the true and proxy sources should be cladal. Another assumption is that there should be no gene flow from sources/target to an outgroup after the target admixture event (*48*). These assumptions can be reduced to one requirement: $f_4$(outgroup$_i$, outgroup$_j$; real source, proxy source) = 0 for all sources and pairs of outgroups. However, in high-throughput tests involving dozens of alternative sources (*30, 54*, and other recent studies) it is hard to ensure that these assumptions are always valid. For instance, Narasimhan *et al.* (*30*) used as proxy sources present-day groups from East Asia and South Asia, regions where ancient DNA data was not available. We investigated *qpAdm* FP and FN rates when the cladality assumption is violated (see Fig. S24 for a graphical representation of the assumptions).

Outcomes of the *qpWave/qpAdm* protocol focused on a single target are shown in Fig. S25 in the FP vs. FN coordinates for four different Houtaomuga ancestry proportions in USR1 and for three different ratios of Mal'ta ancestry in the main source (FAM) and in the target (USR1). Five alternative *p*-value cut-offs were tested: 0.05, 0.01, 0.005, 0.001, 0.0001. Despite using a rather stringent procedure, we often encountered FP rates between 20 and 30% when using *p*-value thresholds most commonly employed in published *qpAdm* analyses (0.01 and 0.05). These high FP rates were encountered only when the cladality assumption (Fig. S24a) was violated, i.e. when FAM received even slightly more Mal'ta gene flow after the divergence from the real source than USR1 did (Fig. S26).

Opposite trends in FP rates were encountered when decreasing the *p*-value cut-off for the "32/30%" and "40/30%" cases (helped in the former case but made things worse in the latter) (Fig. S26). In the case of the mild violation ("32/30%"), false positives were caused by a large proportions of non-fitting *qpWave* models T=S1, e.g. 50% at a cut-off of 0.05 (Fig. S27). We note that this outcome is due to an incorrect interpretation of the *qpWave* result. A non-fitting *qpWave* model T=S1 means that either T or S1 received a gene flow from the outgroups, but here we interpreted the result as a gene flow into T and moved on to testing more complex models for T.

Most of the more complex models T=S1+S2 were fitting (Fig. S27), many of them also demonstrating admixture proportions confidently between 0 and 1 (Fig. S28). In this case decreasing the *p*-value cut-off helped to eliminate the non-fitting models T=S1, and thus drove the FP rate down. In the case of the severe cladality violation ("40/30%"), *qpWave* models T=S1 were uniformly non-fitting (Fig. S27), and the distribution of *p*-values for the 2-way models was highly skewed towards 0 (Fig. S27) All the 2-way models demonstrated admixture proportions confidently between 0 and 1 (Fig. S28). In this case decreasing the *p*-value cut-off increases the number of accepted 2-way models, and this drives the FP rate up. No FP cases were caused by fitting 3-way models USR1 = FAM + Houtaomuga + Mal'ta. Estimated Houtaomuga ancestry proportions across FP iterations were on average 6% and 18% under the "32/30%" and "40/30%" settings, respectively.

Certain outgroup sets also showed opposite effects. When the cladality violation was mild, removing the violating outgroup (the scenario in Fig. S24b) helped, i.e. the "Asian" outgroup set ("rotating set 2") demonstrated lower FP rates (Fig. S26). When the cladality violation was severe, removing the violating outgroup lead to a substantial increase in FP rates (Fig. S26). This behaviour makes method optimization very difficult. An important take-away message here is that a small-scale deviation from cladality of real and proxy sources is more dangerous than a large-scale deviation. In the latter case the method just rejects most 2-way models at *p*=0.05 (but decreasing the *p*-value cut-off is not advisable). However, if the gene flow into the source proxy is small, *qpAdm* applied to a wide collection of alternative sources would likely find a spurious source that is relatively similar to the target. We note that the cladality violation can be revealed when alternative source proxies are also tested as targets, and thus the arguments above are fully applicable to "asymmetric" *qpAdm* protocols that make a distinction between targets and source proxies. In Materials and Methods we propose a protocol that treats all non-outgroup populations equally.

We also note that the ascertainment bias introduced by a global MAF filter skewed *qpWave/qpAdm* *p*-value distributions towards 0 (Fig. S27), and that contributed to the high FP rates observed in our tests. For the simplest model T=S1 the expected false rejection rate was 5% (*p*-value cut-off = 0.05), and the observed rate was 4% in the absence of bias, and 16% in the presence of bias. For the 2-way model T=S1+S2 the observed rejection rate was 4% in the absence of bias and 14% in the presence of bias (Fig. S27). Ascertainment bias of this type, i.e. depletion of rare variants by an explicit or implicit global MAF filter, was not explored before for *f*-statistic-based methods (*45, 49*), and was observed for $f_4$-statistics calculated on popular SNP panels (*31*). Thus, we believe that our simulation results in the presence of bias are relevant for all work on SNP panels in this study and elsewhere.

*e. qpGraph*

Similar effects were observed when fitting the simulated graph to the datasets using *qpGraph*. We emulated the mapping routine used on real data (see Supplementary Text 7b,c). USR1 was mapped on the skeleton graph in the following way. Only one two-way model was tested: $\text{FAM}_{\text{EA}}$ + Mal'ta. This model is consistent with the results by Moreno-Mayar *et al.* (*4*) and was used as a baseline. We tested 25 three-way models $\text{FAM}_{\text{EA}}$ + X + Mal'ta, where X was any possible edge in the anatomically modern human (AMH) sub-graph prior to West Eurasian (Mal'ta, WHG, basal Eurasian) gene flows into Siberians and Americans. Finally, we tested 300 four-way models $FAM_{EA}$ + X +Y + Mal'ta, where X and Y were any possible edges in the anatomically modern human (AMH) sub-graph prior to West Eurasian (Mal'ta, WHG, basal Eurasian) gene flows into Siberians and Americans. The mapping protocol was repeated for four datasets (95,000 and 400,000 sites, with or without the ascertainment bias), 50 simulation iterations, and three setting combinations: 1) all admixture proportions were locked according to the simulated values (Fig. S21a, Table S8); 2) the West Eurasian gene flows into Siberians were locked according to the simulated values and the Mal'ta gene flow into FAM was locked at 32% for all simulation settings; 3) the Eurasian gene flows into Siberians and the Mal'ta gene flow into FAM were unlocked. The Neanderthal gene flows were always locked. The results of this mapping run are summarized in Table S9.

The protocol based on admixture graphs generally resulted in much lower FP and FN rates than *qpAdm* (Fig. S29), if only two expected admixture models were compared (USR1=$FAM_{EA}$ + Mal'ta and USR1=$FAM_{EA}$ + Mal'ta + Houtaomuga) and if an LL difference threshold was used (Table S9). This result is not surprising since knowing the simulated history and fitting it to the data is a huge advantage. When testing all possible ancestry sources for the target (USR1=$FAM_{EA}$ + Mal'ta + X) the distributions of LL and WR difference were much wider on datasets without any Siberian gene flow into USR1 (Fig. S29, Table S9). All source pairs/triplets were not tested using the *qpAdm* protocol described in section d above, so a direct comparison of method performance is impossible in the latter case. Importantly, if the main ancestry source for the target was mismodelled substantially (Mal'ta ancestry in FAM simulated at 40%, but locked at 32% when fitting the graphs), the spurious Houtaomuga gene flow into USR1 was supported by huge likelihood improvements (Table S9). Thus, not only *qpAdm*, but also admixture graph models tend to be problematic if main source proxies are mismodelled.

The same pattern was observed when comparing 3- and 4-way models (Table S9). If Mal'ta ancestry in FAM was simulated at 40%, but locked at 32% when fitting the graphs, models USR1 = $FAM_{EA}$ + Mal'ta + Houtaomuga + Y were supported by substantial likelihood improvements ($95^{th}$ percentiles above 16 units, 400,000 sites) irrespective of the proportion of Houtaomuga ancestry simulated in USR1.

The $95^{th}$ percentile of WR difference did not exceed 0.25 SE for all but 1 of 96 combinations of settings and "null" datasets with the ascertainment bias, even when the main ancestry source proxy (FAM) was severely mismodelled (Table S9). Here the following model comparisons were considered: $FAM_{EA}$ + Mal'ta vs. $FAM_{EA}$ + Mal'ta + X, and $FAM_{EA}$ + Mal'ta + X vs. $FAM_{EA}$ + Mal'ta + X + Y. On the other hand, the $95^{th}$ percentile of LL difference reached 14.4 ln-units when the FAM group was not mismodelled (the "unlocked" settings, 400,000 sites, ascertainment bias) and 45 units when it was mismodelled severely (the "locked32" settings, 400,000 sites, ascertainment bias). The same statistics for the smaller sets of 95,000 sites were 12 and 27 units, respectively. The $95^{th}$ percentile of LL difference did not exceed 13 ln-units for all but 1 out of 72 combinations of settings and "null" datasets with the ascertainment bias, when the main ancestry source proxy (FAM) was *not* mismodelled (Table S9). In practice it is often possible to discard skeleton graphs containing severely mismodelled groups using a WR cut-off (Fig. S30).

In the absence of ascertainment bias, the $95^{th}$ percentile of WR difference varied from 0.11 to 0.67 SE across 16 combinations of settings and "null" datasets (Table S9). Here the following model comparisons were considered: $FAM_{EA}$ + Mal'ta vs. $FAM_{EA}$ + Mal'ta + X. The $95^{th}$ percentile of LL difference reached 13 ln-units when the FAM group was not mismodelled (the "locked" settings, 400,000 sites, no ascertainment bias) and 30 units when it was mismodelled severely (the "locked32" settings, 400,000 sites, no ascertainment bias). The same statistics for the smaller sets of 95,000 sites were 10 and 19 units, respectively. The 95th percentile of LL difference did not exceed 13 ln-units for all 12 combinations of settings and "null" datasets without the ascertainment bias, when the main ancestry source proxy (FAM) was *not* mismodelled (Table S9).

Considering these results on simulated data we propose the following combination of cut-offs. A Siberian gene flow into USR1 will be considered supported by the data if it results in a **WR difference >0.25 SE or an LL difference >13 units** (for both dataset sizes, see Table S9). If all ancestry sources tested show a flat WR profile, i.e. if the most problematic residual is associated with another part of the graph, only the LL metric will be considered. The LL cut-off allows to discard spurious sources in at least 95% of cases for both 2- and 3-way and 3- and 4-way model comparisons in both in the presence and in the absence of ascertainment bias, under an assumption that the main ancestry source is not mismodelled. On simulated data the chosen WR and LL thresholds allow only a high level of Houtaomuga ancestry in USR1 (35% but not 14%) to be detected confidently (for >95% simulation iterations, see Table S9), but allow to keep the FP rate below 5% under all settings tested. We caution that these cut-offs are optimized for a particular dataset size, graph topology and complexity (number of groups and number of model parameters) and are not necessarily optimal for any given dataset and graph.

*f. Measuring absolute fit of admixture graph models*

In Supplementary Text 1j we compared several alternative metrics for assessing absolute model fit: WR and measures of normality for distributions of $f_2$-residuals or all $f$-statistic residuals. We concluded that WR is the most sensitive of those metrics, i.e. allows to distinguish between correct and incorrect ancestry sources better than the other metrics. The second-best metric was a measure of normality for distributions of all $f$-statistic residuals (CCC, concordance correlation coefficient).

Here we explored admixture graph models fitted to the complex simulated histories and visualized the distributions of WR (Fig. S30), CCC and the percentage of all $f$-statistic residuals that lie outside the 95% confidence interval around the standard normal distribution (Fig. S32). We confirmed that WR is slightly more sensitive as compared to CCC on all $f$-statistic residuals, and concluded that CCC on $f_2$-residuals and the percentage of all $f$-statistic residuals that lie outside the 95% confidence interval around the standard normal distribution are not useful on these datasets since they do not allow distinguishing correct and incorrect models for USR1 (Fig. S32). We observe also that WR for correct models fitted to the data with ascertainment bias was in most cases higher than 3 SE (Fig. S30), a threshold value often used for admixture graphs in the literature: 83% and 98% of "unlocked" models on 95,000 and 400,000 sites, respectively. The corresponding 99th WR percentiles were 5.1 and 6.3 SE, respectively. Thus, WR for perfect models on the sets of 95,000 sites lay roughly between 2.5 and 5, and those on the sets of 400,000 sites lay between 3 and 6. For comparison, 99th WR percentiles for the simpler graphs ranged between 4 and 4.2 SE for all datasets sizes and topologies tested (Supplementary Text 1j).

The high WRs observed when correct models are fitted to the complex simulated data can be explained by 1/ difficulty in optimizing edge length and admixture proportions for a complex graph and 2/ ascertainment biases that cause expectations of certain $f_4$-statistics to be shifted away from 0. We reran the graph mapping protocol on bias-free, i.e. completely random, site samples from the complex simulations we observed that WRs dropped substantially: 99th WR percentiles ("unlocked" models) dropped from 5.1 to 4.2 SE on 95,000 sites, and from 6.3 to 4.2

SE on 400,000 sites (Fig. S30). 57% and 59% of "unlocked" models on 95,000 and 400,000 sites, respectively, had WR above 3 SE. We conclude that WR for a perfectly fitting complex model depends on the number of sites only in the presence of ascertainment bias, and WR for a perfectly fitting model drops to 4 SE and below in the absence of ascertainment bias. Given that WRs are affected by graph complexity and ascertainment biases, we propose to use CCC on all *f*-statistic residuals as an alternative measure of absolute model fit that is less affected by these problems.

*g. Conclusions*

Even a small-scale gene flow into an ancestry source proxy that was not shared by a group of interest resulted in very high FP rates for a *qpWave*/*qpAdm* protocol that singles out only one group as a target for modelling. As shown above, the same target can invariably fall between the spurious gene flow sources on a PC plane. An investigator unaware of these problems may easily conclude that an admixture event is well-supported by the data, albeit with a relatively low minor source proportion of 6 to 18%. To mitigate this problem, we propose to avoid *qpWave*/*qpAdm* protocols which single out a set of modelling targets and which are applied to sets of outgroups, potential sources and targets that are not well separated in time. Another way of mitigating the problem is avoiding SNP panels affected by ascertainment bias (*31*) since the bias and assumption violations have synergic effects on *qpAdm* performance.

We were able to find a model ranking scheme that allows to keep the FP rate of the graph mapping procedure below 5% even if a main ancestry source is mismodelled, but we caution that this scheme is applicable only to a particular target and a particular graph topology.

We also suggest that the standard absolute fit threshold for admixture graphs, i.e. WR = 3 SE, is too stringent for complex models. Even in the absence of ascertainment bias, $99^{th}$ WR percentile reached 4.2 SE for a collection of perfectly fitting models (15-populations, 11 unlocked admixture events). In the presence of ascertainment bias WR of a perfectly fitting model depends not only on the number of populations on the graph, but also on the number of sites, and may reach ~5 to 6 SE. Thus, reducing WR of a 15-population model with 11 unlocked admixture events to 3 SE and below may lead to overfitting.

3. Archaeological context of the Houtaomuga cemetery

Northeast China (Manchuria) represents a large geographic region that harbors three provinces including Liaoning (south), Jilin (central) and Heilongjiang (north). It has a markedly continental climate, with cold, dry winters and hot, rainy summers. The winter monsoon brings cold air masses from nearby Siberia, while the summer monsoon brings warm and moist air masses from the south. Located in the central lowland of northeast China lies the Northeast Plain ('Dongbei Pingyuan' in Chinese Pinyin). The plain is drained by the Songhua River and its tributary, the Nen River and was to a large extent a product of erosion from the surrounding highlands with fertile black soils. Although the growing season in this region is quite short, it also is one of the main areas for soybean, maize, rice and wheat cultivation. The Houtaomuga site is located on the gently sloping foothills along the southeast shore of Lake Xinhuangpao (formed by the tributary of the Nen River) in Da'an City, Jilin Province (N45º39′27.5", E123º47′15.1"). The site was first discovered in 1957. From 2011 to 2015 five seasons of excavations had been conducted by the Research Centre for Chinese Frontier Archaeology, Jilin University. A total of 110 burials, 552 pits, 30 ditches and 43 houses were excavated. These burials were divided into seven chronological phases based on the stratigraphy and were radiocarbon dated between ~12000 to 2300 cal yr BP. Burials from phase I to IV showed a clear evidence of sacrificial practices. More than 200 complete or restorable ceramic vessels have been discovered, along with a large number of microlithic tools, bone artifacts, a small number of jade objects as well as large numbers of fish and animal bones. Only a small number of large stone tools like grounding stones, grounding rods and stone axes since phase IV were unearthed. The inhabitants are believed to have practiced a mixed subsistence consisting of hunting-gathering-fishing complemented with limited millet cultivation since phase IV. We successfully obtained genomic data from 22 individuals from the Houtaomuga site that covers at least one individual per chronological phase. As no skeleton was excavated from the seventh phase, it was not discussed in this study.

*Houtaomuga-Phase I (13000-11000 cal yr BP) (n=1 sample)*
Only one individual was excavated from the first phase. An intact skeleton was recovered in an early Neolithic context in layer III in a vertical earthen pit. Samples from this individual were sent to different institutes for radiocarbon dating, and it was finally dated to 13000-11000 cal yr BP. The pits and ditches of this phase often contain large quantity of fish bones and shells, but only a few bones of terrestrial mammals. Houtaomuga phase I pottery is very similar to ceramics found at the Goncharka-I site located in the far southeast of Russia (the lower course of the Amour River).

**M45**

Genetic Sex: Female

Radiocarbon dates for M45:

| AMS ID | Conventional date (BP) | Calibrated date (2σ: 95.4%) (BP) | Materials and notes |
|---|---|---|---|
| ZK-4462 | 11700±30 | 11235-11145 | Fibula bone of M45 |

| | | | |
|---|---|---|---|
| ZK-3800 | 12899±33 | 12940-12850 | Pottery piece excavated from the same stratigraphic layer. |
| ZK-4464 | 11966±52 | 11760-11270 | Hip bone of a dog excavated from the same stratigraphic layer. |

*Houtaomuga-Phase II (7500-8000 cal yr BP) (n=3 samples)*
Three samples from the second phase, namely M54A, M54B and M54C, were buried in one tomb and came from a vertical earthen pit. The calibrated date for M54A was roughly 7500 cal yr BP. Like the first phase, the pits and ditches still mostly contain fish bones and shells, and a relatively large quantity of mammal bones. The lithic artifacts are quite similar to the ones of phase I, mainly composed of microlithic tools, while no agricultural tool has been found. The ceramic vessels of Houtaomuga phase II share their main characteristics with the ones discovered at the Gornoye site (late period of the Gromatuha culture) in the middle course of the Amur River in Russia.

**M54A**
Genetic Sex: Male
Radiocarbon Date: 8400±25 BP; 7430-7320 cal yr BP (ZK-4662)

**M54B**
Genetic Sex: Female
Radiocarbon Date: 8445±25 BP; 7470-7410 cal yr BP (ZK-3799)

**M54C**
Genetic Sex: N/A
Radiocarbon Date: 8380±25 BP; 7420-7350 cal yr BP (ZK-3797)

*Houtaomuga-Phase III (6000-5500 cal yr BP) (n=9 samples)*
Nine samples were selected and sequenced from the third phase (M50, M80, M82, M86, M98, M92, M93, M94, M95) and 2σ calibrated (95.4%) dates for those individuals range between 5900 and 5370 cal yr BP. The burial type was also a vertical earthen pit. However, there is a shift between the third and the first two phases that the pits and ditches begin to contain large quantities of mammal bones. Further, a small number of grounding stones and grounding rods have been discovered, but no tools associated with agricultural practice like stone shovel, hoe or knife had been found. The decoration of the pottery of Houtaomuga phase III shares a lot of traits with pottery discovered in the surrounding region of Jilin-Changchun, with the pottery of the Lower Xinle culture in the middle course of the Liao River, and with the pottery of the Fuhe culture in the west region of the Liao River valley, indicating that cultural interactions shifted to the south since this phase.

**M80**
Genetic Sex: Female
Radiocarbon Date: 6575±25 BP; 5460-5370 cal yr BP (BA-150167)

**M89**
Genetic Sex: Male
Radiocarbon Date: 7080±25 BP; 5940-5870 cal yr BP (BA-150173)

**M50**
Genetic Sex: N/A
Radiocarbon Date: 6970±25 BP; 5900-5650 cal yr BP (ZK-3795)

**M82**
Genetic Sex: N/A
Radiocarbon Date: 7105±30 BP; 5995-5880 cal yr BP (BA-150169)

**M86**
Genetic Sex: Male
Radiocarbon Date: 6770±30 BP; 5550-5470 cal yr BP (BA-150170)

**M92**
Genetic Sex: Female
Radiocarbon Date: 6740±25 BP; 5600-5570 cal yr BP (BA-150176)

**M93**
Genetic Sex: N/A
Radiocarbon Date: 6980±25BP; 5900-5710 cal yr BP (BA-150177)

**M94**
Genetic Sex: Male
Radiocarbon Date: 6745±25BP; 5550-5470 cal yr BP (BA-150178)

**M95**
Genetic Sex: Female
Radiocarbon Date: 6995±25BP; 5900-5720 cal yr BP (BA-150179)

*Houtaomuga-Phase IV (5000 cal yr BP) (n=2 samples)*
Two samples, M73 and M91 were from the fourth phase and were all excavated from vertical earthen pits. The calibrated date for M73 and M91 was around 5500 cal yr BP. Like the third phase, the pits and ditches still often contained mammal bones, mostly from wild ox and deer.

The remains from Houtaomuga phase IV are very close to the ones from the Yushushan site, Xiliang site in Aohan Banner and Haminmangha site in Kezuozhongqi in Inner Mongolia. In this phase primitive agriculture started to emerge based on the plants excavated.

**M91**
Genetic Sex: Female
Radiocarbon Date: 6175±25BP; 5590-5460 cal yr BP (BA-150175)

**M73**
Genetic Sex: Male
Radiocarbon Date: 6555±30BP; 5460-5370 cal yr BP (BA-150165)

*Houtaomuga-Phase V (2500-3000 cal yr BP) (n=1 sample)*
Only one sample (M25B) from the fifth phase was chosen for analysis in this study. All burial types in the fifth phase are vertical earthen pits. The calibrated date for M25B was around 2700 cal yr BP. In contrast to the four phases above, more burial objects were excavated, these included large numbers of bone tools, shells, jades and also a number of copper knives.

**M25B**
Genetic Sex: Male
Radiocarbon Date: 4420±35BP; 2720-2430 cal yr BP (ZK-3812)

*Houtaomuga-Phase VI (2300 cal yr BP) (n=6 samples)*
More than one hundred of skeletons were excavated from the sixth phase, of which six samples (M69A, M69B, M69C, M74A, M74B, M88) who were relatively better preserved by visual inspection were chosen for this genomic study. No direct radiocarbon dating on human skeletons was conducted from this phase, but one charcoal piece (12DHAIV F4) from a house foundation was dated to ca. 2300 cal yr BP. Unlike the burial types of phase I to phase V, a number of catacomb burial types were also discovered.

**12DHAIV F4**
Genetic Sex: N/A
Radiocarbon Date: 4249±23BP; 2360-2300 cal yr BP (ZK-4014)

4. DNA sequencing, read mapping, and contamination estimation

*Laboratory procedures for generating genetic data*

All the 22 petrous bone samples from the Houtaomuga cemetery were screened and laboratory protocols were carried out under strict aDNA guidelines (*82-84*) in dedicated clean facilities specially designed for aDNA studies at the Jilin University, China. We processed petrous bones according to a protocol described by Sirak *et al.* (*85*). DNA extraction was carried out in the clean-room facilities at the Jilin University using an in-solution silica-based protocol (*86*). Briefly, roughly 50mg bone powder from the inner part of petrous bones was digested in 900 µl of 0.5M EDTA, 16.7 µl of Proteinase K (15 mg/ml) and 83.3 µl $ddH_2O$ at 37 °C for 18 hours. The supernatant was then transferred to a MinElute silica spin column (Qiagen) after being fully mixed with 13 ml of a custom binding buffer [5M guanidine hydrochloride (MW 95.53), 40% isopropanol, 90 mM sodium acetate, and 0.05% Tween-20] and washed twice with the PE buffer containing 80% ethanol. DNA was then eluted with 100 µl of the TET buffer (1M Tris-HCl, 0.5M EDTA and 10% Tween-20). Double-stranded libraries were produced with truncated barcoded adaptors following the procedures by Meyer and Kircher (*87*) with minor modifications for some samples. Specifically, we prepared libraries for samples M45 and M69B using the *Bst* enzyme (NEW England Biolabs), while for the rest of the samples we replaced that polymerase with the Q5® High-Fidelity DNA Polymerase (NEW England Biolabs). In order to evaluate the quality of these samples, the libraries were first sequenced at low coverage on an Illumina HiSeq X10 instrument (a 2×150bp run). For samples with more than 10% endogenous human DNA, the libraries were sent for further sequencing aiming at a coverage of 1x. For the oldest Neolithic sample (M45), a further extract was prepared and was treated with uracil-DNA glycosylase (UDG). The library was then sequenced on further 8 full lanes on an HiSeq X10 sequencer, obtaining a high quality genome with a coverage of 45x.

*Sequence data processing*

Raw reads were first quality controlled by fastQC v.0.11.5 (*88*). Adapter sequences were trimmed and overlapping read pairs were merged using AdapterRemoval v2.2.0 (*89*) and sequences shorter than 30 base pairs (bp) were discarded. After adapter sequences were trimmed, we aligned the merged reads to the human reference genome (hg19) using BWA aln/samse v.0.7.12 (*90*) with the '-n 0.01' and '-l 1024' settings to allow for more mismatches and to disable seeding. PCR duplicates were removed by dedup v.0.12.2 (*91*), and sequences with a minimum mapping quality of 30 and base quality >30 were retained by SAMtools (*92*).

*Authentication and quality control*

We used several methods to measure authenticity of the ancient DNA data generated in this study. First, we tabulated the misincorporation patterns at both 5' (C>T) and 3' (G>A) ends (Data S3) using mapDamage v.2.0.6 (*93*). Those patterns are characteristic of ancient DNA. Second, we estimated mitochondrial DNA contamination for all individuals using the calico tool (https://github.com/pontussk/calico), which is based on the idea that mitochondrial DNA will have a single allele, and if the consensus sequence shows a variant that is either absent or present at a low frequency in a reference panel, then it is potentially contaminated. Third, we estimated the contamination of the nuclear genome in males using ANGSD v.0.910 (*94*) by measuring the polymorphism rate on the X chromosome (Data S3).

*Biological sex determination*
We determined the biological sex of the individuals reported in this study by counting the ratios of reads aligned to the Y chromosome and to the X chromosome to the total number of reads aligning to the autosomes. Then the difference between the two ratios was used to determine the genetic sex of our samples.

*Uniparental haplogroup assignment*
Y chromosome haplogroups were examined by aligning the BAM files to the sets of positions in the ISOGG 2016 (http://isogg.org/) and Y-full (https://www.yfull.com/tree/) databases. The Y chromosome haplogroups were than determined according to the most derived SNPs and were further confirmed by checking the corresponding upstream ancestral SNPs. We called mtDNA SNPs relying on reads aligned by SAMtools mpileup against the Revised Cambridge Reference Sequence (rCRS), and we double-checked SNPs by visual inspection of the alignments. Then, mtDNA sequences were assigned to different haplogroups using MitoTool (phylotree build 16).

*Genetic relatedness estimation*
We estimated the genetic relatedness between pairs of individuals reported in this study using the pairwise mismatch rate (pmr) method. This method was first introduced by Kennett *et al.* (*95*). The pmr value equals the number of SNPs for which two individuals share different alleles divided by the number of all SNPs for which both individuals harbor non-missing genotypes. In this sense, pmr between identical individuals (r = 1) is expected to be a half of that between two unrelated individuals (r = 0), and pmr values for the first- (r = 0.5) and second-degree relatives (r = 0.25) are 3/4 and 7/8 of that between unrelated pairs.

#### 5. Building skeleton admixture graphs

The best branching order found (Fig. 3a) is a unique model, and only the position of the 45000 years-old Ust'-Ishim individual (*96*) is unresolved because of a well-known trifurcation (*96, 97*). The deepest Asian branch in the graph is formed by Southeast and South Asian hunter-gatherers (*97-99*) here represented by present-day Onge from the Andaman Islands (India). The second is Tianyuan (*35*), the third is a late Jomon culture individual from Japan dated to ~2600 cal yr BP (*29, 98*), the fourth is an ancient Tibetan Chokhopani individual (*100*), then Southeast Asians (selected individuals having minimal Indian admixture), then First Americans represented by a collection of 13 present-day SAM individuals (Aymara, Mixe, Mixtec, Piapoco, Quechua, Yukpa, Zapotec) (*11*) having low degree of colonial admixture (Fig. S40, Supplementary Text 8), and an apical clade we term Siberian. The deepest branch in the Siberian clade is formed by the Houtaomuga group. The second branch is Baikal hunter-gatherers *sensu* Damgaard *et al.* (*41*) represented by ten 8000-7000 years-old individuals at the Shamanka site, then Paleo-Inuit represented by a 3900 years-old Saqqaq culture individual (*101*), and then present-day West Siberians: Selkups or Kets and Nganasans (*73*). We added by default three West Eurasian flows into each of the Siberian branches, but to keep the graph reasonably simple gene flows below 2% were removed from the model (see a list of West Eurasian flows in Fig. 3b). All Siberian groups on the skeleton graph have Mal'ta ancestry, except for three Houtaomuga individuals. All Siberians on the graph except for Houtaomuga and Saqqaq have WHG ancestry, and only Selkups have basal Eurasian ancestry possibly derived from Indo-Europeans at some point (*50*).

Two versions of the skeleton graph were constructed: a 14-population version lacking the Tianyuan and Onge groups, and a 16-population version including these groups. For constructing the 14-population version of the skeleton graph a dataset based on the 1240K SNP panel (*25*) without transitions was used, and ca. 95,000 sites were available across all groups on the graph. This skeleton graph has the worst $f_4$-statistic residual of 2.6 and includes 11 admixture events (9 unlocked). The model has 54 parameters and is based on 91 pairwise $f_2$-statistics. For constructing the more complex 16-population version of the skeleton graph a dataset based on the 1240K SNP panel (*25*) including transitions was, and ca. 408,000 sites were available across all groups on the graph. We avoided including more than 14 groups on the transition-free dataset to keep the number of sites available close to 100,000. Thus, for making the skeleton more complex transition sites were added. A multi-step procedure used for building these graphs is described below in detail.

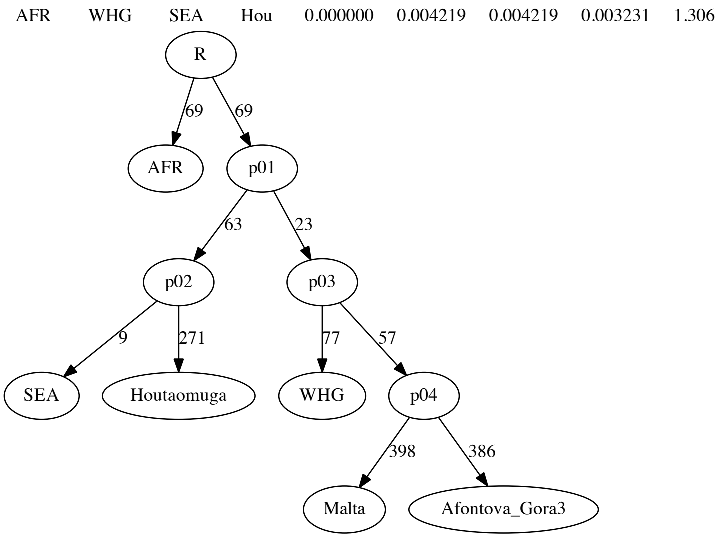


**Fig. S32.** A simple 6-population tree including the final WHG group of 11 individuals. The Houtaomuga group was composed of three individuals without trace West Eurasian ancestry (M45, M54A, M74B).

We started by constructing a simple 6-population tree (Fig. S32) on the transition-free dataset including an African meta-population of 45 present-day individuals (*11*) as an outgroup, Houtaomuga individuals, a Southeast Asian meta-population of 16 present-day individuals (*11*) from the SGDP project (*72*), Mal'ta and Afontova Gora3 (*40*), and a European hunter-gatherer (WHG) group. Houtaomuga individuals were placed on this graph one by one (Fig. S32), and a gene flow from the Mal'ta branch was added. Only individuals for whom inferred gene flow was 0% (HT-M45, HT-M54A, HT-M74B) were taken for subsequent analyses and were merged into one group.

At the next step we tested various individuals and groups on the WHG branch: Bichon, El Miron, Goyet Q116-1, Iberian HGs (Canes, Chan, La Braña), Koros I4971, Koros KO1, Kostenki14, Loschbour, Ostuni1, Ranchot88, Sunghir1, Sunghir2, Sunghir3, Sunghir4, Sunghir6, Vestonice16, and Villabruna. All the individuals/groups except for Kostenki14, Ostuni1, Sunghir1, Sunghir3, and Sunghir4 resulted in WRs below 2 SE, and we merged the following 11 individuals to form the WHG group for subsequent models: Bichon, Canes, Chan, El Miron, GoyetQ116-1, Koros KO1, LaBrana1, Loschbour, Ranchot88, Sunghir6, Villabruna. This composite WHG group on the 5-population tree resulted in a low WR of 1.3 SE. Vestonice16 was excluded from the final WHG group since it had a genetic component derived from the Kostenki-Sunghir clade, and we decided to focus on the WHG proper clade.

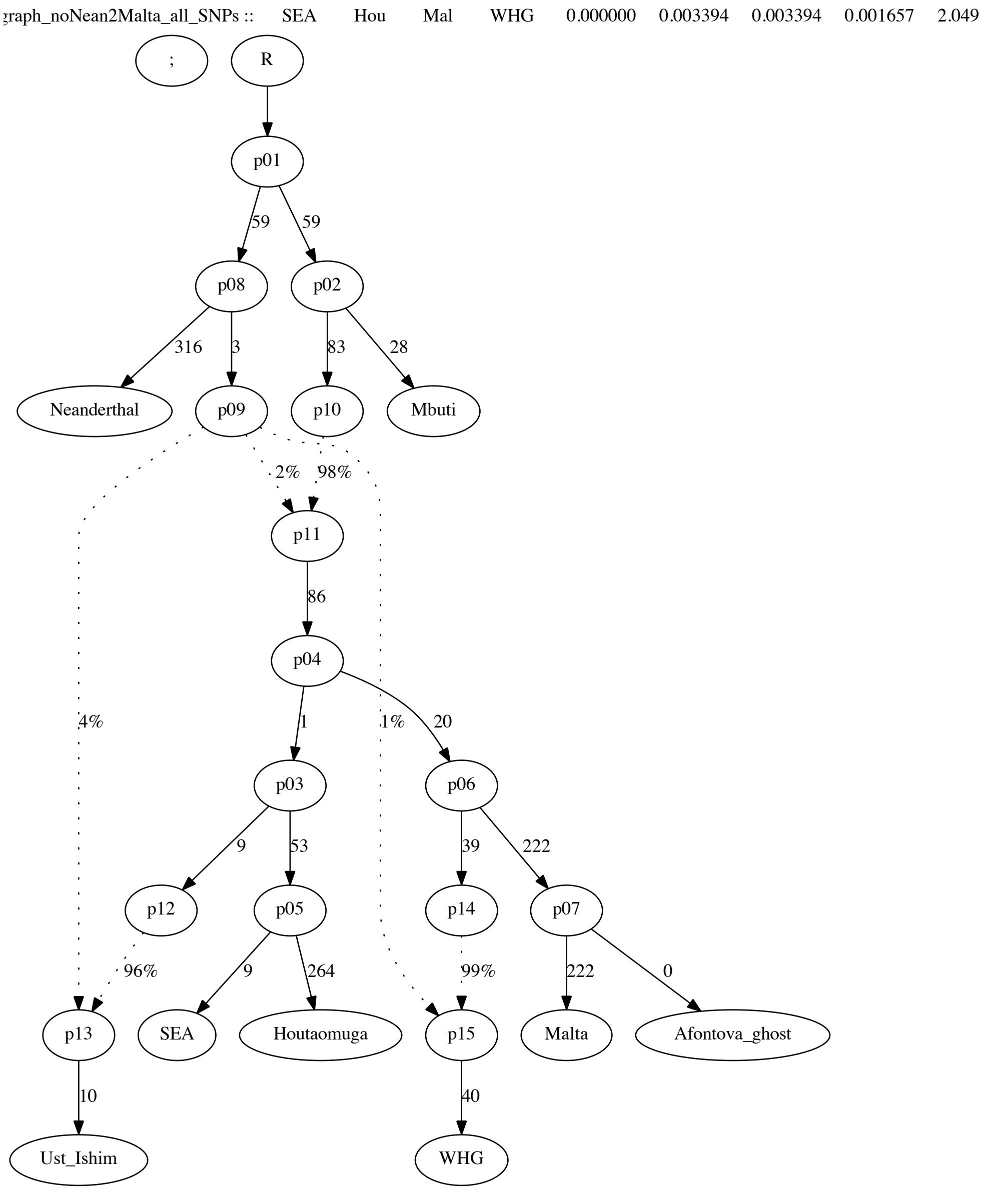


**Fig. S33.** A graph incorporating the Altai Neanderthal and Ust'-Ishim individuals.

Next, we removed the Afontova Gora3 individual to increase the number of sites available and replaced the large African meta-population with

Mbuti from SGDP (5 individuals), added the Ust'-Ishim individual (*96*), the Altai Neanderthal and Neanderthal admixture at the root of non-Africans (locked at 2%). Neanderthal admixture was also added to the Ust'-Ishim and WHG branches, i.e. to the oldest groups following Lipson and Reich (*97*). Consistent with previous results (*96*), Ust'-Ishim effectively forms a trifurcation with Europeans and Asians since three placements gave nearly identical WRs and LLs: as the most basal non-African branch, at the base of the European clade (Malta, WHG), and at the base of the Asian clade (SEA, Houtaomuga). Only the latter model had no trifurcation (i.e. no 0-length edge on the graph "backbone", Fig. S33) and was prioritized for subsequent work. On that graph the best placement of an American group, Karitiana (6 individuals) (*15, 72*), was found to be as a sister-group for Houtaomuga, with a gene flow from a Malta sister-group (*40*). The resulting graph had a WR of 2 SE.

On the graph in Fig. S33 we mapped 37 present-day and ancient Siberian groups as derived from four alternative Asian sources: on the Houtaomuga, Karitiana, SEA branches, or between the SEA and Karitiana split points. Three West Eurasian gene flows were added by default to each Siberian target: 1/ from a Mal'ta sister-group, 2/ from a WHG sister-group splitting before the Neanderthal gene flow into WHG, and 3/ from a basal Eurasian source splitting before the Neanderthal gene flow into non-Africans. The Siberian groups were taken from the following sources: present-day Altaian, Buryat, Even, Ket, Mansi, Nivkh, Tubalar, Ulchi, and Yakut from SGDP (*72*); present-day Altaian, Buryat, Even from Magadan, Even from the Sakha Republic, Evenk, Ket, Khanty, Mansi, Forest Nenets, Tundra Nenets, Nganasan, Selkup, Shor, Tuvinian, and Yakut from Pagani *et al.* (*73*); ancient Botai, Karasuk, Karasuk outlier, Kurma EBA, Lokomotiv EN, Okunevo, Saqqaq, Shamanka EBA, Shamanka EN, Ust'-Belaya EN-EBA, Ust'-Belaya outlier, Ust'-Ida LN, Ust'-Ida BA (*11, 41, 50, 101*). For 31 of 37 targets, the position on the Houtaomuga branch was the only "winning" model according to an LL threshold of 3 units (*11*). Of those best models for 31 targets, 30 models had WR below 3 SE and 18 models had WR below 2 SE. It was impossible to find a sole winning model for the following targets: Altaian, Mansi (2 groups), Tubalar, Tuvinian, Ust'-Belaya outlier. Based on this analysis we conclude that Houtaomuga is the closest relative of Siberians, and that major Asian ancestry lineages of Siberians probably form a clade.

At the previous step the following tree for Asian Siberian lineages was constructed: (Houtaomuga, ((Kurma EBA, Shamanka EBA), (Lokomotiv EN, Shamanka EN))). This topology is distinguished from other possible topologies by huge LL and WR differences, thus this tree is uncontroversial and unites Baikal HG into a clade composed of the Early Neolithic and Early Bronze Age sub-clades. By default, we added Mal'ta, WHG, and basal Eurasian (BE) gene flows into each Siberian branch except for Houtaomuga. Thus, we wanted to focus on the phylogeny of the Asian lineages although we recognize that this is a simplification of real history. To continue building the Siberian clade we mapped 33 present-day and ancient Siberian groups on this 5-population clade: all the groups listed above except for the four groups already incorporated into the graph. Five alternative placements were considered for each mapped target: 1/ as the most basal lineage splitting before Houtaomuga; 2/ as a lineage splitting between Houtaomuga and Baikal HG; 3/ on the Houtaomuga branch; 4/ on the ancestral Baikal EBA branch prior to West Eurasian gene flows; 5/ on the ancestral Baikal EN branch prior to West Eurasian gene flows (Table S10). Each Siberian group was mapped as derived from one Siberian source and three West Eurasian sources.

| no. | topology | classified as a "winner", % of targets | classified as a "sole winner", % of targets | average WR for "sole winners" |
|---|---|---|---|---|
| 1 | (SIB, (Houtaomuga, ((Kurma EBA, Shamanka EBA), (Lokomotiv EN, Shamanka EN)))) | 45% | 9% | 3.3 |
| 2 | (Houtaomuga, (SIB, ((Kurma EBA, Shamanka EBA), (Lokomotiv EN, Shamanka EN)))) | 70% | 30% | 2.7 |
| 3 | ((Houtaomuga, SIB), ((Kurma EBA, Shamanka EBA), (Lokomotiv EN, Shamanka EN))) | 30% | 0% | N/A |
| 4 | (Houtaomuga, ((SIB, (Kurma EBA, Shamanka EBA)), (Lokomotiv EN, Shamanka EN))) | 36% | 18% | 3.0 |
| 5 | (Houtaomuga, ((Kurma EBA, Shamanka EBA), (SIB, (Lokomotiv EN, Shamanka EN)))) | 0% | 0% | N/A |

**Table S10.** Results of mapping 33 Siberian groups onto five alternative positions in the five-population Siberian clade. Each target was mapped as an unadmixed Asian source + three West Eurasian sources. The Houtaomuga group was composed of three individuals without trace West Eurasian ancestry (M45, M54A, M74B).

Topology 5 (the position on the Baikal EN branch) was strongly rejected for all targets, with an average LL difference from the best model of ca. 6,800 units. Topology 4 (on the Baikal EBA branch) was declared as a "sole winner" supported by an LL difference >3 units for 18% of targets. Those unique models had an average WR of 3 SE (Table S10). Topology 3 (on the Houtaomuga branch) was not found among "sole winners". Topology 2 (between the Houtaomuga and Baikal HG branches) was declared as a "sole winner" supported by an LL difference >3 units for 30% of targets. Those unique models had an average WR of 2.7 SE (Table S10), thus their fits were nearly perfect. Topology 1 (the most basal Siberian lineage) was declared as a "sole winner" for 9% of targets (Altaians from SGDP, Buryat reported by Pagani *et al.* (*73*), and Tubalar from SGDP). Those unique models had an average WR of 3.3 SE (Table S10), thus the fits were not optimal. The same analysis was repeated for a group composed of two Houtaomuga individuals (M45, M54A) instead of three, generating nearly identical results (Table S11) to those discussed above. We conclude from this simple mapping experiment that none of the 33 target groups can be confidently derived from a single Asian lineage splitting before Houtaomuga.

| no. | topology | classified as a "winner", % of targets | classified as a "sole winner", % of targets | average WR for "sole winners" |
|---|---|---|---|---|
| 1 | (SIB, (Houtaomuga, ((Kurma EBA, Shamanka EBA), (Lokomotiv EN, Shamanka EN)))) | 39% | 9% | 3.3 |
| 2 | (Houtaomuga, (SIB, ((Kurma EBA, Shamanka EBA), (Lokomotiv EN, Shamanka EN)))) | 73% | 33% | 2.8 |
| 3 | ((Houtaomuga, SIB), ((Kurma EBA, Shamanka EBA), (Lokomotiv EN, Shamanka EN))) | 27% | 0% | N/A |
| 4 | (Houtaomuga, ((SIB, (Kurma EBA, Shamanka EBA)), (Lokomotiv EN, Shamanka EN))) | 36% | 18% | 2.9 |
| 5 | (Houtaomuga, ((Kurma EBA, Shamanka EBA), (SIB, (Lokomotiv EN, Shamanka EN)))) | 0% | 0% | N/A |

**Table S11.** Results of mapping 33 Siberian groups onto five alternative positions in the five-population Siberian clade. Each target was mapped as an unadmixed Asian source + three West Eurasian sources. The Houtaomuga group was composed of two oldest individuals without trace West Eurasian ancestry (M45, M54A).

For the Even and Evenk groups (4 in total) topology 2 was a clear winner supported by an LL difference from the second-best model between 9.6 and 20 units and demonstrating WRs between 2.4 and 3 SE. Thus, the Siberian tree was extended by adding an Evenk/Even clade and the resulting topology was as follows: (Houtaomuga, ((Evenk, Even), ((Kurma EBA, Shamanka EBA), (Lokomotiv EN, Shamanka EN)))). The Evenk (*73*) and Even groups (SGDP) were added one by one and mapping the other groups on the skeleton was repeated, but we do not show these results for brevity. The remaining Siberian groups were mapped on nine alternative positions in this seven-population Siberian clade (Table S12). Again, the most basal position never emerged as a clear "winner", i.e. as a best model differing by >3 LL units from a second-best model. In general, the WR of winning models grew as compared to the previous analysis (see Tables S10, S11), which suggests that most targets represent a mixture of two or more Asian sources or that some groups incorporated into the skeleton graph are admixed in the same way. Only 8 of 279 models tested demonstrated WR <3 SE.

| no. | topology | classified as a "winner", % of targets | classified as a "sole winner", % of targets | average WR for "sole winners" |
|---|---|---|---|---|
| 1 | (SIB, (Houtaomuga, ((Evenk , Even), ((Kurma EBA, Shamanka EBA), (Lokomotiv EN, Shamanka EN))))) | 10% | 0% | N/A |
| 2 | ((Houtaomuga, SIB), ((Evenk , Even), ((Kurma EBA, Shamanka EBA), (Lokomotiv EN, Shamanka EN)))) | 6% | 0% | N/A |
| 3 | (Houtaomuga, (SIB, ((Evenk , Even), ((Kurma EBA, Shamanka EBA), (Lokomotiv EN, Shamanka EN))))) | 19% | 0% | N/A |
| 4 | (Houtaomuga, ((Evenk , Even), (SIB, ((Kurma EBA, Shamanka EBA), (Lokomotiv EN, Shamanka EN))))) | 6% | 0% | N/A |
| 5 | (Houtaomuga, (((Evenk , Even), SIB), ((Kurma EBA, Shamanka EBA), (Lokomotiv EN, Shamanka EN)))) | 55% | 45% | 4.4 |
| 6 | (Houtaomuga, ((Evenk , Even), ((SIB, (Kurma EBA, Shamanka EBA)), (Lokomotiv EN, Shamanka EN)))) | 26% | 23% | 3.2 |
| 7 | (Houtaomuga, ((Evenk , Even), ((Kurma EBA, Shamanka EBA), (SIB, (Lokomotiv EN, Shamanka EN))))) | 0% | 0% | N/A |
| 8 | (Houtaomuga, (((Evenk , SIB), Even)), ((Kurma EBA, Shamanka EBA), (Lokomotiv EN, Shamanka EN))) | 10% | 10% | 3.5 |
| 9 | (Houtaomuga, (((Evenk , (Even, SIB)), ((Kurma EBA, Shamanka EBA), (Lokomotiv EN, Shamanka EN))) | 0% | 0% | N/A |

**Table S12.** Results of mapping 31 Siberian groups onto nine alternative positions in the seven-population Siberian clade. Each target was mapped as an unadmixed Asian source + three West Eurasian sources. The Houtaomuga group was composed of three individuals without trace West Eurasian ancestry (M45, M54A, M74B).

Among three winning models for the Saqqaq Paleo-Inuit (*101*) all had WR <3 SE, but just one had no trifurcation in the Asian clade. That topology was (Houtaomuga, (((Evenk, Even), Saqqaq), ((Kurma EBA, Shamanka EBA), (Lokomotiv EN, Shamanka EN)))). The remaining 30 groups were again mapped on 11 alternative positions in this eight-population Siberian clade (Table S13). The most basal position never emerged as a clear "winner". Only 4 of 330 models tested demonstrated WR <3 SE, suggesting that non-admixed topologies for Siberian Asian lineages on this graph are being exhausted.

Since we cannot be sure that the Asian lineages included into the skeleton (Houtaomuga, (((Evenk, Even), Saqqaq), ((Kurma EBA, Shamanka EBA), (Lokomotiv EN, Shamanka EN)))) indeed form a simple tree, we used a different approach for finding a small set of non-admixed Siberian lineages. Since no group in our set of 37 groups was confidently modelled as derived from a single source diverging deeper that Houtaomuga, we tested all possible population quadruplets of the type (Houtaomuga, ($Siberian_1$, ($Siberian_2$, ($Siberian_3$, $Siberian_4$)))). Each Siberian was modelled with the three West Eurasian gene flows. To limit the number of combinations, we removed some closely related groups from the set of 37 Siberians used above. The resulting set of 27 groups was as follows: present-day Altaian, Buryat, Even, Ket, Mansi, Nivkh, Tubalar, Ulchi, and Yakut (*72*); present-day Evenk, Khanty, Forest Nenets, Tundra Nenets, Nganasan, Shor, and Tuvinian (*73*); ancient Botai, Karasuk, Karasuk outlier, Okunevo, Saqqaq, Shamanka EBA, Shamanka EN, Ust'-Belaya EN-EBA, Ust'-Belaya outlier, Ust'-Ida LN, Ust'-Ida BA (*11, 41, 50, 101*). In total, 210,600 models were tested: 12 alternative branching orders and 17,550 population combinations. The Houtaomuga group was composed of three individuals without trace West Eurasian ancestry (M45, M54A, M74B).

| no. | topology | classified as a "winner", % of targets | classified as a "sole winner", % of targets | average WR for "sole winners" |
|---|---|---|---|---|
| 1 | (SIB, (Houtaomuga, ((Saqqaq, (Evenk , Even)), ((Kurma EBA, Shamanka EBA), (Lokomotiv EN, Shamanka EN))))) | 7% | 0% | N/A |
| 2 | ((Houtaomuga, SIB), ((Saqqaq, (Evenk , Even)), ((Kurma EBA, Shamanka EBA), (Lokomotiv EN, Shamanka EN)))) | 10% | 10% | 3.6 |
| 3 | (Houtaomuga, (SIB, ((Saqqaq, (Evenk , Even)), ((Kurma EBA, Shamanka EBA), (Lokomotiv EN, Shamanka EN))))) | 0% | 0% | N/A |
| 4 | (Houtaomuga, ((Saqqaq, (Evenk , Even)), (SIB, ((Kurma EBA, Shamanka EBA), (Lokomotiv EN, Shamanka EN))))) | 7% | 0% | N/A |
| 5 | (Houtaomuga, ((SIB, (Saqqaq, (Evenk , Even))), ((Kurma EBA, Shamanka EBA), (Lokomotiv EN, Shamanka EN)))) | 17% | 3% | 5.6 |
| 6 | (Houtaomuga, ((Saqqaq, (Evenk , Even)), ((SIB, (Kurma EBA, Shamanka EBA)), (Lokomotiv EN, Shamanka EN)))) | 3% | 0% | N/A |
| 7 | (Houtaomuga, ((Saqqaq, (Evenk , Even)), ((Kurma EBA, Shamanka EBA), (SIB, (Lokomotiv EN, Shamanka EN))))) | 7% | 0% | N/A |
| 8 | (Houtaomuga, ((Saqqaq, (SIB, (Evenk , Even))), ((Kurma EBA, Shamanka EBA), (Lokomotiv EN, Shamanka EN)))) | 27% | 27% | 3.2 |
| 9 | (Houtaomuga, (((Saqqaq, SIB), (Evenk , Even)), ((Kurma EBA, Shamanka EBA), (Lokomotiv EN, Shamanka EN)))) | 0% | 0% | N/A |
| 10 | (Houtaomuga, ((Saqqaq, ((SIB, Evenk ), Even))), ((Kurma EBA, Shamanka EBA), (Lokomotiv EN, Shamanka EN))) | 53% | 47% | 4.3 |
| 11 | (Houtaomuga, ((Saqqaq, (Evenk , (SIB, Even))), ((Kurma EBA, Shamanka EBA), (Lokomotiv EN, Shamanka EN)))) | 7% | 0% | N/A |

**Table S13.** Results of mapping 30 Siberian groups onto 11 alternative positions in the eight-population Siberian clade. Each target was mapped as an unadmixed Asian source + three West Eurasian sources. The Houtaomuga group was composed of three individuals without trace West Eurasian ancestry (M45, M54A, M74B).

For each population combination, models were ranked according to LL, and a combination was considered as "informative" if it satisfied the following conditions: 1/ no trifurcations in the Siberian clade; 2/ LL difference between the best and the second-best model >3 units; 3/ WR <4 SE. Of 17,550 population combinations, 4,038 turned out to be "informative". Then we aligned the branching orders in the informative quadruplets manually, starting from quadruplets demonstrating the largest LL difference between best and second-best models and low WR <3SE. The resulting alignment of non-contradictory branching orders revealed two major sister clades: 1/ a clade including all Baikal HG ancient groups (*41*) and both Karasuk culture groups (*50*); 2/ Kets, Khanty, Mansi, both Nenets groups, and Nganasans. A non-ambiguous placement was not found for all the remaining groups. The following groups tended to diverge deeper than the split point of the Baikal HG and West Siberian clades: Altaian, Botai culture, Buryat, Nivkh and Ulchi, Okunevo culture, Shor and Tubalar, Tuvinian. This analysis, although inconclusive, allowed us to grasp few general properties of the Siberian tree without explicit exploration of all possible topologies for all population combinations.

Next, we proceeded to adding further Asian lineages: Ikawazu Jomon (*98*) and various Tibetans. Before doing that, we optimized the composition of the SEA group. We tested 26 SEA individuals from SGDP (*72*) one by one on a skeleton graph with Dinka (*72*) as the African lineage, Pima (*72*) as the American lineage, and Houtaomuga and Lokomotiv EN (the latter receiving a Mal'ta-related gene flow) as two Siberian groups. We added a Mal'ta-related gene flow into these SEA individuals, and selected only those individuals that demonstrated inferred West Eurasian ancestry of 0% and WR <2 SE. Those were individuals Igorot 2, Miao 1, and She 1, and they comprised an updated SEA group supposedly free of West Eurasian (e.g., Indian) admixture. We mapped the Ikawazu Jomon individual, present-day Tibetans and Sherpa from SGDP (4 individuals), and ancient Tibetans (5 individuals) (*100*) one by one on seven alternative positions in the Asian clade (Table S14). Only one, the most basal, position was a clear winner for 9 of 10 targets, supported by LL differences ranging from 3 to 100 units. WRs of these winning models ranged from 2 to 4.4 SE, 2.9 SE on average. Based on these low WR values we concluded that Jomon. and Tibetans can be approximately modelled as unadmixed lineages.

| no. | topology | classified as a "winner", % of targets | classified as a "sole winner", % of targets | average WR for "sole winners" |
|---|---|---|---|---|
| 1 | (X, (SEA, (FAM, (Houtaomuga, Lokomotiv EN)))) | 100% | 90% | 2.9 |
| 2 | ((SEA, X), (FAM, (Houtaomuga, Lokomotiv EN))) | 10% | 0% | N/A |
| 3 | (SEA, (X, (FAM, (Houtaomuga, Lokomotiv EN)))) | 10% | 0% | N/A |
| 4 | (SEA, ((FAM, X), (Houtaomuga, Lokomotiv EN))) | 0% | 0% | N/A |
| 5 | (SEA, (FAM, (X, (Houtaomuga, Lokomotiv EN)))) | 0% | 0% | N/A |
| 6 | (SEA, (FAM, ((Houtaomuga, X) ,Lokomotiv EN))) | 0% | 0% | N/A |
| 7 | (SEA, (FAM, (Houtaomuga, (Lokomotiv EN, X)))) | 0% | 0% | N/A |

**Table S14.** Results of mapping 10 Tibetan and Jomon groups onto 7 alternative positions in the four-population Asian clade. Each target was mapped as an unadmixed lineage. The Houtaomuga group was composed of three individuals without trace West Eurasian ancestry (M45, M54A, M74B). The SEA group was composed of three individuals (Igorot 2, Miao 1, and She 1, Mallick *et al.* (*72*)).

Then we tested three possible branching orders of the Jomon and Tibetan lineages at the basal position in the Asian clade: 1/ (Jomon, (Tibetan, other Asians)), 2/ (Tibetan, (Jomon, other Asians)), and 3/ ((Tibetan, Jomon), other Asians). The basal position of Jomon was supported by an LL difference of 54 units and a WR difference of 3.7 SE. The winning model had a WR of 2.98 SE and included the Tibetan 1 present-day individual (*72*) as a representative of the Tibetan lineage. We also tested Jomon/Tianyuan (*35*) and Tianyuan/Tibetan combinations in the same way. The basal position of Tianyuan in these combinations was supported by very large LL differences of 133 or 376 units, and model (Tianyuan, (Jomon, (Tibetan, other Asians))) had a WR of 2.99 and no trifurcations in the Asian clade. This topology is shown in Fig. S34. To increase the number of sites available under the "useallsnps: NO" setting, we dropped Tianyuan from subsequent models on the transition-free dataset.

We also performed an exhaustive exploration of all possible branching orders in the Asian clade (but this time using different representatives of some groups): Ikawazu Jomon, an ancient individual Tibetan Chokhopani (*100*), SEA (the three individuals selected above), a meta-population (*11*) of 13 present-day Native Americans mostly from Mexico (Aymara, Mixe, one Mixtec individual without colonial admixture, Piapoco, Quechua, Yukpa, Zapotec), and Houtaomuga (the three individuals selected above). Bantu Tswana (*72*) was used as a representative African group. This adjustment was done to check robustness of the findings.

Unlike the tests above, not only all possible positions of a newly added group on a skeleton phylogeny, but all possible phylogenies for the selected groups were explored. Of 105 models tested, three models were drastically different from the others according to LL (a difference of at least 22 units) and WR (a difference of at least 1.6 SE). The three competing topologies with nearly identical WR and LL values were: 1/ (Jomon, (Tibetan, (SEA, ($FAM_{EA}$, Houtaomuga)))) (the best one according to LL and WR), 2/ (Jomon, ((SEA, Tibetan), ($FAM_{EA}$, Houtaomuga))), and 3/ (Jomon, (SEA, (Tibetan, ($FAM_{EA}$, Houtaomuga)))), where $FAM_{EA}$ is the Asian source group for First Americans (in contrast to the Mal'ta-related source group). Since topologies 2 and 3 resulted in a trifurcation in the Asian clade and topology 1 did not, we selected the latter topology (Fig. S35). The model had a WR of 2.8 SE.

**Fig. S34 (left).** A graph incorporating three basal Asian lineages: the Tianyuan (*35*), Jomon (*98*), and Tibetan 1 individuals (*72*). The Houtaomuga group was composed of three individuals without trace West Eurasian ancestry (M45, M54A, M74B). The SEA group was also composed of three individuals without trace West Eurasian ancestry (Igorot 2, Miao 1, She 1, Mallick *et al.* (*72*)).

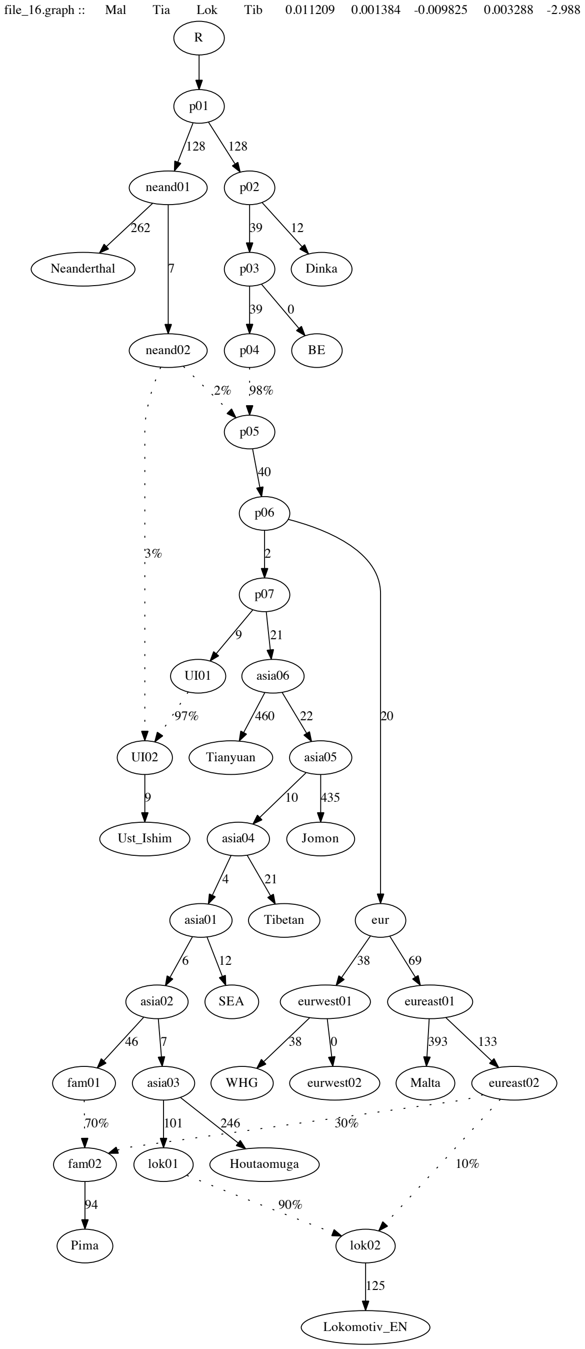


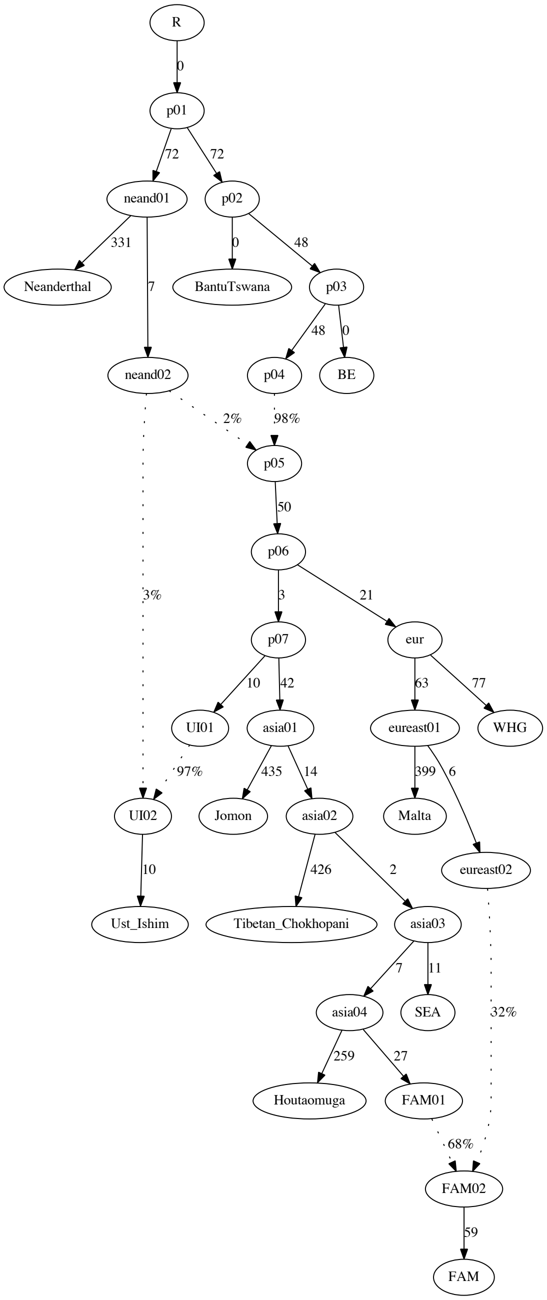


**Fig. S35 (right).** A graph incorporating two basal Asian lineages: the Jomon (*98*) and Tibetan Chokhopani individuals (*100*). The Houtaomuga group was composed of three individuals without trace West Eurasian ancestry (M45, M54A, M74B). The SEA group was also composed of three individuals without trace West Eurasian ancestry (Igorot 2, Miao 1, She 1, Mallick *et al.* (*72*)). The FAM group (*11*) was composed of 13 present-day Native Americans mostly from Mexico.

At this point we returned to building the Siberian clade, hoping that the Jomon and Tibetan groups incorporated into the skeleton will help us to resolve the branching order of Siberian lineages. We selected closely related groups Shamanka EN or Lokomotiv EN as representatives of the Baikal HG clade, Kets (or closely related Selkups) and Nganasans as representatives of the second, West Siberian, clade. We also selected Saqqaq and Houtaomuga as representatives of major archaeological cultures and clearly divergent ancestries (*10*) and tested 105 possible rooted trees for these five lineages. Unlike the previous tests on the Siberian clade, all possible branching orders were explored, but for a particular combination of groups. Similar to the previous tests on the Siberian clade, all Siberians except for Houtaomuga were modelled as derived from one Asian source and three West Eurasian sources. The branching order (Houtaomuga, (Shamanka EN, (Saqqaq, (Selkup, Nganasan)))) was supported by an LL difference of 7.6 units and a WR difference of 0.26 SE vs. a second-best model (Houtaomuga, ((Saqqaq, Shamanka EN), (Selkup, Nganasan))), the WR of the best model was 2.6 SE, and there were no trifurcations in the Asian clade unlike the second-best model. This Siberian topology was selected for further analyses (Fig. S36). West Eurasian gene flows into Siberians of <2% were later removed to keep the number of model parameters reasonably low.

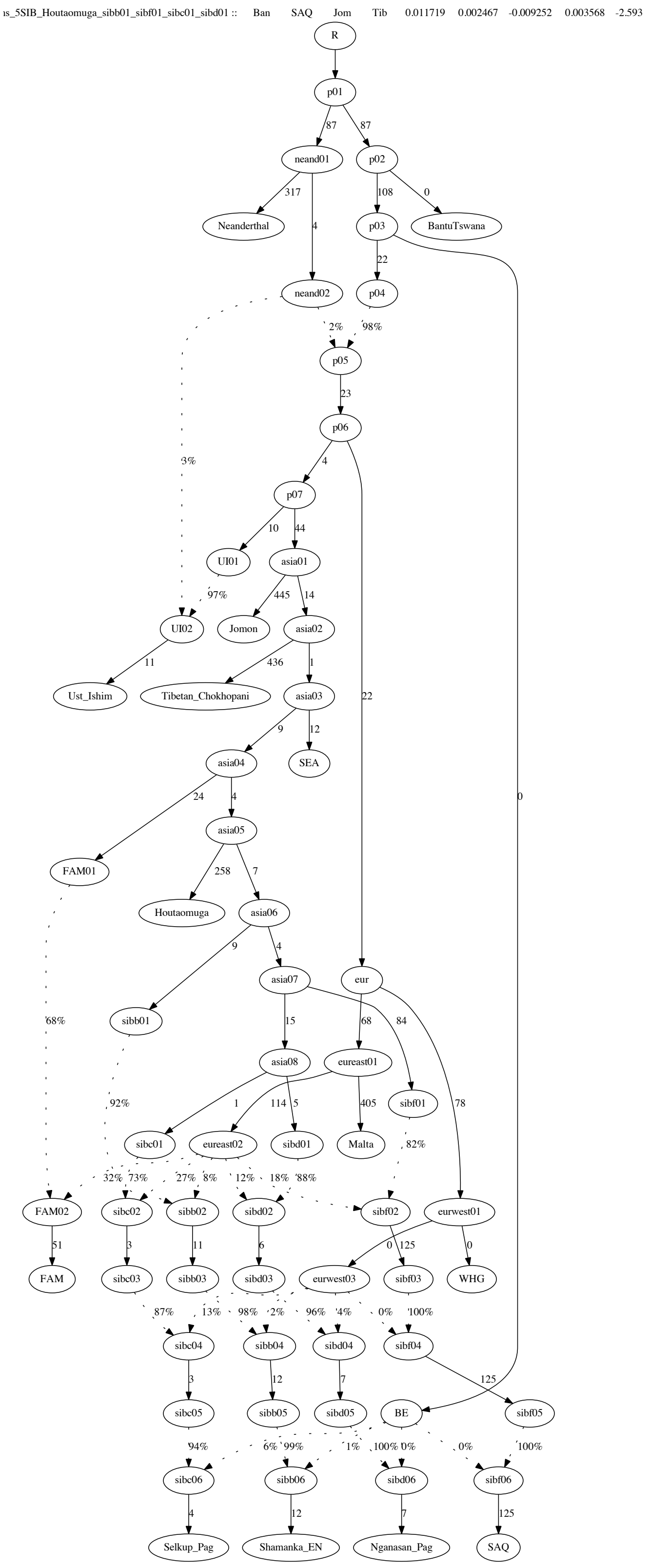


**Fig. S36.** A graph incorporating two basal Asian lineages (Jomon and Tibetan Chokhopani) and five Siberian groups. The Houtaomuga group was composed of three individuals without trace West Eurasian ancestry (M45, M54A, M74B). The SEA group was also composed of three individuals without trace West Eurasian ancestry (Igorot 2, Miao 1, She 1, Mallick *et al.* (*72*)). The FAM group (*11*) was composed of 13 present-day Native Americans mostly from Mexico.

After incorporating the Tibetan and Ikawazu Jomon lineages and constructing the five-population Siberian clade, we tested combinations of groups on the African (21 present-day groups from SGDP), American (Anzick, USR1, 8 present-day groups, and a composite present-day population (*11*)), Baikal HG (the Lokomotiv EN and Shamanka EN groups were tested), Selkup (the Ket SGDP and Selkup groups were tested), and Tibetan branches (present-day Tibetans and Sherpa from SGDP, and ancient Tibetans reported by Jeong *et al.* (*100*) were tested individual by individual, 9 individuals in total). The other branches including Jomon, SEA, Houtaomuga, Saqqaq, and Nganasan were kept invariant. This analysis was done to find a skeleton graph with a low WR and a combination of groups yielding a high number of transversion sites. In total, 8,316 group combinations were tested, under three admixture proportion locking schemes: West Eurasian gene flows into the Siberian and/or Tibetan lineages were locked according to values estimated under a starting model. The resulting models (24,948) were sorted according to the number of trifurcations in the Asian clade and then according to WR. The following combination of variable groups was selected as a "winner" since it had the lowest WR for any combination resulting in no trifurcations in the Asian clade and having 95,000 or more transversion sites available: Bantu Tswana from SGDP (*72*), the meta-population of present-day Native Americans (Aymara, Mixe, one Mixtec individual without colonial admixture, Piapoco, Quechua, Yukpa, Zapotec, Flegontov *et al.* (*11*)), Shamanka EN (*41*), Selkup (*73*), ancient Tibetan Chokhopani (*100*). The WR for that combination was 2.6 SE. This combination was selected as a draft skeleton graph for all subsequent work.

| **Native American population** | **median absolute Z-score (WR)** | **minimal absolute Z-score (WR)** | **number of models with no trifurcatio ns in the Asian clade** | **percentage of models with no trifurcatio ns in the Asian clade** |
|---|---|---|---|---|
| Chane | 3.7 | 2.5 | 525 | 23.1% |
| anc. Anzick (Clovis) | 3.6 | 2.2 | 79 | 3.5% |
| composite FAM | 3.6 | 2.4 | 1392 | 61.4% |
| Huichol | 3.6 | 2.3 | 300 | 13.2% |
| Karitiana | 3.9 | 2.8 | 1429 | 63.0% |
| Mayan | 11.6 | 8.0 | 556 | 24.5% |
| Mixtec | 6.2 | 4.3 | 1194 | 52.6% |
| Nahua | 3.6 | 2.4 | 1307 | 57.6% |
| Pima | 3.5 | 2.2 | 1378 | 60.8% |
| Surui | 3.6 | 2.3 | 967 | 42.6% |
| anc. USR1 | 3.6 | 2.2 | 0 | 0.0% |

**Table S15.** Results for model combinations including 11 alternative Native American groups on the draft skeleton graph. Each American group was represented in 2,268 combinations with other populations on the graph.

Remarkably, among 11 American groups tested (Chane, ancient Anzick, the meta-population of 13 present-day Native Americans mostly from Mexico, Huichol, Karitiana, Mayan, Mixtec, Nahua, Pima, Surui, ancient USR1 pseudohaploid version), models incorporating USR1 invariably had at least one trifurcation in the Asian clade, while for the other groups from 3.5% to 63% of 2,268 population combinations demonstrated no trifurcations (40% on average, see Table S15). The lowest percentage was recorded for the Anzick ancient individual, and the highest for Karitiana. For example, in a model including Juǀ’hoan North, USR1, Shamanka EN, Selkup, and the Tibetan-2 individual, we encountered trifurcations (Tibetan, SEA, (USR1, Siberian)) and (USR1, Houtaomuga, other Siberians). That model was based on ca. 98,000 sites and had a WR of 2.5 SE. In general, the median WR of models incorporating USR1 was on a par with those incorporating other Americans: models incorporating Chane, ancient Anzick, the composite FAM group, Huichol, Karitiana, Nahua, Pima, Surui, or USR1 all had median WR between 3.5 and 3.9 SE and the lowest WR between 2.2 and 2.8 SE (Table S15). Thus, although it is possible to model diverse Siberians and USR1 (derived from an unadmixed Asian source) on a well-fitting graph demonstrating WR down to 2.2 SE, the resulting models are likely sub-optimal since they invariably contain trifurcations.

Here we show the skeleton graph selected at the previous step: its original representation with gene flows into the Tibetan branch locked at 0% (Fig. S37a) and its final cleaned up representation (Fig. S37b).

Then we added the Tianyuan and Onge branches to the Asian clade, and to increase the number of sites available we switched to the full dataset including transitions. We performed an exhaustive test of all possible branching orders in an Asian clade including: Onge, Tianyuan, Ikawazu Jomon, an ancient individual Tibetan Chokhopani (*100*), SEA (the three individuals selected above), the meta-population (*11*) of 13 present-day Native Americans mostly from Mexico, and Houtaomuga (the three individuals selected above). Of 10,395 models tested, two models were significantly different from the others according to LL (a difference of at least 3.4 units), and the same models had no trifurcations in the Asian clade, in contrast to the other models in the same LL range. Nine models were different from the others according to WR (a difference of at least 0.67 SE). Thirteen best topologies ranked by LL are shown in Table S16.

| no. | topology | LL | WR | LL difference vs. the best model | WR difference vs. the best model | trifurcations in the Asian clade |
|---|---|---|---|---|---|---|
| 1 | (Tianyuan, (Onge, (Jomon, (Tibetan_Chokhopani, (SEA, (FAM, Houtaomuga)))))) | 149.2 | 5.81 | | | 0 |
| 2 | (Onge, (Tianyuan, (Jomon, (Tibetan_Chokhopani, (SEA, (FAM, Houtaomuga)))))) | 153.2 | 5.83 | 4.0 | 0.02 | 0 |
| 3 | ((Onge, Tianyuan), (Jomon, (Tibetan_Chokhopani, (SEA, (FAM, Houtaomuga))))) | 156.6 | 5.83 | 7.4 | 0.02 | 1 |
| 4 | (Tianyuan, (Onge, (Jomon, ((FAM, Houtaomuga), (SEA, Tibetan_Chokhopani))))) | 165.2 | 5.81 | 16.0 | 0.00 | 1 |
| 5 | (Tianyuan, (Onge, (Jomon, (SEA, (Tibetan_Chokhopani, (FAM, Houtaomuga)))))) | 165.2 | 5.81 | 16.0 | 0.00 | 1 |
| 6 | (Onge, (Tianyuan, (Jomon, ((FAM, Houtaomuga), (SEA, Tibetan_Chokhopani))))) | 168.4 | 5.83 | 19.2 | 0.02 | 1 |
| 7 | (Onge, (Tianyuan, (Jomon, (SEA, (Tibetan_Chokhopani, (FAM, Houtaomuga)))))) | 168.4 | 5.83 | 19.2 | 0.02 | 1 |
| 8 | ((Onge, Tianyuan), (Jomon, ((FAM, Houtaomuga), (SEA, Tibetan_Chokhopani)))) | 171.8 | 5.83 | 22.7 | 0.02 | 2 |
| 9 | ((Onge, Tianyuan), (Jomon, (SEA, (Tibetan_Chokhopani, (FAM, Houtaomuga))))) | 171.8 | 5.83 | 22.7 | 0.02 | 2 |
| 10 | (Tianyuan, (Onge, (Jomon, (Tibetan_Chokhopani, (Houtaomuga, (FAM, SEA)))))) | 198.2 | 6.47 | 49.0 | 0.67 | 1 |
| 11 | (Tianyuan, (Onge, (Jomon, (Tibetan_Chokhopani, (FAM, (Houtaomuga, SEA)))))) | 198.2 | 6.47 | 49.1 | 0.67 | 1 |
| 12 | (Onge, (Tianyuan, (Jomon, (Tibetan_Chokhopani, (Houtaomuga, (FAM, SEA)))))) | 199.6 | 6.47 | 50.5 | 0.67 | 1 |
| 13 | (Onge, (Tianyuan, (Jomon, (Tibetan, Chokhopani, (FAM, (Houtaomuga, SEA)))))) | 199.6 | 6.47 | 50.5 | 0.67 | 1 |

**Table S16.** Thirteen best topologies for the Asian clade on the full dataset.

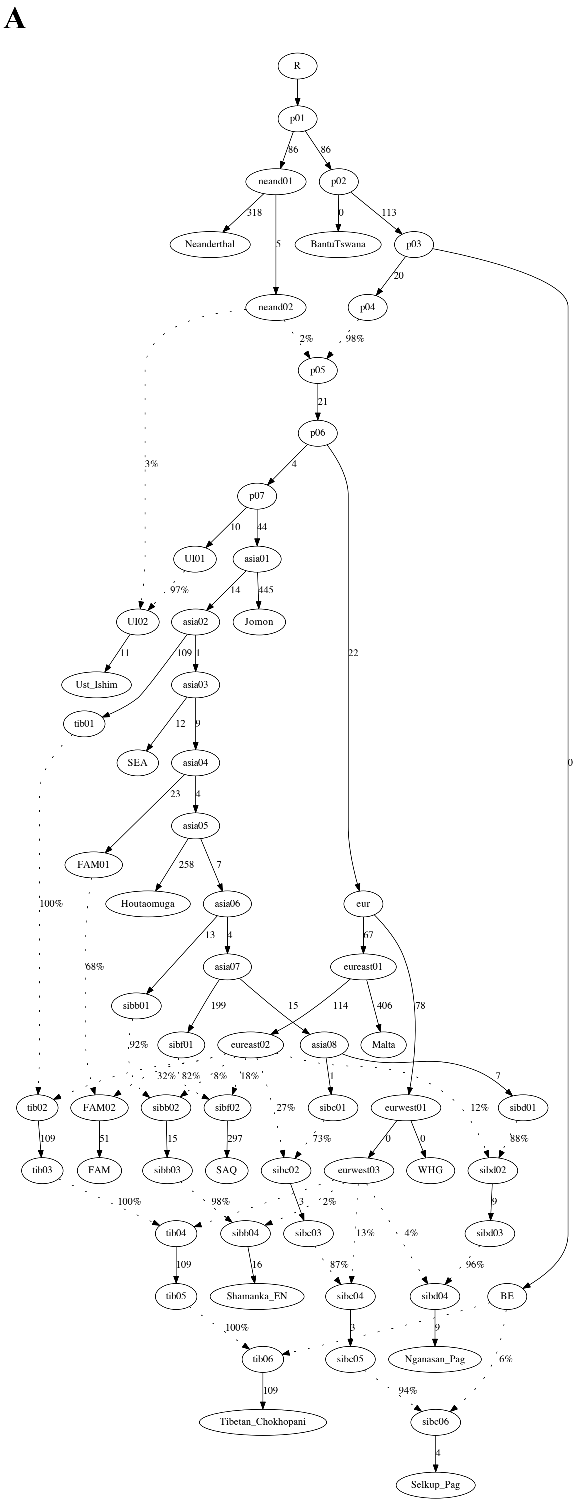


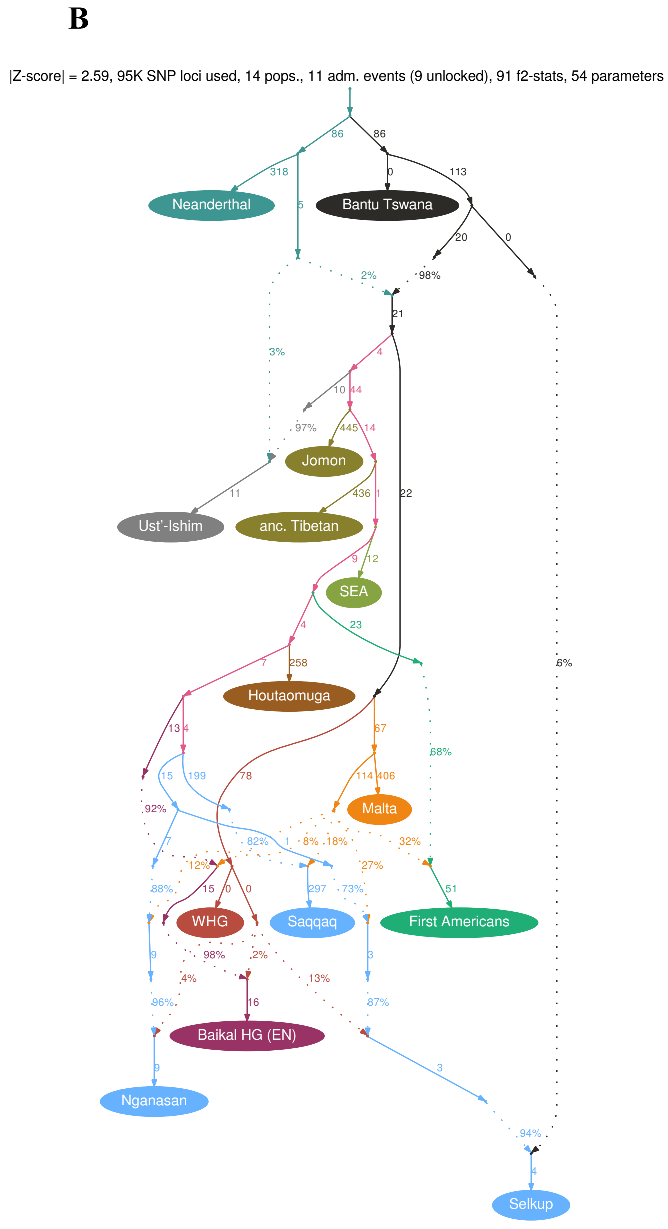


**Fig. S37.** A skeleton graph selected on the transition-free dataset: its original representation with gene flows into the Tibetan branch locked at 0% (**A**) and its final clean representation (**B**).

This topology exploration confirmed that the Tibetan branch likely diverged before the SEA branch since the respective models had significantly different LL. The order of the Tianyuan and Onge branches was hard to resolve, and both highest-ranking models (no. 1 and 2 in Table S16) had no trifurcations in the Asian clade. We note that similar topologies for major Asian lineages

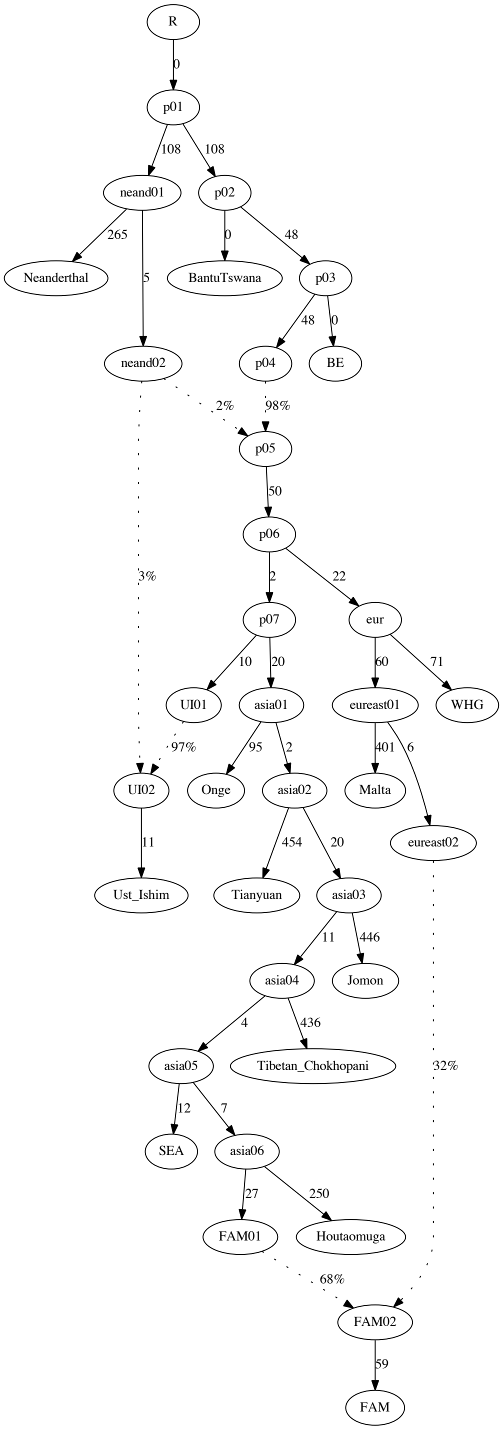


were reported by two studies utilizing *TreeMix*, a conceptually different method. Gakuhari *et al.* (*29*) reported the following topology (only groups we analyzed are shown here): (Ust'-Ishim, ((WHG, Mal'ta), (Onge, (Tianyuan, (Jomon, ((Tibetan Chokhopani, Sherpa), (SEA, (Siberians, $FAM_{EA}$)))))))). This topology is identical to model 2 in Table S16, with the exception of Ust'-Ishim's position. Yang *et al.* (*102*) reported a slightly different topology: (Ust'-Ishim, (Yana [a Mal'ta relative (*10*)], (Tianyuan, (SEA, (Tibetan Chokhopani, Siberians))))). Summarizing these results, we conclude that the branching order of major Asian lineages is uncontroversial, except for the branching order of Onge and Tianyuan, and potentially of Tibetans and SEA. Since we failed to resolve two best branching orders (models 1 and 2 in Table S16), we preferred model 2 since it was consistent with a published finding.

The topology selected is shown in Fig. S38. When the five-population Siberian sub-graph was grafted onto this Asian graph, a final skeleton (Fig. S39) had a WR of 5.9 SE on the full dataset. Considering the results on simulated data where in the presence of ascertainment bias WR for a very similar correct graph on 400K sites ranged from ~3 to 6 SE (Fig. S30), we decided to stop improving the skeleton further to avoid overfitting. Thus, we generated two similar skeleton graphs: the 14-population graph on the transition-free dataset of 95K sites (Fig. S37), and the 16-population graph on the full dataset of 408K sites (Fig. S39). These two skeleton graphs were used in all subsequent mapping work.

**Fig. S38.** A graph (model 2 in Table S16) incorporating four basal Asian lineages: the Onge (*72*), Tianyuan (*35*), Jomon (*98*) and Tibetan Chokhopani individuals (*100*). The Houtaomuga group was composed of three individuals without trace West Eurasian ancestry (M45, M54A, M74B). The SEA group was also composed of three individuals without trace West Eurasian ancestry (Igorot 2, Miao 1, She 1, Mallick *et al.* (*72*)). The FAM group (*11*) was composed of 13 present-day Native Americans mostly from Mexico.

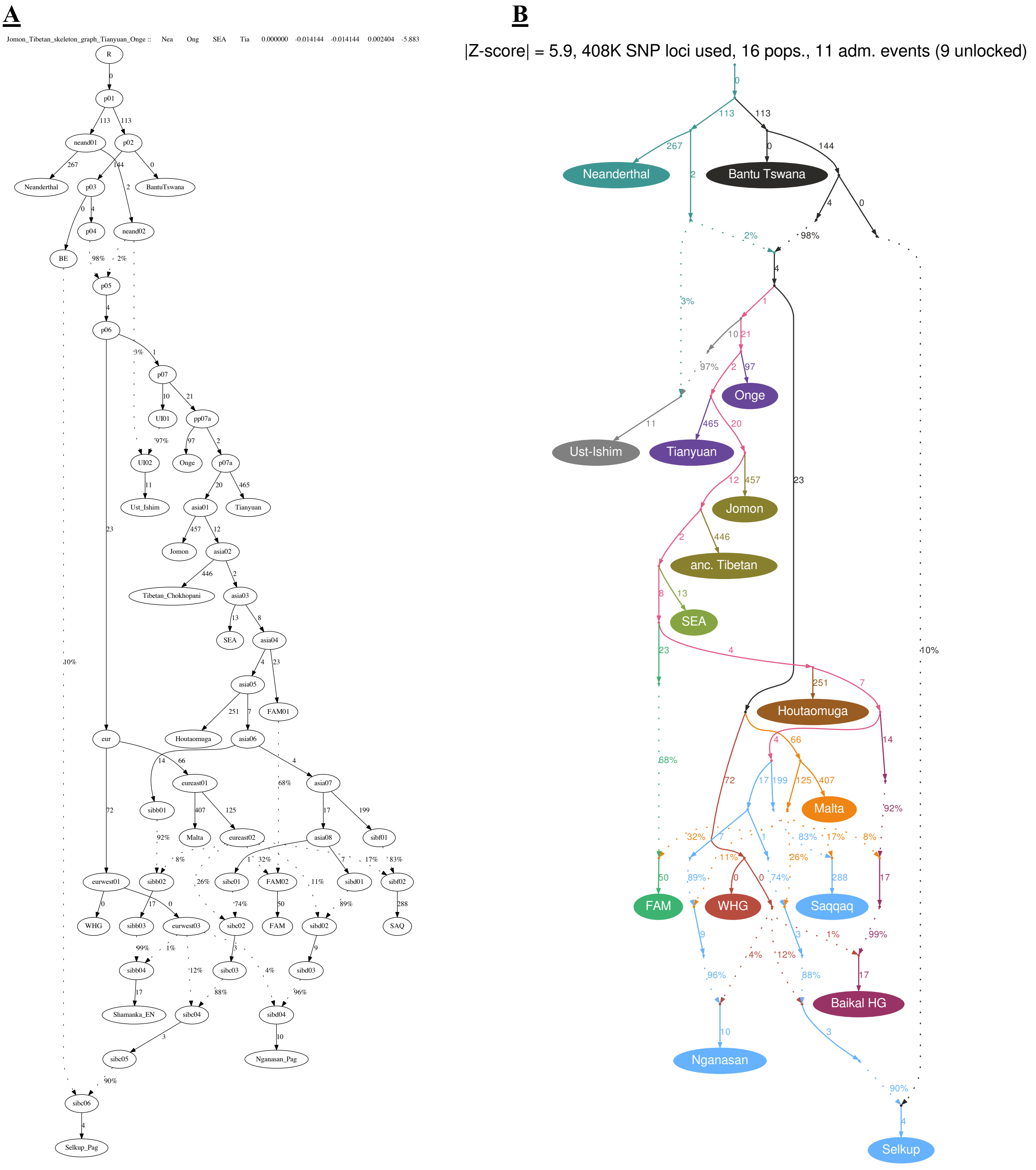


**Fig. S39.** A skeleton graph selected on the full dataset: its original representation (**A**) and its final clean representation (**B**).

6. Principal component analysis

We ran PCA analysis for 526 ancient and present-day individuals mapped as targets on the skeleton graphs, 51 ancient and present-day individuals comprising the groups on the 16-population skeleton graph (except for the Altai Neanderthal, Africans, and Onge), and 88 additional individuals (mostly present-day ones) from Southeast Asia, Europe and the Caucasus. The latter group of individuals was added to make the dataset composition more balanced. We used either the dataset without transitions or the full dataset, and tested three PCA protocols: 1) 10 PCs based on both ancient and present-day individuals, PLINK v. 1.90b6.10; 2) 10 PCs based on both ancient and present-day individuals, smartPCA v.16000, "shrink mode" turned on; 3) 10 PCs based on present-day individuals, ancient individuals projected, smartPCA v. 16000, "shrink mode" turned on. For the first setup we visualized PC1, 2, 3, and 4 using color and marker size (Fig. S40a,b). For the second setup we visualized PC1, 2, and 3 using color (Fig. S40c,d). For the third setup we visualized PC1, 2, 3, and 5 using color and marker size (Fig. S40e,f). Some PCs were not visualized because they separated just few individuals from the others. The distribution of individuals on the PC1 vs. PC2 plane was very much similar across all protocols, and here we focus on the results of the first setup (PLINK) since in this case one PC (PC4) revealed an important Mal'ta-related cline reported before (*30, 41*). The USR1 and Trail Creek American individuals were substantially shifted towards the Asian cline irrespective of the protocol used. This shift was observed before using MDS (*4*). Here we describe all individuals lying between the First American cluster and the Asian cline in 4 dimensions and interpret the PCA embedding in the light of the results on simulated data (Supplementary Text 2).

A prominent cline described before (*11*) extends from unadmixed Asians (Houtaomuga, Southeast Asians) to unadmixed First Americans (Fig. S40a,b). The Saqqaq Paleo-Inuit, related Chukotko-Kamchatkan speakers (*11*) and the Magadan Bronze Age individuals are slightly shifted towards Americans. This shift of the Chukotko-Kamchatkan speakers and the Magadan Bronze Age individuals is explained by the FAM ancestry detected in both of them (Data S10). It was shown before that FAM ancestry in the Chukotko-Kamchatkan speakers is derived from a bidirectional admixture with early Eskimos in Chukotka (*11*).

Further groups on the cline are Chukotkan Yupik (Eskimo-Aleut speakers) and Chukchi, ancient Chukotkan Eskimos and present-day Inuit, ancient Aleuts and an ancient Alaskan Athabaskan. Varying proportions of FAM ancestry in these groups were explained by a back and forth migration model across the Bering Strait (*11*). Another small cluster lying closer to FAM is composed of ancient and present-day Athabaskans, and of ~6000 cal yr BP Big Bar and 939 individuals from Canada. We found highly significant Paleo-Inuit admixture signals for all Athabaskans, but no significant signals were detected for Big Bar and 939 (Supplementary Text 8, Data S6). The Kennewick and ASO CK-13 individuals lie even closer to the main FAM cluster. Present-day Cree and Tsimshian (labelled "NAM") are shifted from the northern North American cluster (Athabaskans, Big Bar, 939, ASO, Kennewick) towards Europeans, in line with substantial European admixture in this group (Data S6). Four Mixtec and Mayan individuals are shifted from the main FAM cluster towards the European cluster, in line with substantial West Eurasian admixture detected (Data S6).

Looking at PC4 (6% of variance explained) we notice a distinct cline between American and Asians composed of some ~10000 cal yr BP individuals at the Lapa do Santo and Sumidouro sites (Brazil), USR1, Trail Creek, and Kolyma1. In the fourth dimension these individuals are shifted towards a Mal'ta-related cline (Fig. S40a,b). The Mal'ta-related cline is separated from the main European-Asian cline mainly in the 2nd and 4th dimensions and its extremes are the Mal'ta and Houtaomuga individuals. The following clusters lie on the cline: 1) West Siberian hunter-gatherers (*30*), Botai culture (*27, 41*), Dali EBA (*30*); 2) Okunevo culture (*41*); 3) Baikal Bronze Age (*41*); 4) Baikal Early Neolithic (*41*).

Importantly, the simulated USR1 individual was positioned between Americans and Asians in the absence of any Siberian gene flow (Fig. S32), and also shifted towards Mal'ta along PC4 (Data S2). Thus, these features of the USR1 and Kolyma1 individuals' PC positions cannot be interpreted as potential signals of gene flow. The same applies to PC positions of NAM individuals (939, Big Bar, ASO CK-13, Kennewick).

Ancestry proportions extracted from highest-ranking admixture graph models for both Asian and American targets (Supplementary Text 8) were mapped onto the PC1/PC2/PC4 space. PC4 was selected because it separates the groups on the Mal'ta-Siberian cline (Fig. S40). See results for selected ancestral lineages in Fig. S42 and raw data in Data S10 and S8-1. Non-random distributions of ancestry proportions in the PC space are observed for all components shown here. Mal'ta-related ancestry is common in Eurasia and America, as expected, and indeed reaches a maximum in a cluster formed by West Siberian hunter-gatherers, Botai culture individuals, and a Dali EBA individual. The First American ancestry was found only in individuals forming a planar cline in the PC1/PC2/PC4 space and occupying an intermediate position between the Asian and SAM clusters. This cline is subdivided into several clusters reported before (*11*): Chukotko-Kamchatkan speakers and Paleo-Inuit, Eskimo-Aleut speakers, Na-Dene speakers (represented by Athabaskans), other NAM groups (Fig. S42). Substantial Houtaomuga ancestry proportions were found along three clines (Fig. S42): 1) the cline extending towards WHG and present-day Europeans; 2) the cline extending towards Mal'ta; 3) the cline formed by the Kolyma1 and USR1 individuals.

#### 7. Applying the admixture graph mapping approach to the USR1 individual

*a. Introduction*

We showed on simulated data that a particular approach to inferring admixture, namely mapping a target on a skeleton admixture graph, is preferable as compared to protocols based on *qpWave*/*qpAdm* and *D*-statistics. A major advantage of graph mapping over these protocols is the possibility to compare alternative ancestry sources in a statistically sound way. Moreover, a model-comparison protocol resistant to overfitting was developed for admixture graphs (Supplementary Text 11). The graph mapping protocol also surpassed the *qpWave*/*qpAdm* protocol in sensitivity to gene flow and in the ability to resolve alternative ancestry sources (Supplementary Text 1). For these reasons we decided to reassess the original results on USR1 (4) taking advantage of the graph mapping approach and the skeleton graphs described in Supplementary Text 5.

*b. Mapping USR1 on the 14-population graph using the dataset without transitions*

First, we mapped USR1 on the simpler 14-population graph (Fig. S37) and used the dataset without transitions in order to mitigate at least some ancient DNA biases. A problem of this approach was that about 95,000 sites were left even for high-coverage targets, if the mathematically dubious "useallsnps: YES" option was avoided. In the next sub-section, we also show results for the more complex 16-population graph (Fig. S39) and the dataset including transitions (from ca. 377,000 to 408,000 sites available under the "useallsnps: NO" setting).

The USR1 individual (pseudohaploid and diploid versions) was mapped on the graph in the following way. By default, a gene flow from a Mal'ta-related lineage was added to account for the well-attested signal in USR1 and other Americans (*4, 10*). Only one two-way model was tested: the Asian source group for First Americans ($FAM_{EA}$) + Mal'ta. This model is consistent with Moreno-Mayar *et al.* (*4*) and was used as a baseline. The baseline model demonstrated worst *f*-statistic residuals between 3.9 and 4.8 SE, depending on settings and the USR1 genome version (Data S4a), but the distributions of *f*-statistic residuals conformed to the standard normal distribution (CCC = 0.99). Thus, the baseline model fitted the data according to the criterion developed in Supplementary Text 2f (CCC > 0.95). We tested 29 three-way models $FAM_{EA}$ + X + Mal'ta, where X was any possible edge in the anatomically modern human (AMH) sub-graph prior to West Eurasian (Mal'ta, WHG, basal Eurasian) gene flows into Siberians and Americans. Finally, we tested 406 four-way models $FAM_{EA}$ + X +Y + Mal'ta, where X and Y were any possible edges in the anatomically modern human (AMH) sub-graph prior to West Eurasian gene flows into Siberians and Americans. The results of this analysis are summarized in Data S4a.

In order to take account for systematic errors, we repeated the mapping protocol under eight setting combinations. First, West Eurasian gene flows into Siberians and FAM were either locked according to the values estimated under the skeleton model (Fig. S37) or left freely variable (the Neanderthal gene flows were always locked according to the values estimated under the skeleton graph). These setups correspond to those tested on simulated genetic data in Supplementary Text 2e. Second, the Houtaomuga group was composed of either two oldest Early Neolithic individuals (M45, M54A), or an Iron Age individual (M74B) having no West Eurasian admixture was added too (the same group compositions were tested in Supplementary Texts 5, 8-12). Third, either the "useallsnps: NO" or "useallsnps: YES" settings were used. The former setting was prioritized for target groups/individuals having more than 80,000 sites in common with the groups on the skeleton graph, and USR1 had from 81,903 to 94,853 such sites, depending on the Houtaomuga group composition and the USR1 genome version used. The "useallsnps: YES" setting was prioritized only for target groups/individuals having less than 80,000 sites in common with the groups on the skeleton graph. In the case of USR1, the "useallsnps: YES" setting resulted in 183,606 – 183,632 sites being used. In summary, three pairs of settings produced eight setting combinations.

The following thresholds optimized on simulated data were used. First, models within the same class (i.e. having identical number of parameters) were ranked by log-likelihood, and a threshold of 3 ln-units was used for defining a group of models indistinguishable from the best one ("top" models). This approach followed Flegontov *et al.* (*11*). Models classified as "top" under at least 3 out of 4 prioritized settings ("useallsnps: NO" in this case) were considered as "best" models in its class. A model with *n+1* gene flows was favored over a model with *n* gene flows if at least under **1 of 4** prioritized settings a **WR difference of >0.25 SE** intervals **OR** an **LL difference of >13 ln-units** was observed. According to the results on simulated data in the presence of ascertainment bias, we do not expect to see a WR difference >0.25 SE for >5% of datasets simulated under the null hypothesis, even if the main ancestry source proxy for USR1, i.e. FAM, is substantially mismodelled (Supplementary Text 2e, Table S9). If FAM is not mismodelled, we also do not expect to see an LL difference >13 units for >5% of datasets simulated under the null hypothesis, both in the presence and in the absence of ascertainment bias (Table S9). As an auxiliary criterion for model selection, we counted 0-length drift edges within the Asian clade which resulted in trifurcations. Trifurcations are believed to arise much more often as a result of topology misspecification than as a reflection of historical reality (*11*). We favored models having the lowest number of such 0-length edges under the prioritized setting combinations. Since 83% of correct 14-population graphs demonstrated WRs above 3 SE on simulated datasets of 95,000 sites with ascertainment bias and 57% of correct graphs demonstrated WRs above 3 SE in the absence of ascertainment bias, and since WR depended on dataset size in the presence of ascertainment bias (Fig. S30), the absolute value of the WR (Z-score) was not an important statistic for our model selection procedure. Similar mapping protocols were used throughout this study.

Among the three-way models for both USR1 genotype versions, only one model per target was classified as "best" (Data S4a): 65% $FAM_{EA}$ + 19% "basal Mal'ta" + 17% Mal'ta (the proportions are shown for the pseudohaploid target version). "Basal Mal'ta" is a source diverging deeper than the Mal'ta sister group contributing ancestry to all other Siberians and Americans, but not as deep as the WHG and Mal'ta split point. We note that the ratio of the Mal'ta and "basal Mal'ta" source proportions is unconstrained on our skeleton graph since just 3 and not 4 or more groups split off on the path between these ancestry sources (Lipson 2020 preprint). The model "$FAM_{EA}$ + Mal'ta + basal Mal'ta" was consistently classified as "top" under all 8 setting combinations, and as the only "top" model under 6 out of 8 setting combinations (these are results for the pseudohaploid target version). In other words, this model tended to be significantly different from all the other models of its class according to likelihood. LL improvement over the baseline model $FAM_{EA}$ + Mal'ta ranged from 11.7 to 20.6 log-units, and the WR improvement was 0.78-0.96 under "useallsnps: NO" settings. Most of those LL difference values and all WR difference values are significant by our standards.

Among the four-way models for both USR1 versions, five models were classified as "best" under the "useallsnps: NO" settings (Data S4a). Only one of those models demonstrated a significant improvement in WR over the 3-way class, i.e. an improvement exceeding 0.25 SE: "$FAM_{EA}$ + "basal Mal'ta" + Saqqaq" for the diploid USR1 version and "Houtaomuga + $FAM_{EA}$ +

"basal Mal'ta" for the pseudohaploid USR1 version. The WR improvements ranged from 0.28 to 0.57 SE under the prioritized settings (Data S4a). The latter model also had no "backbone" 0-length edges in the Asian clade under all 4 prioritized setting combinations and for both target versions. The LL improvement associated with the "$FAM_{EA}$ + basal Mal'ta + Saqqaq" model ranged from 5.7 to 6.5 units (for the diploid USR1 version under the prioritized settings). The LL improvement associated with the "Houtaomuga + $FAM_{EA}$ + basal Mal'ta" model ranged from 5.8 to 8.8 units (for the pseudohaploid USR1 version under the prioritized settings).

LL differences of this magnitude were often encountered in the absence of Siberian gene flow into the simulated USR1, especially if the main ancestry source proxy (FAM) was mismodelled, but WR differences of the observed magnitude were rare (Table S9). When 2- and 3-way models were compared, just 2.3% of graph mapping runs on simulated data with ascertainment bias showed a WR improvement above 0.25 SE, and the largest difference observed was 0.5 SE (95,000 sites, different *qpGraph* setups, 25 alternative 3-way models tested per run, 400 runs in total). If we consider the "$FAM_{EA}$ + Mal'ta + Houtaomuga" model only, just one of 400 runs (0.25%) showed a WR difference above 0.25 SE (0.47 SE). Moreover, if we consider the "$FAM_{EA}$ + Mal'ta + Houtaomuga" vs. "$FAM_{EA}$ + Mal'ta" comparison only and do not consider the "mismodelled FAM" scenarios, just 2.7% of 300 runs showed an LL difference above 5.7 units (the lowest difference observed on real data), and the largest difference observed on simulated data was 10 units.

When 3- and 4-way models were compared, 1.1% of runs on simulated data with ascertainment bias showed a WR improvement above 0.25 SE, and the largest difference observed was 1 SE (95,000 sites, different *qpGraph* setups and 5 simulated Houtaomuga admixture proportions in USR1, 300 alternative 4-way models tested per run, 2,000 runs in total). If we compared the "$FAM_{EA}$ + Mal'ta + Houtaomuga" and "$FAM_{EA}$ + Mal'ta + Houtaomuga+Y" models only, 3.7% of runs showed a WR difference above 0.25 SE (up to 1.7 SE).

If we consider all 3- and 4-way models and do not consider the "mismodelled FAM" scenarios, 14.3% of 1,500 runs showed an LL difference above 5.7 units (the lowest difference observed on real data), and the largest difference observed on simulated data was 19.9 units. If we compared the "$FAM_{EA}$ + Mal'ta + Houtaomuga" and "$FAM_{EA}$ + Mal'ta + Houtaomuga+Y" models only, 25.9% of runs showed an LL difference above 5.7 units, and the largest difference observed on simulated data was 38 units.

The results in the paragraphs above are based on simulated data with ascertainment bias. In the absence of ascertainment bias, 6.5% of graph mapping runs on simulated data showed a WR improvement above 0.25 SE, and the largest difference observed was 1.2 SE (95,000 sites, 400 runs in total). If we consider the "$FAM_{EA}$ + Mal'ta + Houtaomuga" model only, just three of 400 runs (0.75%) showed a WR difference above 0.25 SE, and the largest difference was 0.42 SE. When 2- and 3-way models were compared and the "mismodelled FAM" scenarios were not considered, 18.3% of runs on simulated data showed an LL improvement above 5.7 units, and the largest difference observed was 14.8 units (95,000 sites, different *qpGraph* setups, 300 runs in total). If we compared the "$FAM_{EA}$ + Mal'ta + Houtaomuga" and "$FAM_{EA}$ + Mal'ta + Houtaomuga+Y" models only, 4% of runs showed an LL difference above 5.7 units (up to 10.1 units).

We note that FAM mismodelling on simulated data was associated with high absolute WR values above ca. 10 SE (Fig. S30a), and we did not observe that on real data, where WR for the best model ranged from 2.6 to 2.8 SE (Data S4a). The worst-case results observed on simulated data (95,000 sites with ascertainment bias) in the absence of FAM mismodelling were the following: 1/ an LL improvement of 16.5 units and a WR improvement of 0.47 SE at the same time (all 2- and 3-way models compared, 300 runs); 2/ an LL improvement of 19.9 units and a WR improvement of 1 SE (all 3- and 4-way models compared, 1,500 runs). The worst-case results observed on simulated data with no ascertainment bias (95,000 sites) were the following: an LL improvement of 13.7 units and a WR improvement of 1.2 SE at the same time (all 2- and 3-way models compared, 300 runs). We conclude that it is not impossible, but unlikely to observe WR and LL improvements of the observed magnitude that would support a false signal of Siberian gene flow into a divergent American group such as USR1. A caveat is that the simulated data did not reproduce all the aspects of the real data (missing data, pseudohaploid samples, pairwise $F_{ST}$).

In our view, the evidence is clear enough for proclaiming only one "winning" model for the pseudohaploid USR1 version, namely: 35% Houtaomuga + 28% $FAM_{EA}$ + 20% "basal Mal'ta" + 17% Mal'ta. Standard deviations of the admixture proportions across the prioritized settings were <1%. The best models for each class are shown in Fig. S43.

Even given an informative phylogeny it is hard to determine the direction of a gene flow if an admixture proportion is below 10% (Fig. S6, Table S2). Under these conditions it is often possible to detect the existence of a gene flow relying on an LL difference cut-off of 10 units (Tables S1, S2), but impossible to determine its direction relying on an LL difference cut-off of 3 units (*11*). We tested this possibility and explored the effect of reversing the direction of the Houtaomuga-USR1 gene flow in the "USR1 = $FAM_{EA}$ + Houtaomuga + basal Mal'ta + Mal'ta" model. The model where the USR1 branch prior to the West Eurasian flows contributed to Houtaomuga was not favored according to an LL cut-off of 3 units and a WR cut-off of 0.1 SE. We note that since the models compared have the same number of parameters, the thresholds used here are different from those used above for comparing models differing by one admixture event. The LL difference between models with the opposite gene flow directions ranged from 3.6 to 5.4 units (4.4 units on average across 8 setting combinations), and the WR difference ranged from 0.38 to 0.49 SE (0.44 SE on average across 4 "useallsnps: NO" setting combinations). We also mapped the Houtaomuga group as unadmixed or as a 2-way mixture of any possible AMH sources on a skeleton graph including both the FAM group (with a Mal'ta-related gene flow) and USR1 (with both Mal'ta and "basal Mal'ta" flows). First, the admixed models for Houtaomuga were not supported: the LL difference for admixed vs. non-admixed Houtaomuga models ranged from 2.7 to 4.7 units (3.9 units on average across 8 setting combinations). Second, the highest-ranking admixed models for Houtaomuga had a likelihood significantly higher than the "standard model" with Houtaomuga gene flow into USR1 and the same number of parameters: the LL difference ranged from 3.2 to 4.4 units (3.7 units on average across 4 "useallsnps: NO" setting combinations).

We note that our results should be interpreted in a phylogenetic sense: when we mention, e.g., an "ARB" ancestry source, that means any source on the ARB branch after its split from the Asian stem. Considering that our method can probe relatively star-like phylogenies, the genetic drift shared by the actual ancestry source and the "anchor lineage" represented on a skeleton graph may be small. These arguments should be considered when interpreting our results from the archaeological perspective.

*c. Mapping USR1 on the 16-population graph using the dataset containing transitions*
To validate the results further, we mapped both USR1 versions on a more complex graph including the Tianyuan and Onge branches (Fig. S39) and used a larger dataset including transitions. Under the "useallsnps: NO" setting 377,481 – 407,556 sites were available, and we decided to skip the mathematically incorrect "useallsnps: YES" setting. We tested one model $FAM_{EA}$ + Mal'ta, 33 models $FAM_{EA}$ + X + Mal'ta, and 528 models $FAM_{EA}$ + X +Y + Mal'ta, all under 4 setting combinations. The WR of this more complex skeleton graph on a larger dataset was high (~5.8 SE), but it lies within the 95% CI for the basic skeleton graph fitted to simulated data with ascertainment bias (400,000 sites, no simulated Tianyuan and Onge branches, Fig. S30). Thus, the high WR does not necessarily suggest that the graph does not fit the data (Supplementary Text 2f). However, the baseline 2-way model for USR1 was mildly non-fitting (CCC = 0.93) according to the criterion developed in Supplementary Text 2f (CCC > 0.95). As we have shown above on simulated data (Fig. S6, Table S2), LL ranking remains precise even if WR of a skeleton graph is very large. In other words, fitting problems in another part of the graph do not necessarily influence mapping on a clade of interest. On the other hand, it is important that main ancestry sources for a target are not mismodelled (Table S9).

According to the results on simulated data with ascertainment bias (400,000 sites, the 14-population skeleton graph), we do not expect to see a WR difference >0.25 SE for >5% of datasets simulated under the null hypothesis, even if FAM is substantially mismodelled (Supplementary Text 2e, Table S9). If FAM is not mismodelled, we also do not expect to see an LL difference >13 units for >5% of datasets simulated under the null hypothesis (Table S9). Although the simulated graph was simpler than the graph tested on the full real dataset (it lacked the Tianyuan and Onge lineages), we extrapolated those results to the more complex graph used in this section.

In the case of the diploid USR1 version, one 3-way admixture model was supported ($FAM_{EA}$ + Mal'ta + "basal Mal'ta"), but no 4-way model was supported according to the LL difference cut-off of 13 units (Data S4b). The group of best 4-way models consisted of models "$FAM_{EA}$ + Mal'ta + Houtaomuga or Saqqaq + X", and the LL improvements associated with those models ranged from 8.7 to 13 units. In the case of the pseudohaploid USR1 version, only one 3-way model ($FAM_{EA}$ + Mal'ta + "basal Mal'ta") was classified as top, and LL improvement ranged from 20.1 to 22.6 units. Only one 4-way model (Houtaomuga + $FAM_{EA}$ + "basal Mal'ta" + Mal'ta) was classified as top under all 4 setting combinations, and LL improvement ranged from 13.6 to 17.7 units. The best models for each class are shown in Fig. S44.

LL differences of this magnitude were encountered in the absence of Siberian gene flow into the simulated USR1 only if the main ancestry source proxy (FAM) was severely mismodelled (Table S9). The simulated results discussed here are based on the 14-population skeleton graph and 400,000 sites with ascertainment bias. When 2- and 3-way models were compared and the "mismodelled FAM" scenarios were not considered, just 1.3% of 300 runs showed an LL difference above 13.6 units (the lowest difference observed for the pseudohaploid USR1 version), and the largest difference observed was 15.4 units. If we consider the models "$FAM_{EA}$ + Mal'ta + Houtaomuga" only, the largest LL difference observed was 9.5 units. We note that FAM mismodelling on simulated data was associated with high absolute WR values above ca. 15 SE (Fig. S30), and we did not observe that on real data, where WR for the best model ranged from 5.7 to 6.2 SE (Data S4b). WR differences observed in the absence of simulated gene flow were generally lower on 400K datasets as compared to 95K datasets (Table S9).

When we compared all 3- and 4-way models and did not consider the "mismodelled FAM" scenarios, 1.7% of 1,500 runs showed an LL difference above 13.6 units (the lowest difference observed on real pseudohaploid data), and the largest difference observed on simulated data was 21.3 units. If we compared the "$FAM_{EA}$ + Mal'ta + Houtaomuga" and "$FAM_{EA}$ + Mal'ta + Houtaomuga+Y" models only, 3.1% of 1,500 runs showed an LL difference above 13.6 units, and the largest difference observed on simulated data was 21.8 units.

The results in the paragraphs above are based on simulated data with ascertainment bias. In the absence of ascertainment bias, when 2- and 3-way models were compared and the "mismodelled FAM" scenarios were not considered, 1% of runs on simulated data showed an LL improvement above 13.6 units, and the largest difference observed was 20.2 units (400,000 sites, different *qpGraph* setups, 300 runs in total). If we compared the "$FAM_{EA}$ + Mal'ta + Houtaomuga" and "$FAM_{EA}$ + Mal'ta + Houtaomuga+Y" models only, no runs showed an LL difference above 13.6 units (the largest difference was 6.1 units).

Here we also explored the effect of reversing the direction of the Houtaomuga-USR1 gene flow in the "USR1 = $FAM_{EA}$ + Houtaomuga + basal Mal'ta + Mal'ta" model. The model where the USR1 branch prior to the West Eurasian flows contributed to Houtaomuga was not favored according to an LL cut-off of 3 units. The LL difference between models with the opposite gene flow directions ranged from 7.3 to 9.3 units (8.3 units on average across 4 setting combinations). We also mapped the Houtaomuga group as unadmixed or as a 2-way mixture of any possible AMH sources on a skeleton graph including both the FAM group and USR1. Admixed models for Houtaomuga were not supported according to the cut-off of 13 log-units: the LL difference ranged from 5.6 to 9.8 units (7.4 units on average across 4 setting combinations). Second, the highest-ranking admixed models for Houtaomuga had a likelihood significantly lower than the "standard model" with Houtaomuga gene flow into USR1 and the same number of parameters: the LL difference ranged from 7.4 to 9.4 units (8 units on average across 4 setting combinations).

*d. Assessing absolute fits of admixture graph models to the data*

Ranking alternative models and assessing if a model fits the data are different questions. As shown in Supplementary Texts 1j and 2f, using a WR threshold of 3 SE as a criterion of absolute fit is probably incorrect since 57% and 59% of correct models on 95,000 and 400,000 sites without ascertainment bias, respectively, had WR above 3 SE, and even higher WRs were reported for the same models in the presence of ascertainment bias (Fig. S30). Moreover, as discussed in Supplementary Text 2f, WR for a perfect model fitted to the data depends on the number of groups and, in the presence of ascertainment bias, on the number of sites. Thus, using a universal WR cutoff above 3 SE is also problematic. We suggest that a model with normally distributed residuals should be considered fitting the data. As we have shown on simulated data, the best way of assessing normality is calculating the concordance correlation coefficient that measures how close a quantile-quantile plot based on all *f*-statistic residuals is to the diagonal, i.e. to the standard normal distribution (Supplementary Texts 1j and 2f, Fig. S31).

The following models from sections *b* and *c* above were considered: the baseline model USR1 = $FAM_{EA}$ + Mal'ta; FAM $_{EA}$+ "basal Mal'ta" + Mal'ta; $FAM_{EA}$ + Houtaomuga + "basal Mal'ta" + Mal'ta. In the case of the dataset without transitions and the 14-population skeleton graph, distributions of all *f*-statistic residuals under "useallsnps: NO" or "useallsnps: YES" settings did not deviate from normality according to CCC: CCC remained above 0.98 for all model types, target versions, and settings. A CCC cut-off optimized on simulated data in Supplementary Text 2f is 0.95. Visual inspection of quantile-quantile plots for the pseudohaploid

USR1 version supports a conclusion that all the models listed above fit the data (Fig. S45a,b). Residual distributions for the same models on the more complex skeleton graph including Tianyuan and Onge and the larger dataset including transitions deviated from normality (Fig. S45c) due to interactions between these newly added groups and other Asian branches such as Southeast Asians. CCC was 0.93 for the 2-way model and between 0.940 and 0.946 for the 3- and 4-way models (both target versions, all settings). As shown in Supplementary Text 1, ranking models on an incorrect skeleton graph is not affected if the clade of interest is modelled correctly.

In summary, we have demonstrated that the baseline model for USR1 (*4*) fits the data reasonably well. However, that result does not imply that more complex models for USR1 are over-fitting the data. For instance, absolute fit metrics on simulated data failed to distinguish the correct ancestry source (Houtaomuga) and the other sources in the Siberian clade (Fig. S31). In other words, CCC is tolerant to sub-optimal ancestry sources (see also Figs. S19, S20).

*e. Exploring other mapping protocols and skeleton graphs for USR1*

To see if results are robust to details of the mapping protocol, we tested its further modifications. First, we mapped the pseudohaploid USR1 version on the 14-population graph (dataset without transitions), but this time only the $FAM_{EA}$ source was added by default, and all possible unadmixed ancestry sources on the graph were tested (not only those in the AMH clade). We tested 35 models $FAM_{EA}$ + X, 595 models $FAM_{EA}$ + X +Y, and 6,545 models $FAM_{EA}$ + X + Y + Z, all under 8 setting combinations. In addition, for one setting combination (West Eurasian gene flows into Siberians unlocked, "useallsnps: NO", 2 Houtaomuga ind.), 52,360 5-way models were tested (Data S4c). Here we counted trifurcations not only within the Asian clade, but on the whole graph. Among the 2-way models, only one model was classified as "best" (Data S4c): 65% $FAM_{EA}$ + 35% Mal'ta. Among the 3-way models, only one model was classified as "best" under both "useallsnps: NO" and "useallsnps: YES" settings: 37% $FAM_{EA}$ + 36% Mal'ta + 28% Houtaomuga. LL improvement over the previous model $FAM_{EA}$ + Mal'ta was non-significant according to our criteria under the "useallsnps: NO" setting combinations (LL difference ranged between 4.8 and 8.7 units), but WR improvement was significant under those settings (from 0.29 to 0.62 SE under 3 of 4 setting combinations). Remarkably, for this model there were no trifurcations on the whole graph under the "useallsnps: NO" settings. 4- and 5-way models were not supported by LL or WR differences above the selected thresholds. Thus, when the Mal'ta-like ancestry source in USR1 is not forced to be identical to that contributing to FAM and Siberians, the final model becomes simpler as no "basal Mal'ta" contribution is required. The simplification (all Mal'ta-like gene flows on the skeleton graph radiated from one node) was introduced on purpose to avoid an extremely complex model.

We repeated the same type of mapping protocol for the pseudohaploid USR1 version, the full dataset, and the 16-population graph including Tianyuan and Onge: $FAM_{EA}$ was the only default source, and 2-, 3, and 4-way models were tested (Data S4d). Among the 3-way models again only one model was classified as "best": 36% Houtaomuga + 35% Mal'ta + 29% $FAM_{EA}$. Log-likelihood improvement over the less complex model $FAM_{EA}$ + Mal'ta was significant (13.6 – 17.3 units) under all 4 setting combinations, and the WR profile was flat across different model classes. Like in the previous mapping round, 4-way models demonstrated no significant LL improvement (LL difference was up to 8.5 units, i.e. below the cut-off value of 13 units) (Data S4d).

Finally, we added the Yana Upper Paleolithic group (diploid or alternatively pseudohaploid data for two individuals *(10)*) to the 16-population skeleton graph including Tianyuan and Onge (on the full dataset including transitions). As expected, Yana was confidently mapped on the Mal'ta branch, but an additional gene flow from the Tianyuan branch got significant support (only one top model was supported by WR difference between 1.57 – 1.75 SE and LL difference between 70.5 and 78.6 log-units, depending on the settings and the type of Yana data used). This new skeleton including a pseudohaploid Yana group was again used for mapping USR1 (the pseudohaploid version). We tested 741 3-way models $FAM_{EA}$ + X + Y and 9,139 4-way models $FAM_{EA}$ + X + Y + Z on the full dataset under 4 setting combinations. In the 3-way class, only one model was classified as top: 37% Houtaomuga + 35% Mal'ta + 29% $FAM_{EA}$. 4-way models were not supported according to the LL difference cut-off of 13 units (LL difference was up to 9 units).

*f. Exploring the likelihood space of admixture proportions in USR1*

We took one of the best models found above for USR1, "Houtaomuga + $FAM_{EA}$ + "basal Mal'ta" + Mal'ta", and explored a three-dimensional likelihood space of admixture proportions. We varied the intensities of three gene flows (Houtaomuga, "basal Mal'ta", and Mal'ta) in 5% increments (from 0% to 100%) and then took 8,421 distinct combinations of resulting admixture proportions in the target. The analyses was repeated for 4 setting/dataset combinations: 1) the 14-population graph, no transitions, West Eurasian gene flows into Siberians and FAM locked according to the values estimated under the skeleton model; 2) the 14-population graph, no transitions, West Eurasian gene flows into Siberians not locked; 3) the 16-population graph, transitions included, West Eurasian gene flows into Siberians and FAM locked; 4) the 16-population graph, transitions included, West Eurasian gene flows into Siberians not locked. The "useallsnps: NO" setting was used, and two individuals comprised the Houtaomuga group. For each analysis run, models were classified into "top" (LL within 3 units of the best one), "questionable" (LL within 3 to 5 units from the best value), and "rejected" (LL >5 units different from the best one). Top models were assigned a rank of 1, questionable models were assigned a rank of 0.5, and rejected models were assigned a rank of 0. Summing up the ranks across 4 dataset/setting combinations, we got final model ranks (Data S4e). We considered models having a final rank >2 (under at least one setting combination these models were classified as "top"), and among those the inferred ancestry proportions in USR1 varied from 54.4% $FAM_{EA}$ + 9.6% Houtaomuga to 28.8% $FAM_{EA}$ + 35.2% Houtaomuga. The "basal Mal'ta" proportion varied from 16% to 21.3%, and the Mal'ta proportion from 15% to 20%. Lower Houtaomuga admixture proportions (3.2% and 6.4%) were encountered among models having a lower rank, but most of those were never classified as "top", in other words were significantly different from the best model under given settings. Only one of these latter models was classified as "top" under one setting combination: 57.4% $FAM_{EA}$ + 6.4% Houtaomuga. Thus, we conclude that ~6% is a lower bound on the Houtaomuga ancestry proportion in USR1, and just a small fraction of the likelihood space is occupied by acceptable models (see a 3D visualization in Data S5).

*g. Does the First American ancestor of USR1 form the deepest branch in the American clade?*

Finally, we attempted to check if the FAM source population of USR1 indeed represents a branch of the American clade splitting deeper than the well-documented South/North American and northern North American FAM clades, as shown by Moreno-Mayar *et al.* (*4*). For this purpose, we removed the original FAM group and grafted onto the 16-population skeleton graph an American clade composed of 4 individuals: USR1; an ancient Athabaskan (I5321, 700 cal yr BP) as a representative of the northern North American clade; Anzick (Clovis, 12500 cal yr BP) and an ancient Peruvian IL7 (1800 cal yr BP) as representatives of the North/South American clade. In addition to the Mal'ta gene flow, other supported gene flows were added to the USR1 lineage (from the Houtaomuga and "basal Malta" branches), to the ancient Athabaskan I5321 (from the Saqqaq branch), and to Clovis (from the Saqqaq and "basal Malta" branches). For justification of these models see Supplementary Text 8. To explore the parameter space thoroughly, we tested 24 setting combinations for each American clade topology: 1) the West Eurasian gene flows into Siberians were either locked according to the values inferred under the skeleton graph or unlocked; 2) any gene flows into the four American individuals were either locked or unlocked; 3) transitions were excluded or kept; 4) "useallsnps: NO" or "YES" settings were used; 5) the Houtaomuga group was composed of two or three individuals as detailed above. The "useallsnps: YES" setting was not used in combination with the larger dataset containing transitions, but all other setting combinations were tested. All possible branching orders of the $FAM_{EA}$ sources were tested, i.e. 15 alternative topologies. The following topology classes were defined: 1/ two sister clades ((A, B), (C, D)); 2/ the $FAM_{EA}$ source population of USR1 occupying an apical (A, (B, (USR1, C))), intermediate (A, (USR1, (B, C))), or a basal position (USR1, (A, (B, C))). Under each setting combination, models were classified into "top" (LL within 3 units of the best one), "questionable" (LL within 3 to 5 units from the best value), and "rejected" (LL >5 units different from the best one), and the number of trifurcations in the FAM clade was counted.

The topology (I5321, (USR1, (Anzick, IL7))) was classified as "top" under 20 out of 24 setting combinations (Data S4f). The topology ((I5321, USR1), (Anzick, IL7)) emerged as a second-best model and was classified as "top" under 16 setting combinations. A topology including the basal USR1 position (USR1, (I5321, (Anzick, IL7))) was classified as "top" under 12 setting combinations, and two other topologies with the basal USR1 position never emerged in the "top" class. Remarkably, the only acceptable basal USR1 topology (USR1, (I5321, (Anzick, IL7))) yielded trifurcations in the FAM clade under 22 of 24 setting combinations. In contrast, the topology ((I5321, USR1), (Anzick, IL7)) demonstrated by far the lowest number of trifurcations, under 5 of 24 setting combinations.

Since the complex model for Anzick was not supported on the realigned version of this genome (Data S6), we also repeated the procedure described above taking a simple $FAM_{EA}$ + Mal'ta model for Anzick. The topology (I5321, (USR1, (Anzick, IL7))) was classified as "top" under 18 out of 24 setting combinations (Data S4g). The topology ((I5321, USR1), (Anzick, IL7)) was again a second-best model and was classified as "top" under 16 setting combinations. A topology including the basal USR1 position (USR1, (I5321, (Anzick, IL7))) was classified as "top" under 10 setting combinations, and two other topologies with the basal USR1 position again never emerged in the "top" class. The only acceptable basal USR1 topology (USR1, (I5321, (Anzick, IL7))) yielded trifurcations in the FAM clade under 21 of 24 setting combinations. In contrast, the topology ((I5321, USR1), (Anzick, IL7)) demonstrated no trifurcations under any setting combination tested. The second-best model according to this criterion was much worse, demonstrating trifurcations under 12 setting combinations.

We conclude that forcing the basal position of the $FAM_{EA}$ source in USR1 almost invariably leads to model rejection or to trifurcations, suggesting that the basal position reported before for USR1 (*4*) is an artefact resulting from the unmodelled Siberian gene flow in USR1. Consequently, the very early split date reported for USR1 in the same publication is probably an artefact too. Since the topology ((I5321, USR1), (Anzick, IL7)) was not only the second most supported topology, but also was much less prone to trifurcations than the other topologies, we consider it a "winner" in this model competition (Fig. S46). This topology implies that the FAM source in USR1 belongs to the northern North American clade, which is not surprising given that the burial is located in Alaska.

#### 8. Asian gene flow signals in Native Americans

We explored other ancient and present-day Americans using the graph mapping approach applied to the USR1 individual. We mapped 84 ancient individuals and 13 present-day groups on the 14-population graph (the dataset without transitions, 8 setting combinations, "$FAM_{EA}$ + Mal'ta", "$FAM_{EA}$ + X + Mal'ta", "$FAM_{EA}$ + X + Y + Mal'ta" models) or alternatively on the 16-population graph (the full dataset, 4 setting combinations, "$FAM_{EA}$ + Mal'ta", "$FAM_{EA}$ + X + Mal'ta", "$FAM_{EA}$ + X + Y + Mal'ta" models). All ancient Americans having at least 30% of the 1240K panel (*25*) autosomal sites covered and published prior to 2019 were included into this analysis, and the data were obtained from the following source: https://reich.hms.harvard.edu/downloadable-genotypes-present-day-and-ancient-dna-data-compiled-published-papers. Only one dataset published in 2019 was included (*11*). Most ancient individuals analyzed here were published by Posth *et al.* (*14*) (33 ind.), Moreno-Mayar *et al.* (*13*) (14 ind.), and Scheib *et al.* (*18*) (13 ind.)

In the case of the full dataset where only the "useallsnps: NO" setting was used, site counts ranged from 134 to 412 thousand (309 thousand on average). In the case of the dataset without transitions and the "useallsnps: NO" setting, site counts ranged from 32 to 96 thousand (68 thousand on average). When the "useallsnps: YES" setting was used, 184 thousand sites were available. In about a half of cases (see also section 9) winning models were the same across all settings. The "useallsnps" setting influenced the results in many remaining cases, and it had clearly a stronger influence than the other settings tested. For this reason, we decided to split targets into two groups according to their coverage. For targets having less than 80,000 sites available under the "useallsnps: NO" setting, the interpretation was based on that setting only. For the other targets, the interpretation was based on the "useallsnps: YES" setting. Model ranking and model selection exactly followed the protocol described for USR1 above. A model with *n+1* gene flows was favored over a model with *n* gene flows if at least under 1 of 4 settings a WR difference of >0.25 SE intervals OR an LL difference of >13 log-units was observed.

Best models selected for Americans are summarized in Data S6 (all models tested are not shown). Results for 70 targets out of 101 (pseudohaploid USR1 + 100 other targets) were stable across both dataset/skeleton graph combinations (Data S6), i.e. the sets of winning models were identical. We were able to detect a number of known gene flows, and we accomplished that in an automated and unsupervised way. First, Paleo-Inuit admixture well-documented in Eskimo-Aleut speakers (*4, 11, 17*) and in Na-Dene speakers (*11, 17*) was revealed. Models "$FAM_{EA}$ + Saqqaq" were supported by an LL difference of >13 units and were unique solutions, i.e. significantly different from the other models in the same class (LL difference >3 units). For example, the Saqqaq gene flow into an ancient Aleut I0719.SG (*11*) (~85,000 sites under the "useallsnps: NO" setting on the dataset without transitions) was supported by a WR difference between 5 and 5.3 SE and an LL difference between 79 and 93 units (across 8 setting combinations, the 14-population graph). Adding yet another gene flow to that individual was not supported under any setting combination. In the case of an ancient Athabaskan I5321 (*11*) dated to ~700 cal yr BP (~85,000 sites under the "useallsnps: NO" setting on the dataset without transitions), the Saqqaq gene flow was also supported strongly: by a WR difference between 0.7 and 0.9 SE and an LL difference between 12 and 15 units (across 8 setting combinations, the 14-population graph). The Saqqaq (Paleo-Inuit) admixture proportion averaged across four "useallsnps: NO" setting combinations was 22%, in good agreement with a previous study (*11*).

We also found a strong signal corresponding to a well-known instance of colonial European admixture in present-day Aleuts (*33*). The best model for present-day Aleuts was 20% $FAM_{\underline{EA}}$ + 22% Saqqaq + 40% "basal WHG" + 18% Mal'ta (the 14-population graph, dataset without transitions, Data S6). Remarkably, both the $FAM_{\underline{EA}}$/Saqqaq ratio and the European admixture proportion manifested here as a WHG-related source were in perfect agreement with previous studies (*11, 33*). A very similar result was obtained on the full dataset (Data S6). Other known signals of West Eurasian gene flow (*11*) were found in the following present-day groups: Tsimshians and Cree (three individuals merged under a "NAM" label), Mixtec, and Mayans. In Mayans and Mixtec an African signal was also found, and it was previously reported too (*11*).

The "population Y" signal is manifested as "attraction" of a subset of ancient and present-day native Amazonians to the Onge/Papuan/Australian clade (*13, 34*), or alternatively to Tianyuan (*35*). This signal remains controversial (*13, 14*). We detected the Tianyuan component in 1 out of 6 ancient Brazilians from Lapa do Santo (9000-10000 cal yr BP) analyzed here (Data S6). These individuals were previously analyzed by Posth *et al.* (*14*), and no signal was found. The Tianyuan admixture proportion inferred by us was low, 6%, but the signal was above the selected LL threshold and specific for the Tianyuan branch. The Tianyuan-related component in an individual CP22 (~246,000 sites under the "useallsnps: NO" setting on the full dataset) was supported by an LL difference between 23 and 27 units (across 4 "useallsnps: NO" setting combinations). That model contained no trifurcations in the Asian clade and was a unique solution in its model class. Adding yet another gene flow to that individual was not supported under any setting combination (LL difference <13 units).

Three of four ancient Brazilians from Sumidouro (~10000 cal yr BP) analyzed here showed evidence of gene flows from deeply diverging sources, but several sources were almost equally supported, and "basal Mal'ta" was one of them in the case of the Sumidouro6 and Sumidouro7 individuals (Data S6). In contrast, for the highest-coverage individual Sumidouro5 (~408,000 sites under the "useallsnps: NO" setting on the full dataset) only the ancestry sources on the Onge and Tianyuan branches and on the adjacent stem edges were among the winning models (Data S6). The Onge source was nominally the best one and was supported by an LL difference between 16 and 18 units (across 4 "useallsnps: NO" setting combinations). That model contained no trifurcations in the Asian clade. Adding yet another gene flow to that individual was not supported under any setting combination (LL difference <13 units). The Onge- or Tianyuan-related admixture proportions averaged across 4 "useallsnps: NO" setting combinations were 7% or 8% (Data S6). The cases described above serve as positive controls, and our results suggests that the graph mapping approach can detect signals that remained elusive for simpler methods like single *f*- and *D*-statistics (*15, 33*).

We also found previously unreported admixture signals for two American individuals apart from USR1. Here we focus on most stringent results, i.e. on models that were declared as "winners" for both dataset/skeleton graph combinations. Only the Tianyuan and Onge sources were treated differently since they were included into one skeleton graph only. This high-confidence subset of extra-American ancestry sources is shown on a map in Fig. S47. Tianyuan-like ancestry was found at a low level (2%) in an ancient individual from Belize dated at 9300 cal yr BP (*14*) (Data S6). The Tianyuan-related component in an individual I3443 (~193,000 sites under the "useallsnps: NO" setting on the full dataset) was supported by an LL difference between 15 and 17 units (across 4 "useallsnps: NO" setting combinations). That model contained no trifurcations in the Asian clade and was a unique solution in its model class. Adding yet another gene flow to that individual was not supported under any setting combination (LL difference <13 units). Tianyuan forms the deepest known lineage in the East Asian clade (our results and Gakuhari *et al*. (*29*)) predating the American-Siberian split. Therefore, it is not unlikely that the first American settlement wave was not homogeneous genetically and included groups differentially related to the Tianyuan lineage.

Paleo-Inuit ancestry was found at a low level (5%) in an ancient individual from California dated at ca. 740 cal yr BP (*18*) (Data S6). This component in an individual SM-02 (~352,000 sites under the "useallsnps: NO" setting on the full dataset) was supported by an LL difference between 18.5 and 19.2 units (across 4 "useallsnps: NO" setting combinations). That model contained no trifurcations in the Asian clade and was a unique solution in its model class. Adding yet another gene flow to that individual was not supported under any setting combination (LL difference <13 units). Since this individual postdates the arrival of Southern Athabaskans into the southwestern USA (*11*), this signal may be explained by diffusion of Paleo-Inuit ancestry typical for Athabaskan speakers across the region.

Remarkably, no significant signals were found in an individual from the Trail Creek Cave in Alaska dated at 9000 cal yr BP (*13*). This low-coverage individual was shown to be related to USR1 (*4*). This result can be attributed to the low number of sites available in the full dataset (~145,000 sites).

#### 9. Applying the admixture graph mapping approach to the Kolyma1 and Ust'-Kyakhta individuals

A high-coverage genome (14×) of an individual from the Duvanny Yar site at the Kolyma River (Kolyma1) is another sample potentially very important for studying the settlement of America and the potential Siberian-American gene flow due to his geographic location in West Beringia and his date of 9800 cal yr BP (*10*). This individual was shown to be the closest relative of First Americans found in Asia to date, was interpreted as a typical representative of Paleo-Siberians whose ancestry today survives in an admixed form in Paleo-Inuit, Koryaks, Yukaghirs, Ket, some Uralic speakers, and was demonstrated to form a clade with USR1 or FAM that excludes Neo-Siberians (mostly Turkic and Tungusic speakers) (*10*).

Later data for an older individual (from the Ust-Kyakhta-3 site (UKY) just south to the Lake Baikal in Buryatia, Russia) closely related to Kolyma1 was published: targeted enrichment data on the 1240K SNP panel (*20*). The Ust'-Kyakhta individual was dated to 14050–13770 cal yr BP. Both Kolyma1 and Ust'-Kyakhta individuals were modelled as mixtures between independent basal FAM lineages (having Mal'ta ancestry) and Siberians (*20*).

We mapped the Kolyma1 individual (*10*), alternatively the diploid and pseudohaploid versions, on the basic skeleton graph and used the dataset without transitions. Kolyma1 was mapped as X (any AMH source) + Mal'ta (30 models under 8 setting combinations), then as X + Y + Mal'ta (380 models), and as X + Y + Z + Mal'ta (3,895 models). "Useallsnps: NO" settings were prioritized since the number of sites available under this setting exceeded 90 thousand. For both Kolyma1 versions, there were three 2-way models significantly different from the other models: Kolyma1 was mapped on the Houtaomuga branch or on the edges before or after the Houtaomuga split point (Data S7a). For both Kolyma1 versions, there were four or five 3-way models significantly different from the other 3-way models and also showing a significant improvement as compared to the best 2-way model. LL difference was >13 units and WR difference was >0.58 SE for the diploid version under prioritized settings and >10 units and >0.6 SE for the pseudohaploid version under prioritized settings. LL difference exceeding 13 units and WR difference exceeding 0.25 SE is significant according to our criteria. 4-way models were not supported by significant LL or WR differences. There were four "winning" models common for both genome versions. All of them included the Saqqaq branch as a minor ancestry source (3-15%), and $FAM_{EA}$, or the Houtaomuga branch, or the edges before and after the Houtaomuga split point (Data S7a). The proportion of the latter source ranged from 59% to 69%.

Reversing the direction of the Saqqaq => Kolyma1 gene flow in the "Houtaomuga + Saqqaq + Mal'ta" model resulted in negligible LL and WR differences (up to 1.2 units and 0.01 SE across all 8 setting combinations and both genome versions). The Kolyma1 admixture proportion in Saqqaq ranged from 3% to 7% (5% on average). Looking from the historical perspective, we believe that the gene flow direction from Kolyma1 to Saqqaq branch is more likely. The former individual is more than two times older than the latter (9800 vs. 3900 cal yr BP), and we doubt that the Paleo-Inuit branch diverged from the Siberian stem much more than 10 thousand years ago (see a discussion of Paleo-Inuit origins and their possible predecessors among Siberian archaeological cultures in Flegontov *et al.* (*11*)).

This analysis was repeated for the full dataset and the more complex skeleton graph including Tianyuan and Onge (Data S7b). For the diploid genome version, there were three 4-way models significantly different from the other models of the same class: $FAM_{EA}$ + the Houtaomuga branch or the edges before or after the Houtaomuga split point + Saqqaq + Mal'ta.

These models were supported by LL improvements from 10 to 13 units under 2 out of 4 setting combinations. Just one model had no trifurcations in the Asian clade under all settings, and that was 46% $FAM_{EA}$ + 18% Houtaomuga + 10% Saqqaq + 26% Mal'ta. For the pseudohaploid genome version, the same model was among 4 "winning" models: 45% $FAM_{EA}$ + 25% Houtaomuga + 5% Saqqaq + 25% Mal'ta (Data S7b). This model was supported by LL difference of up to 13.7 units and had no trifurcations in the Asian clade under all settings. Reversing the direction of Saqqaq-Kolyma1 gene flow in the "$FAM_{EA}$ + Houtaomuga + Saqqaq + Mal'ta" model resulted in negligible LL difference (up to 1.9 units across all 4 setting combinations and both genome versions). The Kolyma1 admixture proportion in Saqqaq ranged from 4% to 8% (6% on average). According to this collection of models, in Kolyma1 the Houtaomuga ancestry proportion ranged from 37% to 55% (45% on average across 4 settings and both genome versions), and the $FAM_{EA}$ ancestry proportion ranged from 19% to 38% (29% on average).

We mapped the Ust'-Kyakhta individual (a pseudohaploid version) on both skeleton graph versions in the same manner as the Kolyma1 individual. "Useallsnps: NO" settings were prioritized. First, we consider mapping on the 14-population graph and the transition-free dataset (Data S7a). There were three 2-way models significantly different from the other models: Ust'-Kyakhta was mapped on the $FAM_{EA}$ branch or on the edges before or after the $FAM_{EA}$ split point (Data S7a). There was only one 3-way model significantly different from the other 3-way models and also showing a significant LL and WR improvement as compared to the best 2-way model: 76% $FAM_{EA}$ + 20% Mal'ta + 4% Saqqaq. LL difference was between 14 and 15.5 units and WR difference was >0.58 SE under prioritized settings. Two 4-way models were supported by significant WR differences between 0.27 and 0.46 SE under prioritized settings: "$FAM_{EA}$ + Saqqaq + basal Mal'ta + Mal'ta" and "$FAM_{EA}$ + Baikal HG + Saqqaq + Mal'ta". However, LL differences were not significant between the best 3- and 4-way models under prioritized settings.

Reversing the direction of the Saqqaq => Ust'-Kyakhta gene flow in the "$FAM_{EA}$ + Mal'ta + Saqqaq" model resulted in negligible LL and WR differences (up to 0.6 units and 0.1 SE across all 8 setting combinations). The Ust'-Kyakhta admixture proportion in Saqqaq ranged from 4% to 5%. As in the case of Kolyma1, we believe that the gene flow direction from Ust'-Kyakhta to Saqqaq branch is more likely because of the great difference in the dates of these individuals. On the other hand, reversing the direction of the $FAM_{EA}$ => Ust'-Kyakhta gene flow in the "$FAM_{EA}$ + Mal'ta + Saqqaq" model resulted in LL differences ca. 60 units and WR differences above 1.4 SE. Thus, the reversed gene flow direction was strongly rejected in this case.

This analysis was repeated for the full dataset and the 16-population skeleton graph (Data S7b). Although the same models "$FAM_{EA}$ + Mal'ta", "$FAM_{EA}$ + Mal'ta + Saqqaq" and "$FAM_{EA}$ + Saqqaq + basal Mal'ta + Mal'ta" were the winners in their classes, there were no significant LL or WR differences between the classes. The simplest model "77% $FAM_{EA}$ + 23% Mal'ta" was the sole winner in its class under two of four setting combinations (Data S7b). Reversing the direction of the Saqqaq => Ust'-Kyakhta gene flow in the "$FAM_{EA}$ + Mal'ta + Saqqaq" model resulted in negligible LL differences (up to 0.5 units across 4 setting combinations). Reversing the direction of the $FAM_{EA}$ => Ust'-Kyakhta gene flow in the "$FAM_{EA}$ + Mal'ta + Saqqaq" model resulted in LL differences from 72 to 77 units and WR differences up to 1.0 SE. Thus, the reversed gene flow direction was strongly rejected.

Next, we co-analyzed the five key members of the proto-American clade on one graph, extending the analysis in Supplementary Text 7, section g. For this purpose, we removed the original FAM group and grafted onto the 16-population skeleton graph an American clade composed of 5 individuals: Ust'-Kyakhta, the pseudohaploid Kolyma version; the pseudohaploid USR1 version; an ancient Athabaskan (I5321, 700 cal yr BP) as a representative of the NNA clade; an ancient Peruvian (IL7, 1800 cal yr BP) as a representative of the SNA clade. In addition to the Mal'ta gene flow, other supported gene flows were added to the Kolyma1 lineage (from the Houtaomuga branch), USR1 lineage (from the Houtaomuga and "basal Malta" branches), and to the ancient Athabaskan I5321 (from the Saqqaq branch). For simplicity, the Ust'-Kyakhta/Kolyma-related gene flow to the Saqqaq branch was omitted, and the Houtaomuga-related ancestry sources in USR1 and Kolyma were derived from the same node. We tested one setting combination: 1) the West Eurasian gene flows into Siberians and Americans were unlocked; 2) transitions were kept; 3) "useallsnps: NO"; 4) the Houtaomuga group was composed of three individuals as detailed above. All possible branching orders of the FAM sources were tested, i.e. 105 alternative topologies.

Twenty-five best topologies for the American clade (ranked by LL) are shown in Table S19. There is a group of 21 models that is separated by a wide LL gap from the other models, >120 ln-units. These models have nearly identical WRs and all include the Ust'-Kyakhta and/or Kolyma individuals as the deepest branch in the clade we term proto-American. Just two of those 21 topologies have no trifurcations in the proto-American clade, and one of those is indistinguishable from the best LL-ranked topology according to both LL and WR: ((Kolyma, Ust'-Kyakhta), (SNA, (NNA, USR1))). This winning topology is in perfect agreement with the results reported in Supplementary Text 7, section g, i.e. with the placement of the FAM source of USR1 in the NNA clade. The winning topology is visualized in a simplified form in Fig. S48.

In summary, we tentatively support main conclusions by Sikora *et al.* (*10*). First, the Kolyma1 and/or Ust'-Kyakhta lineages indeed contributed ancestry to the whole Paleo-Inuit-related clade including Chukotko-Kamchatkan speakers, albeit at a low level according to our results. Second, Kolyma1 is partially descended from the FAM branch. If Kolyma1 indeed is a $FAM_{EA}$ + Houtaomuga mixture (not counting the non-debatable Mal'ta-like contribution), then this individual might represent a remnant of the migration that gave rise to USR1. Another conclusion by Sikora *et al.* (*10*) we do not support: according to our results Kolyma1 is closer to the Amur River Basin population cluster and Evens or Evenks (i.e. to Neo-Siberians) who have substantial proportions of the Houtaomuga component, and not to Paleo-Siberians having the Saqqaq, Ket/Selkup and Nganasan components (see Data S10).

We support the result by Yu *et al.* (*20*) that the Ust'-Kyakhta individual descends from the FAM lineage but question another result: Siberian admixture in Ust'-Kyakhta. According to our graph mapping results, Ust'-Kyakhta is best interpreted as an unadmixed proto-American lineage that contributed ancestry to some later Siberians (Paleo-Inuit). Although the direction of the Ust'-Kyakhta/Saqqaq gene flow was impossible to resolve, we argue based on parsimony that the Ust'-Kyakhta=>Saqqaq direction is more likely. Thus, the Kolyma1 and Ust'-Kyakhta individuals are related, but distinct, since there is support for the ARB-related admixture in Kolyma, but not in Ust'-Kyakhta.

Yu *et al.* (*20*) presented the following admixture graph topology for the deepest FAM lineages: (Ust'-Kyakhta [admixed with Siberians], (Kolyma [admixed], (USR1 [unadmixed], (NNA, SNA)))). In contrast to our results, USR1 was modelled as an unadmixed lineage in that study. An exhaustive exploration of all possible topologies for the FAM source populations of these five groups revealed three topologies indistinguishable according to LL and WR (Table S19), but only one of those had not trifurcations in the American clade. A conservative interpretation of our results is that either of these two topologies is correct: 1/ ((Ust'-Kyakhta [unadmixed], Kolyma [admixed]), (SNA, (USR1 [admixed], NNA))); 2/ (Ust'-Kyakhta [unadmixed], (Kolyma [admixed]), (SNA, (USR1 [admixed], NNA)))). The simplest interpretation of this topology is that USR1 is an NNA sub-group admixed with Siberians, and Kolyma and Ust'-Kyakhta represent groups that share at least a part of the standstill history with Native Americans but have never left Asia. The finding of Ust'-Kyakhta deep within Asia suggests that the standstill has actually happened far from Beringia.

10. *qpAdm* analysis of the USR1 individual

We ran *qpAdm* as suggested by Harney *et al.* (*49*), using a "rotating" set of outgroups and testing initially the least complex and then more complex models for the target. We ran *qpAdm* on the dataset without transitions and on the full dataset, and used the following "rotating" set of outgroups: the Altai Neanderthal individual, present-day Bantu Tswana, the Ust'-Ishim individual, the Mal'ta individual, a composite group of West European hunter-gatherers (WHG), the Jomon individual, the Tibetan Chokhopani individual, a composite group of present-day Southeast Asians (SEA), a composite group of present-day First Americans (FAM), the Shamanka early Neolithic (EN) group, the Saqqaq individual, present-day Selkups, present-day Nganasans, two or three oldest Houtaomuga individuals, and the USR1 individual. The group composition matched that used for *qpGraph* analyses above (Supplementary Text 7). We first tested all possible pairs of groups using *qpWave*, and then tested all possible 2- and 3-way admixed models for USR1, Houtaomuga, and FAM. Each test was repeated for pseudo-haploid and diploid USR1 versions (*4*), and for two alternative compositions of the Houtaomuga group (two or three individuals). Thus, four alternative setups were tested in parallel. The "useallsnps NO" setting was used for all tests.

Initially we tested the cladality of USR1 and various FAM groups using *qpWave*: a group of non-Amazonian present-day SNA individuals (Aymara, Mixe, Mixtec, Piapoco, Quechua, Yukpa, Zapotec) (*11*) used for most admixture graph analyses in this study, 45 alternative groups composed of ancient SNA and NNA individuals published mostly by Posth *et al.* (*14*), Moreno-Mayar *et al.* (*13*), and Scheib *et al.* (*18*), and 11 alternative present-day SNA groups partially overlapping with the composite SNA group. The cladality of First Americans and USR1 was rejected for 29 to 33 of 57 groups, depending on settings (*p*-values from $6\times10^{-7}$ to 0.01, Data S8a). The FP rate of this simple *qpWave* test on simulated data was 7% in the presence of ascertainment bias (*p*-value cut-off = 0.01, 30% Malta ancestry in FAM and USR1, 100 simulated datasets, 400,000 randomly sampled sites with MAF >5%) and <2% in the absence of ascertainment bias (Materials and Methods).

To test if the FAM or USR1 group is responsible for the cladality violation, we tested all possible 2- and 3-way admixed models for USR1, the composite FAM group of 13 individuals, and the Houtaomuga group as targets. All group pairs were inconsistent with being a clade with respect to the outgroups, given a *p*-value cut-off of 0.01 (see the pairwise *qpWave* results in Data S8b). Among 273 2-way models tested using *qpAdm* for the USR1, FAM, and Houtaomuga targets, no model satisfied the following conditions (following Narasimhan *et al.* (*30*)): *p*-value > 0.01 and inferred admixture proportions ± 2 SE lie between 0 and 1 for at least one setup. Among 1,092 3-way models tested using *qpAdm* for the same targets, just one model satisfied these conditions: USR1 = 45±20% FAM + 35±13% Houtaomuga + 20±7% Mal'ta. The conditions were satisfied when the diploid version of the USR1 genome was used, and the *p*-value for the pseudohaploid USR1 version was slightly below 0.01 (Data S8d). The condition "inferred admixture proportions ± 2 SE lie between 0 and 1" was satisfied under all four setups. Here we focus on results for the full dataset, which demonstrated better resolution due to the higher number of sites used (Data S8b-d). Results for the transition-free dataset are also presented in the same spreadsheet (Data S8e-g), and they did not contradict the results for the full dataset.

The FP rate of the whole *qpWave/qpAdm* procedure (cladality testing and further tests of 2- and 3-way models) on simulated data was 5% in the presence of ascertainment bias (*p*-value cut-off = 0.01, 30% Malta ancestry in FAM and USR1, 100 simulated datasets, 400,000 randomly sampled sites with MAF >5%) and <2% in the absence of ascertainment bias (Materials and Methods). In other words, for 5 out of 100 datasets simulated under the null hypothesis i/ the cladality of FAM and USR1 was rejected and ii/ only one of three alternative targets (FAM, USR1, Houtaomuga) was successfully modeled as a 2- or 3-way mixture of all possible sources according to the criteria outlined above. The *qpWave* and *qpAdm* analyses performed here suggest that the USR1 individual is admixed, contrary to Moreno-Mayar *et al.* (*4*).

11. A model comparison method for *qpGraph* resistant to overfitting

In this section we approach the model comparison problem from a different angle. Model likelihood almost always grows when unnecessary additional parameters are added, and that is a well-known problem of overfitting. Previously we tried to explore the LL and WR difference distributions observed under the null hypothesis on simulated data and set our thresholds accordingly. Here we propose a more general and less labor-intensive approach. We attempted to eliminate the overfitting problem by calculating in-sample and out-of-sample likelihood scores, which allow for fair comparisons of models of different complexity. We further test whether the difference in out-of-sample likelihood scores of two models is significant by bootstrap resampling across SNP blocks.

These methods are part of a new implementation of AdmixTools (https://uqrmaie1.github.io/admixtools/index.html, Maier *et al.* in preparation), and work as follows: $f_3$-statistics are calculated on around 700 5 cM SNP blocks. Instead of following the usual approach of fitting the *qpGraph* model using all SNP blocks, we randomly draw a subset of SNP blocks with replacement until the total number of blocks is reached. This constitutes one bootstrap iteration. The drawn set of SNP blocks forms the training set, while all other blocks form the test set. The training set is used for fitting the models, while the test set is used only to compute the final likelihood scores of both models. The final likelihood score is based on the difference between fitted $f_3$-statistics for the training set on one side, and estimated $f_3$-statistics for the test set on the other side. This procedure is repeated many times to yield a bootstrap distribution of likelihood score differences of two models, from which confidence intervals and a $p$-value can be obtained. We note that the covariance matrix of $f_3$-statistics, which is also necessary to compute the *qpGraph* likelihood score, is computed using all SNPs. This reduces noise without leading to overfitting.

We tested the new approach on the complex simulated graphs. The "diag" setting was kept at 0.0001, to match the setting used for all *qpGraph* runs in this study. It is important to keep this setting constant to ensure that LL values are comparable across datasets of the same size and containing the same number of groups. We performed 200 bootstrap iterations on simulated data (400,000 sites sampled with the 5% MAF filter, graphs with "unlocked" admixture proportions), comparing the baseline model for USR1 to 3-way models "$FAM_{EA}$ + Mal'ta + basal Mal'ta" and "$FAM_{EA}$ + Mal'ta + Houtaomuga", and those three models were compared to a 4-way model "$FAM_{EA}$ + Mal'ta + basal Mal'ta + Houtaomuga". We generated distributions of model-comparison $p$-values on 130 simulated datasets having various proportions of Houtaomuga ancestry in USR1: 0, 3.5, 7, 14, and 35% (Fig. 3e, Table S20).

Model-comparison $p$-values fluctuated above 0.1 for data simulated under the 2-way model and for two model pairs: baseline vs. "$FAM_{EA}$ + Mal'ta + Houtaomuga", and "$FAM_{EA}$ + Mal'ta + basal Mal'ta" (Fig. 3e, Table S20). We used empirical $p$-values since the distributions of LL differences were not normal (Fig. 3c, Fig. S49), and we used one-tailed tests since we did not expect that a simpler model would ever demonstrate a significantly better fit than a more complex model. When 14% Houtaomuga ancestry in USR1 was simulated, the model-comparison $p$-values on simulated datasets remained mostly below 0.1, and the median $p$-value fell below the statistical significance threshold of 0.05, but statistical significance was invariably reached only when 35% Houtaomuga ancestry in USR1 was simulated (Fig. 3e, Table S20). LL difference calculated using the standard approach on the same simulated data (35% Houtaomuga ancestry in USR1) invariably exceeded 100 ln-units and WR difference exceeded 3 SE (see 5$^{th}$

percentiles in Table S9). Given that $F_{ST}$ between the simulated groups was high (0.08-0.09 between FAM and USR1, 0.13-0.14 between USR1 and Houtaomuga under the baseline model, and 0.15-0.16 between FAM and Houtaomuga), we conclude that the new test is conservative, i.e. that statistical significance is reached only for strong admixture signals.

We then analyzed the real data (the full dataset, ca. 133,000 – 408,000 sites) under the same settings (graphs with "unlocked" admixture proportions, 500 bootstrap iterations, diagonal = 0.0001, 10 starting combinations of admixture weights tested for each model) (Table S20). To test if the results are dependent on the composition of source and target groups, we applied the new model-comparison approach to admixture graphs including different compositions of the American source: the composite group of non-Amazonian present-day SNA individuals (Aymara, Mixe, Mixtec, Piapoco, Quechua, Yukpa, Zapotec) (*11*), 44 alternative groups composed of ancient SNA and NNA individuals published mostly by Posth *et al.* (*14*), Moreno-Mayar *et al.* (*13*), and Scheib *et al.* (*18*), and 11 alternative present-day SNA groups partially overlapping with the composite FAM group (Table S20). FAM groups with unequivocal evidence of colonial or Paleo-Inuit admixture (Data S6) were not considered for this analysis. Different versions of the USR1 target (pseudohaploid and diploid) and Siberian source were also tested (two or three Houtaomuga individuals lacking West Eurasian admixture) (Table S20).

27 of 60 target/source combinations and 15 of 21 combinations having at least 400,000 sites available (under the "useallsnps NO" setting) showed a signal inconsistent with 0% ARB ancestry in USR1 (Table S20), i.e. demonstrated an empirical *p*-value below 0.1 for at least one of the two model comparisons informative for the ARB gene flow (baseline vs. "$FAM_{EA}$ + Mal'ta + Houtaomuga", and "$FAM_{EA}$ + Mal'ta + basal Mal'ta" vs. "$FAM_{EA}$ + Mal'ta + basal Mal'ta + Houtaomuga"). Results for the original FAM group of 13 present-day SNA individuals, 3 Houtaomuga individuals and the pseudohaploid USR1 version were statistically significant at $p$=0.032 (the baseline vs. "$FAM_{EA}$ + Mal'ta + Houtaomuga" model comparison, 500 bootstrap replicates) (Fig. 3c, Fig. S49). The baseline vs. "$FAM_{EA}$ + Mal'ta + basal Mal'ta + Houtaomuga" comparison on the same data demonstrated a much lower *p*-value = 0.002. This application of a new model comparison approach resistant to overfitting supports our main conclusion that the population represented by the USR1 individual experienced a gene flow from a Siberian source.

12. A more realistic graph model of the USR1 history

When constructing the skeleton graphs (Supplementary Text 5) and conducting mapping (Supplementary Texts 7, 8, 9), we used certain conventions that helped to streamline and standardize the modelling process, but those conventions make the resulting models unrealistic in certain ways. West Eurasian gene flows added to Siberian and American lineages that require them were always right below the leaf nodes, and they were added separately to each Siberian and American branch. For instance, there was no Mal'ta-related gene flow into the common ancestor of First Americans and USR1.

To check if the results generated using these skeleton graphs hold for more realistic topologies, we performed another graph model comparison. We assembled a small American clade composed of USR1 (pseudohaploid version), an NNA individual (the ancient Athabaskan I5321 used in topology tests above in Supplementary Texts 7 and 9), and an SNA individual (the ancient Peruvian IL7 used in topology tests above in Supplementary Texts 7 and 9) or group (the FAM group of 13 present-day individuals). That clade was grafted onto the 14-population skeleton graph, and a shared Mal'ta-related gene flow was added at the basis of the American clade. Three Houtaomuga individuals were used, and only the Neanderthal gene flows were locked (Fig. S50). The same graphs were fitted to the 1240K-based dataset without transitions and to the full 1240K-based dataset used above. Thus, four population/dataset combinations were tested. A gene flow from the Asian source of Paleo-Inuit was added to the Athabaskan branch by default.

First, the baseline models for USR1 were tested: USR1 as a member of the American clade in one of three possible positions, and without any gene flows unique to the USR1 branch. Next, either the "basal Mal'ta" or the Houtaomuga gene flow was added to the USR1 branch (3-way models). Then 4-way models were tested that included gene flows in the order Mal'ta (shared), Houtaomuga, "basal Mal'ta". Another class of 4-way models included a Houtaomuga- and basal Mal'ta-related lineages mixing first and then admixing into USR1. In all cases three possible topologies of the First American sources were tested.

The "basal Mal'ta" gene flow to USR1 was supported by LL and WR differences vs. the baseline model (Table S21) that exceeded the thresholds established in Supplementary Text 2 albeit for a subtly different graph. The Houtaomuga gene flow was not supported when added first, but when added on top of the "basal Mal'ta" flow it was supported by WR differences on the dataset without transitions and by LL differences on the full dataset (Table S21). For any 2- and 3-way model the basal position of the First American source of USR1 was supported by significant LL differences (>3 ln-units) and by the fact that the other positions resulted in a trifurcation. In the case of the 4-way models, the branching order of the First American sources was impossible to resolve based on LL or WR, but the basal position of $FAM_{USR1}$ invariably resulted in a trifurcation, in contrast to the other topologies (Table S21). The order of gene flows had no influence on LL or WR of the 4-way models. Reversing the Houtaomuga => USR1 gene flow in the context of the 4-way models resulted in significantly worse fits.

In general, these results are very similar to those of graph mapping and branching order explorations presented above in Supplementary Texts 7, 8, and 9. We conclude that introducing a Mal'ta gene flow shared by all First Americans has little impact on the outcomes of the model comparison procedure. We use the model shown in Fig. S50 as a final model featured in simplified form in Fig. 3a.

13. Automated graph space search and a bias-free dataset

All graph building protocols described above had imposed directionality because a target was mapped on a skeleton, and only some gene flows found in this way were flipped to test their direction. This is a limitation, and to overcome it we took advantage of an automated graph optimization algorithm that is similar in its spirit to the *TreeMix* software (Pickrell and Pritchard 2012) and that is implemented in *ADMIXTOOLS 2* (https://github.com/uqrmaie1/admixtools). We used the "find_graphs" function to identify admixture graphs with good fits. For a fixed set of populations and a fixed number of admixture events, "find_graphs" searches the space of possible admixture graphs in the following way: random admixture graphs are generated, and the likelihood score for each one is evaluated. A fraction of graphs is then selected to enter the next generation, such that graphs with better scores are selected. The selected graphs are then randomly modified by one out of a number of so-called "mutation functions". The single best graph is in addition passed to the next generation without being modified. All selected and possibly modified graphs are then evaluated, and the resulting scores are again used as a selection criterion for the next generation. This process is repeated for a number of generations, until the best likelihood score reaches a plateau, and a winning graph is identified. In order to avoid getting stuck in a local optimum, and to assess the robustness of the approach, the "find_graphs" function can be repeatedly run, each time starting with different random graphs. If the winning graphs in independent runs are similar to each other, it suggests that the winning graph is close to the global optimum.

Another important limitation is that most analyses were performed on two SNP sets derived from the 1240K SNP panel (*25*) and on a limited set of populations. We constructed a drastically different dataset that is expected to be free of most types of bias: 1/ uneven deamination damage across ancient DNA samples that leads to ancient DNA attraction; 2/ reference bias (Günther and Nettelblad 2019), 3/ ascertainment bias caused by the paucity of rare variants (Supplementary Text 2) that is typical for SNP panels (*31*); 4/ background selection and biased gene conversion which possibly affect 97% of variable sites in the human genome (*32*); 5/ various levels of missing data across groups. We focused only on neutrally evolving variants as defined by Pouyet *et al*. (*32*): G⇔C and A⇔T mutations lying withing regions with a high recombination rate (1.5 cM/Mb according to the Yoruba recombination map). These sites are all transversions, which automatically takes care of bias no. 1. Moreover, we considered only shotgun-sequenced ancient genomes having a coverage of 5x and higher. Thus, we do not expect to see attraction between ancient vs. present-day groups, and we do not expect to see ascertainment bias since sites were not selected based on allele frequency. To take care of the reference bias, an appropriate PileupCaller protocol (https://github.com/stschiff/sequenceTools) was used, and pseudohaploid genotypes were called at each site. In short, pseudohaploid genotypes were called within regions mapped by Pouyet *et al*. (*32*), at biallelic G⇔C and A⇔T sites listed in dbSNP (https://ftp.ncbi.nih.gov/snp/organisms/human_9606_b151_GRCh37p13/VCF/00-All.vcf.gz). No missing data were allowed at the group level. Sixty-nine individuals grouped into 55 groups were taken into the analysis (Data S9). The resulting dataset was composed of 1,270,644 variable sites. The population composition of the dataset was drastically different from the 1240K datasets in this study, but it incorporated all relevant groups: nine SNA individuals, USR1, Kolyma1, an Early Neolithic ARB individual from the Devil's Gate site (*10*), two Upper Paleolithic individuals from the Yana site (*10*), the Saqqaq Paleo-Inuit, the Tibetan Chokhopani individual, an Andamanese individual, etc. (Data S9).

Before exploring the graph space, we first constructed a Eurasian-American PCA plot to check if the neutral dataset recovers the same global population structure as the 1240K dataset (Fig. S51). Before running PCA, linked sites were pruned from both datasets using the following function in *PLINK v.2*: --indep-pairwise 2000 500 0.5. Ten PCs were calculated using *PLINK v. 1.90b6.10* on 59 Eurasian and American individuals (excluding the Andamanese, Africans, and archaic humans). The 1240K dataset was taken directly from the following source: https://reich.hms.harvard.edu/downloadable-genotypes-present-day-and-ancient-dna-data-compiled-published-papers. We observe the same pattern across both datasets, with arguably a better resolution of major genetic clusters in the case of the neutral dataset (Ust'-Ishim vs. other Eurasians, the Tibetan Chokhopani vs. Siberians).

We also visualized $f_4$-statistic biases. All possible $f_4$-statistics (1,023,165 statistics) were calculated (Data S9) for both datasets using *ADMIXTOOLS 2* (starting from $f_2$-statistics, with an option "afprod=TRUE", on 5 cM blocks). For these calculations only sites genotyped across all 55 groups were used. The first collection of biplots shows a general overview of $f_4$-statistics and their Z-scores (Fig. S52). The second collection of biplots (Fig. S53) shows that the most biased statistics include two African groups and an archaic human, while statistics including Americans and Siberians are relatively unbiased. These results are in line with those reported by Bergström *et al.* (*31*), who compared $f_4$-statistic calculated on shotgun data for 929 present-day individuals, on two Illumina panels and the HumanOrigins panel, and also compared statistics calculated on rare and common variants. See also Fig. S53e which shows that calculating $f$-statistics on common variants reproduces the biases seen when comparing the 1240K panel and the set of neutrally evolving sites derived from shotgun data.

Next, a space of relatively simple graphs incorporating 13 groups (listed in Data S9) and 6 admixture events was explored in an automated way. After removing missing data across groups, 1,058,768 SNPs remained, and 240,506 were polymorphic. Only the latter group of sites was used for fitting admixture graphs. Per-block $f_2$-statistics were calculated using *ADMIXTOOLS 2* and stored (537 SNP blocks of 5 cM were generated). Subsequently $f_3$-statistics were calculated based on the stored $f_2$-statistics and admixture graphs were fitted to the $f_3$-statistics, generating fit scores (log-likelihoods). Four independent graph exploration runs were started in parallel from sets of random graphs and consisted of 5,000 replicates each. 3,000 best-ranking graphs were inherited from the previous replicate and were used to generate 5,000 graphs for the subsequent replicate by randomly applying one of five graph modification procedures described above. The diagonal was set at 0.0001, Denisovan was set as an outgroup, and to speed up the process only one initial combination of admixture weights was tried for each graph. In total, up to 25 million graphs were explored in each run, which constitutes $7.61\times10^{-21}$ of the graph space.

All four graph exploration runs reached a likelihood score plateau, and results (the best-scoring models) for two runs are shown in Fig. S54. The four graphs generated in an automated way on the set of neutrally evolving sites were very similar, and included several well-known admixture events: for instance, the Neanderthal gene flow into all non-Africans, the West Eurasian gene flow into Native Americans, and the basal Eurasian gene flow into Caucasian and Middle Eastern groups. On the other hand, two "unconventional" admixture events shared by all four "winning" graphs were as follows: a 10% gene flow from an American or Siberian source into the Caucasian hunter-gatherer (Kotias), and a 14-17% gene flow from the USR1 branch into the Yana branch. The latter event seems implausible from historical perspective since the Yana individuals are dated to 31600 cal yr BP (*10*), but the USR1 individual is much younger (11500 cal yr BP), and >30000 cal yr BP is an improbable age for the divergence of American sub-clades (4). We hypothesized that this event is a conflation of two separate events: an Asian gene flow into the Yana group that was reported before (*10*) and the "basal Mal'ta" gene flow into the USR1 lineage documented in the preceding sections of this study.

In order to resolve the Yana/USR1 interaction better, we added two groups potentially informative for this relationship: the Saqqaq Paleo-Inuit (*101*) and the Mesolithic Kolyma individual (*10*). The SNA group was also replaced (see the group composition of this new model in Data S9). We recalculated *f*-statistics for the new set of groups and sites (231,753 polymorphic sites), but this time we did not resort to the automated topology exploration method since the graph space grows too quickly with any group or admixture event added. We performed manual exploration of most likely branching points for the newly added groups, manually tested additional admixture events, and incorporated branching orders and gene flows know from the literature and revealed in this study above: e.g., the ARB and "basal Mal'ta" gene flows into the USR1 lineage and the attraction between ancient Brazilians from the Sumidouro site and Andamanese (*13*). When adding an admixture event, several alternative sources and both gene flow directions were routinely tested. Models were ranked according to LL, and a threshold of 3 ln-units was used as a significance threshold for models having the same number of parameters (*11*). The resulting model is shown in Fig. S55.

The model shown in Fig. S55 fits the data (WR = 2.66 SE) and incorporates the Siberian gene flow into USR1, although its source is not located on the ARB branch (since the source on the ARB branch results in a trifurcation). The Kolyma individual was modelled as a Siberian/American mixture, to reflect other results in this study and the results reported by Yu *et al.* (*20*). The model also incorporates the interaction between Americans and Andamanese (*13*), although the gene flow direction to the Andamanese and not to Americans was supported by an LL difference of 4.6 units. The model incorporates the Asian admixture in Yana reported by Sikora *et al.* (*10*) and the well-documented Yana/Mal'ta-related gene flows into Americans and Paleo-Inuit. The gene flow from Kolyma to Saqqaq suggested by Sikora *et al.* (*10*) and elsewhere in this study was not supported (LL difference of 60.7 units).

We explored the following derivatives of the model shown in Fig. S55. First, gene flows were dropped one by one, resulting in 18 graphs. Likelihood scores of the resulting graphs were invariably worse than that of the original model, and the lowest LL differences observed were 8.8 ln-units (dropping the Siberian=>USR1 gene flow, WR difference = 0.88 SE), 9.5 ln-units (dropping the American=>Andamanese gene flow, WR difference = 0.54 SE), and 21.4 units (dropping the proto-Kolyma=>CHG gene flow, WR difference = 0.69 SE). The LL and WR difference thresholds optimized in Supplementary Text 2 are likely not applicable to this drastically different dataset, so we performed the model-comparison bootstrapping procedure described in Supplementary Text 11. The highest *p*-value = 0.05 (based on 500 bootstrap replicates) among the three model pairs tested corresponded to the Siberian gene flow dropped. Thus, the gene flow is moderately significant, similar to the result observed for the 1240K-derived dataset above.

Next, we flipped the direction of gene flows one by one, wherever that was possible, resulting in 16 graphs. Likelihood scores of the resulting graphs were invariably worse than that of the original model, and the lowest LL differences observed were 7 ln-units (flipping the American=>Andamanese gene flow), 7.7 ln-units (flipping the Siberian=>USR1 gene flow), and 10.7 units (flipping the basal Asian=>Yana gene flow). All these LL differences are significant according to the threshold value of 3 units (*11*). Finally, we added one admixture event at all possible positions, resulting in 159 graphs. Of those, 109 demonstrated an improved likelihood score, however the improvements were marginal. The largest improvement observed (7.4 units for the Ust'-Ishim=>Devil's Cave N gene flow, WR improvement = 0.19 SE) was not supported by the model-comparison bootstrapping procedure (*p*-value = 0.146, 500 bootstrap replicates).

The graph shown in Fig. S55 was also fitted to the original 1240K dataset (807,106 polymorphic sites including transitions). The graph had a relatively poor fit: WR = 4.57 SE. Flipping the gene flows one by one produced a rather unexpected result: a large LL difference (20.3 units) supported the non-African=>Neanderthal gene flow direction. Large LL differences (up to 22 units) also supported gene flows from various non-Africans to the South African group. We suppose that these results are caused by the biased *f*-statistics including African and archaic groups (Fig. S53b).

#### 14. Linguistic implications of the gene flow between Siberians and Ancient Beringians

##### *a. Potential Chukotko-Kamchatkan-like linguistic substrate in Alaska*

Alexei S. Kassian

Below I evaluate some words of Tanaina, a modern Athabaskan (Dene) language spoken in coastal Alaska which belong to basic vocabulary yet have no etymological connection to other Athabaskan languages. These words probably originate from a pre-Athabaskan substrate language. Several basic terms (including Swadesh items) demonstrate phonetic and semantic closeness to the reconstructed Proto-Chukotko-Kamchatkan vocabulary. The assumed pre-Tanaina substrate language might either belong to the Chukotko-Kamchatkan family or be an outlier in the Chukotko-Kamchatkan–Nivkh clade.

Tanaina, a.k.a. Dena'ina, is an endangered language of the Athabaskan group of the Na-Dene family (*103, 104*). It consists of several dialects spoken in the southern part of Alaska. Modern Tanaina's territory partially overlaps with the area of the Denali archaeological complex. A specific feature of Tanaina is that it lacks inherited Athapaskan roots for some semantic concepts (including Swadesh items) using non-etymologizable expressions instead. These cases were collected by James Kari (*104*: xxi, *105*: 545, *106*: 59), who called them "elite replacements", explaining them as either substrate loans or taboo replacements. Actually, the Tanaina stems analyzed by Kari can be divided into two classes.

The first class consists of nominalized verbal forms, which have superseded original roots; e.g., ‘head' is expressed as ‘tip that extends', and so on. In these cases, we are most probably dealing with normal gradual evolution of the original lexicon during natural language development. Note that we are not aware of specific evidence that such replacements were indeed taboo-driven in Tanaina.

The second class consists of Tanaina words, which are morphologically unanalyzable and unetymologizable. For example, ‘fire' is expressed by the enigmatic form *tazʔi* in the bulk of the Tanaina dialects, whereas the inherited Athabaskan term *qʰən* ‘fire' is only used in the Upper Inlet dialect. In these cases, it is likely that we are dealing with remnants of a substrate language which was previously spoken in this territory and later assimilated by Tanaina centuries ago. For the sake of convenience one can label this potential substrate language as "Pre-Tanaina".

The potential Pre-Tanaina words partially belong to the basic vocabulary. First of all, these include several Swadesh items:

- *ʁəs* ‘bone' (*104*: 12).
- (*ə*)*łtʰuʁ* ‘eye' (*104*: 88).
- *tazʔi* ‘fire' (*104*: 248).
- *č’ix* ‘hair (of head)' (*104*: 87).
- *miłni* ~ *piłni* ~ *vinłni* ‘water' (*104*: 121, 293).
- *k’is-ən* ‘woman; girl' (*104*: 72), *k’is-i* ‘female animal' (*104*: 13).

Other Tanaina basic terms of unclear origin are:

- *čʰik’a* ‘wood, dry wood, stick, firewood' (*104*: 63, 250).
- *iƛ’* ‘clothing (in general)' (*104*: xxi, 165).

- *izin* ‘arrow' (*104*: xxi, 207).
- *k’aƛ’* ‘snow showers with moist large flakes' (*104*: xxi, 153).
- *qʰičʰi* ‘old woman, old lady' (*104*: 73).
- *χenzi ~ χeɬni* ‘coma, unconscious' (*104*: 99).
- *yusti* ‘open fire, fireplace, outdoor hearth' (*104*: 248).

More cultural Pre-Tanaina words are:

- *ʔa-* (incorporated) ‘to hunt bear at night' (*104*: xxi).
- *ʔək* ‘to act as a shaman' (*104*: 307).
- *čanču* ‘medicine object, dream object kept as source of power; shaman's spirit that can travel' (*104*: 306).
- *ič* ‘a contagious disease (unidentified)' (*104*: 99).
- *kəš ~ kʰəš* ‘webbing holes in snowshoe frame' (*104*: 239).
- *nənli ~ nəli ~ ənli ~ nəlni* ‘stream bath' (*104*: 221).
- *niqitay* ‘large skin boat with flat-bottom pole frame (downstream use)' (*104*: 243).

Tanaina names of endemic species form a specific subset of the potential substrate vocabulary:

- *aqema* ‘arctic ground squirrel' (*104*: 8).
- *k’=čʰəʁuxa* ‘marten, pine marten' (*104*: 5), indefinite possessive prefix *k’-*.
- *k’=qʰušiya* ‘marmot' (*104*: 5), indefinite possessive prefix *k’-*.
- *palušuk* ‘chiton (mollusk)' (*104*: 22).
- *qʰančʰi* ‘porcupine' (*104*: 7).
- *qʰiyχi* ‘marmot' (*104*: 5).
- *qʰunša* ‘arctic ground squirrel' (*104*: 8).
- *tʰaza* ‘sea lion' (*104*: 9).

Out of the 13 aforementioned words from Pre-Tanaina basic vocabulary, 8 stems have phonetically similar and semantically compatible counterparts in the Chukotko-Kamchatkan family.

The Chukotko-Kamchatkan family consists of two branches (sub-families): Chukotian and Itelmen (a.k.a. Kamchatkan). A genetic relationship between Chukotian and Itelmen is generally accepted by experts (*107-113*). The opposite opinion was expressed by Worth (*114*) and Volodin (*115, 116*: 224–228), who supposed that the observed Chukotian-Itelmen matches, which cover not only basic vocabulary but also some main grammatical exponents (*109*), are contact loans from one language group to another. Worth-Volodin's scenario, however, clearly contradicts the theory of language contacts which predicts that cultural vocabulary is borrowed first, whereas basic vocabulary and main grammatical exponents are most protected from borrowing (*117*).

My Proto-Chukotian reconstruction is based on Fortescue's (*110*) etymological dictionary as well as on main lexicographic sources on synchronic Chukotian languages. Reconstruction of Proto-Itelmen is a less trivial task, because 18$^{th}$-19$^{th}$ century data on the extinct Eastern and Southern Itelmen languages are not very reliable and consistent. I generally follow Mudrak's reconstruction of Proto-Itelmen (*112, 118, 119*) with emendations and/or simplifications, if needed.

There are too many unsolved obstacles in the Chukotian-Itelmen comparison at the current stage of research which makes reliable phonological reconstruction of a Proto-Chukotko-Kamchatkan vocabulary impossible. Instead we are forced to use separate Proto-Chukotian and Proto-Itelmen vocabularies.

The most striking case of coincidence between Pre-Tanaina and Chukotko-Kamchatkan with direct semantic match is:

(1) *miłni, piłni, vinłni* ‘water' (*104*: 121, 293), the variants are distributed among the Tanaina dialects. Initial *m-* points to a non-inherited form, it was denasalized *m* > *p~v* in some dialects in order to avoid a foreign nasal labial phoneme as, for instance, in Tanaina *milo* ~ *vilo* ‘soap' < Russian *mɨlo* ‘id.' or *muka* ~ *puka* ‘flour (food)' < Russian *muka* ‘id.'. ► Cf. Proto-Chukotian **mi-məl* (partial reduplication) ‘water' (*110*: 99).

Another interesting case is:

(2) (*ə*)*łt*$^{h}$*uʁ* ‘eye' (*104*: 88). ► The Tanaina stem *łt*$^{h}$*uʁ* may contain cognates of Chukotko-Kamchatkan **lV* ‘eye', reduplicated pl. **lV-lV* (> Chukotian **lə-lä* ‘eye', Itelmen **lo-* ‘eye', pl. **lu-l-*, *110*: 163; *118*) and the Chukotko-Kamchatkan plural or rather dual ending *-*t(V)* (110: 426, 433). Final *-ʁ* remains without explication, however.

Other cases with weaker phonetic compatibility and/or without direct semantic matches are:

(3) *k'aƛ'* ‘snow showers with moist large flakes' (*104*: xxi, 153). ► Chukotko-Kamchatkan **qəl* ‘snow (esp. on ground)' (> Chukotian **ʁəl-ʁəl*, Itelmen **qal-ʔal*, reduplication in both cases, *110*: 243). Dissimilation *q-q* > *q-ʔ* in Itelmen suggests how the ejectivity in the Athabaskan form might emerge.
(4) *ʁəs* ‘bone' (*104*: 12). ► Itelmen **kas(-)x-* ‘shank, shin' (*118*). For a similar semantic shift, cf., e.g., Icelandic *leggur* ‘shank; bone (of arm or leg)'.
(5) *k'is-ən* ‘woman, girl' (*104*: 72), *k'is-i* ‘female animal' (*104*: 13); final *-ən* and *-i* are human and non-human nominalizers respectively. ► Chukotian **čakə-ɣet* ‘sister' (*110*: 42), Itelmen **čik-* ‘girl, young woman' (*118*). A metathesis?
*(6) iƛ'* ‘clothing (in general)' (*104*: xxi, 165). ► Chukotian **iðʁə-n* ‘parka', Itelmen **eč'* ‘clothing' in the complex negated stems ‘naked' and ‘to get undressed' (*110*: 94; *118*).
(7) *č'ix* ‘hair (of head)' (*104*: 87). ► Chukotian **ðəɣ* ‘fur' (*110*: 64).
(8) *yusti* ‘open fire, fireplace, outdoor hearth' (*104*: 248). ► Chukotko-Kamchatkan **uyi* ‘to make fire, cook' (> Chukotian **uyi*, Itelmen **uyi*, *110*: 305). The assumed nominal suffix *-st-* in the Tanaina form requires additional explanation.

The lexical similarities offered above may indicate that the assumed linguistic substrate of Tanaina (Pre-Tanaina) represented a close relative of the Chukotko-Kamchatkan family. In other words, it might be a trace of an ancient migration from Siberia to Alaska which probably predated the Athabaskan radiation.

Assessing the offered phonetic and semantic similarities between Proto-Chukotko-Kamchatkan and a recent Tanaina substrate, one should keep in mind that we are dealing with a very long chronological distance probably comparable, say, with the distance between Proto-Balto-Slavic (a representative of the Indo-European family) and modern Finnish (a representative of the Uralic family). The split between Indo-European and Proto-Uralic branches is dated back to *ca.* 8-10 millennia BC (*120*). Out of 7 Proto-Indo-European–Proto-Uralic lexical etymologies within the Swadesh 110-item wordlist (*121*: 320), 5 roots have Proto-Balto-Slavic and modern Finnish descendants without semantic changes, but with very different degree of phonetic similarity:

- 'I (obl.)': Proto-Balto-Slavic **me-n-* / Finnish *minä*.
- 'name': Proto-Balto-Slavic **inmen-* / Finnish *nimi*.
- 'thou': Proto-Balto-Slavic **tu:* / Finnish *sinä*.
- 'water': Proto-Balto-Slavic **vad-a:* / Finnish *vesi*.
- 'who?': Proto-Balto-Slavic **k-* / Finnish *kuka*.

Note that I intentionally do not use the hyphen in the Finnish forms *mi-nä* 'I', *si-nä* 'thou' and *ku-ka* 'who', since we know nothing about morphology of the hypothetical Tanaina substrate and are not able to propose a synchronous morphological analysis of the Tanaina substrate forms.

Can the observed phonetic similarities between Pre-Tanaina and Chukotko-Kamchatkan be chance coincidences? Definitely yes, but unfortunately we do not have available tools to estimate the probability of getting these similarities due to chance. Weighted permutation test as developed in Changmai *et al.* forthcoming returns insignificant $p = 0.333$, if 6 available Swadesh items of Pre-Tanaina and Proto-Chukotko-Kamchatkan are compared (Table S22). It is not expected, however, to obtain a significant $p$-value for such a time distance since the accumulated phonetic mutations should seriously obscure original phonetic similarities. For examples, the permutation test applied to the Proto-Balto-Slavic and modern Finnish 110-item wordlists also returns insignificant $p >> 0.05$, but comparison between the wordlists with a shorter time distance, Proto-Indo-European and Proto-Uralic, yields significant $p = 0.019$ or 0.005 depending on the procedure (*121*).

Nevertheless, from the traditional linguistic point of view the available matches, especially Tanaina *miłni* 'water' / Proto-Chukotian **mi-məl* 'water', looks persuasive enough to hypothesize about a Chukotko-Kamchatkan-like substrate in Alaska.

| meaning | Tanaina | Proto-Chuk. | Proto-Itelmen | Proto-Nivkh |
|---|---|---|---|---|
| 'bone' | ʁəs | *qətʁəm | *ktχʷəm | *ŋa=ɲɣəv |
| 'eye' | łtʰuʁ ~ əłtʰuʁ | *lə-lä | *lo- | *ɲŋaɣ |
| 'fire' | tazʔi | *milɣə- | *xi=młx | *tu-ɣr ~ *tuɣ-r |
| 'hair' | č'ix | *kəð=wir | *k'ʷimi | *ŋa=mɣ |
| 'water' | miłni | *mi=məl | *iʔ | *ȶaʁ |
| 'woman' | k'is-ən | *ŋäv- | *ŋim-sx | *taŋq |

Table S22. Six Swadesh item of the assumed Tanaina substrate language (Pre-Tanaina) and their Proto-Chukotian, Proto-Itelmen and Proto-Nivkh semantic counterparts.

This substrate scenario raises a question how the assumed Tanaina substrate correlates with Nivkh, a compact language family spoken in the Amur river region and on Sakhalin island, since it is likely that Nivkh is the closest linguistic relative of Chukotko-Kamchatkan (*36*). See Changmai *et al.* forthcoming for probabilistic evaluation. In this case, lexical matches between Pre-Tanaina and Proto-Nivkh also could be revealed, but I failed to find convincing etymologies between the Pre-Tanaina forms and the Proto-Nivkh basic vocabulary (as reconstructed by Fortescue (*122*)). The factual absence of recognizable matches between Pre-Tanaina and Proto-Nivkh allows for at least two explications. First, the cognate words in the Tanaina substrate and/or Proto-Nivkh might have mutated to such a degree that currently are not discernible to the naked eye. Second, since the number of Pre-Tanaina words known to us is very modest (e.g., out of 110 Swadesh items, only 6 words are available), it would not be surprising if all historical Pre-Tanaina–Nivkh cognates fell into a non-attested part of the Pre-Tanaina vocabulary. If we assume that historically there were, say, 10 Pre-Tanaina–Proto-Nivkh etymological matches within the Swadesh 110-item wordlist, the probability of missing all these etymologies in the random subset of 6 concepts (as in the Tanaina case where only 6 Swadesh items likely belong to a substrate language) is very high: $p = 0.557$.

In summary, the available linguistic data (if we are not dealing with a series of chance coincidences) suggest that the assumed pre-Athabaskan substrate language might either belong to the Chukotko-Kamchatkan family or be an outlier in the Chukotko-Kamchatkan–Nivkh clade.

##### *b. Modeling diverse Asian origins for indigenous North American language families*

by Edward Vajda

The discovery of a Late Pleistocene gene flow from Northeast Asia into Northern Native Americans (NNA) dated to at least ~12,000 yBP that in this publication is termed Houtaomuga-related (from the eponymous archaeological site) has implications for understanding linguistic prehistory on both sides of Bering Strait. The genes in question are shared on the Asian side by contemporary speakers of Nivkh, Tungusic, Japanese, and Korean languages, but in North America this genetic component was found only in a 11,500-years-old individual USR1, whose genome is the best-quality representative of the Denali archaeological culture. A signal found in a much later ancient Athabaskan individual (ca. 730 yBP, Data S6) was supported on one dataset only, and thus unequivocal evidence for the survival of this component in recent and contemporary peoples is lacking. But given the location of USR1 in Alaska and subsequent development and contacts of the Denali culture attested archaeologically, we expect to find Houtaomuga-related genetic heritage among speakers of four major primary families (Algic, Salishan, Eskaleut, Na-Dene) and several smaller primary families or isolates indigenous to the Pacific Northwest (Tsimshianic, Wakashan, Haida, and possibly also Chimakuan and Kutenai). "Primary family", "family", and "isolate" – a language family with only one member – all refer to genealogical taxons with no currently proven deeper relatives. The goal here is to triangulate data from linguistics, genetics, folklore studies, and archaeology to assess whether chronologically deeper genealogical relationships might exist among these families that would parallel this human genetic link. Several distinct scenarios for the genesis of linguistic diversity in the North America and its more ancient roots in Eurasia are identified.

While archaeology, genetics and the study of folklore motifs have already yielded many facts of potential relevance for modeling the prehistoric origins of indigenous New World language families, several factors continue to argue against the likelihood that language relatedness can be firmly demonstrated at time depths older than the Early to Mid-Holocene. Languages change at variable and unpredictable rates. Attempts to determine a separation time between languages known to be genealogically related based on equating the percentage of shared cognates with a particular time depth (glottochronology) are unreliable. Only when parallel information from historical records, human genetics, or archaeology happens to be available has it been possible to determine the timing of a language family's breakup. One primary family dated with the help of extra-linguistic methods is Indo-European (in the sense of Indo-Hittite), for which archaeological parallels help demonstrate a time depth range of 5800 to 6500 yBP (*123*). Possibly the oldest universally accepted primary family proven using the Comparative Method is Afroasiatic, which contains Arabic, Hebrew, Amharic, Berber and many other languages of northwestern Africa, and is claimed to have a time depth of ~8,000 years (*124*). The apparent temporal limitations of the Comparative Method cannot be overcome by substituting human genetic findings for linguistic evidence. The lack of strong correlation between population genetics and language relatedness over time precludes the reliability of the former in contributing actual evidence of linguistic relatedness. The best that can be achieved at present in exploring the Old-World origins of New World language families from an interdisciplinary perspective is a clear articulation of which linguistic scenarios are indicated as plausible by the currently known facts from other scientific disciplines, and which are not. In other words, which speculative hypotheses of distant language relatedness have parallel support from human genetics and archeology, and which do not?

The discovery of a Late Pleistocene Northeast Asian gene flow into at least some NNA (Northern Native American) populations opens the logical possibility that one or more indigenous language families of northwestern North America may descend not from an ancestor spoken in Beringia throughout the Beringian Standstill but rather from one brought after the end of the standstill only about 16,000 years ago. It had previously been thought that all First Peoples of the Americas (defined as indigenous New World populations whose ancestors entered North America from Asia in Pleistocene times) descended from a population that had remained isolated in Beringia during the Last Glacial Maximum from ~23,000 to ~15,500 yBP, a period called the Beringian Standstill. This population later contributed to the genomes of indigenous populations across the Americas. Given the time depth involved, any linguistic connections between contemporary Old-World populations and First Peoples of the Americas mediated through descent from a language spoken in Beringia during the Standstill would necessarily date back deeper than ~23,000 yBP, far too ancient to be demonstrated using accepted comparative linguistic methods. The newly discovered human genetic link between certain populations of Northeast Asia and North America at the much younger time depth of at least 12,000 years ago opens new possibilities for investigating cross-Bering linguistic connections, though the time depth involved may still be too remote to evaluate them with full confidence.

Aside from this gene flow, the ancient Beringian population appears to have been isolated from contact with other populations of Eurasia throughout the last 10,000 years of the Late Pleistocene. At the time of European contact in 1492, the indigenous peoples of the Americas spoke over 120 primary language families (including many isolates) spread from Alaska to Tierra del Fuego. All of these peoples, including speakers of Eskaleut and Na-Dene, possess at least some genetic ancestry derived from the population living in Beringia during the Standstill.

The population ancestral to all Native Americans underwent a further split ~15,500 yBP into Southern Native Americans (SNA) and Northern Native Americans (NNA) (*13, 15*). SNA groups exclusively populated all of South and Central America, as well as much of North America. NNA genes are found in the remaining indigenous populations of northern North America. Looking from the vantage of genetics and archaeology, it is therefore possible that every primary language family of the New World descends from a language (or several languages) spoken in Beringia during the later phase of the Standstill and later spread south into the Americas by either NNA or SNA populations.

For SNA populations, a language origin from Beringia early in the Standstill remains the only logical possibility so far indicated. This encompasses over 90% of the indigenous primary language families documented in the Americas (about 115). If it is true that language genealogy cannot be reliably traced back beyond eight or ten thousand years, then it would not be possible to determine whether SNA ancestors brought multiple distinct proto-languages into the Americas from Beringia in the Late Pleistocene or only one. The Early Beringian population derived about 70% of their genetic ancestry from East Asians and 30% from a deep branch of the European clade termed Ancient North Eurasians or ANE (*13, 15, 40*). An analysis of the comparative folklore motifs presented in Berezkin (*125*) suggests that several founding groups with distinct cultures emerged from the post-Standstill Beringian bottleneck to give rise to New World populations. It is therefore plausible that multiple distinct languages connected with either the ANE or East Asian genetic components of Beringia's population were spoken in Beringia at the beginning of the Standstill and that the descendant(s) of more than one of them were brought into the Americas in the Late Pleistocene. The language(s) spoken by the earliest SNA populations in the Americas might therefore descend from: 1) a single language of ANE origin; 2) a single language of ancient East Asian origin, or 3) multiple languages from both or either origin.

These same possibilities equally apply to the linguistic origins of NNA populations, given their shared origin with SNA populations earlier in Pleistocene Beringia at a time depth of ~15,000 yBP. As is true of languages spoken by SNA populations, all languages spoken by populations with NNA genes could likewise derive from a proto-language (or several proto-languages) with ancestral roots present in Beringia during the early phase of the Standstill. However, there exists a second possibility for the linguistic provenance of NNA populations that is not indicated for SNA populations: some language families spoken by NNA populations may descend from a proto-language connected instead with the newly discovered Northeast Asian gene flow brought into Late Beringia after the end of the standstill. The ramifications of this possibility for historical-comparative linguistics will be explored in detail below.

Only two primary language families spoken by populations with NNA genes show additional possibilities. Proto-Na-Dene could have entered Alaska much later, during the Mid-Holocene about 5,000 years ago in connection with the so-called Paleo-Inuit (Paleo-Eskimo) migration from Siberia (*11*), rather than deriving from a source in either Ancient Beringia or Late Pleistocene Northeast Asia. Finally, for Eskaleut speakers there exists both this possibility as well as a fourth: while Proto-Eskaleut probably first arrived in North America ~5000 yBP, the population directly ancestral to Yupik- and Inuit-speaking groups may have crossed Bering Strait only ~2000 yBP.

Table S23 summarizes the four logical possibilities for indigenous New World language origins so far suggested by genetics and archaeology:

**Table S23. Plausible Origins of Indigenous New World Language Families**

**Hypothesis One: Early Beringian Pleistocene Origin**: The language descends from one or more proto-languages spoken in Beringia already at the earliest phase of the Standstill with deeper origins connected either with ANE or Ancient East Asians. New World language families with Early Beringian origins would therefore be separated by a time depth of at least ~23,000 years from any Old-World relatives and probably not recoverable by the Comparative Method. For indigenous languages spoken by populations with SNA genetic origin (about 115 of the recognized 123 primary New World families), this is the only possibility currently indicated by human genetic evidence. Therefore, the Early Beringian origin hypothesis encompasses the vast majority of indigenous language families of the Americas. Although a linguistic origin in Early Beringia is likewise theoretically possible for the seven to nine primary families spoken by populations with the NNA genetic component (Eskaleut, Na-Dene, Haida, Tsimshianic, Wakashan, Salishan, Algic, as well as possibly Quileute and Kutenai), there exist additional possibilities to account for their Old World origins.

**Hypothesis Two: Late Pleistocene Northeast Asian Origin**: This hypothesis is new and considered here for the first time. The language descends from a proto-language not present in Early Beringia but rather brought from Northeast Asia after the end of the standstill about 16,000 years ago, and therefore potentially separated from possible linguistic relatives in Northeast Asia at approximately the same time depth. This "Trans North Pacific" hypothesis appears plausible only for languages spoken by NNA populations, including Na-Dene and Eskaleut, and not to populations in the southern or eastern areas of North America or in Central or South America, for which only Hypothesis One applies.

**Hypothesis Three: Mid-Holocene Siberian Origin**: The ancestral language arrived in Alaska ~5000 yBP in conjunction with the so-called Paleo-Inuit migration, which contributed at least 50% genetic ancestry to Eskaleut speakers and 30-40% to early Na-Dene speakers (*11*). This hypothesis is not applicable to other NNA populations, including Haida speakers, who lack this gene flow except where later admixed with neighboring Tlingit or Dene speakers. Based on the sharp linguistic difference between Eskaleut and Na-Dene, which are universally recognized as completely unrelated, a Mid-Holocene Siberian origin for both families is plausible only if two linguistically distinct groups crossed Bering Strait and entered Alaska at this time (*126*).

**Hypothesis Four: Late-Holocene Siberian Origin**: The ancestral language was brought across Bering Strait during the past 2000 years by an Asian population closely related to that involved in Hypothesis Three. This scenario is applicable only for Eskaleut and probably only for its Yupik-Inuit branch, if Aleut had already been brought into Alaska by the migration from Siberia that occurred ~5000 yBP. See a discussion of the back and forth movement of Eskaleut ancestors across the Bering Strait and the special position of Aleuts in Flegontov *et al.* (*11*).

Genetics and archaeology thus indicate four distinct possibilities for the origin of language families spoken by NNA populations in North America. How can findings from comparative linguistics help choose between them? Identifying language genealogies across Bering Strait would narrow down these possibilities. Success in demonstrating cross-Bering language relationships using the Comparative Method, assumed not applicable at time depths earlier than the Holocene, would support a Holocene-Era entry into the Americas for both Na-Dene and Eskaleut. Na-Dene shows increasingly robust evidence of relatedness to the Yeniseian language family in Siberia (Dene-Yeniseian), while Eskaleut appears related to Uralic and Yukaghir in what has been called the Uralo-Siberian family (*126*). The fact that Eskaleut and Na-Dene have never plausibly been related either to one another or to any of the other primary families spoken in the Americas could be explained if both Eskaleut and Na-Dene were brought into Alaska during the Mid or Late Holocene (*126*). A firm demonstration of Dene-Yeniseian and Uralo-Siberian would support a mid-Holocene entry into Alaska of proto-Na-Dene and Proto-Eskaleut and would also attest to the unrelatedness of each of these two families to other indigenous North American families, whose speakers share no Mid-Holocene Siberian gene flow, except where due to late admixture with adjacent Tlingit, Dene, or Inuit-speaking populations. For the remaining five (or possibly seven) primary language families spoken by populations with NNA genes, the data from genetics do not reveal a Holocene-era entry and therefore indicate only a choice between Hypothesis One and Hypothesis Two.

The Trans-North-Pacific Hypothesis (Hypothesis Two) is intriguing since a time depth of ~16,000 years rather than ~23,000 years holds more potential for tracing relatives in the Old World. Of languages spoken by contemporary Asian peoples with a human genetic link to NNA populations, Japanese, Korean and Tungusic show a deep typological patterning with the so-called Altaic (or Uralo-Altaic) band of language families that also encompasses Mongolic and Turkic. None of these languages show structural affinities with North American languages, and it is plausible that the Late Pleistocene Amur River Basin ("Houtaomuga") genetic component present in contemporary Japanese, Korean and Tungusic populations represents a substrate unconnected with the origin of the languages these populations speak today. Only Nivkh and Chukotko-Kamchatkan, which themselves may be genealogically related (*36, 127*), remain as possible candidates for a distant relationship to languages spoken by NNA populations. Both families have already been investigated by a number of linguists working independently of one another as potential relatives to Algic, Salish, Wakashan, and possibly also Chimakuan and Kutenai (*108, 127-129*). It is not clear whether Chimakuan and Kutenai speakers actually possess the NNA genetic component, but their locations make this plausible. The Chimakuan microfamily contains the closely related Quileute and Chemakum languages, both indigenous to Washington's Olympic Peninsula. The Kutenai (also spelled Ktunaxa or Kootenai) language isolate is spoken in Montana and adjacent areas of Canada. Kutenai has been plausibly but not definitively linked to Salishan based on both lexical and morphological evidence (*130*). Dryer (*131*) investigated Kutenai parallels to Algic morphosyntactic patterns that may be due to deep genealogical relatedness. Another isolate occasionally claimed to be related to Algic is the extinct Beothuk of Newfoundland (*132*). Nothing definitive has yet been published about Beothuk genetics, but the scant attestations of Beothuk vocabulary do not indicate a relationship with Algonquian. In any event, the homeland of Algic speakers is more plausibly located far to the east, near the Columbia Plateau in the area where there is a primary split between Blackfoot (Siksika) and the remaining Algonquian languages (*133*). The Siouan languages of the upper Great Plains likely have their ancestral homeland in southeastern North America, far from NNA populations. Finally, the various Penutian languages of eastern and southern Washington State do not pattern with Pacific Northwest coastal language families in most regards and instead appear connected with languages farther south in Oregon and interior California. Siouan,

Penutian, Iroquoian and other language families spoken farther south in North America do not appear to have any genealogical link with languages spoken by NNA populations.

What linguistic features could have survived for 16,000 years that might indicate a genealogical relationship between languages spoken by NNA populations in North America and Nivkh or Chukotko-Kamchatkan languages in northeastern Asia? The Comparative Method relies on a combination of evidence from cognate vocabulary and homologous grammatical systems, unified by a system of interlocking sound correspondences. Claims of genealogical relatedness between languages based entirely on resemblances in basic vocabulary or on typological similarities without cognate morphological combinations cannot be regarded as convincing evidence of genealogical language relatedness (*134*). As mentioned already, if by eight to ten thousand years the pace of language change generally erodes the quantity of evidence needed to fully demonstrate language relatedness, it might simply not be possible to prove linguistic genealogy deeper than the Holocene, even if some evidence of language relatedness did survive from that time. However, if at least partial evidence of this linguistic link survives in contemporary languages, what might this be? It is likely that some cognates in basic vocabulary roots have survived in detectable form among languages with Pleistocene-era genealogical relationships. But finding them based purely on phonetic resemblance without any overarching morphological comparison is likely to yield either too many false matches (due purely to coincidence or, in some cases, to ancient borrowing) resulting in too few positive matches to posit a verifiable system of sound correspondences. Even when examining basic vocabulary, sorting out false lookalikes from true cognates necessarily requires a convincing and systematic morphological analysis to justify the division of root morphemes from fused affixal elements. For this reason, sound correspondences calculated from putative cognates in basic vocabulary without parallel evidence of an inherited morphological system have never proven a single language family. A recent study of primary families in northern Asia (*135*), concluded that grammatical homologies are in fact more persistent over time than phonological and lexical patterns.

When attempting a preliminary hypothesis of possible distant language relationship (either areal or genealogical) it is therefore preferable to search first for shared morphological patterns of the type explored by Nichols (*124*) and Fortescue (*108*). These include unusual combinations of morphosyntactic traits (like the rare combination of prefixes with SOV syntax and postpositions shared between Yeniseian and Na-Dene). Other potential evidence includes first- and second-person pronoun configurations such as the M/T pattern in Eurasia or the N/M pattern in the western portions of the Americas. Both patterns could possibly be remnants of ancient linguistic genealogies. Another possibility includes shared inflectional systems of case endings, tense/aspect/mood affixes, or other idiosyncratic homologies in complex word structure, including specific types of polysynthetic verb structures. Though this sort of "typological-comparative" approach cannot replace the Comparative Method in proving genealogical relationship, its results can furnish preliminary and speculative support for certain possibilities over others and point the way toward more intensive comparison that includes both vocabulary and morphological systems.

As already mentioned above, Japanese, Korean, and the Tungusic languages pattern typologically with Turkic and Mongolic that are spoken farther west in Eurasia, rather than with Native American families. All of these families are strongly suffixing and agglutinating; they allow strings of inflectional suffixes in which each morpheme expresses a single, separate grammatical meaning (such as ‘plural', or ‘locative'), unlike the fusional Indo-European languages, where words have one at most two inflectional suffixes, usually with multiple meanings (such as ‘singular' + ‘locative' being fused in a single suffix and ‘plural' + ‘locative' fused in a completely different suffixal form). These shared patterns arose at least in part due to extensive language contact but may also reflect distant genealogical relationship between some or all of these language – a problem still not resolved by historical linguistics. The B/S pattern of first- and second-person singular personal pronouns shared between Tungusic, Mongolic and Turkic has never been adequately explained as arising either from language contact or through sheer coincidence. The Altaic B/S pattern, in turn, has been linked genealogically with the M/T pattern shared by Indo-European, Uralic, Yukaghir, Eskaleut and possibly Chukotko-Kamchatkan. Languages of North America's Subarctic regions or Pacific Northwest Coast show no discernable affinities with any of these languages. The most plausible conclusion is that the Amur River Basin genetic component in Asian populations speaking Japanese, Korean, or Tungusic in the latter groups is a genetic substrate not historically connected with their language origins.

The situation is quite different with Nivkh and Chukotko-Kamchatkan. Both Nivkh and Chukotko-Kamchatkan are polysynthetic, "Paleosiberian" languages with verb structures that display typological affinities with certain languages spoken in North America. Neither of these primary language families fit the broader Uralo-Altaic suffixal-agglutinating type, though both share oblique case systems that pattern typologically with other Inner and Northern Asian languages. Chukotko-Kamchatkan also appears to exhibit the M/T pronoun pattern that appears to tie it to Uralic, Yukaghir and Eskaleut, though other forms of first- and second-person pronoun markers also exist in the family and it is not clear which should be reconstructed as primary. There has been extensive areal contact between Eskaleut and Chukotko-Kamchatkan, presumably occurring in the Late Holocene (*126*) that must be sorted out before attempting a deeper comparison with Nivkh or languages of northwestern North America. Fortescue (*36*) has already worked out a promising body of preliminary evidence for a genealogical connection between Nivkh and Chukotko-Kamchatkan that includes lexical cognates, grammatical homologies and the beginnings of systematic sound correspondences. Importantly, none of this putative evidence can be found in Yeniseian (also considered a "Paleosiberian" language) or in Na-Dene; instead, both of these families share a completely different type of polysynthetic verb structure with a prefixing templatic grid that contrasts starkly with that of all other families in Northern Asia and North America (*136*). Similarities between Nivkh and Chukotko-Kamchatkan shared with North American languages thus skip over both Eskaleut and Na-Dene, a fact that is explainable if the latter two families entered North America during the Mid- or Late Holocene. The comparison with languages spoken by the remaining NNA populations undertaken below will consider Nivkh and Chukotko-Kamchatkan together, though these two families have yet to be fully proven as genealogically related.

The most promising possibility of determining a linguistic parallel to the Late Pleistocene gene flow into NNA populations concerns Nivkh and Chukotko-Kamchatkan on the Asian side and Algic and Salishan in North America. The resulting hypothesis of genealogical relationship could be called "the Trans North Pacific Hypothesis" or "Trans North Pacific Phylum", with the use of the word "phylum" indicating a time depth that might not be amenable to conclusive demonstration using the Comparative Method. Several important prior studies have already been published working toward this goal. As earlier mentioned, Fortescue (*36*) has made a plausible case that Chukotko-Kamchatkan languages could be distantly related to Nivkh, citing morphological homologies as well as lexical cognates with the beginnings of a system of sound correspondences. Bakker (*137*) describes a range of idiosyncratic structural homologies between Algic and Salishan, some of which also subsume Kutenai. Nikolaev (*128, 129*) has compiled an impressive list of potential lexical cognates shared between Algic, Salishan and Nivkh, upon which basis he posits a system of interlocking sound correspondences. Lexical correspondences with Chukotko-Kamchatkan, however, are identified by Nikolaev as ancient loanwords rather than potential cognates. Although none of this work has yet conclusively proven the relationship between any of these primary families, it is instructive to note that all of it was conducted before the discovery of any human genetic connection between the populations speaking these languages, in much the same way that the Dene-Yeniseian linguistic hypothesis was formulated prior to the discovery of a Siberian genetic component shared between modern Kets and Na-Dene speaking populations.

Other Pacific Northwest language families spoken by populations with NNA genes are less likely to be connected with this hypothesized "Trans North Pacific Phylum". Wakashan and Chimakuan are known to form a complex *Sprachbund* with Salishan, with intensive language contact between the three groups going back many centuries (*138*). The idea that these three language families are genealogically related, i.e. Sapir's "Mosan" hypothesis (*139*), has been almost universally abandoned. Nikolaev (*128, 129*) claims that Wakashan and Chimakuan are genealogically related to Nivkh and Salish-Algic at a deeper time level, but such a genealogy would contradict the known human genetic and archaeological parallels. If all of these languages are genealogically related, the families on the North American side would almost certainly form a taxon together, as would Nivkh and Chukotko-Kamchatkan on the Asian side. The status of Wakashan and Chimakuan remains that of two separate primary families linked with Salishan (and to a lesser extent with other Pacific Northwest language families) through a pervasive areal contact.

The two remaining primary families of the greater Pacific Northwest language area are Tsimshianic and Haida. Both appear to be genealogically unrelated to one another as well as to the other families of the region. The inclusion of Haida in Na-Dene has almost been universally abandoned, with shared features between Haida and Tlingit better explained as arising through long-term contact. The suffixing Haida verb is fundamentally different than the prefixing Na-Dene (or deeper Dene-Yeniseian) finite verb complex, with its many shared idiosyncratic homologies (*136*). Tsimshianic has been hypothesized to be somehow connected to the Penutian languages spoken farther south in western North America. One shared feature of these languages is the N/M pronoun pattern found widely in western portions of the Americas. If this pattern is indeed a relic of distant language relatedness in some or all of the primary families in which it appears, the language family (or phylum) in question would probably date back to the end of the Pleistocene, possibly spreading in connection with the Clovis Culture, though other scenarios are plausible, as well. In any event, Tsimshianic and Haida appear to be relics in the Pacific Northwest that descend from languages originating in ancient Beringia rather than brought from Northeast Asia in the Late Pleistocene. The same also may be true of Wakashan and Chimakuan. The ancestors of all of these languages may have been spoken by NNA populations that did not originally mix with these Asian newcomers, the mixing of genes (as well as linguistic traits) occurring only later in the broader Pacific Northwest convergence zone.

Table S24 below presents a theoretically plausible layering of language connections across Bering Strait. The term "family" is used for Holocene-Era genealogical units that might be proven using the traditional Comparative Method, while the term "phylum" refers to hypothetical unity dating back to Pleistocene times that may not be fully demonstrable.

Table S24. Hypothesized Cross-Bering Linguistic Layers (youngest to oldest)

A. Uralo-Siberian Family, containing Uralic, Yukaghir, Eskaleut (historically connected with Mid- to Late-Holocene gene flow out of northeastern Asia)
B. Dene-Yeniseian Family, linking Yeniseian with Tlingit, Eyak and Dene languages (associated with Mid-Holocene gene flow out of central Siberia)
C. Trans North Pacific Phylum, linking Nivkh and Chukotko-Kamchatkan with Algic and Salishan, and possibly also with Wakashan, Chimakuan, and/or Kutenai (historically connected with the hypothetical Late Pleistocene gene flow between proto-Siberians and proto-Americans that happened in Siberia or in Beringia)
D. Languages descending from phyla originating in the Pleistocene Beringian Standstill, including Tsimshianic, Haida, and all language families south of the Pacific Northwest (presumably the ancestors of all of these families have Ancient Beringian origins with possible deeper connections either to ANE or Ancient East Asians)

The languages subsumed under D in Table S24 are presently classified into at least 110 primary families that cannot plausibly be linked to any known Holocene or Late Pleistocene migration out of Eurasia. These languages, with roots in the Beringian Standstill, might represent a single phylum or multiple phyla, depending upon how many distinct proto-languages were brought south into the Americas during the Late Pleistocene. The sheer typological diversity of documented families that are not connectable to groups A, B or C in Table S24, together with the multiple distinct folklore connections with Old World groups observed between the peoples who speak them (*125*), suggests that ancestral Native Americans populations brought several proto-languages into the Americas from an earlier source in ancient Beringia. Furthermore, these languages may have had completely distinct origins in ancient northern Eurasia or East Asia. In any case, the existence of a Trans North Pacific Phylum with its distinct, non-Beringian origin would already demonstrate the untenability of Greenberg's (*132*) Amerind Hypothesis, which claimed that all indigenous New World languages except Eskaleut and Na-Dene (into which Haida was erroneously included) comprised a single phylum.

Although linguistics cannot provide the degree of confidence found in archaeological and human genetic studies of ancient prehistory dating back to Pleistocene times, it still can contribute something to the emerging synthesis about the early peopling of the Americas. The language layering across Bering Strait described here involves the logical consideration of speculative possibilities rather than actual evidence or proof. The foregoing discussion has modeled a variety of logically possible scenarios for how distinct population movements out of Beringia or Asia could be reflected in North America's linguistic diversity. In particular it has identified a possible linguistic parallel matching the gene flow between proto-Siberians and proto-Americans ~16,000 yBP with what has been called the "Trans North Pacific language phylum" encompassing Nivkh and Chukotko-Kamchatkan on the Asian side and Salishan and Algic (as well as possibly Kutenai, and less likely also Wakashan and Chimakuan) in North America. This hypothesis could also explain the existence of shared Trans North Pacific folklore motifs such as the Raven Creator. The degree of affinity in both folklore and language typology would be difficult to explain as arising either from Holocene-Era contact, for which there is no evidence, or surviving from Ancient Beringian times, which would not explain the discrete patterning on both sides of the North Pacific Rim.

Finally, it should be noted that there is no evidence that Eskaleut, Na-Dene, or Algic are genealogically related. The stark differences between these major language families spoken by populations with the NNA genetic component argues against their ancestral unity and favors the scenario of multiple Asian origins for the populations and languages indigenous to northern North America.

**Fig. S1.**

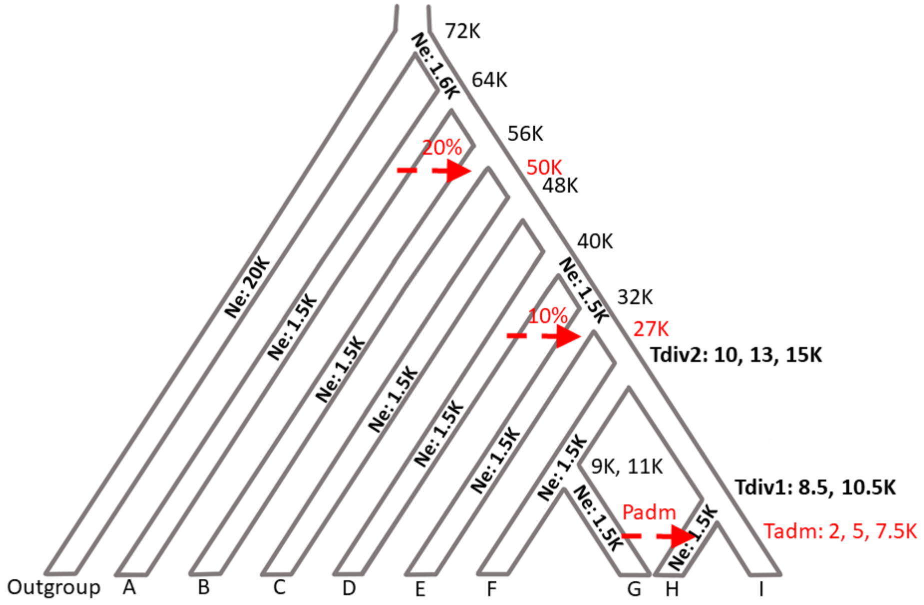


A simulated "information-poor" demographic history for 10 groups (5 individuals each) used for testing the sensitivity of *qpGraph*, *qpAdm*, and *D*-statistics to gene flow. Effective population sizes ($N_e$) in thousands of individuals are shown on each branch. The time in thousands of years ago (K) is shown on the right. Gene flow events are shown in red. The following divergence and admixture dates were variable: the divergence of the FG and HI clades ($T_{div2}$, 10, 13, and 15 ka), the divergence of the F and G populations ($T_{div1}$, 8.5, 10.5 ka), the G=>H gene flow ($T_{adm}$, 2.5, 5, and 7.5 ka). Populations F, G, H, and I form the target clade, and H = I + G is the target admixture event. Only one population is differentially related to I and G, that is F. The simulated admixture proportions were variable: 0%, 5%, 10%, 20%, 30%, 40%, 50%, 60%, 70%, 80%, 90%, 95%, and 100% G ancestry in H.

**Fig. S2.**

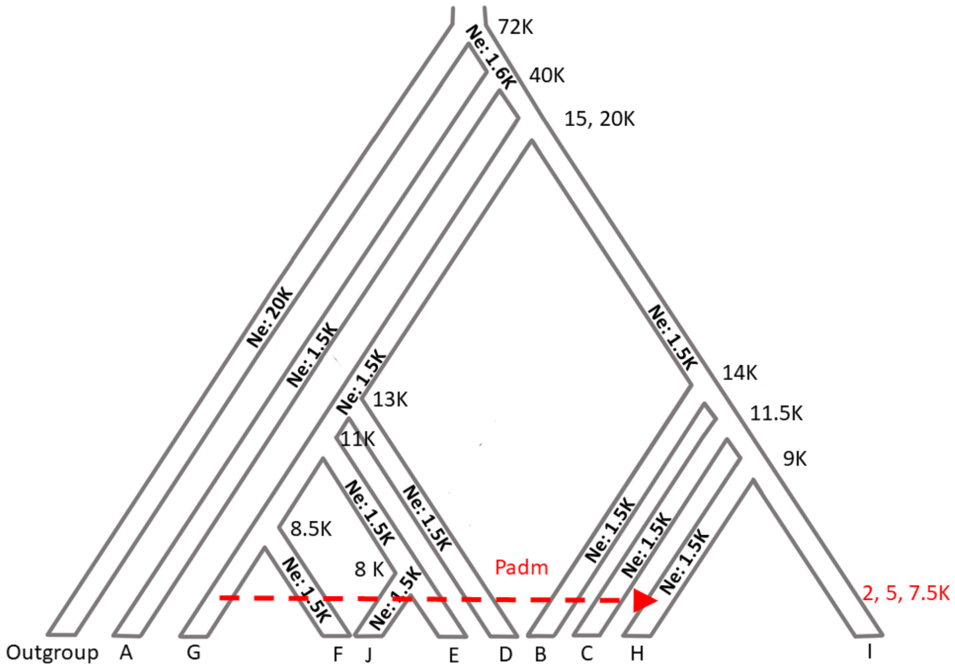


A simulated "information-rich" demographic history for 11 groups (5 individuals each) used for testing the sensitivity of *qpGraph*, *qpAdm*, and *D*-statistics to gene flow. Effective population sizes ($N_e$) in thousands of individuals are shown on each branch. The time in thousands of years ago (K) is shown next branch splits. The gene flow event is shown in red. The following dates were variable: the divergence of the DEFGJ and BCHI clades ($T_{div2}$, 15 or 20 ka) and the G=>H gene flow ($T_{adm}$, 2.5, 5, and 7.5 ka). Populations B through J form the target clade, and H = I + G is the target admixture event. Six populations are differentially related to I and G, those are B, C, D, E, J, and F. The simulated admixture proportions were variable: 0%, 5%, 10%, 20%, 30%, 40%, 50%, 60%, 70%, 80%, 90%, 95%, and 100% G ancestry in H.

**Fig. S3.**

**A** 95K sites

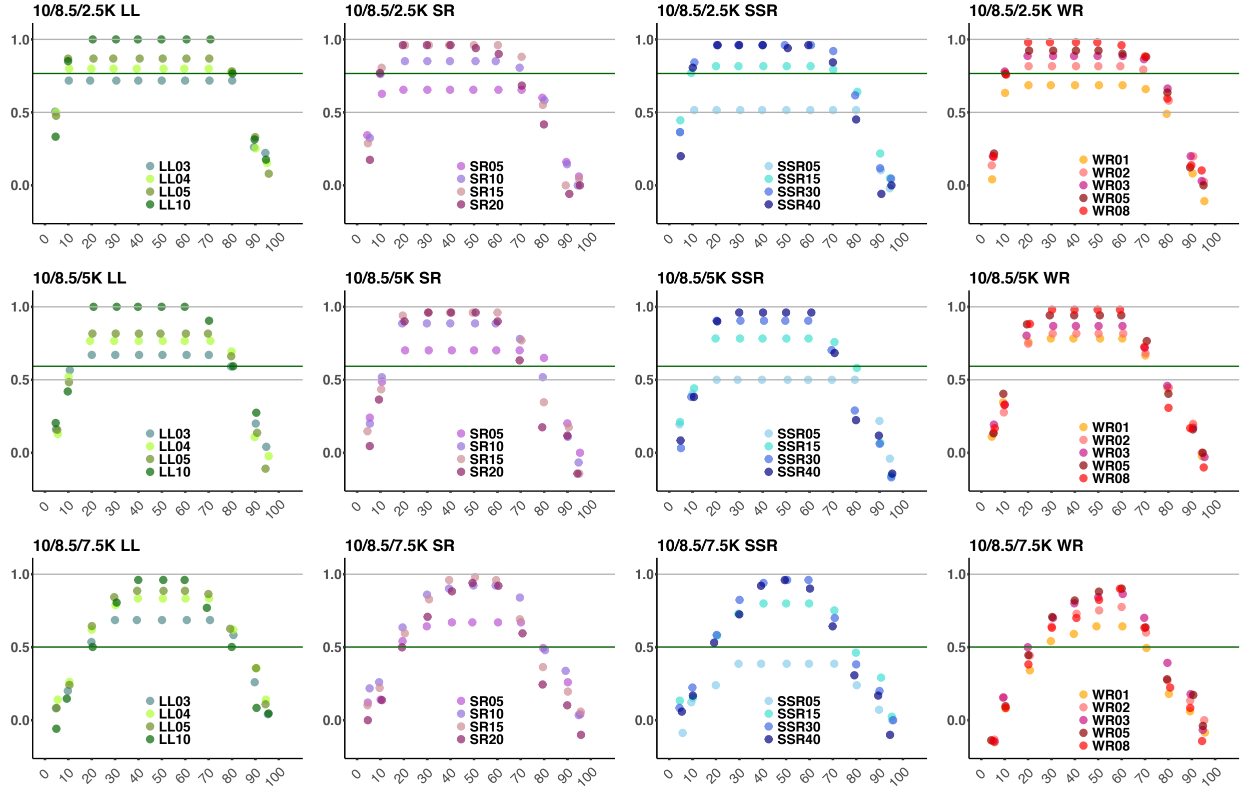


**B** 400K sites

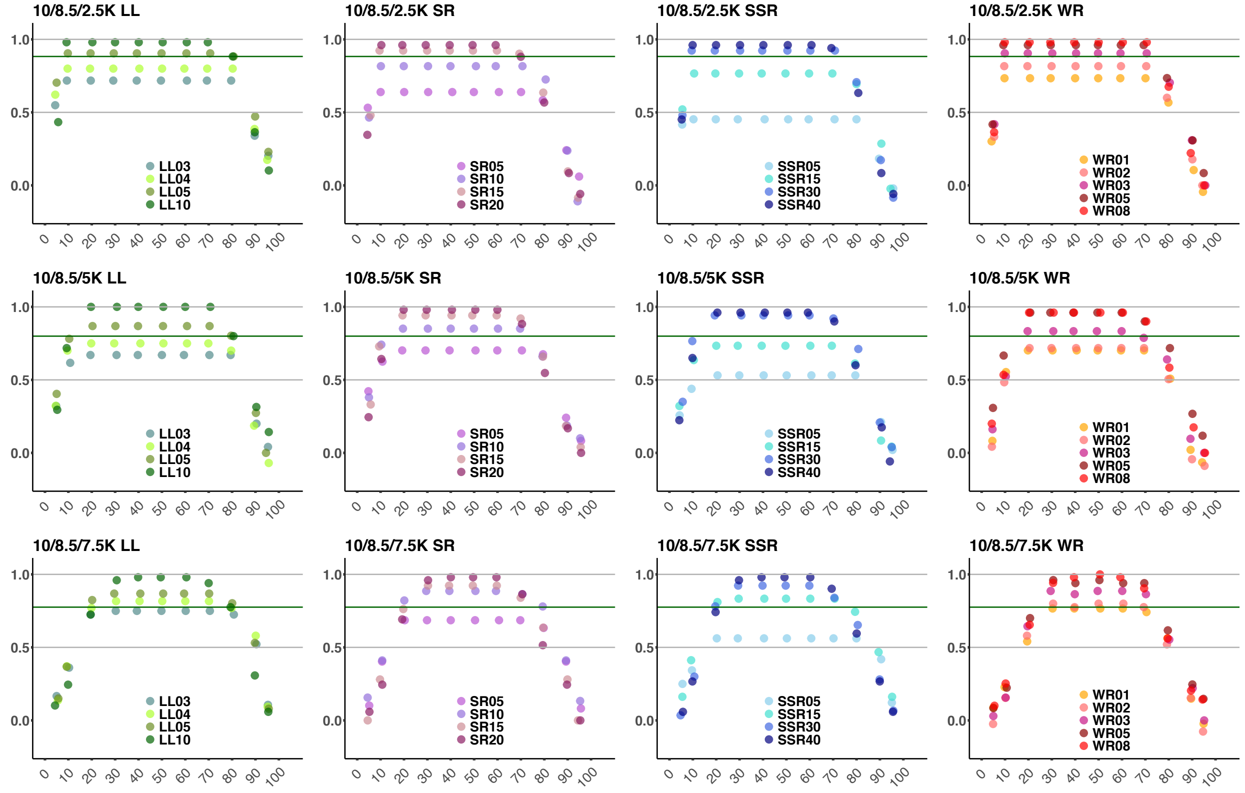

**C** 95K sites

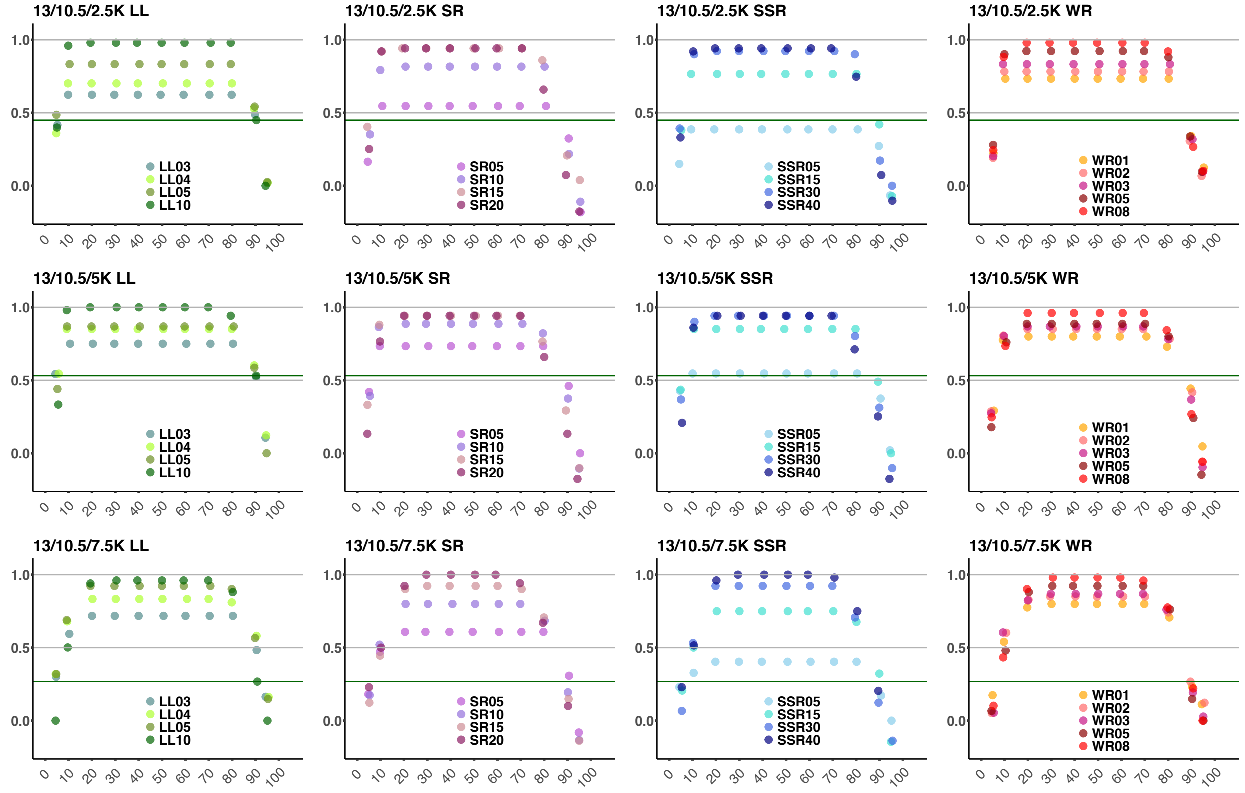


**D** 400K sites

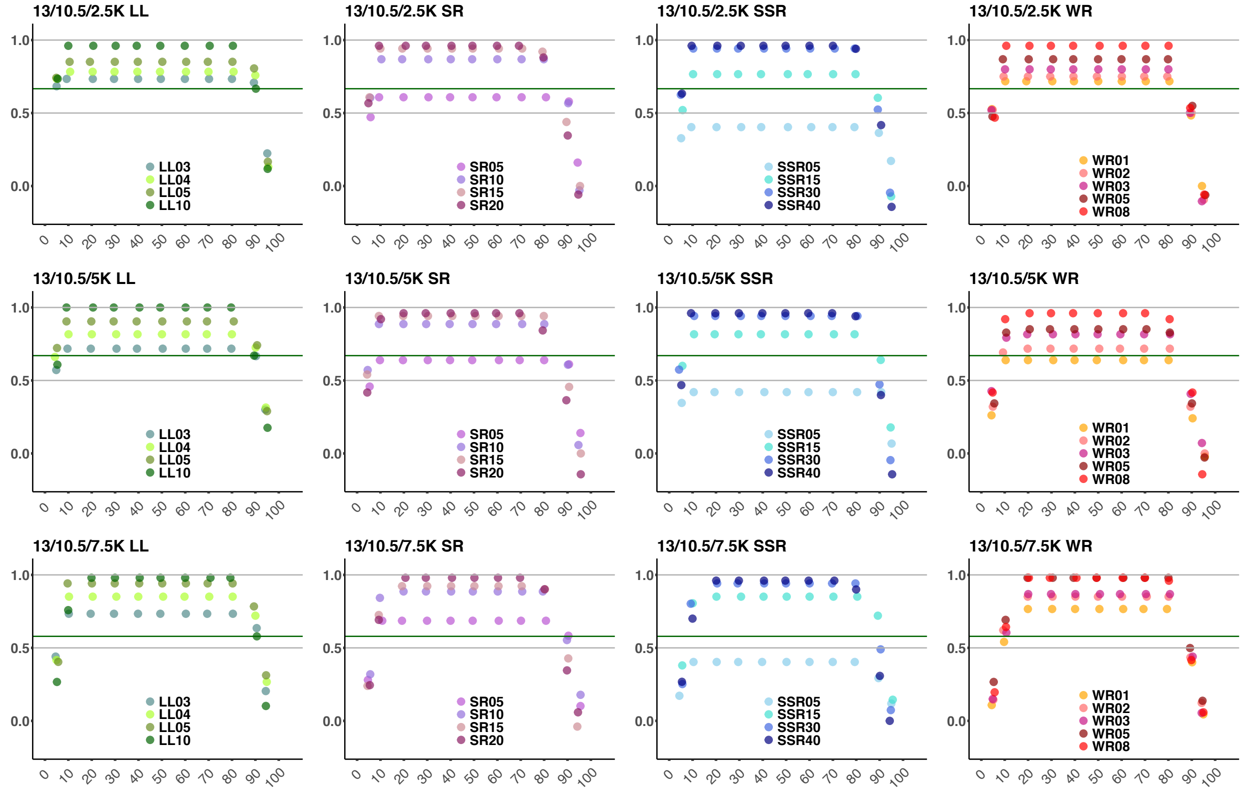

**E** 95K sites

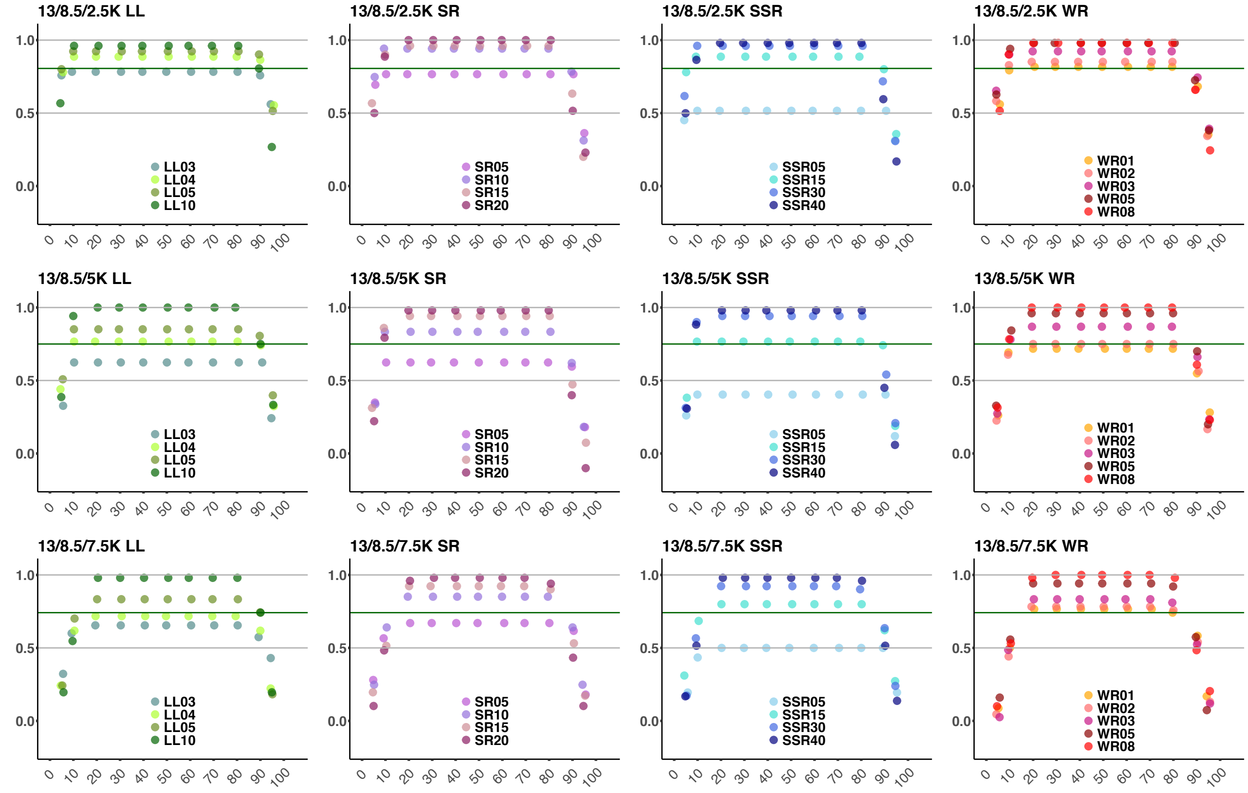


**F** 400K sites

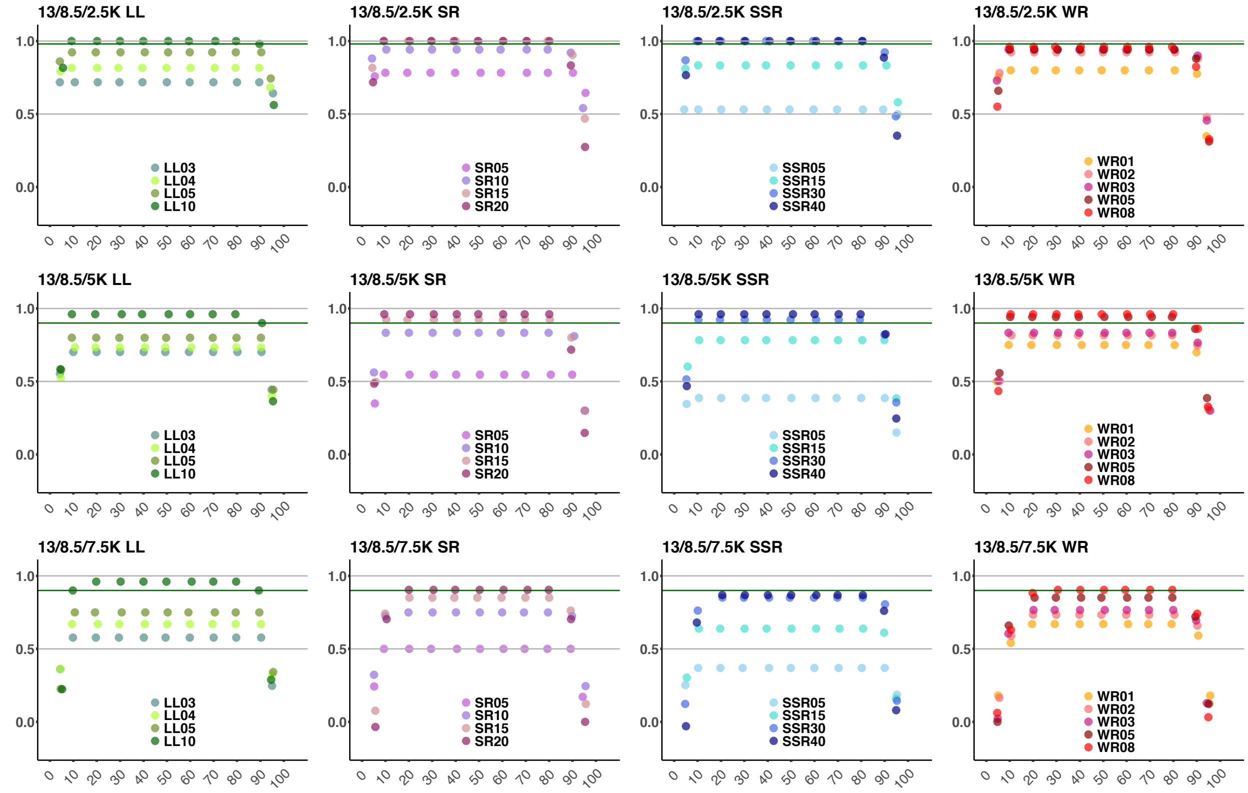

**G** 95K sites

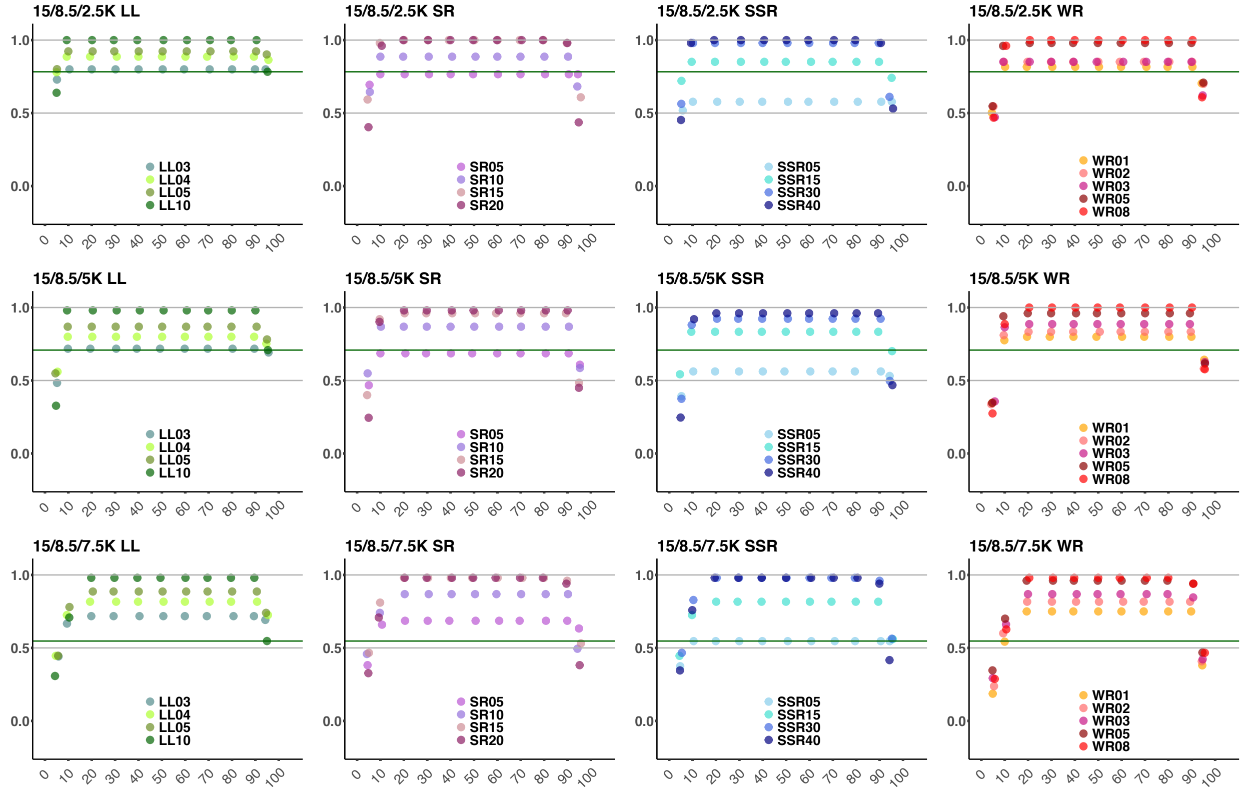


**H** 400K sites

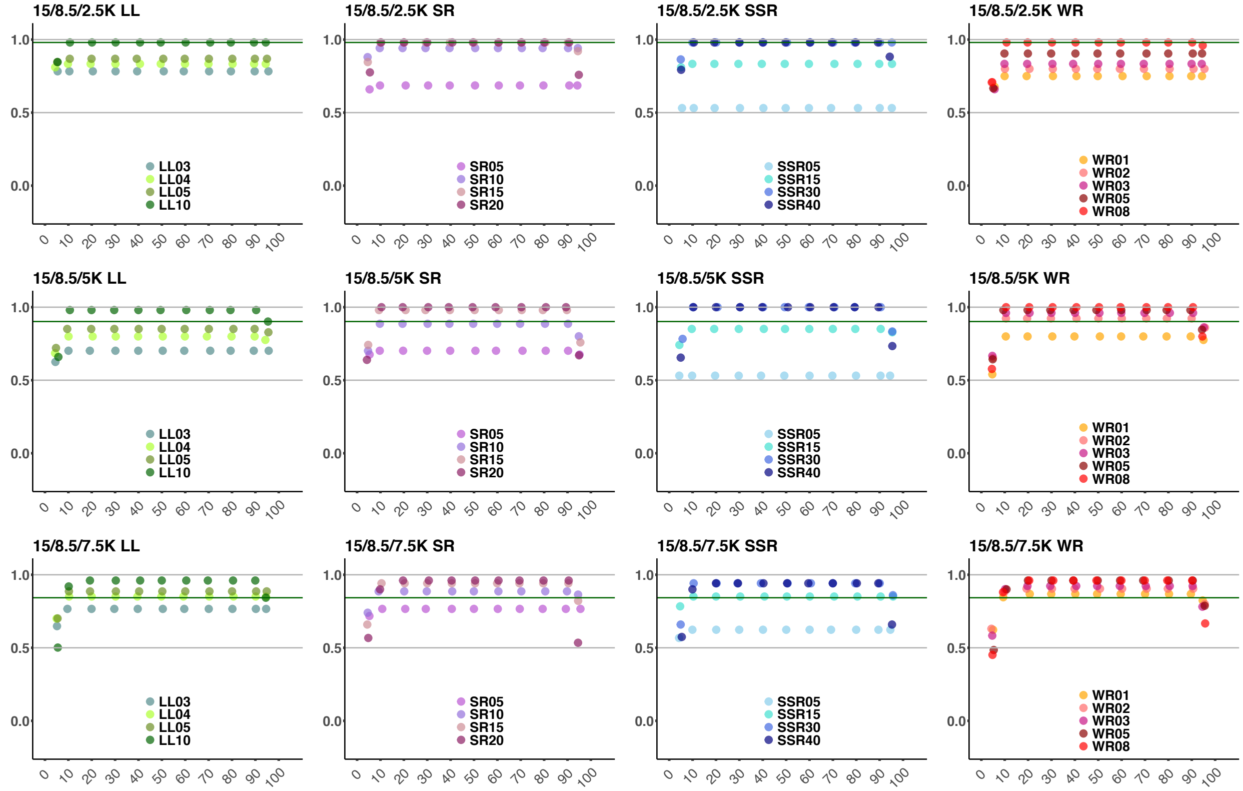

Performance of various classifiers color-coded according to the legends under the "mapping" setup, "information-poor" graphs, two datasets (**A,C,I,G**, 95,000 sites; **B,D,F,H**, 400,000 sites), and four divergence date combinations (**A,B**, 10 and 8.5 ka; **C,D**, 13 and 10.5 ka; **E,F**, 13 and 8.5 ka; **G,H**, 15 and 8.5 ka). The following threshold values were used: 3, 4, 5, or 10 ln-units difference for the LL classifier (abbreviated as "LL03", "LL04", etc.); 5, 10, 15, or 20% difference for the SR classifier (abbreviated in a similar way); 5, 15, 30, or 40% difference for the SSR classifier; 0.1, 0.2, 0.3, 0.5, or 0.8 SE intervals for the WR classifier. The plots show simulated gene flow proportions between 5% and 95% (x-axis) vs. MCC (y-axis). The grey lines show classifier performance that was considered acceptable (MCC between 0.5 and 1), and the dark green lines are drawn to make comparison of the classifiers easier. The green lines mark MCC values for the LL10 classifier under certain "non-optimal" intensities of gene flow where performance of classifiers was highly variable. MCC was undefined in some cases due to division by 0.

**Fig. S4.**

**A**

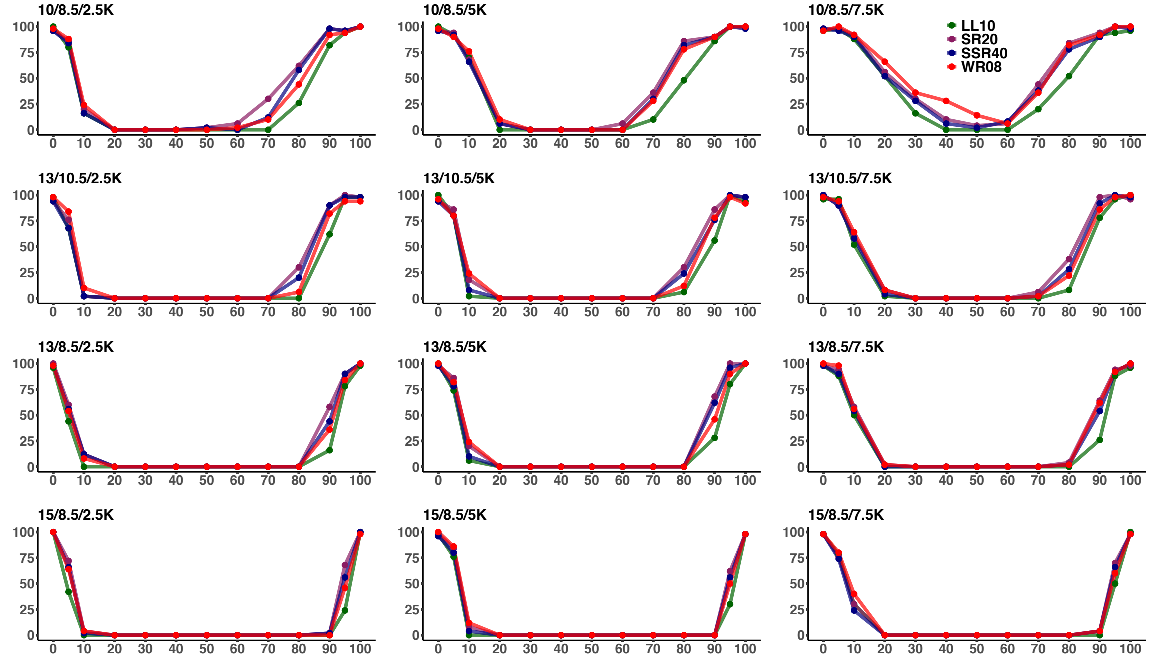


**B**

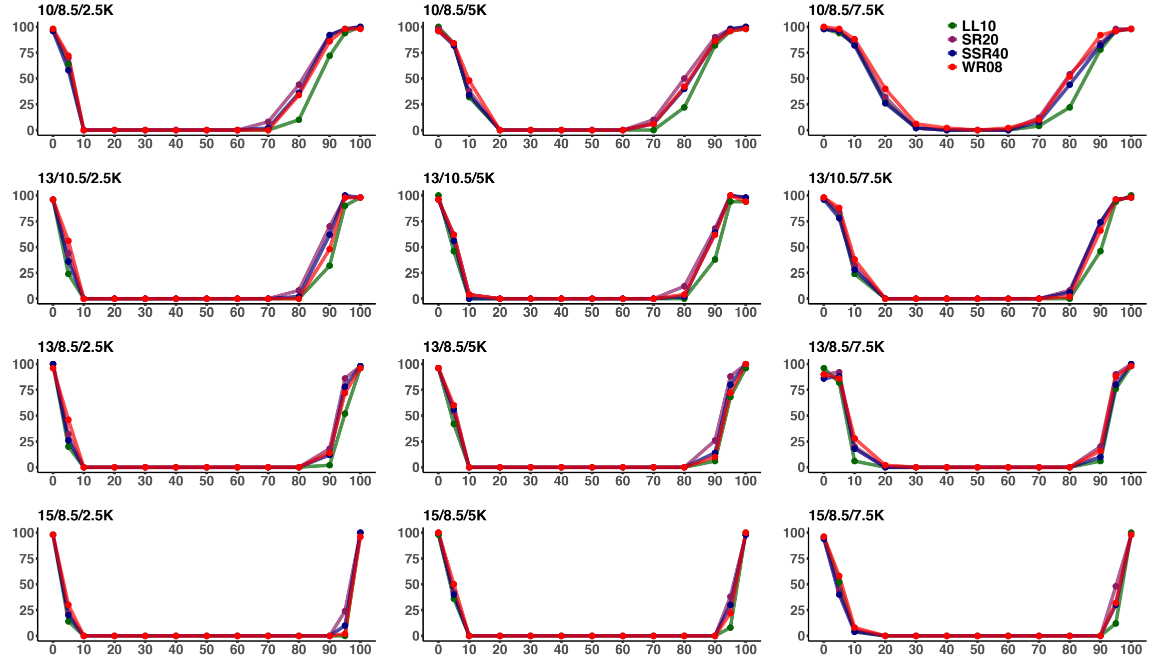


The percentage of iterations ("information-poor" graphs) that gave a negative result: i.e. where the highest-ranking unadmixed model for group H was not worse as compared to the highest-ranking pair of sources according to four alternative metric/threshold combinations abbreviated as LL10 (dark green), SR20 (dark violet), SSR40 (dark blue), and WR08 (red). The settings, i.e. combinations of admixture and divergence dates in the clade of interest in thousands of years ago, are indicated above each plot. The following datasets were used: (**A**) 95,000 sites; (**B**) 400,000 sites.

### Fig. S5.

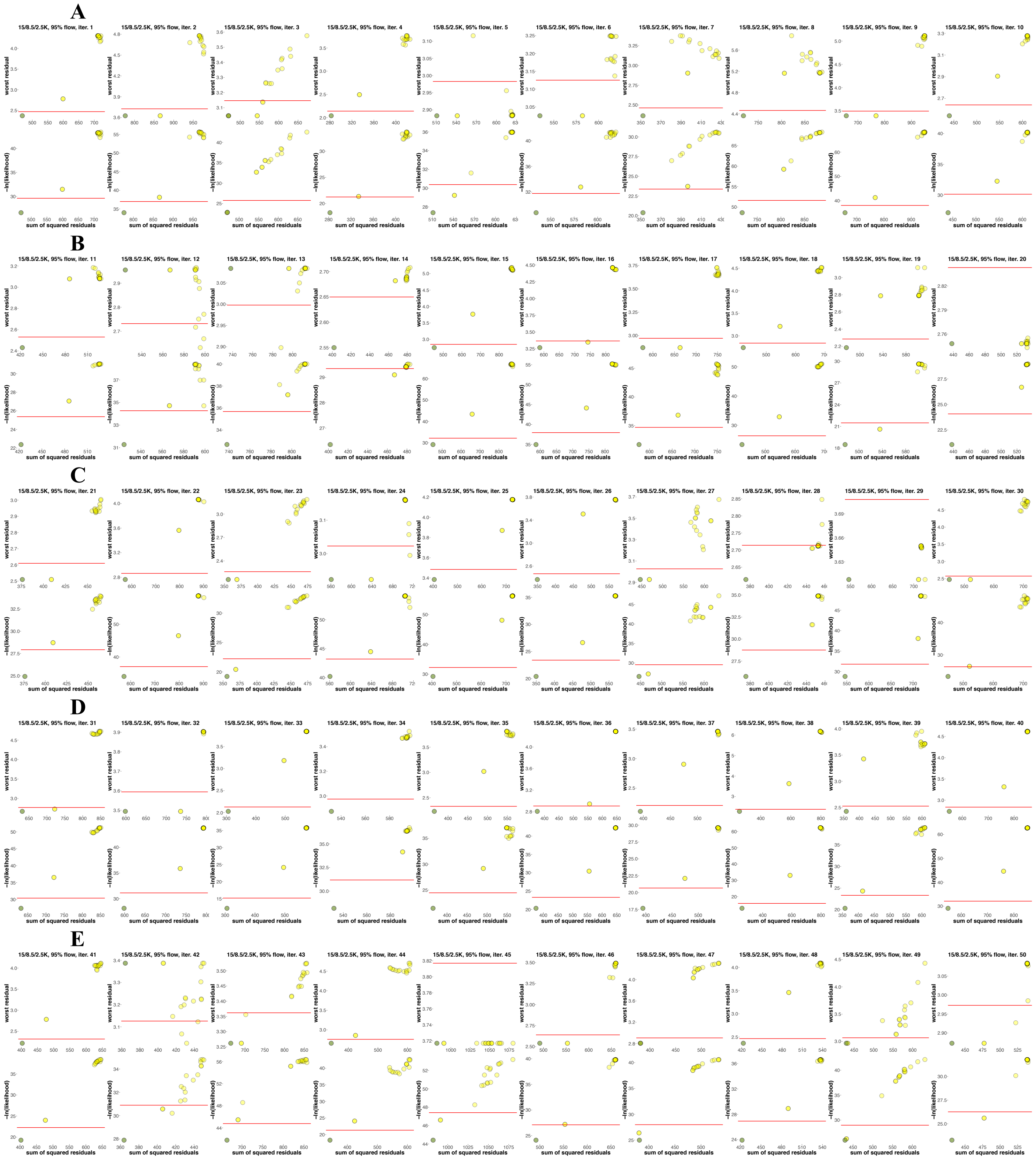
A
15/8.5/2.5K, 95% flow, iter. 1
15/8.5/2.5K, 95% flow, iter. 2
15/8.5/2.5K, 95% flow, iter. 3
15/8.5/2.5K, 95% flow, iter. 4
15/8.5/2.5K, 95% flow, iter. 5
15/8.5/2.5K, 95% flow, iter. 6
15/8.5/2.5K, 95% flow, iter. 7
15/8.5/2.5K, 95% flow, iter. 8
15/8.5/2.5K, 95% flow, iter. 9
15/8.5/2.5K, 95% flow, iter. 10
B
15/8.5/2.5K, 95% flow, iter. 11
15/8.5/2.5K, 95% flow, iter. 12
15/8.5/2.5K, 95% flow, iter. 13
15/8.5/2.5K, 95% flow, iter. 14
15/8.5/2.5K, 95% flow, iter. 15
15/8.5/2.5K, 95% flow, iter. 16
15/8.5/2.5K, 95% flow, iter. 17
15/8.5/2.5K, 95% flow, iter. 18
15/8.5/2.5K, 95% flow, iter. 19
15/8.5/2.5K, 95% flow, iter. 20
C
15/8.5/2.5K, 95% flow, iter. 21
15/8.5/2.5K, 95% flow, iter. 22
15/8.5/2.5K, 95% flow, iter. 23
15/8.5/2.5K, 95% flow, iter. 24
15/8.5/2.5K, 95% flow, iter. 25
15/8.5/2.5K, 95% flow, iter. 26
15/8.5/2.5K, 95% flow, iter. 27
15/8.5/2.5K, 95% flow, iter. 28
15/8.5/2.5K, 95% flow, iter. 29
15/8.5/2.5K, 95% flow, iter. 30
D
15/8.5/2.5K, 95% flow, iter. 31
15/8.5/2.5K, 95% flow, iter. 32
15/8.5/2.5K, 95% flow, iter. 33
15/8.5/2.5K, 95% flow, iter. 34
15/8.5/2.5K, 95% flow, iter. 35
15/8.5/2.5K, 95% flow, iter. 36
15/8.5/2.5K, 95% flow, iter. 37
15/8.5/2.5K, 95% flow, iter. 38
15/8.5/2.5K, 95% flow, iter. 39
15/8.5/2.5K, 95% flow, iter. 40
E
15/8.5/2.5K, 95% flow, iter. 41
15/8.5/2.5K, 95% flow, iter. 42
15/8.5/2.5K, 95% flow, iter. 43
15/8.5/2.5K, 95% flow, iter. 44
15/8.5/2.5K, 95% flow, iter. 45
15/8.5/2.5K, 95% flow, iter. 46
15/8.5/2.5K, 95% flow, iter. 47
15/8.5/2.5K, 95% flow, iter. 48
15/8.5/2.5K, 95% flow, iter. 49
15/8.5/2.5K, 95% flow, iter. 50
worst residual
−ln(likelihood)
sum of squared residuals

F

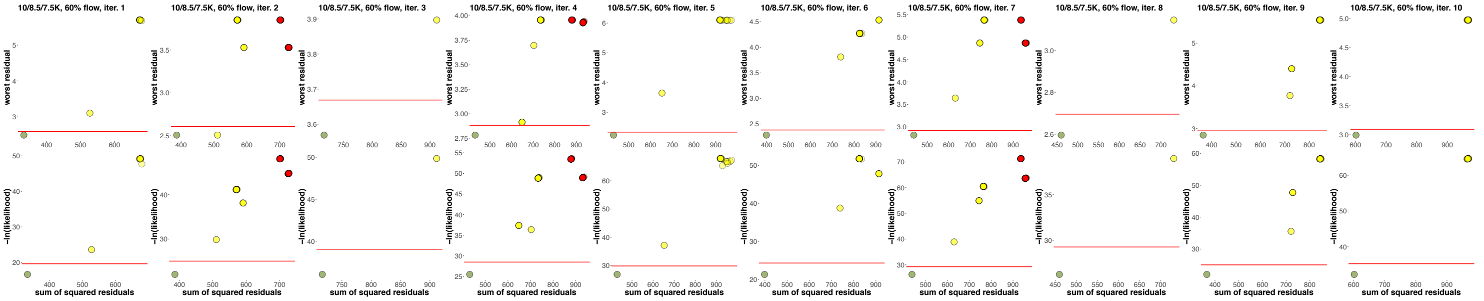

10/8.5/7.5K, 60% flow, iter. 1
10/8.5/7.5K, 60% flow, iter. 2
10/8.5/7.5K, 60% flow, iter. 3
10/8.5/7.5K, 60% flow, iter. 4
10/8.5/7.5K, 60% flow, iter. 5
10/8.5/7.5K, 60% flow, iter. 6
10/8.5/7.5K, 60% flow, iter. 7
10/8.5/7.5K, 60% flow, iter. 8
10/8.5/7.5K, 60% flow, iter. 9
10/8.5/7.5K, 60% flow, iter. 10
worst residual
−ln(likelihood)
sum of squared residuals


G

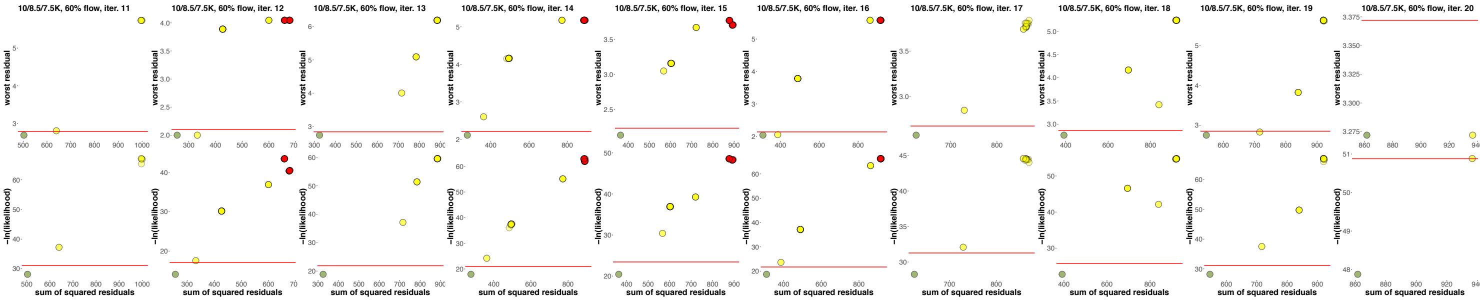

10/8.5/7.5K, 60% flow, iter. 11
10/8.5/7.5K, 60% flow, iter. 12
10/8.5/7.5K, 60% flow, iter. 13
10/8.5/7.5K, 60% flow, iter. 14
10/8.5/7.5K, 60% flow, iter. 15
10/8.5/7.5K, 60% flow, iter. 16
10/8.5/7.5K, 60% flow, iter. 17
10/8.5/7.5K, 60% flow, iter. 18
10/8.5/7.5K, 60% flow, iter. 19
10/8.5/7.5K, 60% flow, iter. 20
worst residual
−ln(likelihood)
sum of squared residuals


H

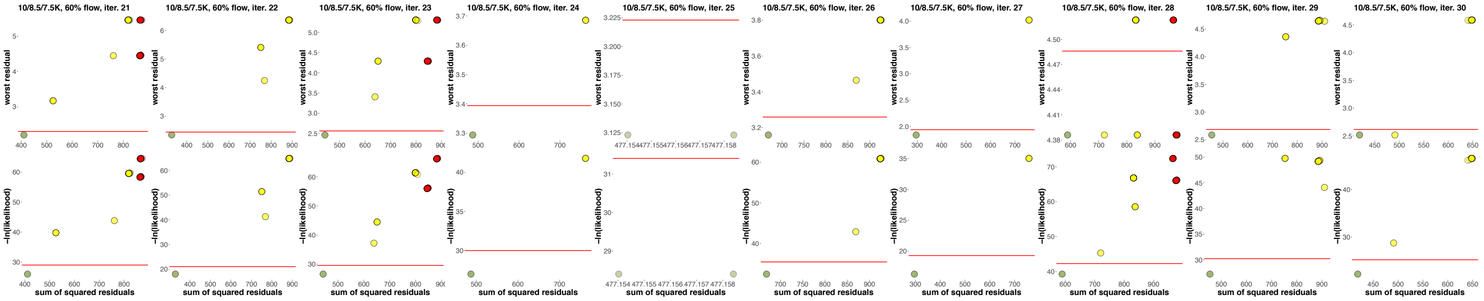

10/8.5/7.5K, 60% flow, iter. 21
10/8.5/7.5K, 60% flow, iter. 22
10/8.5/7.5K, 60% flow, iter. 23
10/8.5/7.5K, 60% flow, iter. 24
10/8.5/7.5K, 60% flow, iter. 25
10/8.5/7.5K, 60% flow, iter. 26
10/8.5/7.5K, 60% flow, iter. 27
10/8.5/7.5K, 60% flow, iter. 28
10/8.5/7.5K, 60% flow, iter. 29
10/8.5/7.5K, 60% flow, iter. 30
worst residual
−ln(likelihood)
sum of squared residuals


I

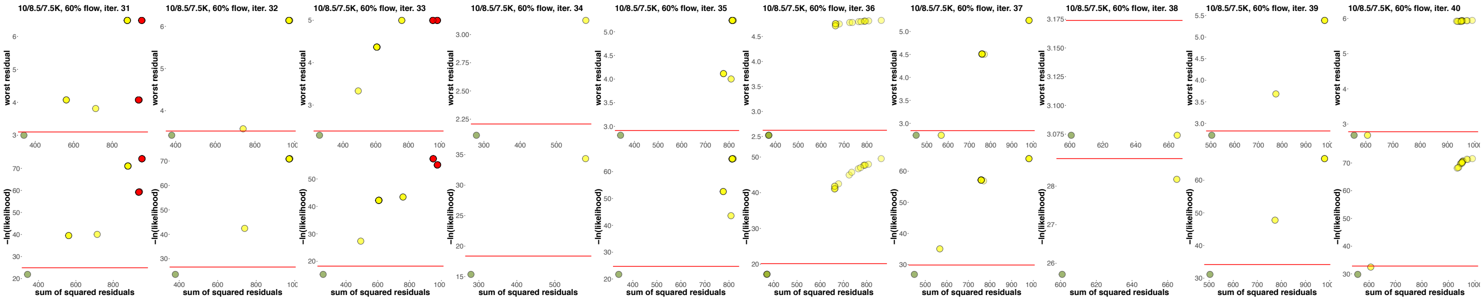

10/8.5/7.5K, 60% flow, iter. 31
10/8.5/7.5K, 60% flow, iter. 32
10/8.5/7.5K, 60% flow, iter. 33
10/8.5/7.5K, 60% flow, iter. 34
10/8.5/7.5K, 60% flow, iter. 35
10/8.5/7.5K, 60% flow, iter. 36
10/8.5/7.5K, 60% flow, iter. 37
10/8.5/7.5K, 60% flow, iter. 38
10/8.5/7.5K, 60% flow, iter. 39
10/8.5/7.5K, 60% flow, iter. 40
worst residual
−ln(likelihood)
sum of squared residuals


J

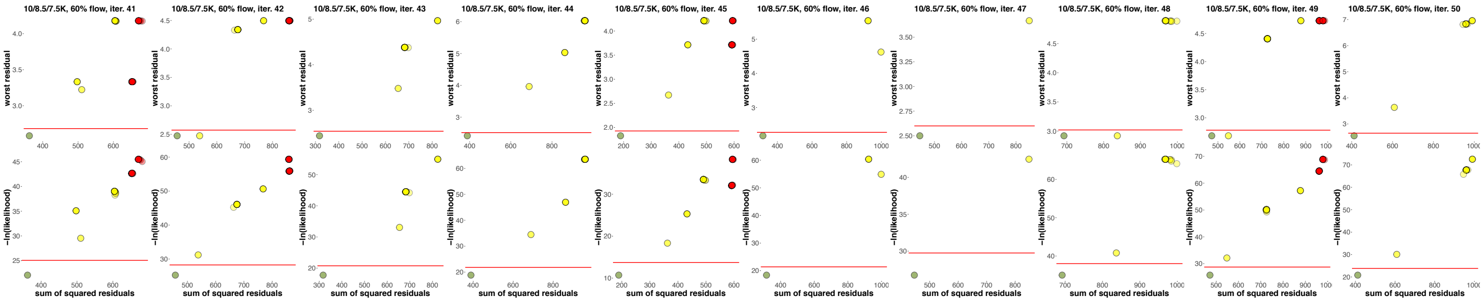

10/8.5/7.5K, 60% flow, iter. 41
10/8.5/7.5K, 60% flow, iter. 42
10/8.5/7.5K, 60% flow, iter. 43
10/8.5/7.5K, 60% flow, iter. 44
10/8.5/7.5K, 60% flow, iter. 45
10/8.5/7.5K, 60% flow, iter. 46
10/8.5/7.5K, 60% flow, iter. 47
10/8.5/7.5K, 60% flow, iter. 48
10/8.5/7.5K, 60% flow, iter. 49
10/8.5/7.5K, 60% flow, iter. 50
worst residual
−ln(likelihood)
sum of squared residuals

SSR vs. WR (top panels) and SSR vs. LL (bottom panels) biplots for 50 mapping iterations under high- and low-resolution settings: 15/8.5/2.5 ka (**A-E**) and 10/8.5/7.5 ka (**F-J**), respectively. Ancestry proportions of 95% (**A-E**) or 60% (**F-J**) G in H were simulated. The "information-poor" graphs (95,000 sites) and 171 alternative 2-way admixed models for the target group H were tested, and only top-ranking models are shown in each plot. Each ancestry source pair is shown as a dot and colored as follows: red, models where both sources are incorrect; yellow, models where only one source is correct; green, models where both sources are correct. We did not pay attention to admixture proportions inferred by *qpGraph*, only to source combinations. The settings, i.e. combinations of admixture and divergence dates in the clade of interest in thousands of years ago, are indicated above each plot. The horizontal red lines in the SSR vs. LL plots show *the lowest LL + 3 ln-units* threshold value considered as a statistical significance cut-off for ranking models having the same number of parameters (see Flegontov *et al*. (*11*) for further details). The horizontal red lines in the SSR vs. WR plots show *the lowest WR + 0.1 SE* threshold value.

**Fig. S6.**

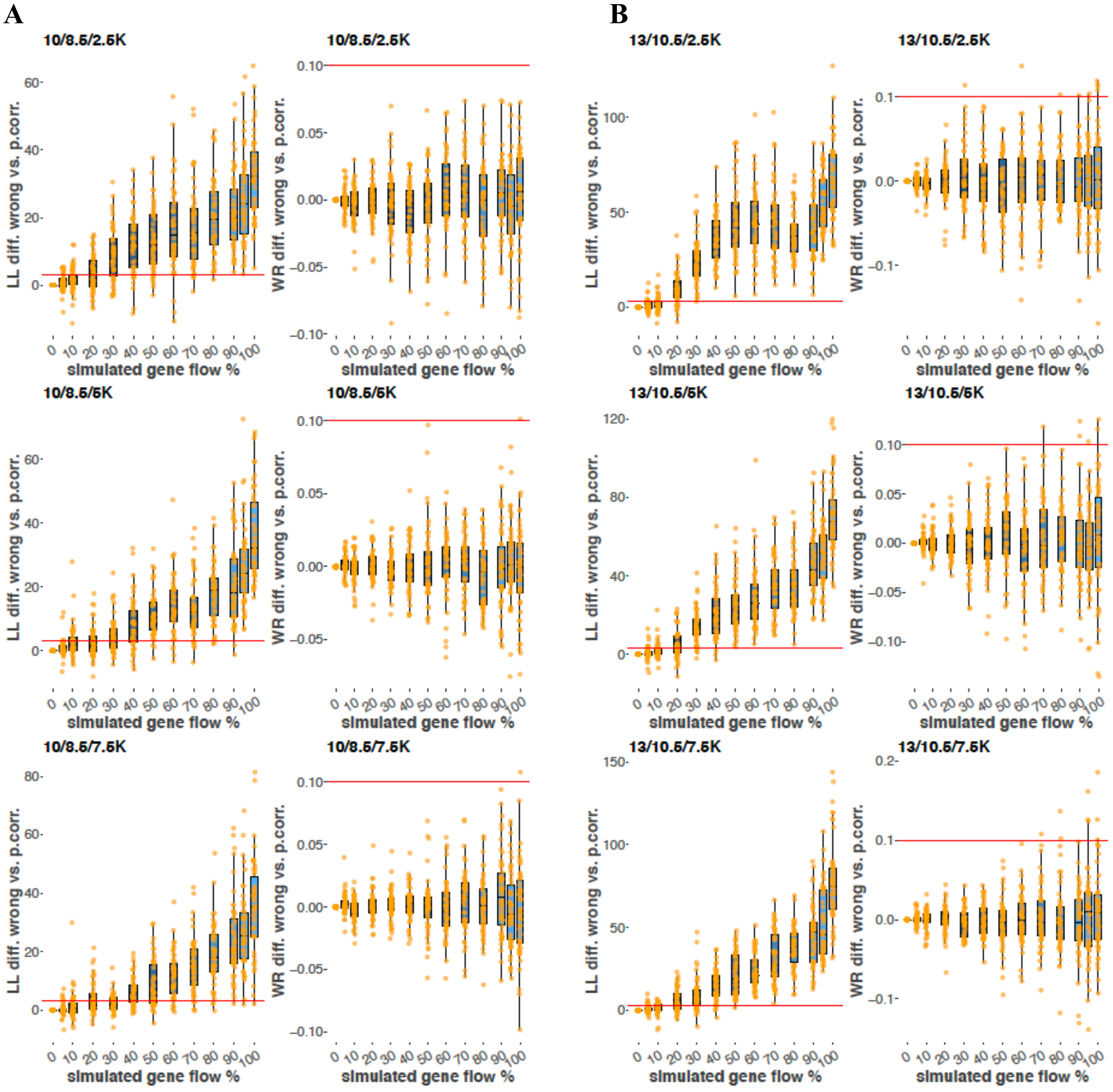
A
B
10/8.5/2.5K
13/10.5/2.5K
10/8.5/5K
13/10.5/5K
10/8.5/7.5K
13/10.5/7.5K
LL diff. wrong vs. p.corr.
WR diff. wrong vs. p.corr.
simulated gene flow %

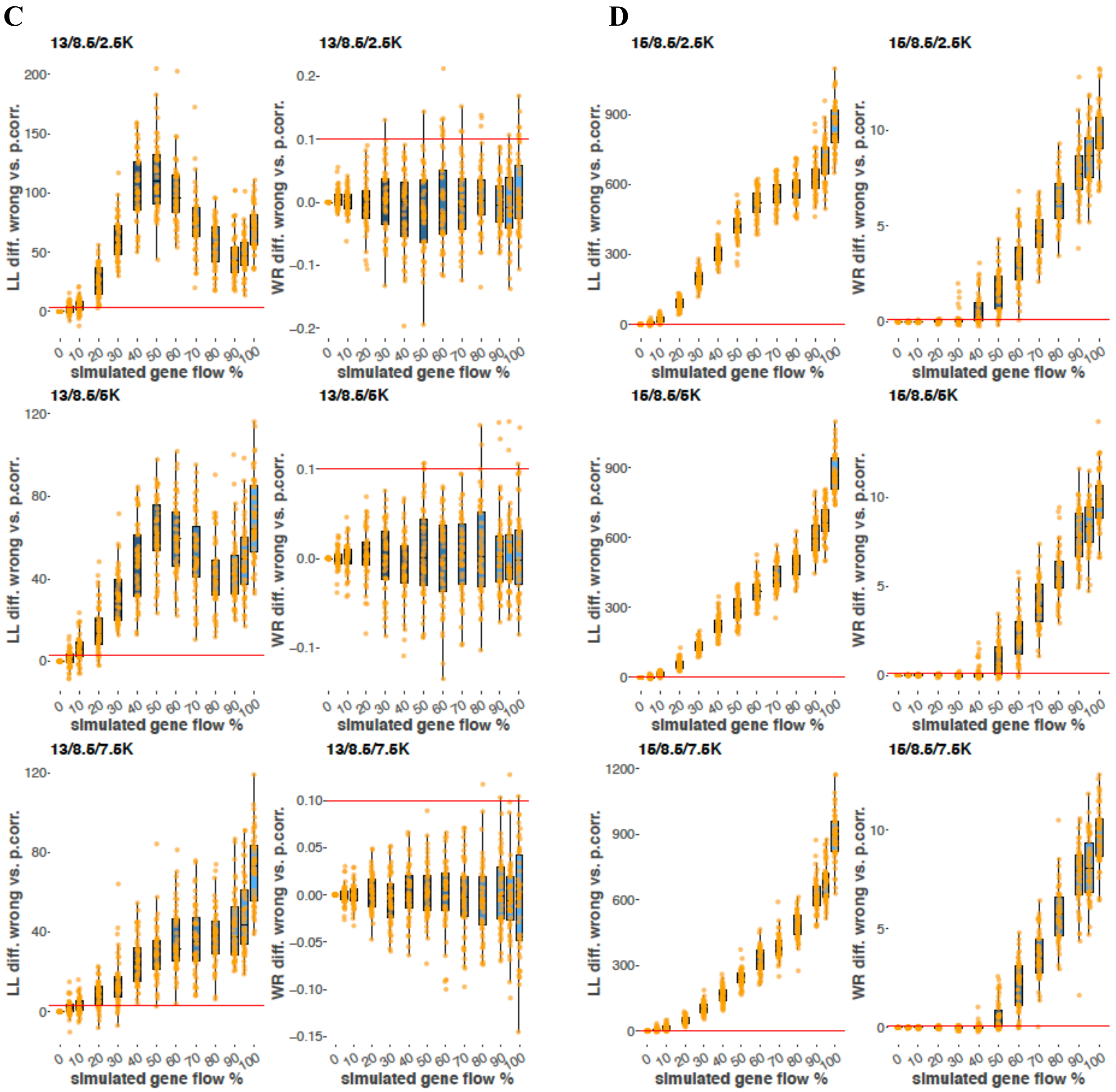


Boxplots illustrating differences between "partially correct 2" (G→H flow) and "incorrect 2" graphs (H→G flow) according to the LL (left-hand plots) and WR metrics (right-hand plots). The plots are based on "information-poor" graphs, 95,000 sites. Simulated gene flow proportions are shown on the x-axes, and each orange dot represents a simulation iteration. The settings, i.e. combinations of divergence and admixture dates in the clade of interest in thousands of years ago, are indicated above each plot: (**A**) 10/8.5 ka, (**B**) 13/10.5 ka, (**C**) 13/8.5 ka, (**D**) 15/8.5 ka. The horizontal red lines in the LL plots show *the lowest LL + 3 ln-units* threshold value considered as a statistical significance cut-off for ranking models having the same number of parameters (see Flegontov *et al.* (*11*) for further details). The horizontal red lines in the WR plots show *the lowest WR + 0.1 SE* threshold value.

**Fig. S7.**

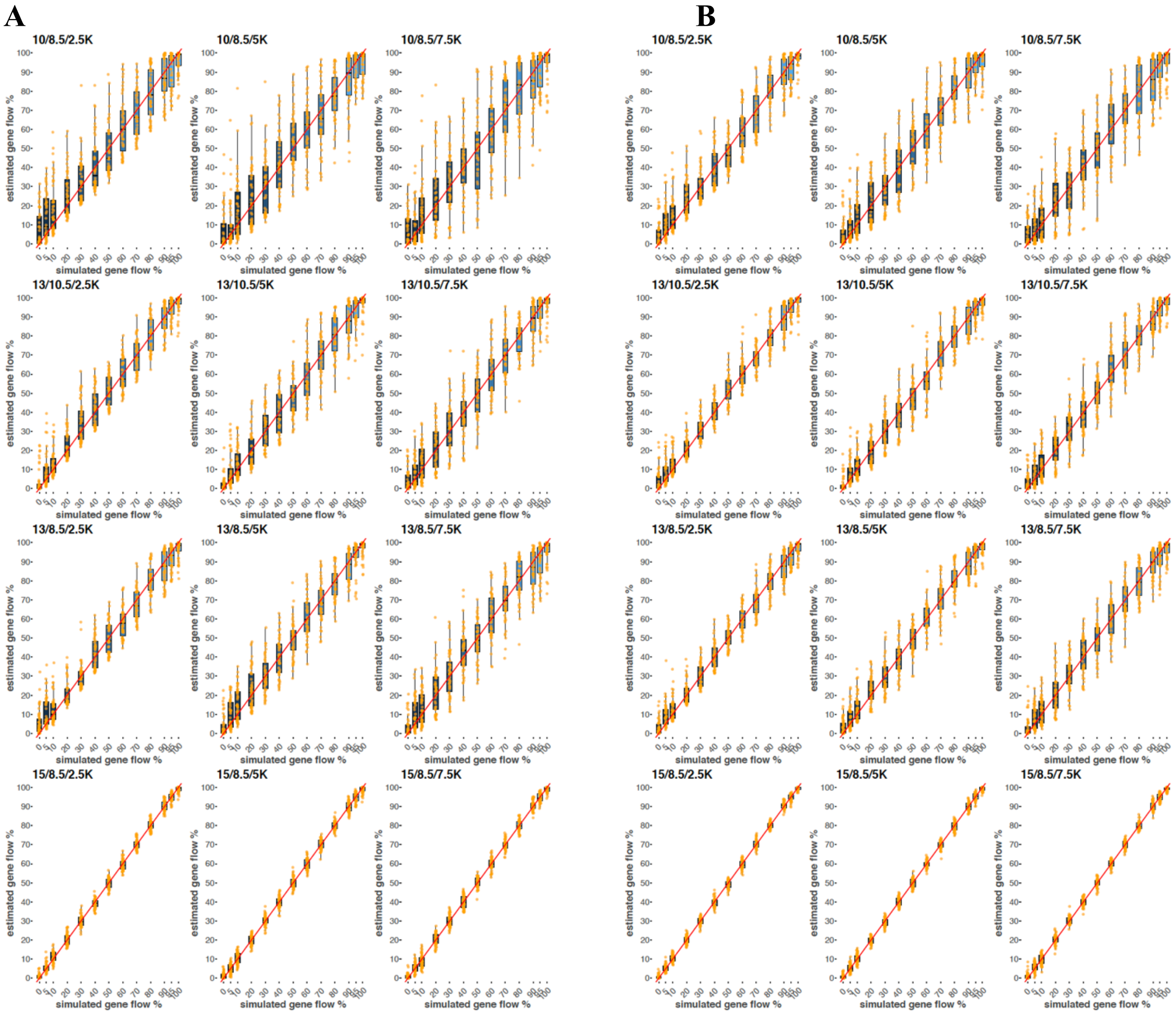


Boxplots illustrating estimated admixture proportions for "information-poor" graphs, 95,000 (**A**) or 400,000 sites (**B**). Only the correct H = I + G models were considered. Simulated gene flow proportions are shown on the x-axes, and each orange dot represents a simulation iteration. The settings, i.e. combinations of divergence and admixture dates in the clade of interest in thousands of years ago, are indicated above each plot. Diagonals are shown in red.

**Fig. S8.**

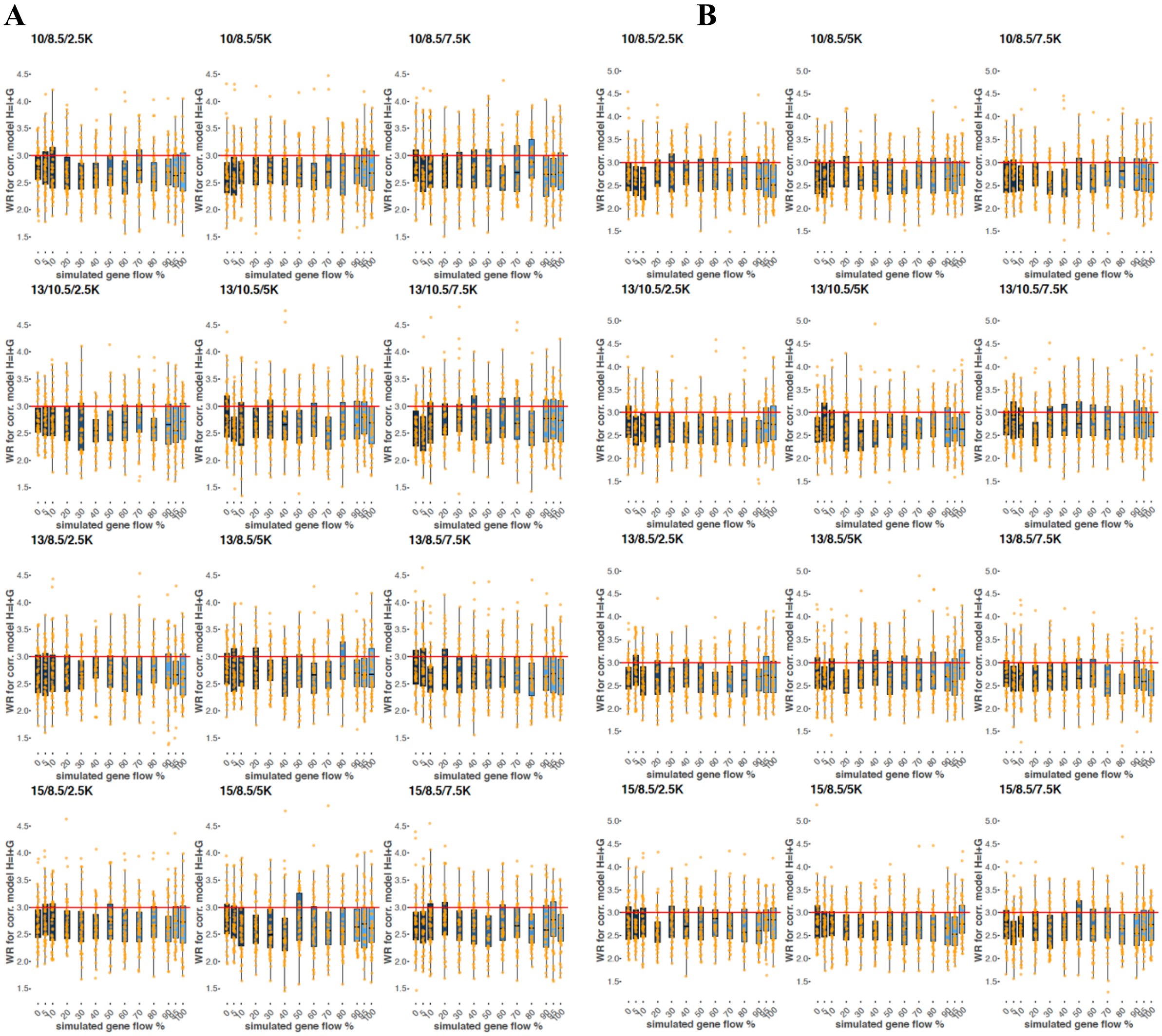


Boxplots illustrating the worst residuals for "information-poor" graphs, 95,000 (**A**) or 400,000 sites (**B**). Only the correct H = I + G models were considered (or H = I/H = G models for 0% and 100% G admixture proportions, respectively). Simulated admixture proportions are shown on the x-axes, and each orange dot represents a simulation iteration. The settings, i.e. combinations of divergence and admixture dates in the clade of interest in thousands of years ago, are indicated above each plot.

**Fig. S9.**

**A**

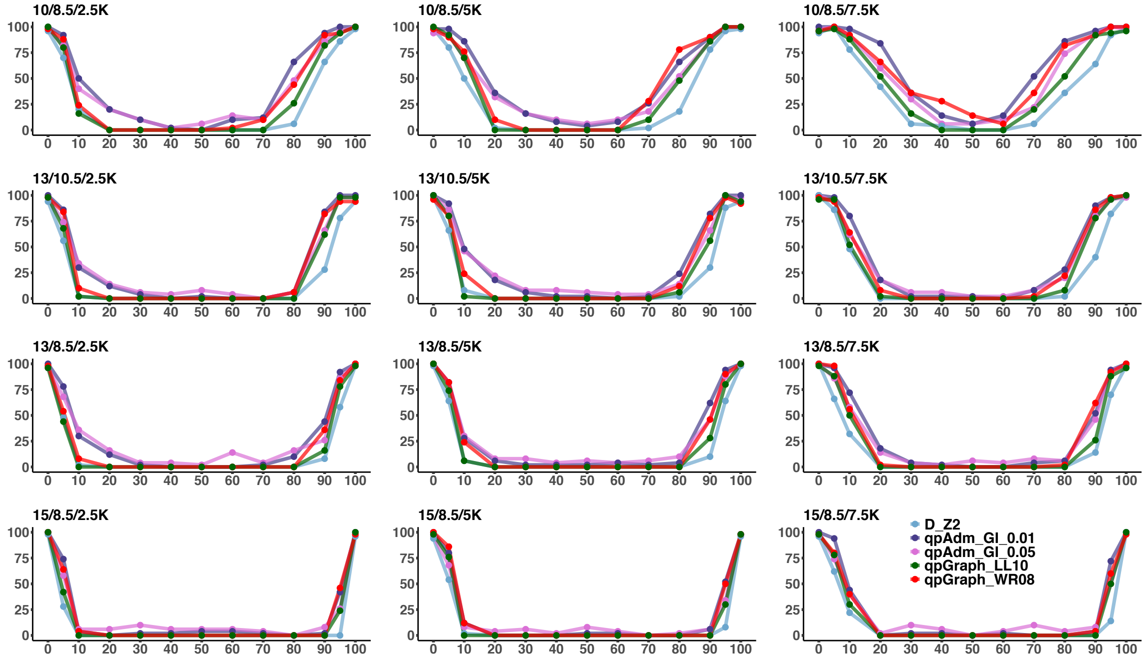


**B**

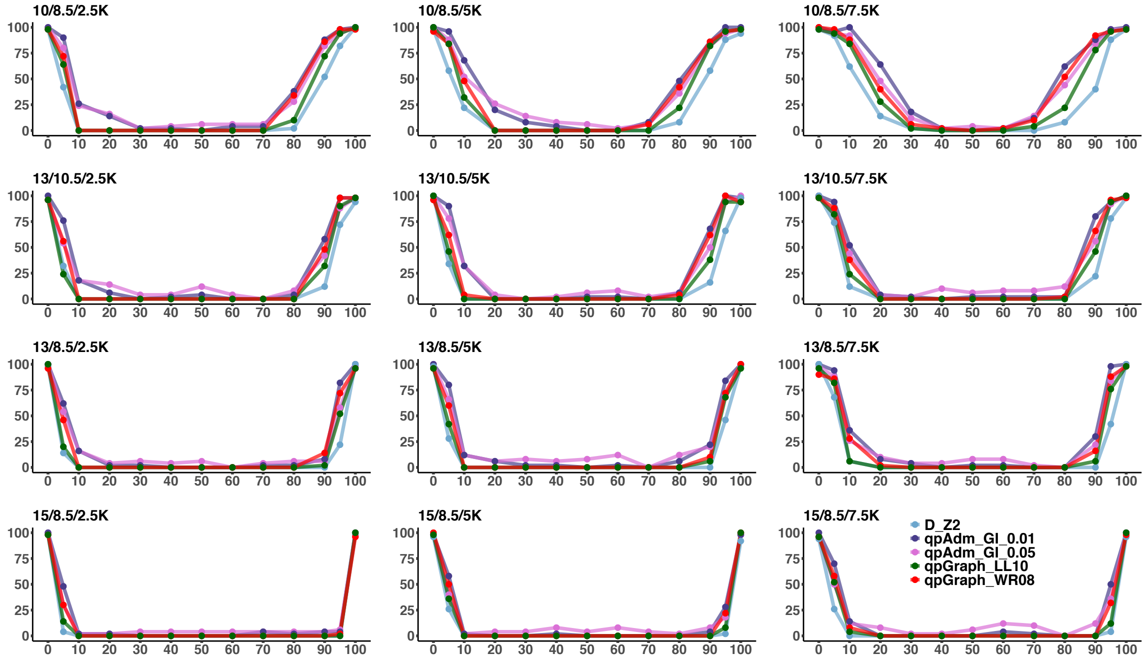


The fraction of iterations ("information-poor" demographic history) that gave a negative result according to *D*-statistics with a Z-score threshold of 2 (blue), *qpAdm*/*qpWave* with G or I in the outgroup set and a *p*-value threshold of 0.01 (dark violet) or 0.05 (orchid), and according to the *qpGraph* LL10 (dark green) and *qpGraph* WR0.8 classifiers (red). Sets of 95,000 (**A**) or 400,000 sites (**B**) were used. The settings, i.e. combinations of divergence and admixture dates in the clade of interest in thousands of years ago, are indicated above each plot.

Fig. S10.

A

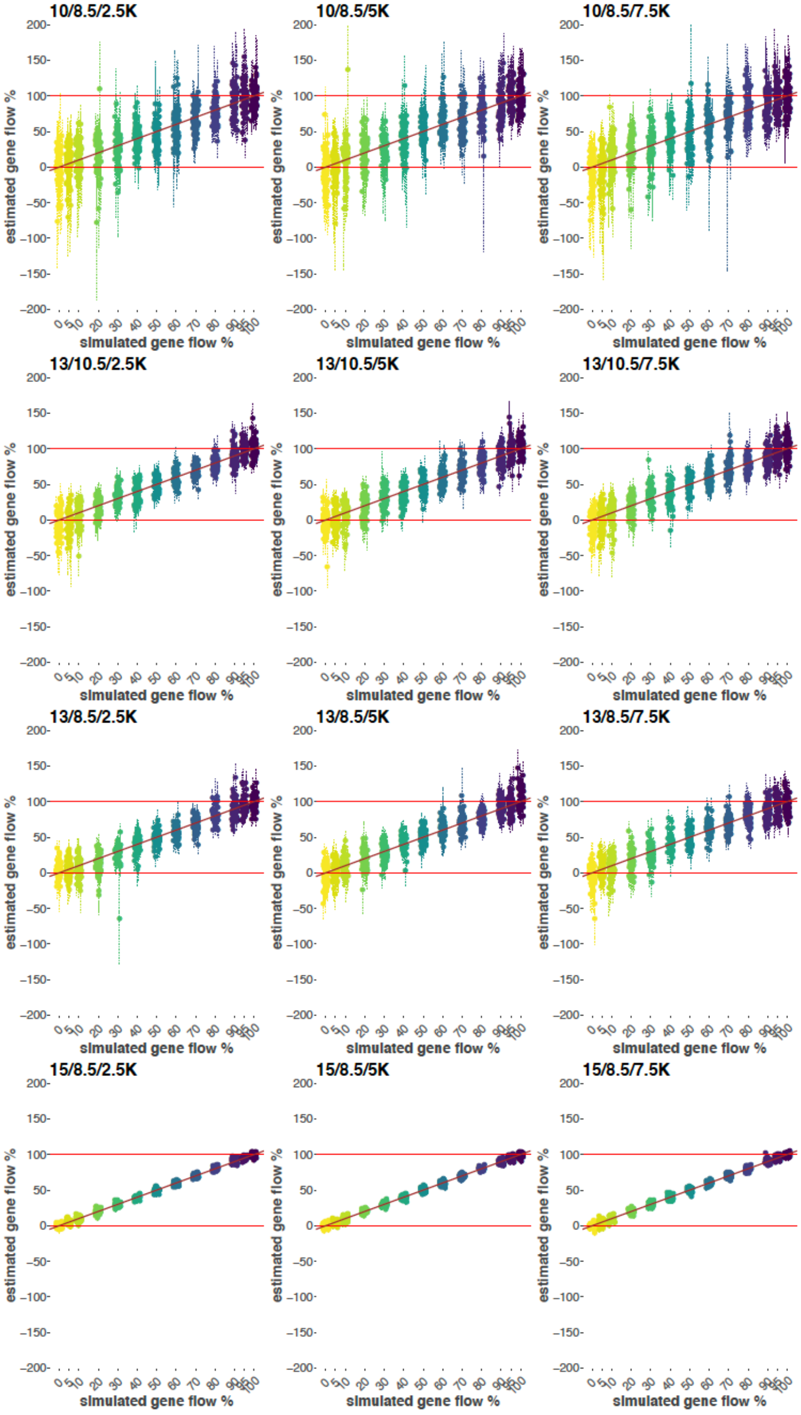
10/8.5/2.5K
10/8.5/5K
10/8.5/7.5K
13/10.5/2.5K
13/10.5/5K
13/10.5/7.5K
13/8.5/2.5K
13/8.5/5K
13/8.5/7.5K
15/8.5/2.5K
15/8.5/5K
15/8.5/7.5K
estimated gene flow %
simulated gene flow %

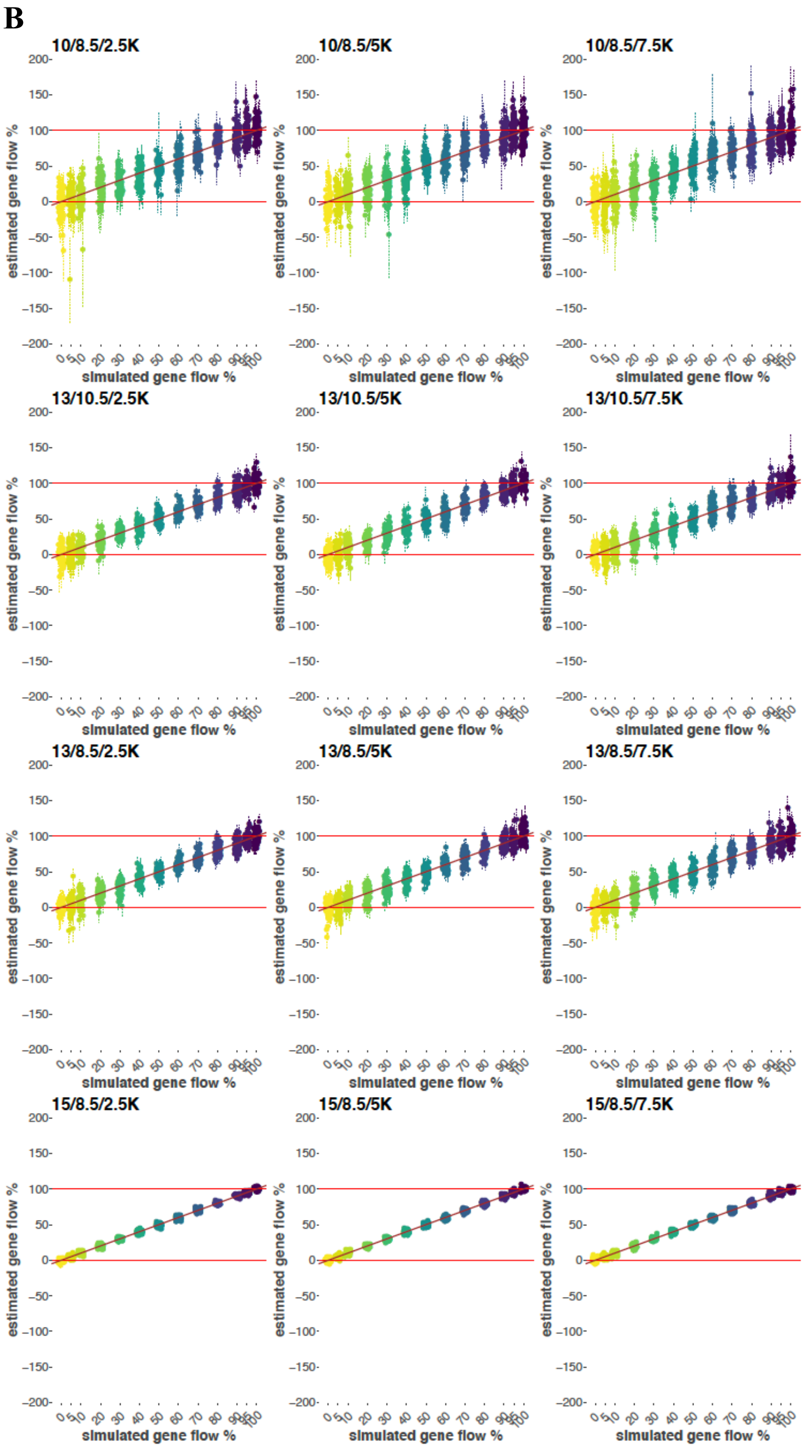


Dot plots illustrating admixture proportions estimated using *qpAdm* ("information-poor" demographic history) on sets of 95,000 (**A**) or 400,000 sites (**B**). Simulated G gene flow proportions are shown on the x-axes and coded by the color gradient. Estimated G ancestry proportions in H are shown on the y-axes, each dot represents a simulation iteration, and single SE intervals are shown as thin dotted lines. The settings, i.e. combinations of divergence and admixture dates in the clade of interest in thousands of years ago, are indicated above each plot. Diagonals are shown in brown.

**Fig. S11.**

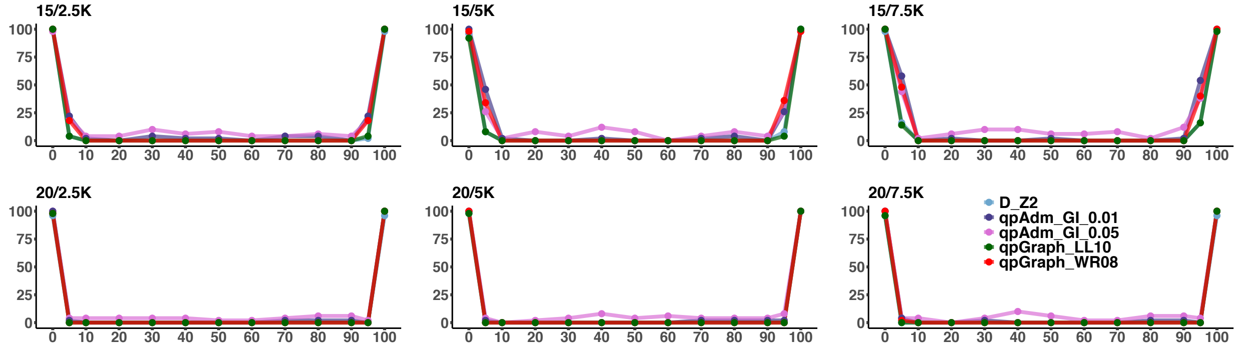


The fraction of iterations ("information-rich" demographic history, 95,000 sites) that gave a negative result according to *D*-statistics with a Z-score threshold of 2 (blue), *qpAdm*/*qpWave* with G or I in the outgroup set and a *p*-value threshold of 0.01 (dark violet) or 0.05 (orchid), and according to the *qpGraph* LL10 (dark green) and *qpGraph* WR0.8 classifiers (red). The settings, i.e. combinations of divergence and admixture dates in the clade of interest in thousands of years ago, are indicated above each plot.

**Fig. S12.**

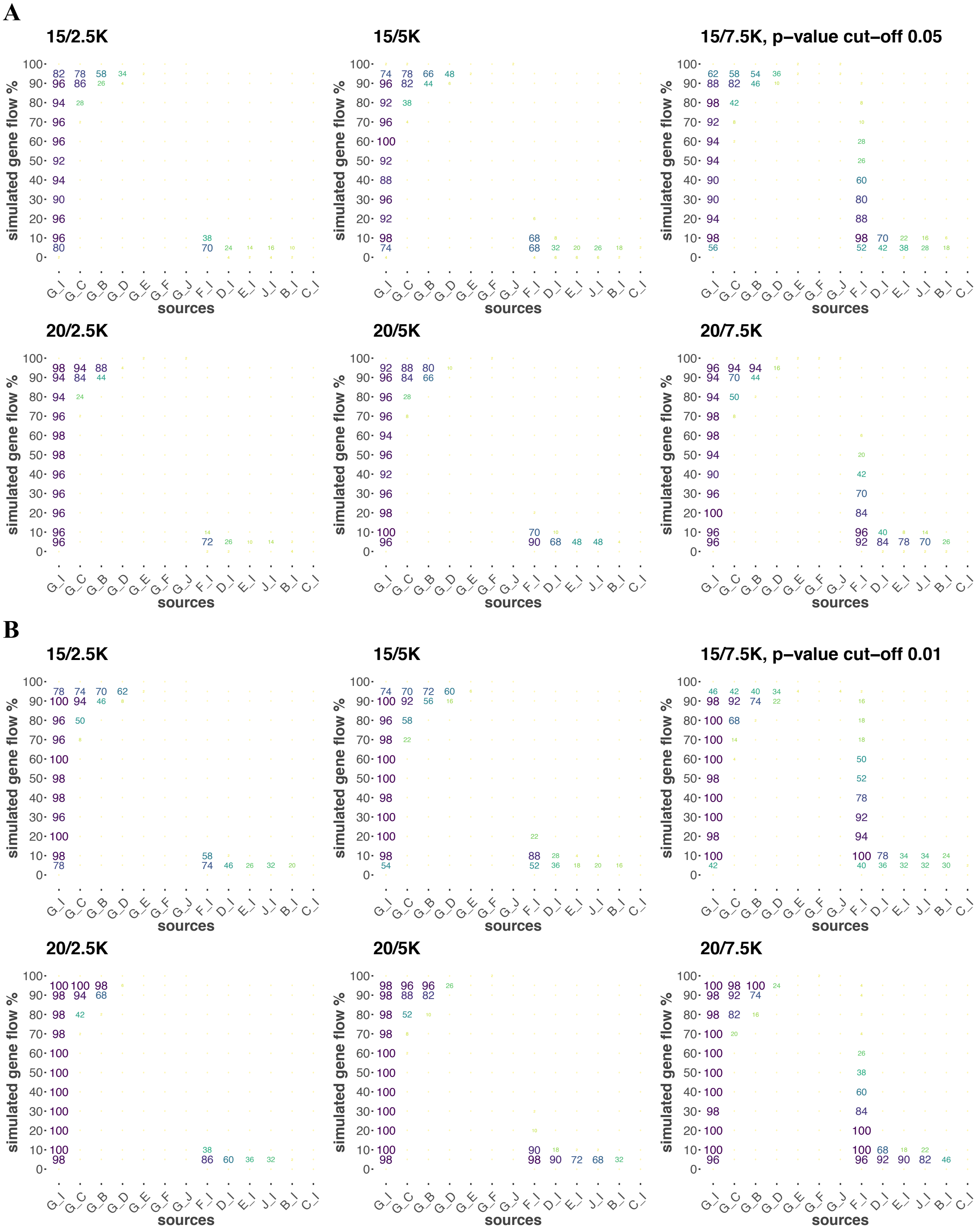
A
15/2.5K
15/5K
15/7.5K, p–value cut–off 0.05
20/2.5K
20/5K
20/7.5K
B
15/2.5K
15/5K
15/7.5K, p–value cut–off 0.01
20/2.5K
20/5K
20/7.5K
simulated gene flow %
sources

The percentage of iterations ("information-rich" demographic history, 95,000 sites) that gave a positive result according to *qpWave*/*qpAdm* classifiers with *p*-value thresholds of 0.05 (**A**) or 0.01 (**B**). Percentages are coded by color and size of the digits. Alternative source pairs are labelled on the x-axis and shown in columns. Results for the correct source pair G+I are shown in the leftmost column. Simulated G admixture proportions are shown on the y-axis. The settings, i.e. combinations of divergence and admixture dates in the clade of interest in thousands of years ago, are indicated above each plot.

**Fig. S13.**

**A**

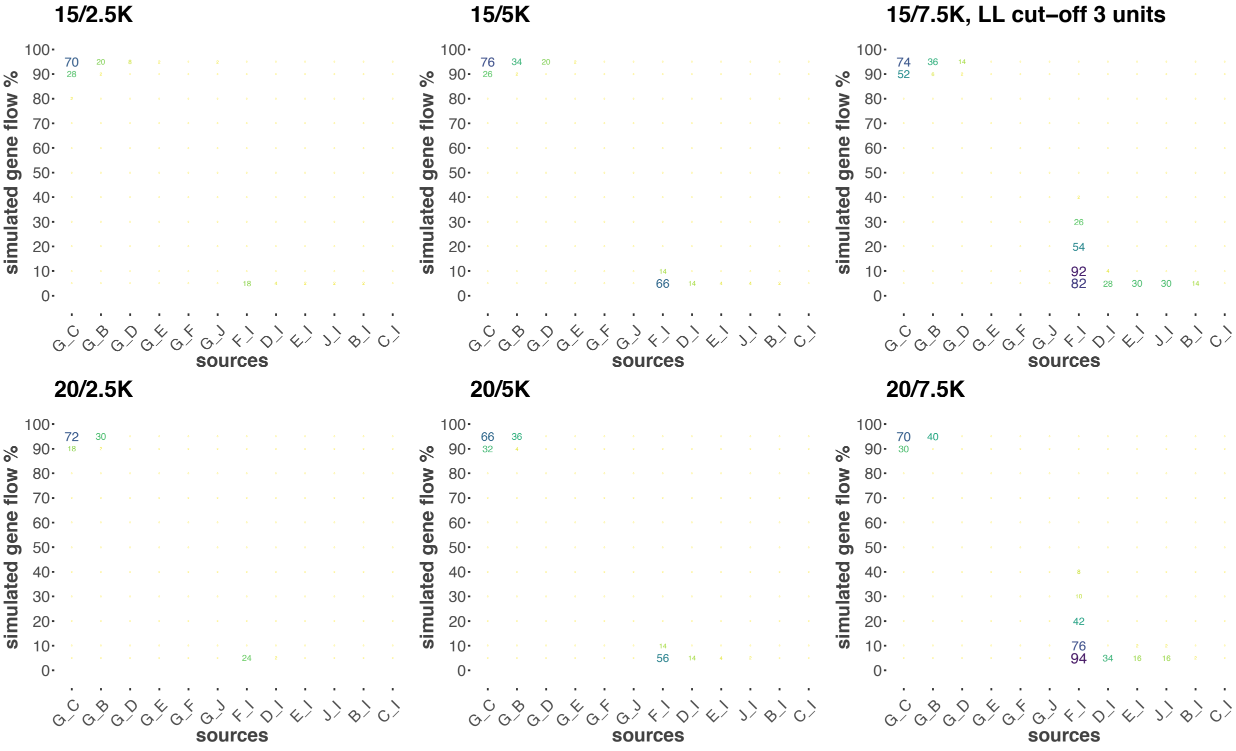
15/2.5K
15/5K
15/7.5K, LL cut–off 3 units
20/2.5K
20/5K
20/7.5K
simulated gene flow %
sources
G_C G_B G_D G_E G_F G_J F_I D_I E_I J_I B_I C_I

**B**

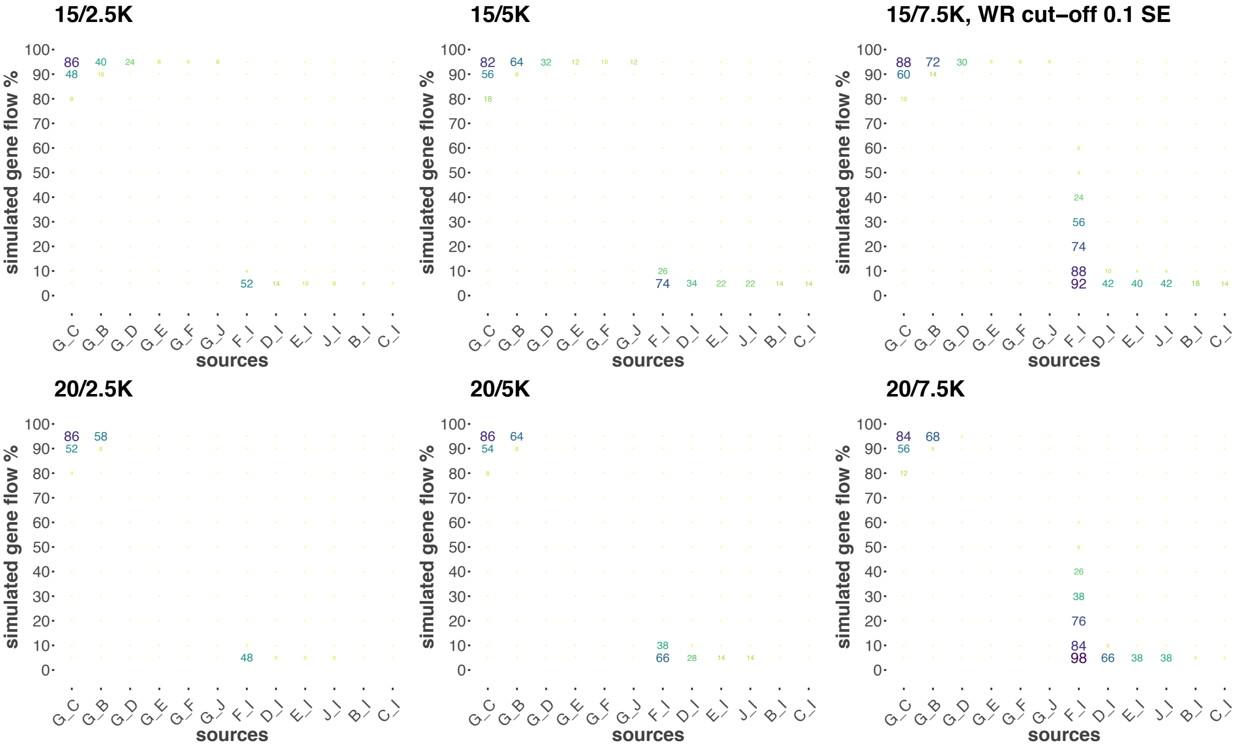
15/2.5K
15/5K
15/7.5K, WR cut–off 0.1 SE
20/2.5K
20/5K
20/7.5K
simulated gene flow %
sources
G_C G_B G_D G_E G_F G_J F_I D_I E_I J_I B_I C_I

The percentage of iterations ("information-rich" demographic history, 95,000 sites) where partially correct source pairs were indistinguishable from the correct source pair G+I according to a LL threshold of 3 units (**A**) or a WR threshold of 0.1 (**B**). Percentages are coded by color and size of the digits. Alternative source pairs are labelled on the x-axis and shown in columns. Simulated G admixture proportions are shown on the y-axis. The settings, i.e. combinations of divergence and admixture dates in the clade of interest in thousands of years ago, are indicated above each plot.

**Fig. S14.**

**A**

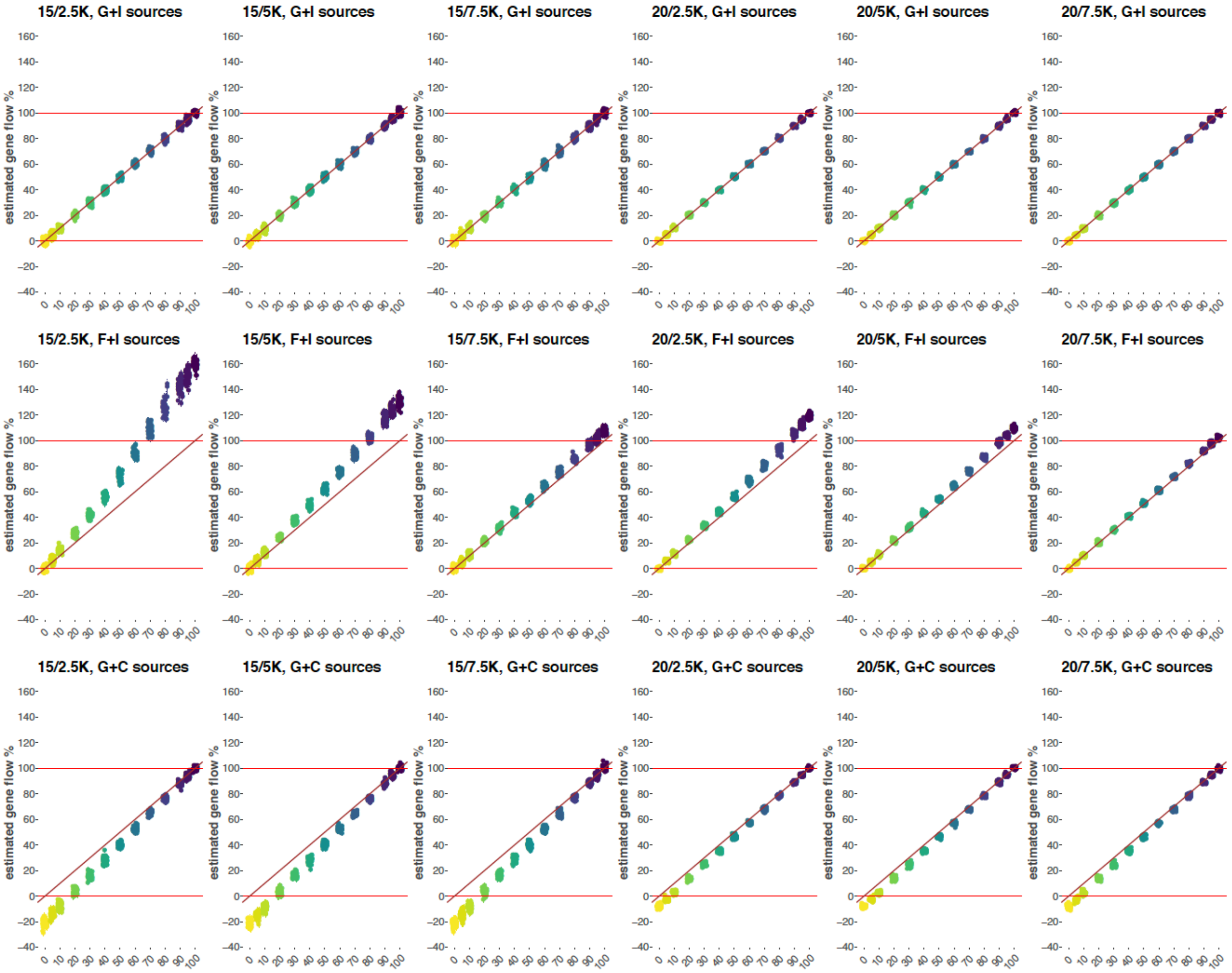
15/2.5K, G+I sources
15/5K, G+I sources
15/7.5K, G+I sources
20/2.5K, G+I sources
20/5K, G+I sources
20/7.5K, G+I sources
15/2.5K, F+I sources
15/5K, F+I sources
15/7.5K, F+I sources
20/2.5K, F+I sources
20/5K, F+I sources
20/7.5K, F+I sources
15/2.5K, G+C sources
15/5K, G+C sources
15/7.5K, G+C sources
20/2.5K, G+C sources
20/5K, G+C sources
20/7.5K, G+C sources
estimated gene flow %

**B**

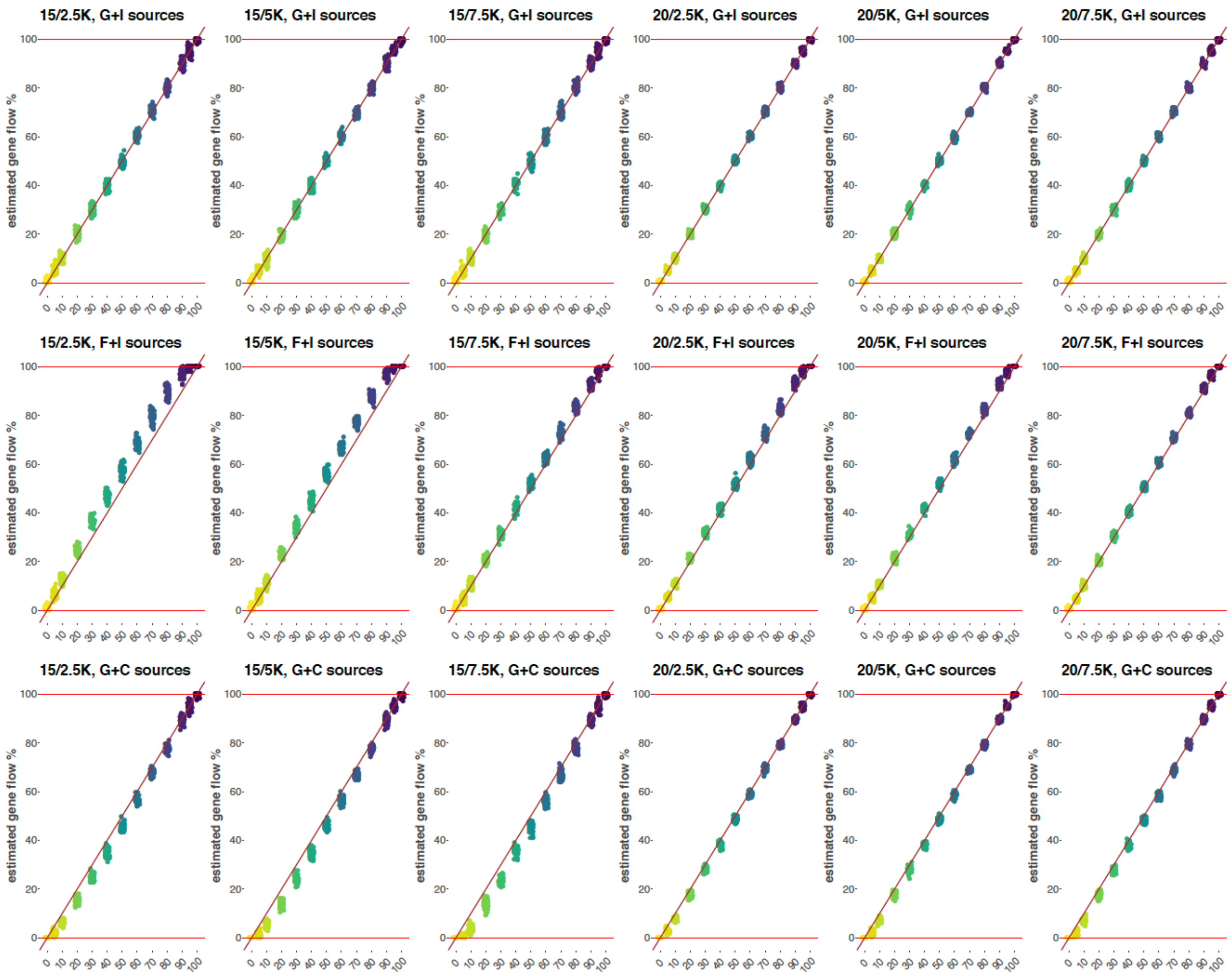


Dot plots illustrating admixture proportions estimated for the "information-rich" demographic history (95,000 sites) using *qpAdm* (**A**) or *qpGraph* (**B**). Result for the correct source pair G+I and for two partially correct source pairs F+I and G+C are shown. Simulated G gene flow proportions are shown on the x-axes and coded by the color gradient. Estimated G ancestry proportions in H are shown on the y-axes, each dot represents a simulation iteration, and single SE intervals are shown as thin dotted lines (**A**). The settings, i.e. combinations of divergence and admixture dates in the clade of interest in thousands of years ago, are indicated above each plot. Diagonals are shown in brown.

**Fig. S15.**

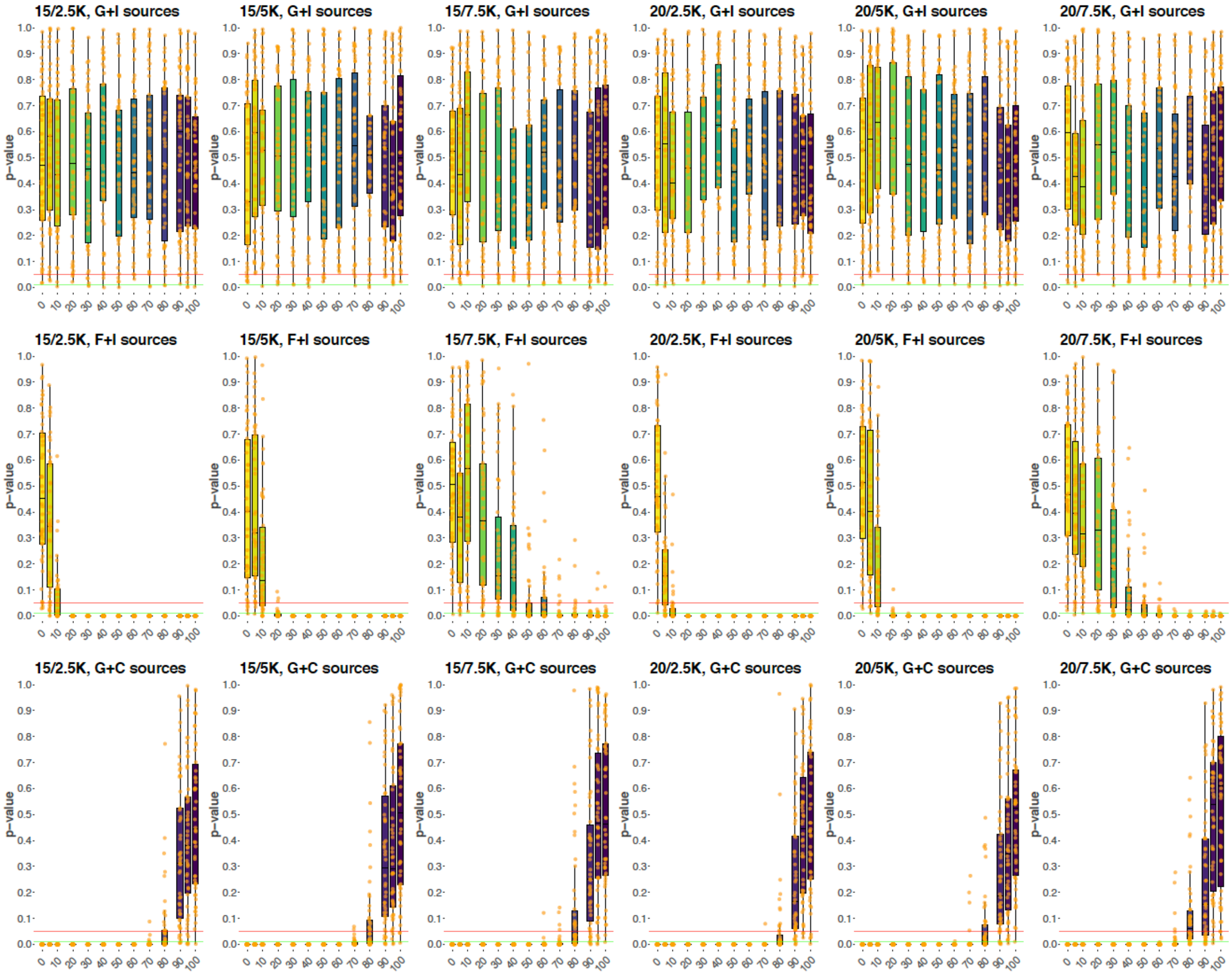


Boxplots illustrating the distribution of *p*-values for the H = G + I, H = F + I, and H = G + C *qpAdm* models ("information-rich" demographic history, 95,000 sites). Simulated G gene flow proportions are shown on the x-axes and coded by the color gradient, and *p*-values are shown on the y-axes. Each dot represents a simulation iteration. The settings, i.e. combinations of divergence and admixture dates in the clade of interest in thousands of years ago, are indicated above each plot. The horizontal red and green lines correspond to the 0.05 and 0.01 *p*-value cut-offs, respectively.

**Fig. S16.**

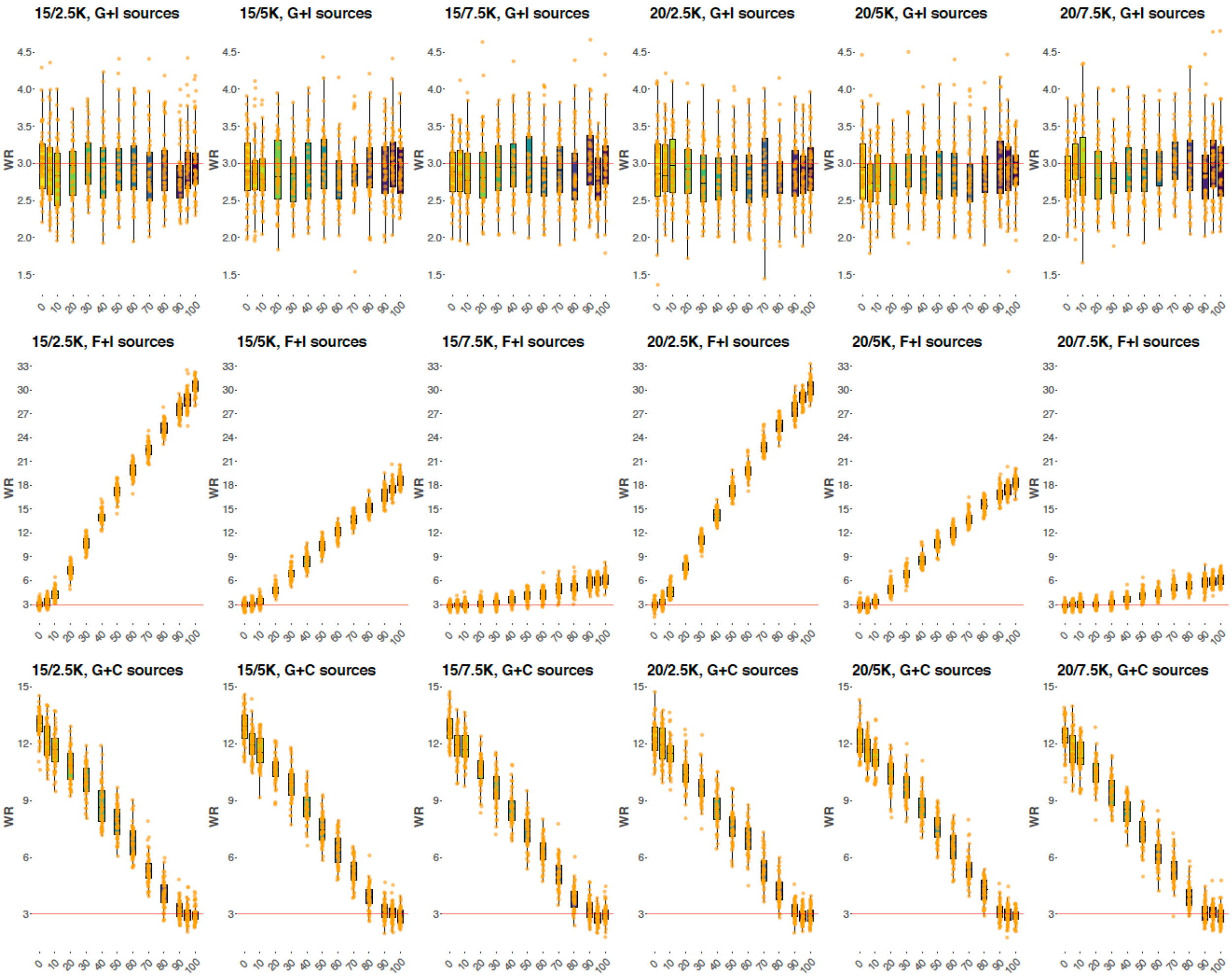


Boxplots illustrating the worst admixture graph residuals ("information rich" graphs, 95,000 sites) for the correct H = G + I models (or H = I/H = G models for 0% and 100% G admixture proportions, respectively) or for partially correct models H = F + I and H = C + G. Simulated admixture proportions are shown on the x-axes, and each orange dot represents a simulation iteration. The settings, i.e. combinations of divergence and admixture dates in the clade of interest in thousands of years ago, are indicated above each plot.

**Fig. S17.**

**A**

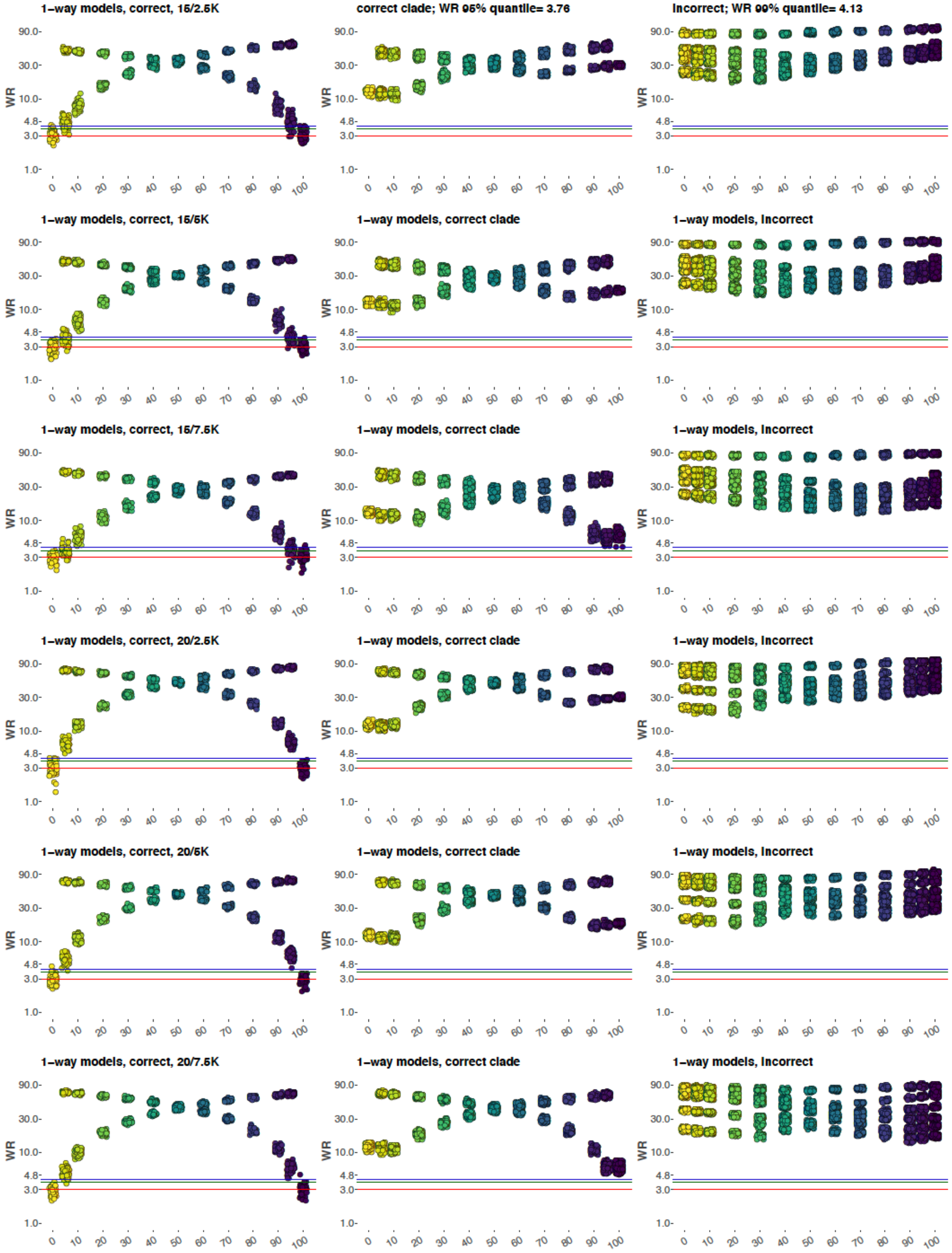

1–way models, correct, 15/2.5K
correct clade; WR 95% quantile= 3.76
incorrect; WR 99% quantile= 4.13
1–way models, correct, 15/5K
1–way models, correct clade
1–way models, incorrect
1–way models, correct, 15/7.5K
1–way models, correct clade
1–way models, incorrect
1–way models, correct, 20/2.5K
1–way models, correct clade
1–way models, incorrect
1–way models, correct, 20/5K
1–way models, correct clade
1–way models, incorrect
1–way models, correct, 20/7.5K
1–way models, correct clade
1–way models, incorrect
WR
90.0
30.0
10.0
4.8
3.0
1.0
0 10 20 30 40 50 60 70 80 90 100

**B**

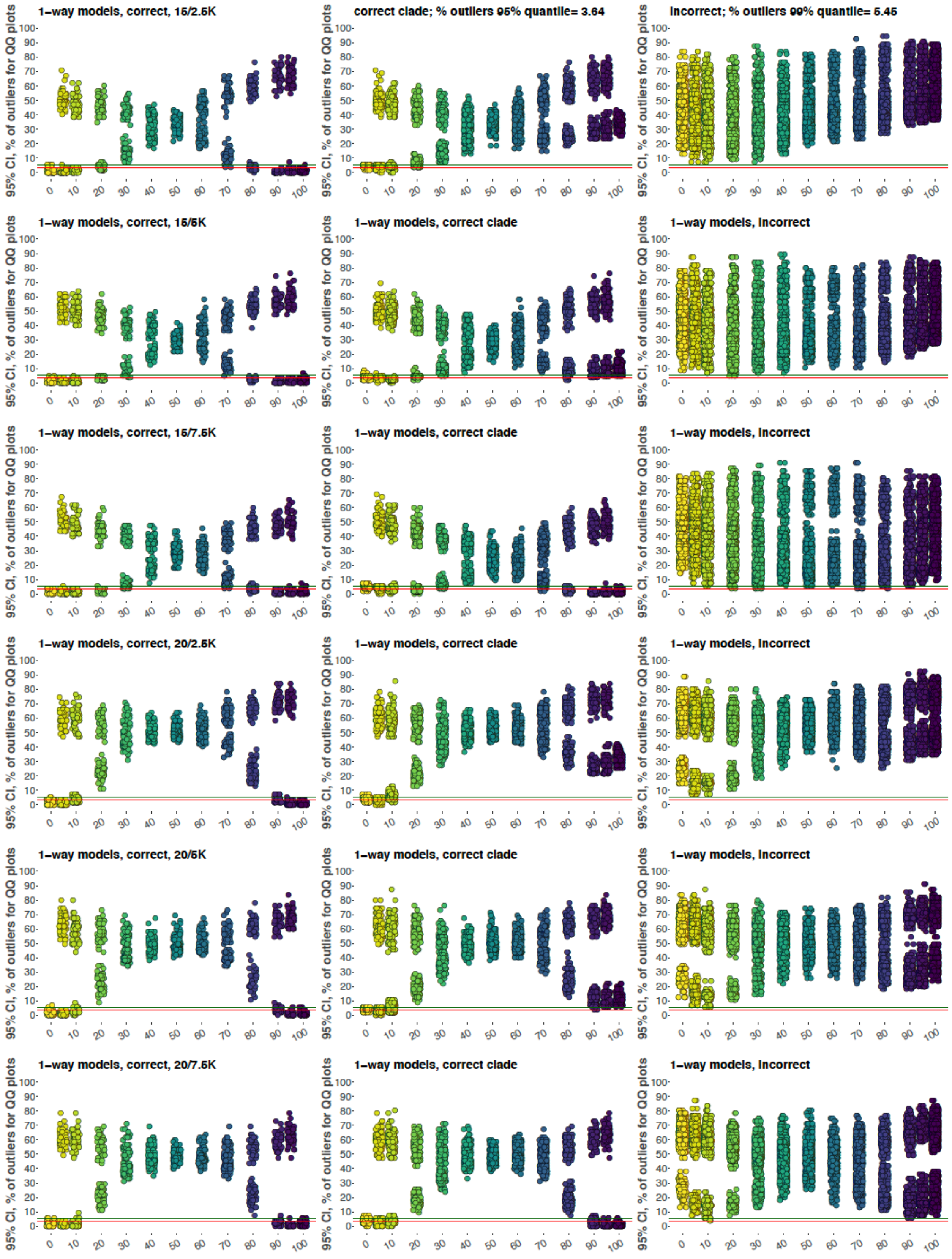
1–way models, correct, 15/2.5K
correct clade; % outliers 95% quantile= 3.64
incorrect; % outliers 99% quantile= 5.45
1–way models, correct, 15/5K
1–way models, correct clade
1–way models, incorrect
1–way models, correct, 15/7.5K
1–way models, correct clade
1–way models, incorrect
1–way models, correct, 20/2.5K
1–way models, correct clade
1–way models, incorrect
1–way models, correct, 20/5K
1–way models, correct clade
1–way models, incorrect
1–way models, correct, 20/7.5K
1–way models, correct clade
1–way models, incorrect
95% CI, % of outliers for QQ plots
0 10 20 30 40 50 60 70 80 90 100
0 10 20 30 40 50 60 70 80 90 100

**C**

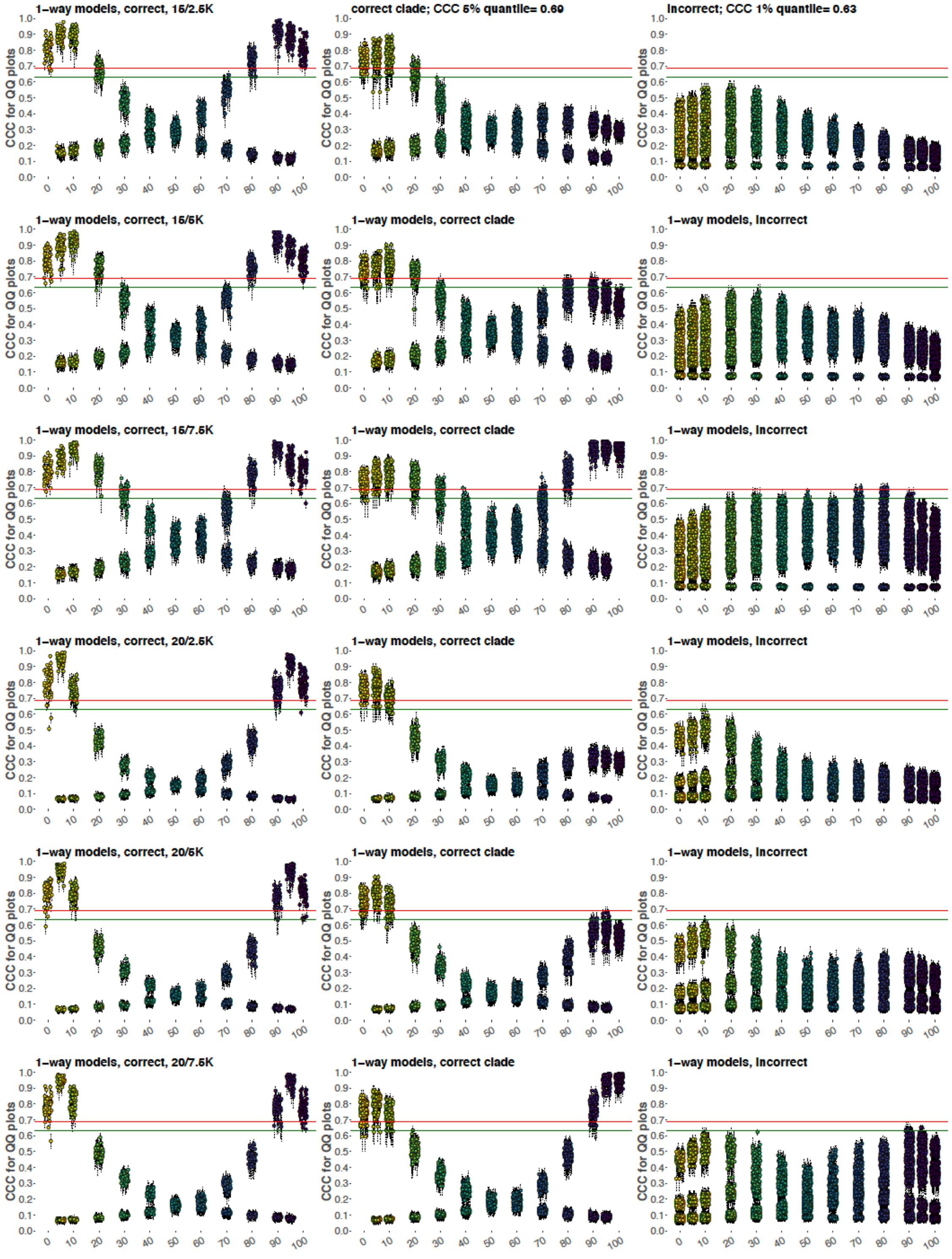
1–way models, correct, 15/2.5K
correct clade; CCC 5% quantile= 0.69
Incorrect; CCC 1% quantile= 0.63
1–way models, correct, 15/5K
1–way models, correct clade
1–way models, Incorrect
1–way models, correct, 15/7.5K
1–way models, correct clade
1–way models, Incorrect
1–way models, correct, 20/2.5K
1–way models, correct clade
1–way models, Incorrect
1–way models, correct, 20/5K
1–way models, correct clade
1–way models, Incorrect
1–way models, correct, 20/7.5K
1–way models, correct clade
1–way models, Incorrect
CCC for QQ plots

Testing three alternative methods for assessing absolute model fit to the data on the information-rich graph, 95,000 sites and unadmixed models for group H. (**A**) Worst $f_4$-statistic residuals for correct sources, sources closely related to the correct ones ("correct clade"), and incorrect sources. Iterations are grouped by simulated G admixture proportions on the x-axis, and worst residual values are plotted on the y-axis in log10 scale. The traditional threshold of 3 SE is shown with red lines, and 95% and 99% quantiles calculated on the combined dataset without admixture (0% and 100% G proportion in H, all settings combined) are shown with green and blue lines, respectively. The 95% quantile = 3.76 SE, and the 99% quantile = 4.13 SE. (**B**) Percentage of $f_2$-statistic residuals that lie outside of the 95% confidence interval around the standard normal distribution. 95% and 99% quantiles calculated on the combined dataset without admixture are shown with red and green lines, respectively. The 95% quantile = 3.6%, and the 99% quantile = 5.5%. (**C**) Concordance correlation coefficients (and their 95% confidence intervals) for $f_2$-statistic residuals vs. the standard normal distribution. 5% and 1% quantiles calculated on the combined dataset without admixture are shown with red and green lines, respectively. The 5% quantile = 0.69, and the 1% quantile = 0.63.

**Fig. S18.**

**A**

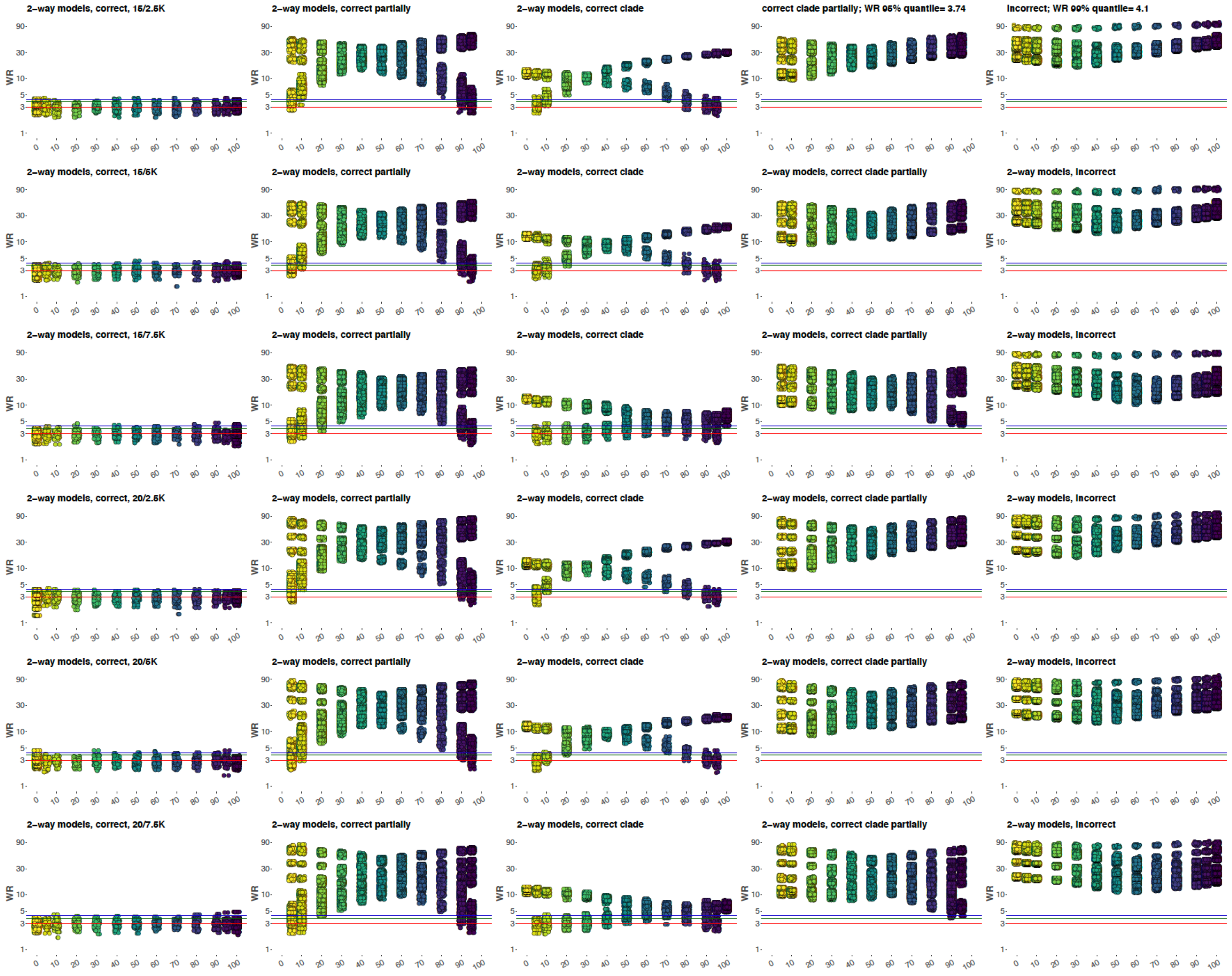
2–way models, correct, 15/2.5K
2–way models, correct partially
2–way models, correct clade
correct clade partially; WR 95% quantile= 3.74
incorrect; WR 99% quantile= 4.1
2–way models, correct, 15/5K
2–way models, correct clade partially
2–way models, incorrect
2–way models, correct, 15/7.5K
2–way models, correct, 20/2.5K
2–way models, correct, 20/5K
2–way models, correct, 20/7.5K
WR

**B**

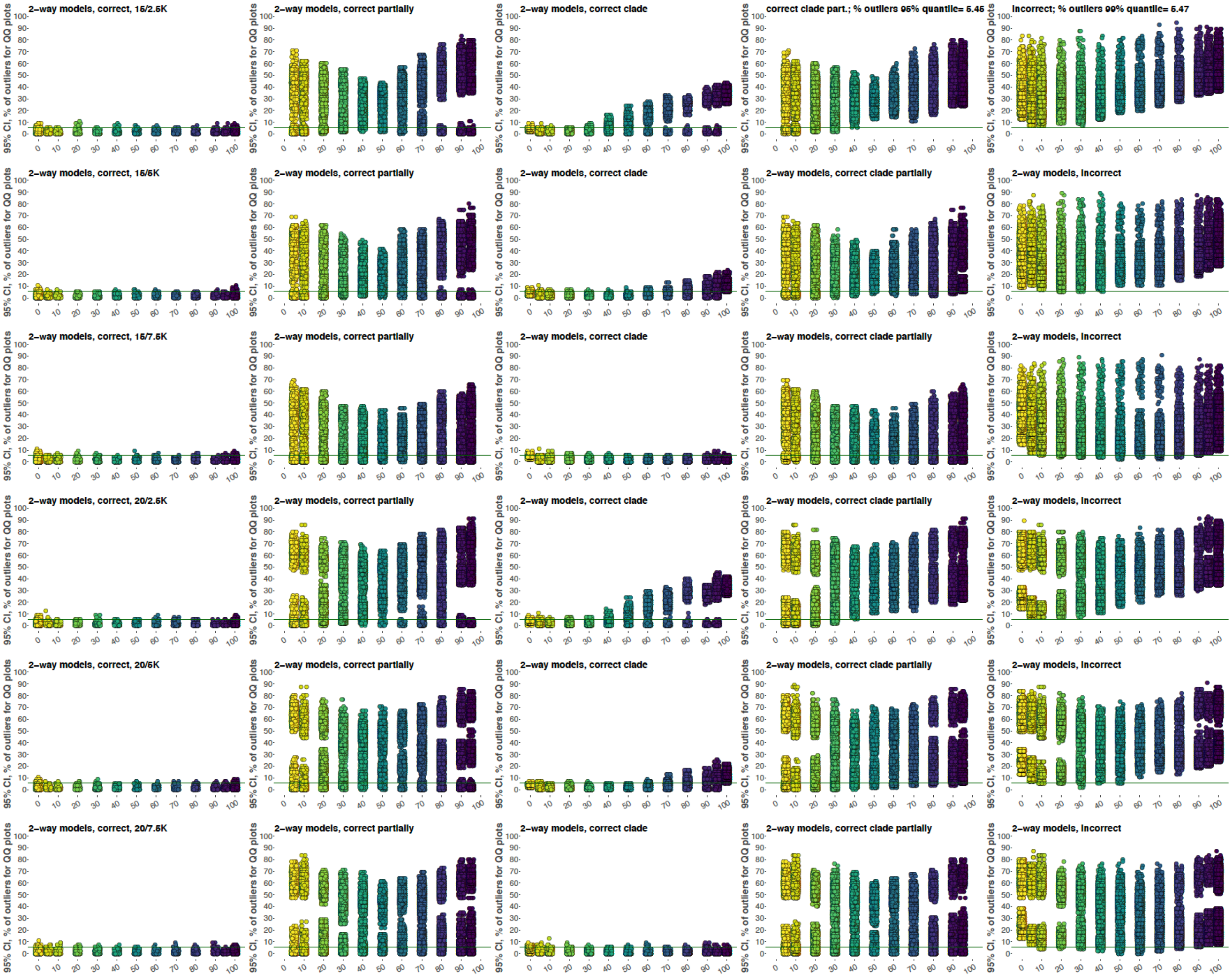
2–way models, correct, 15/2.5K
2–way models, correct partially
2–way models, correct clade
correct clade part.; % outliers 95% quantile= 5.45
incorrect; % outliers 99% quantile= 5.47
2–way models, correct, 15/5K
2–way models, correct partially
2–way models, correct clade
2–way models, correct clade partially
2–way models, incorrect
2–way models, correct, 15/7.5K
2–way models, correct partially
2–way models, correct clade
2–way models, correct clade partially
2–way models, incorrect
2–way models, correct, 20/2.5K
2–way models, correct partially
2–way models, correct clade
2–way models, correct clade partially
2–way models, incorrect
2–way models, correct, 20/5K
2–way models, correct partially
2–way models, correct clade
2–way models, correct clade partially
2–way models, incorrect
2–way models, correct, 20/7.5K
2–way models, correct partially
2–way models, correct clade
2–way models, correct clade partially
2–way models, incorrect
95% CI, % of outliers for QQ plots

C

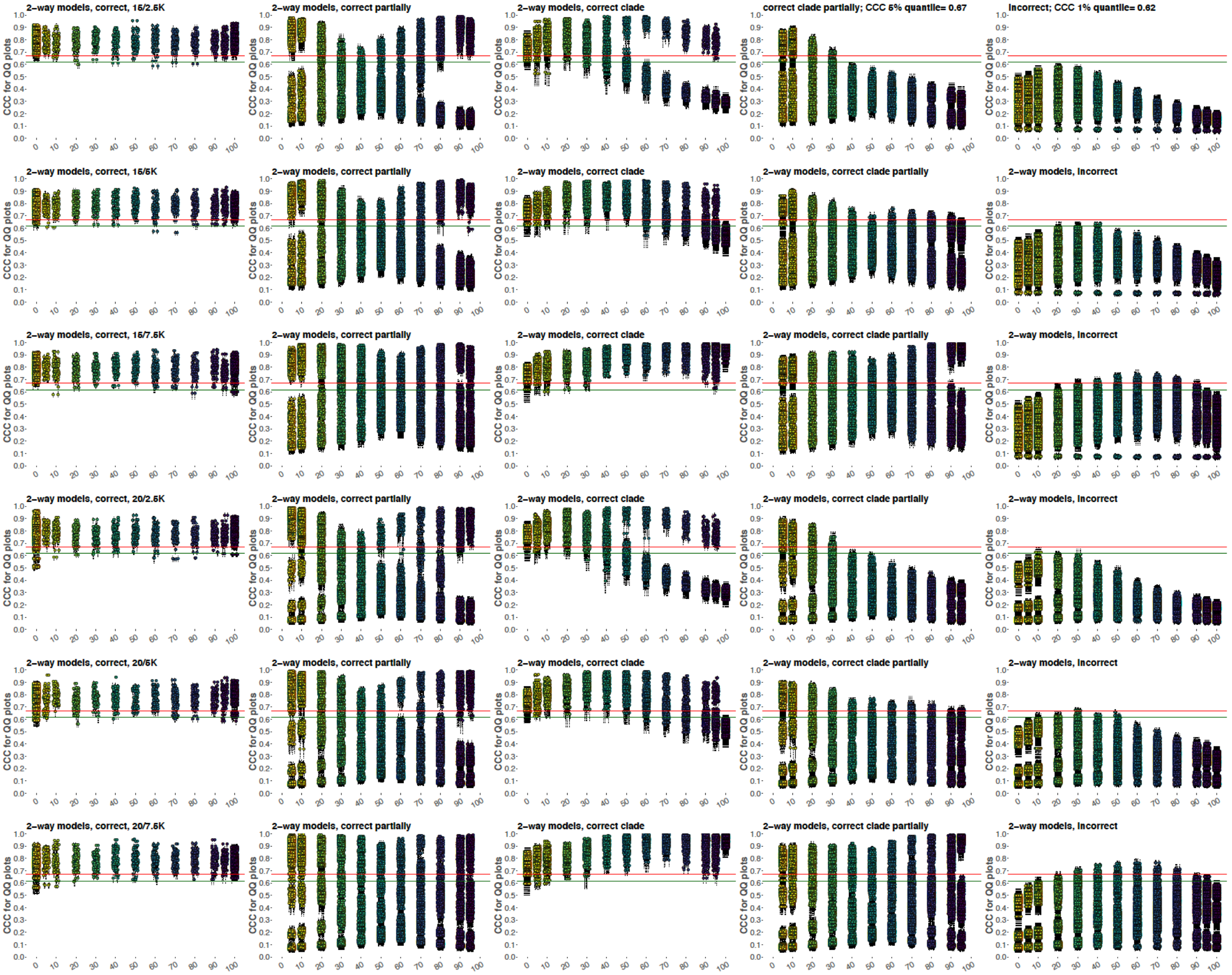

2–way models, correct, 15/2.5K
2–way models, correct partially
2–way models, correct clade
correct clade partially; CCC 5% quantile= 0.67
incorrect; CCC 1% quantile= 0.62
2–way models, correct, 15/5K
2–way models, correct partially
2–way models, correct clade
2–way models, correct clade partially
2–way models, incorrect
2–way models, correct, 15/7.5K
2–way models, correct partially
2–way models, correct clade
2–way models, correct clade partially
2–way models, incorrect
2–way models, correct, 20/2.5K
2–way models, correct partially
2–way models, correct clade
2–way models, correct clade partially
2–way models, incorrect
2–way models, correct, 20/5K
2–way models, correct partially
2–way models, correct clade
2–way models, correct clade partially
2–way models, incorrect
2–way models, correct, 20/7.5K
2–way models, correct partially
2–way models, correct clade
2–way models, correct clade partially
2–way models, incorrect
CCC for QQ plots

Testing three alternative methods for assessing absolute model fit to the data on the information-rich graph, 95,000 sites and 2-way admixed models for group H. (**A**) Worst $f_4$-statistic residuals for correct source pairs, pairs with only one correct source ("correct partially"), pairs with one ("correct clade partially") or both sources ("correct clade") closely related to the correct ones, and incorrect pairs. Iterations are grouped by simulated G admixture proportions on the x-axis, and worst residual values are plotted on the y-axis in log10 scale. The traditional threshold of 3 SE is shown with red lines, and 95% and 99% quantiles calculated on the combined dataset with simulated admixture (5% to 95% G proportion in H, all settings combined) are shown with green and blue lines, respectively. The 95% quantile = 3.74 SE, and the 99% quantile = 4.1 SE. (**B**) Percentage of $f_2$-statistic residuals that lie outside of the 95% confidence interval around the standard normal distribution. 95% and 99% quantiles calculated on the combined dataset with simulated admixture are shown with red and green lines, respectively. The 95% quantile = 5.45%, and the 99% quantile = 5.47%. (**C**) Concordance correlation coefficients (and their 95% confidence intervals) for $f_2$-statistic residuals vs. the standard normal distribution. 5% and 1% quantiles calculated on the combined dataset with simulated admixture are shown with red and green lines, respectively. The 5% quantile = 0.67, and the 1% quantile = 0.62.

**Fig. S19.**

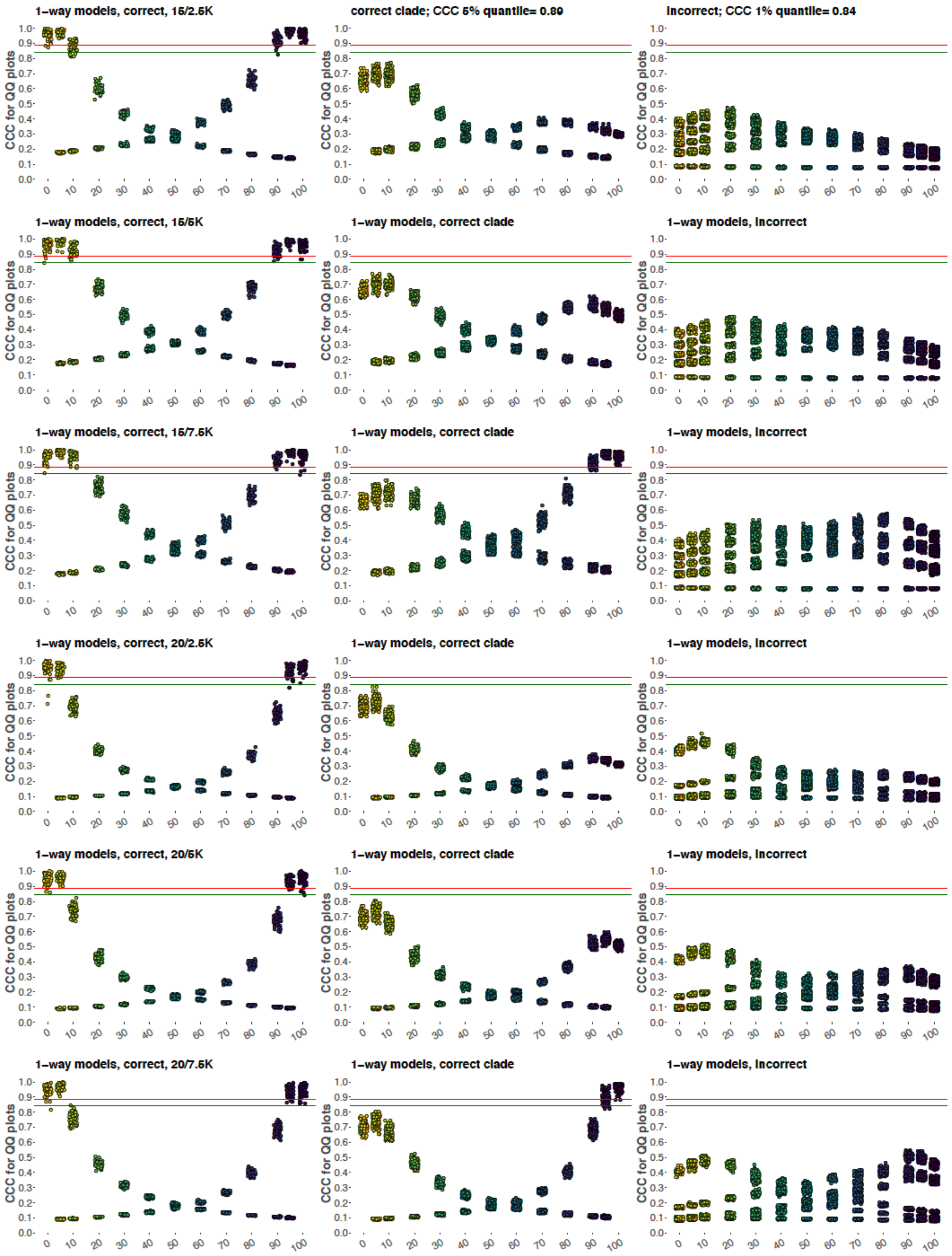
1−way models, correct, 15/2.5K
correct clade; CCC 5% quantile= 0.89
incorrect; CCC 1% quantile= 0.84
1−way models, correct, 15/5K
1−way models, correct clade
1−way models, incorrect
1−way models, correct, 15/7.5K
1−way models, correct clade
1−way models, incorrect
1−way models, correct, 20/2.5K
1−way models, correct clade
1−way models, incorrect
1−way models, correct, 20/5K
1−way models, correct clade
1−way models, incorrect
1−way models, correct, 20/7.5K
1−way models, correct clade
1−way models, incorrect
CCC for QQ plots
0.0 0.1 0.2 0.3 0.4 0.5 0.6 0.7 0.8 0.9 1.0
0 10 20 30 40 50 60 70 80 90 100

Testing an alternative method (based on all *f*-statistics) for assessing absolute model fit to the data on the information-rich graph, 95,000 sites and unadmixed models for group H Concordance correlation coefficients (and their 95% confidence intervals) for $f_2/f_3/f_4$-statistic residuals vs. the standard normal distribution are plotted. 5% and 1% quantiles calculated on the combined dataset without admixture are shown with red and green lines, respectively. The 5% quantile = 0.89, and the 1% quantile = 0.84.

**Fig. S20.**

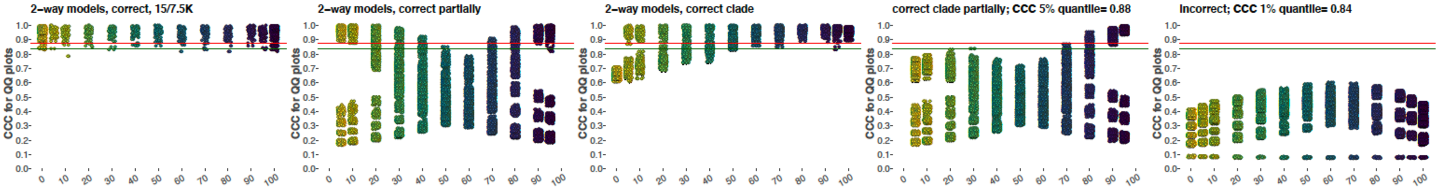


Assessing absolute model fit to the data on the information-rich graph, 95,000 sites and 2-way admixed models for group H. Concordance correlation coefficients (and their 95% confidence intervals) for $f_2/f_3/f_4$-statistic residuals vs. the standard normal distribution. 5% and 1% quantiles calculated on the combined dataset with simulated admixture are shown with red and green lines, respectively. The 5% quantile = 0.88, and the 1% quantile = 0.84.

**Fig. S21.**

**A**

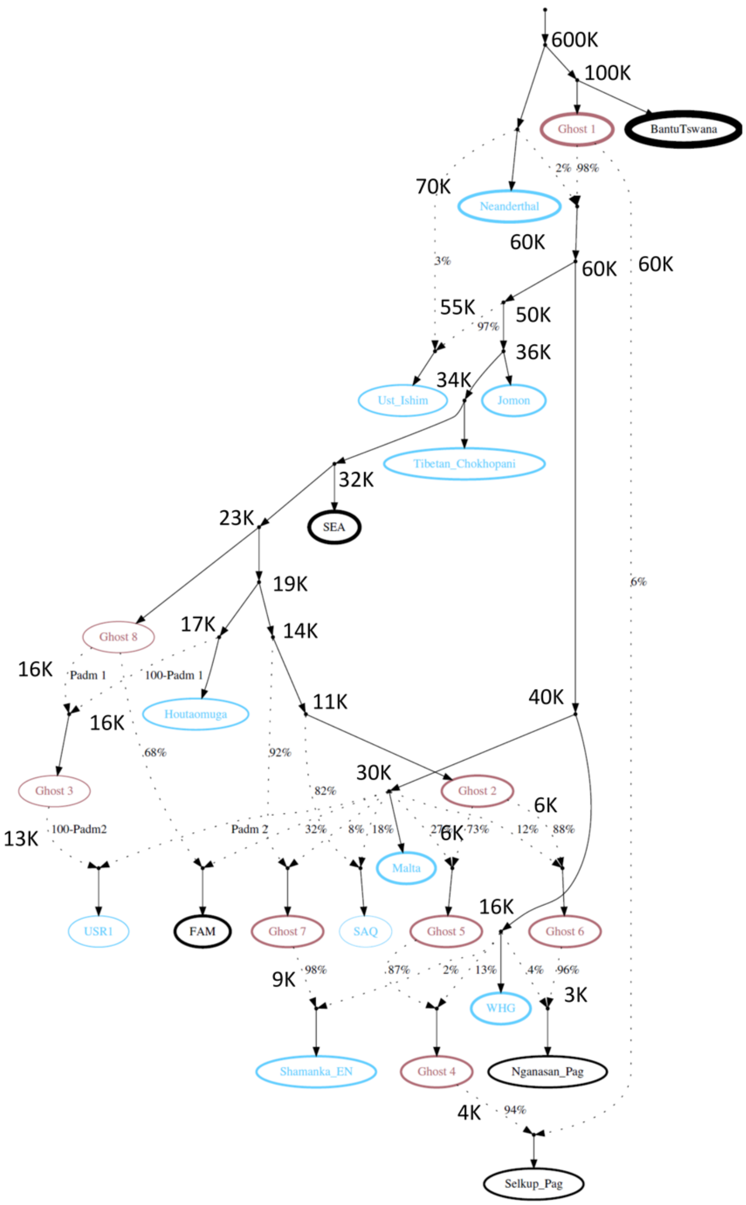


**B**

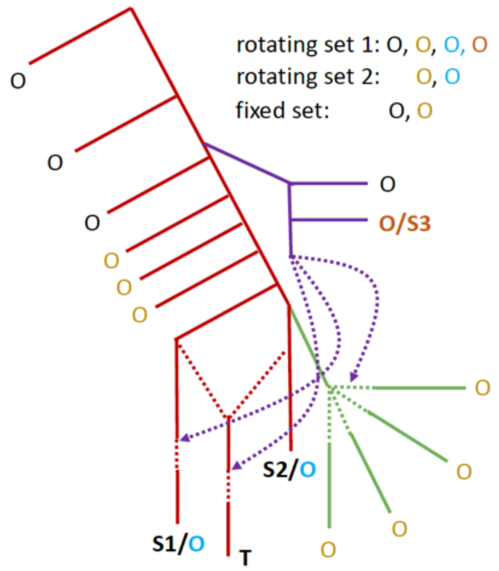


A simulated "realistic" demographic history (**A**) for 15 groups that approximates the skeleton admixture graph used for studying Siberian admixture in Americans, although with some simplifications. Group sizes and dates matched those for the skeleton graph, however for simplicity all simulated individuals were diploid and had no missing data. Effective population sizes are coded by the thickness of population label outlines. Dates in thousands of years ago (K) are shown beside each real or "ghost" group, or gene flow source. Ancient groups are shown in blue, ghost groups in purple, and present-day groups in black. The simulated proportion of Houtaomuga-related ancestry in USR1 was variable (0, 3.5, 7, 14, & 35%), as well as the proportion of Mal'ta-related ancestry in First Americans (FAM, 30, 32, & 40%). All the other admixture proportions and divergence dates were invariant. (**B**) A highly simplified representation of the simulated topology. The following abbreviations are used: O, outgroup; S1, S2 S3, ancestry source proxies for the target T. Three alternative outgroup sets are shown in different colors. Only one clade of outgroups is differentially related to S1 and S2 (green branches).

**Fig. S22.**

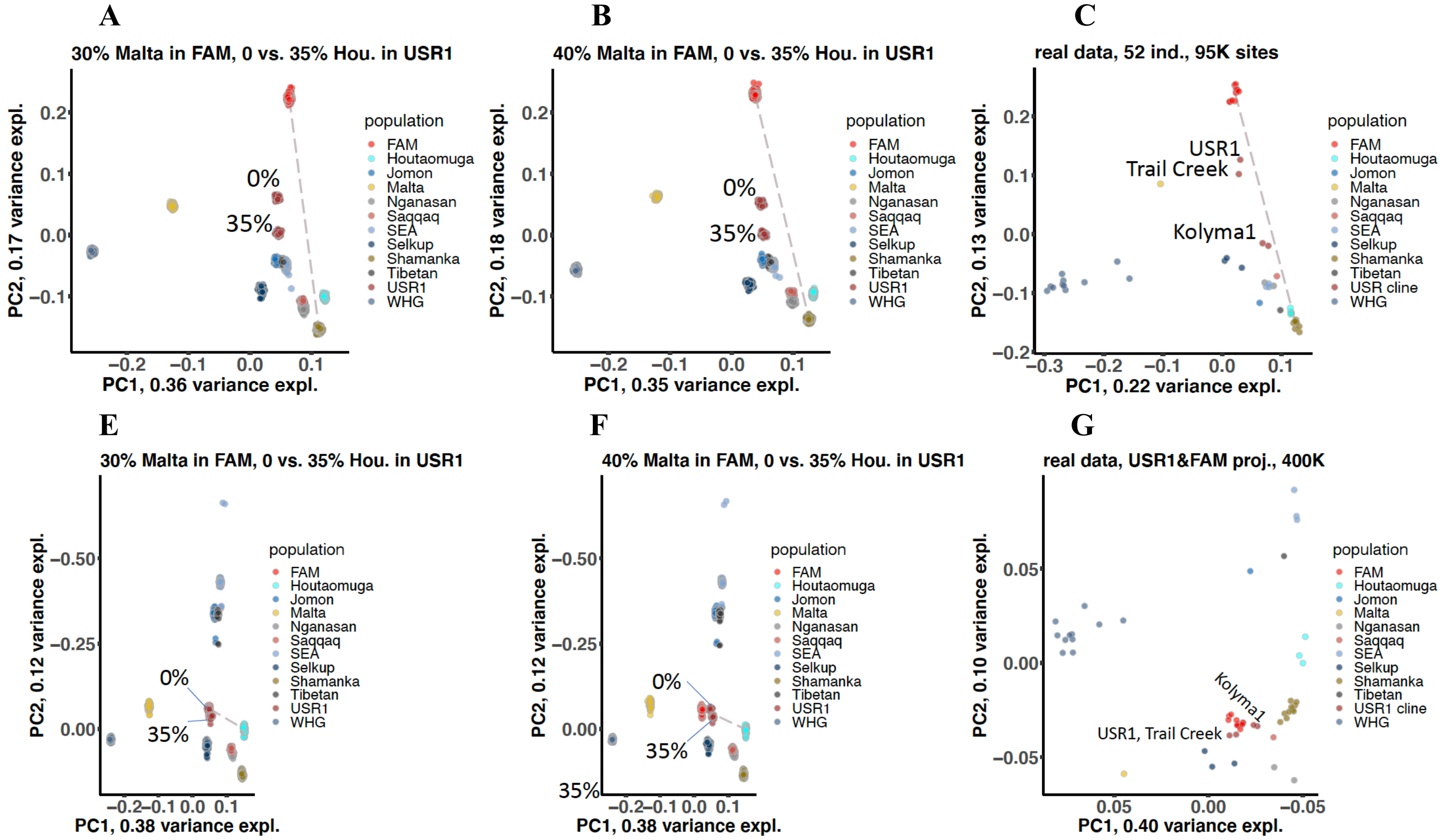


Panels **A-C** show PC1/PC2 plots constructed using *PLINK* on simulated (**A,B**) and real data (**C**). PCs were calculated using nearly identical sets of individuals: 50 simulated ind. (all individuals except for the Neanderthal, Africans, and Ust'-Ishim) and the same number of real counterparts. In the case of the real data, 3 USR1-related individuals were added: Trail Creek and two versions of the Kolyma1 genome. Dataset sizes in terms of sites were also identical, 95,000. Centroids of the FAM and Shamanka EN clusters are connected with a dashed line. Plots for 100 iterations were overlapped. Panels **D-F** show PC1/PC2 plots constructed using *smartPCA* on simulated (**D,E**) and real data (**F**). PCs were calculated using 36 simulated ind. (95,000 sites) or 657 real individuals from Eurasia (~400,000 sites present across all groups plotted). In contrast to panels **A-C**, American groups were projected on the PCs: FAM and USR1; FAM and USR1-related individuals on real data. Panels **A** and **B, E** and **F** differ in the proportion of Malta ancestry in the FAM group. Clusters of USR1 points are labelled according to the simulated proportion of Houtaomuga ancestry. USR1-related individuals are labelled in panels **C** and **G**.

**Fig. S23.**

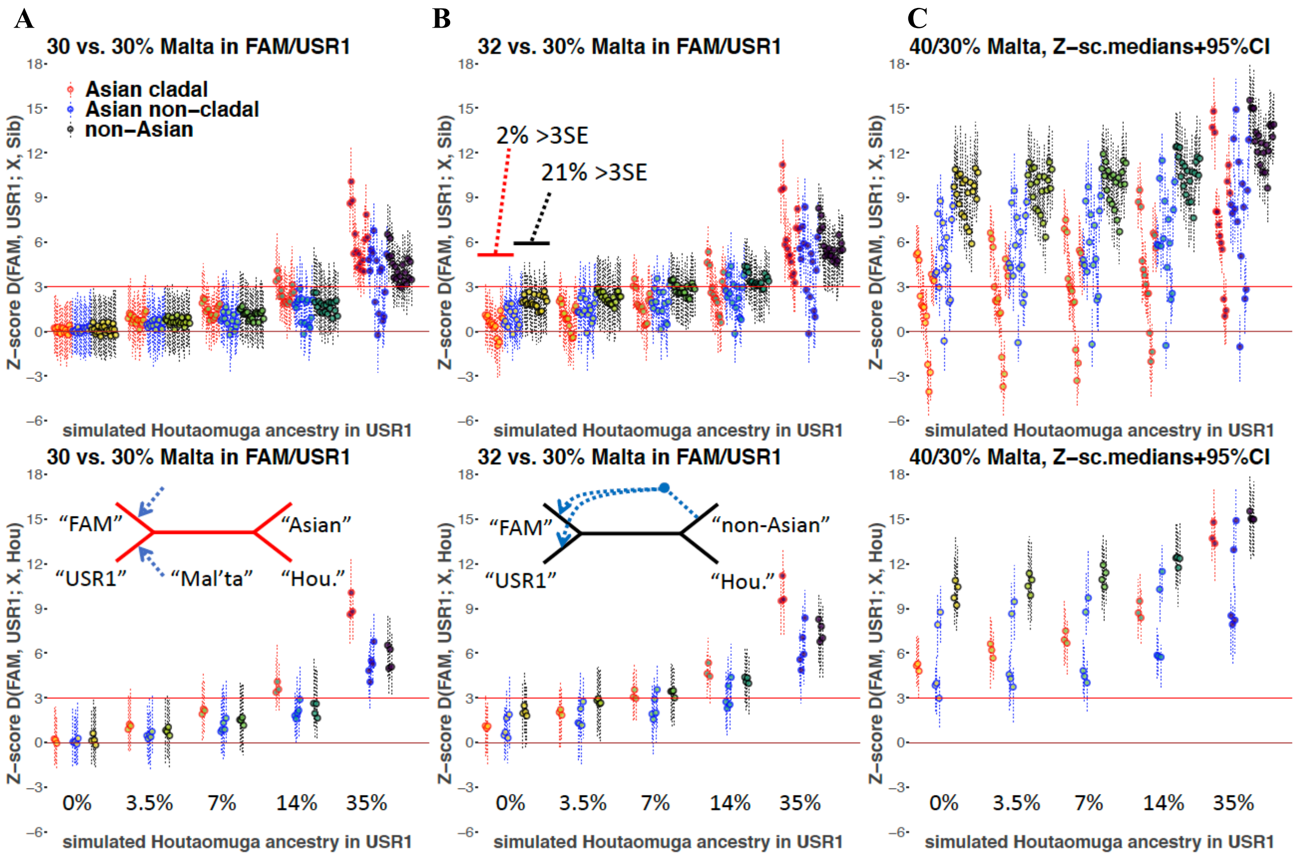


*D*-statistics of the form *D*(FAM, USR1; X, Siberian) or *D*(FAM, USR1; X, Houtaomuga) calculated on simulated data (the complex history, 400,000 sites with the ascertainment bias) are shown in the upper and lower panels, respectively. Median Z-scores and 95% confidence intervals are shown across simulation iterations. Outgroups X are divided into 3 classes color-coded according to the legend: Asians fully cladal with the Siberian ancestry source in USR1 (Houtaomuga); Asians receiving additional gene flows, i.e. not fully cladal with Houtaomuga; and non-Asians. Schematic unrooted trees illustrate the graph topology for the "Asian cladal" and "non-Asian" classes.

**Fig. S24.**

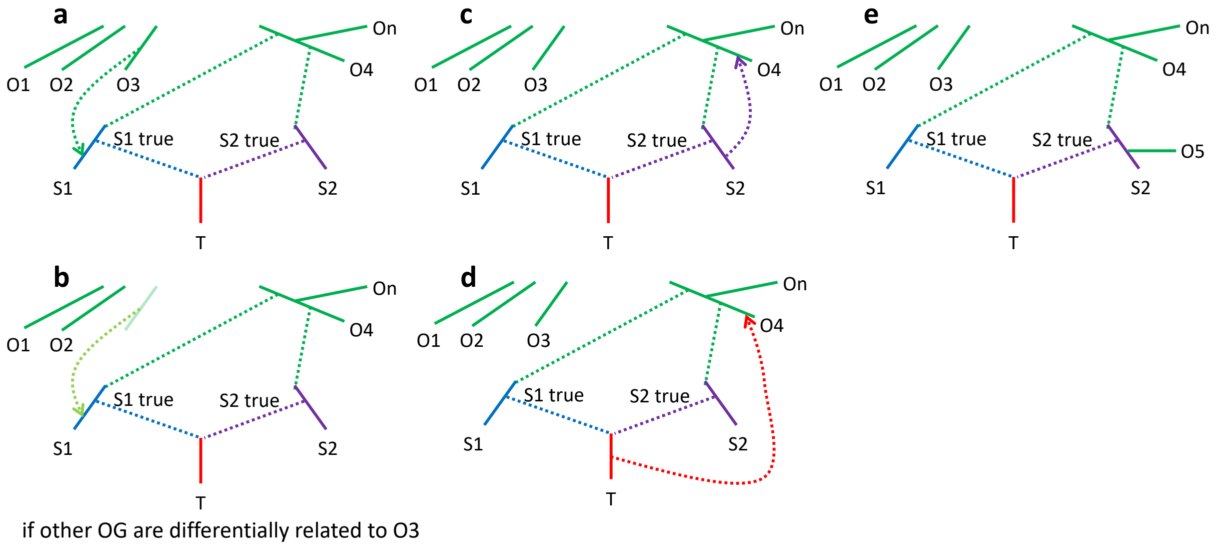


Graphs showing an exhaustive list of assumption violations for a *qpAdm* setup which prioritizes one target group. (**A**) A gene flow from an outgroup to a source proxy after the divergence of the latter from the true source. (**B**) A gene flow from an unsampled source to a source proxy after the divergence of the latter from the true source. This case is problematic if the outgroups are differentially related to the unsampled source. (**C**) A gene flow from a source proxy to an outgroup after the divergence of the former from the true source. (**D**) A gene flow from a target to an outgroup. (**E**) An outgroup cladal with a source proxy.

**Fig. S25.**

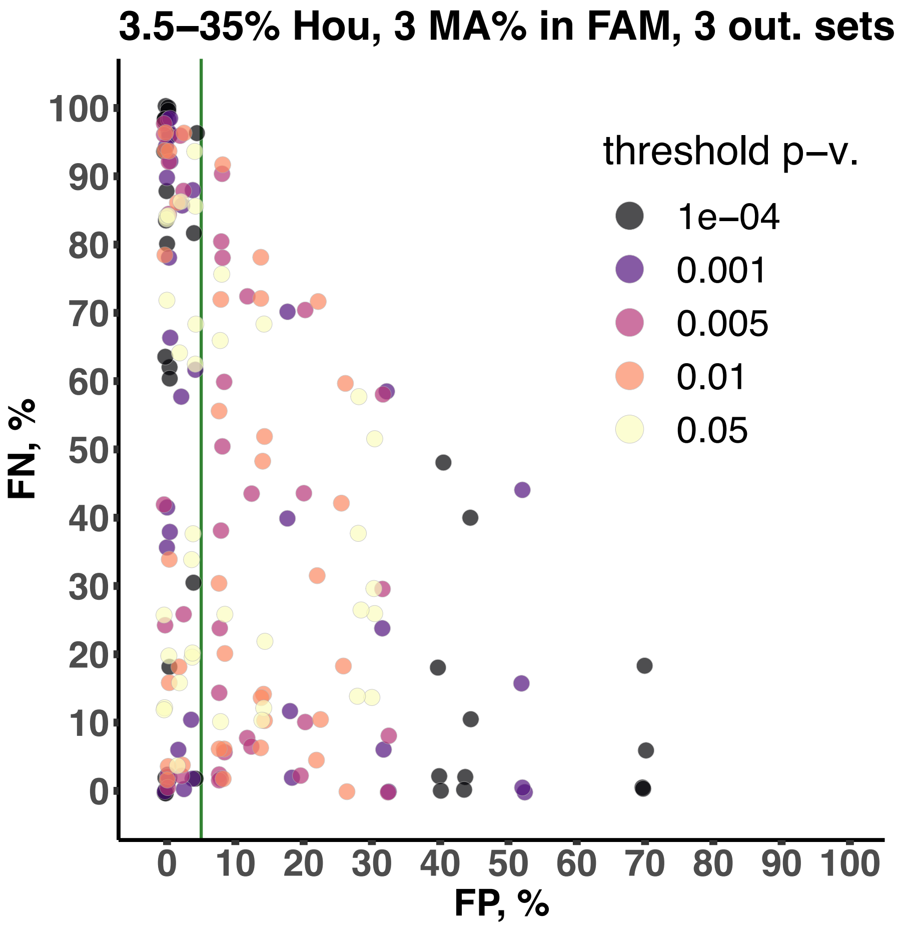


FP and FN rates of the *qpWave*/*qpAdm* procedure that prioritizes one target group for various proportions of Houtaomuga ancestry in USR1 and Mal'ta ancestry ratios in FAM and USR1 (the complex history, 400,000 sites with the ascertainment bias). Each point corresponds to 50 iterations of a particular type of simulated history, given a certain outgroup set, and a *p*-value threshold (color-coded according to the legend). Low-level jitter was added to aid the visualization. Very much similar results were obtained when the condition on inferred admixture proportions was relaxed (the proportions should be non-negative, and not the proportions ± 2SE).

**Fig. S26.**

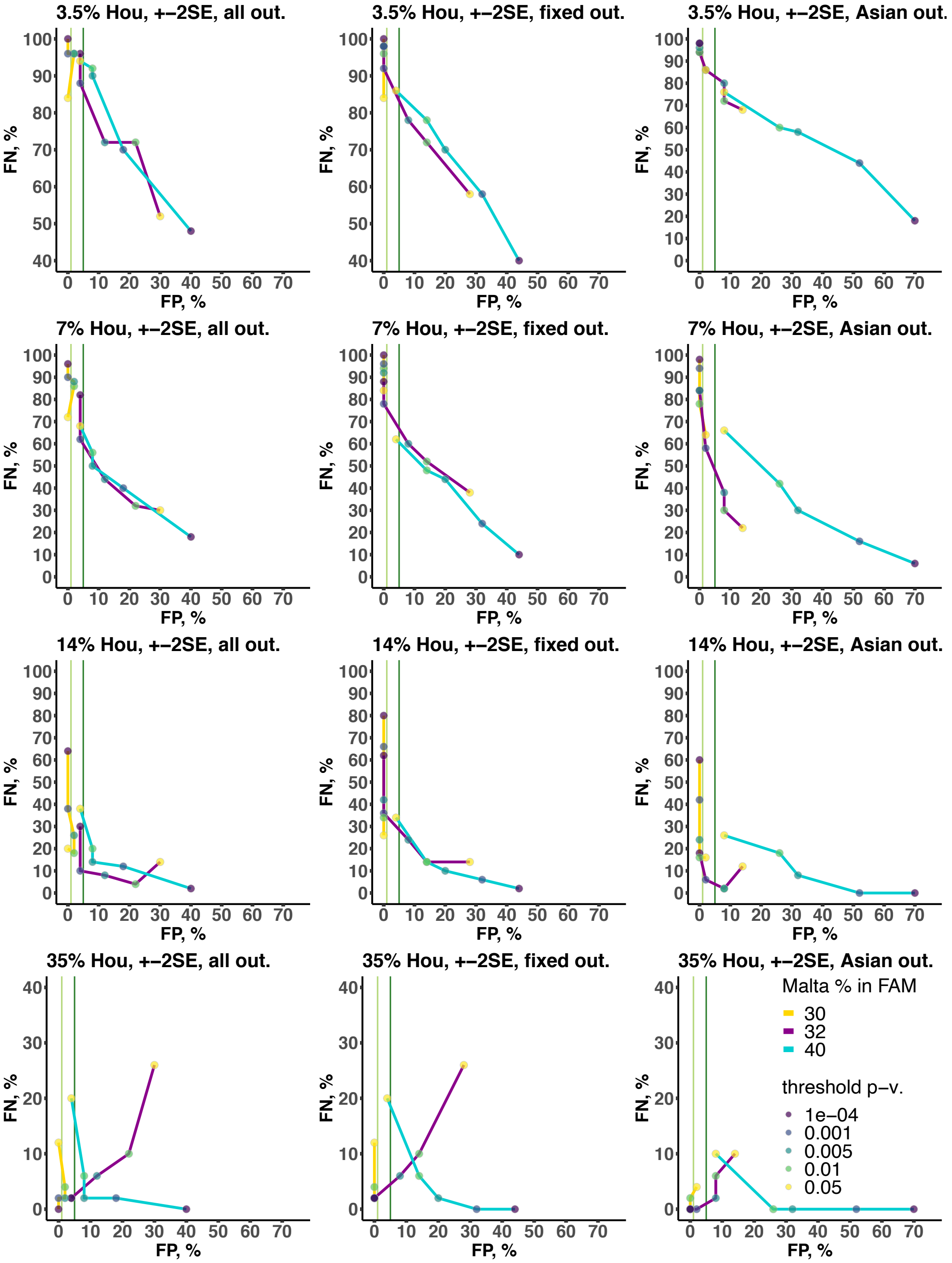


A detailed look at the FP and FN rates of the *qpAdm* procedure that prioritizes one target group (the complex history, 400,000 sites with the ascertainment bias). The following settings were tested: from 3.5% to 35% Houtaomuga ancestry in USR1, various levels of cladality violation (coded by line color), and various *p*-value thresholds (coded by point color). Results for the rotating outgroup set 1 are shown on the left, fixed set in the middle, and rotating set 2 on the left.

**Fig. S27.**

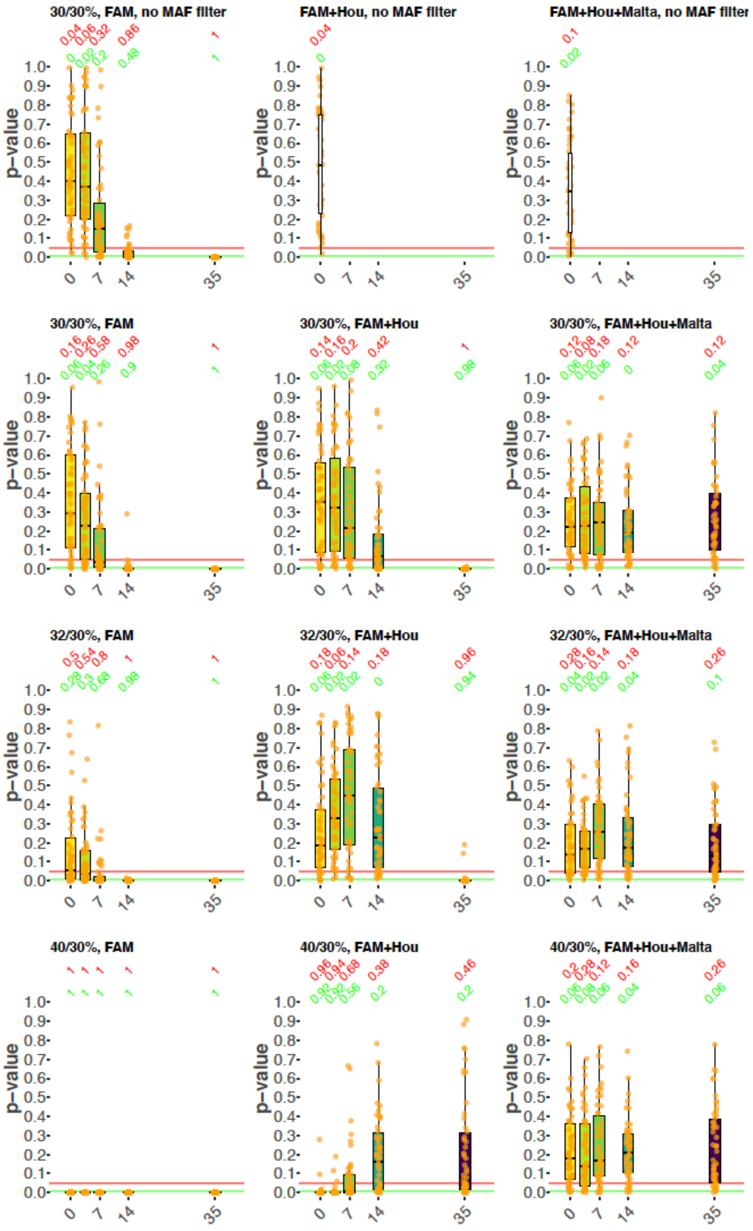


Boxplots illustrating the distribution of *p*-values for *qpAdm* models (the complex demographic history, 400,000 sites). Results for 1-way models USR1=FAM are shown on the left, 2-way models USR1=FAM+Houtaomuga in the middle, and for 3-way models USR1=FAM+Houtaomuga+Mal'ta on the right. Results for different ratios of simulated Mal'ta ancestry in FAM and USR1 are shown. Results for datasets without the ascertainment bias are shown in the top row, and results for datasets with ascertainment bias are shown in the other rows. Simulated Houtaomuga gene flow proportions are shown on the x-axes and coded by the color gradient, and *p*-values are shown on the y-axes. Each dot represents a simulation iteration. The horizontal red and green lines correspond to the 0.05 and 0.01 *p*-value cut-offs, respectively, and the numbers above the plots show the fractions of iterations demonstrating *p*-values below these cut-offs.

**Fig. S28.**

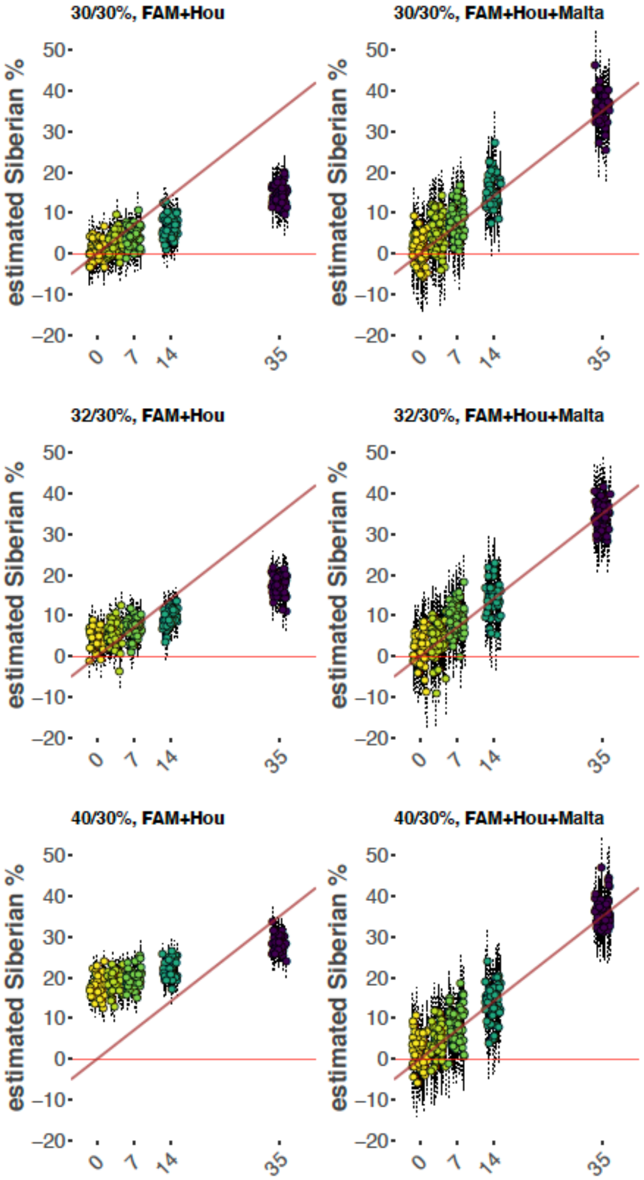


Dot plots illustrating admixture proportions estimated for the complex demographic history (400,000 sites with the ascertainment bias) using *qpAdm*. Results for 2-way models USR1=FAM+Houtaomuga are shown on the left, and for 3-way models USR1=FAM+Houtaomuga+Mal'ta on the right. Results for different ratios of simulated Mal'ta ancestry in FAM and USR1 are shown. Simulated Houtaomuga gene flow proportions are shown on the x-axes and coded by the color gradient. Estimated Houtaomuga ancestry proportions in USR1 are shown on the y-axes, each dot represents a simulation iteration, and double SE intervals are shown as thin dotted lines. Diagonals are shown in brown.

**Fig. S29.**

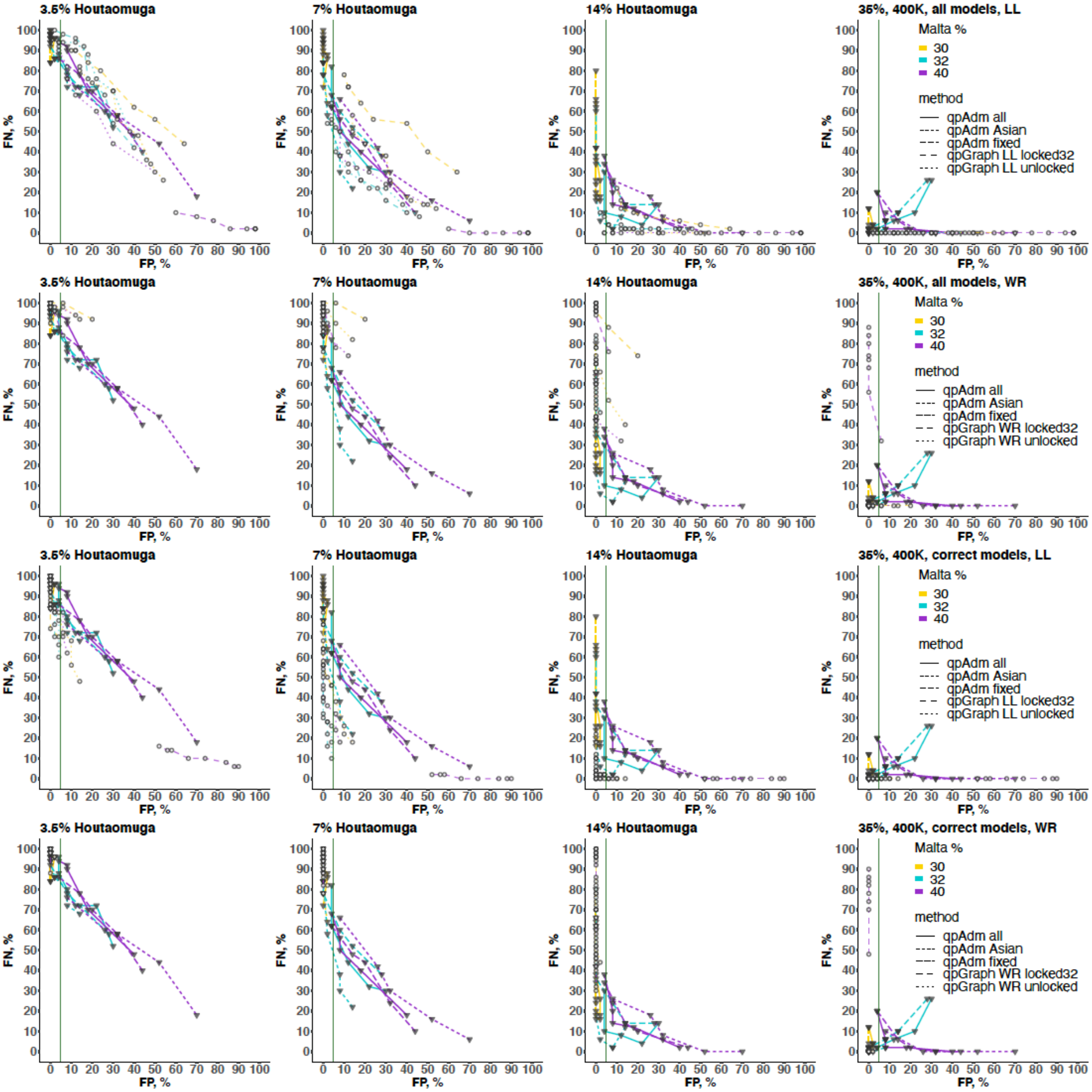


A detailed look at the FP and FN rates of the *qpAdm* procedure that prioritizes one target group and the graph mapping procedure (the complex history, 400,000 sites with the ascertainment bias). The following settings were tested: from 3.5% to 35% Houtaomuga ancestry in USR1, various levels of cladality violation (coded by line color). *qpAdm* points are shown as grey triangles, and *qpGraph* points as white circles. In the case of graph mapping, either only the baseline 2-way and the expected 3-way model were compared (see plots labelled "correct models"), or all possible 3-way models were tested, and the best 3-way model was compared to the baseline 2-way model (see plots labelled "all models"). Only the former protocol is directly comparable to the *qpAdm* tests performed here. Result for either the LL or WR metric are shown. Various *p*-value thresholds were used for the *qpAdm* protocol (see Fig. S26), as well as LL thresholds (from 4 to 11 ln-units) and WR thresholds for the *qpGraph* protocol (from 0.1 to 0.8 SE). *qpAdm* outgroup sets (all = "rotating set 1", Asian = "rotating set 2", fixed = fixed set) and *qpGraph* settings ("locked32" and "unlocked") are coded by line type.

**Fig. S30.**

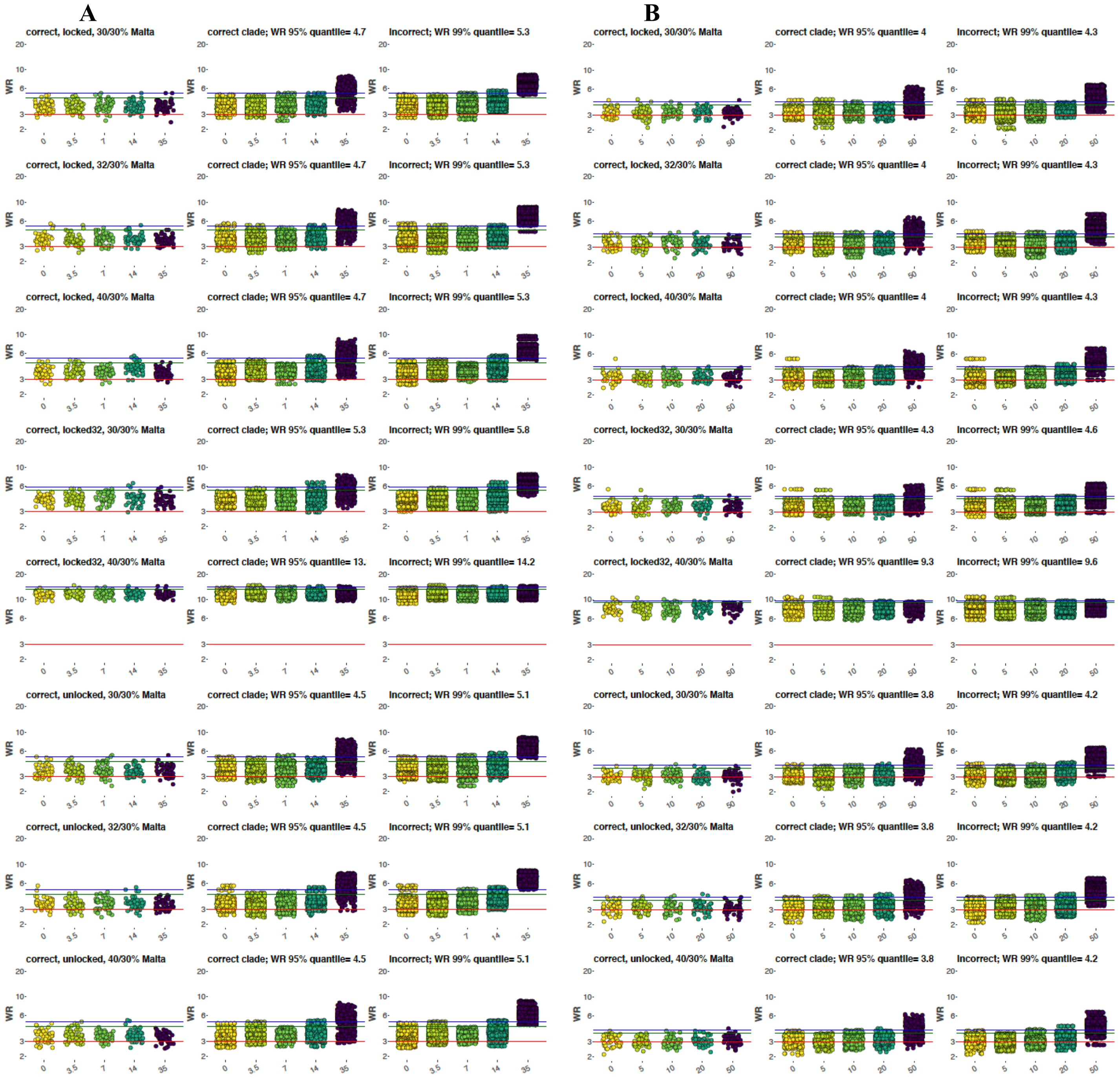
A
correct, locked, 30/30% Malta
correct clade; WR 95% quantile= 4.7
Incorrect; WR 99% quantile= 5.3
correct, locked, 32/30% Malta
correct clade; WR 95% quantile= 4.7
Incorrect; WR 99% quantile= 5.3
correct, locked, 40/30% Malta
correct clade; WR 95% quantile= 4.7
Incorrect; WR 99% quantile= 5.3
correct, locked32, 30/30% Malta
correct clade; WR 95% quantile= 5.3
Incorrect; WR 99% quantile= 5.8
correct, locked32, 40/30% Malta
correct clade; WR 95% quantile= 13.
Incorrect; WR 99% quantile= 14.2
correct, unlocked, 30/30% Malta
correct clade; WR 95% quantile= 4.5
Incorrect; WR 99% quantile= 5.1
correct, unlocked, 32/30% Malta
correct clade; WR 95% quantile= 4.5
Incorrect; WR 99% quantile= 5.1
correct, unlocked, 40/30% Malta
correct clade; WR 95% quantile= 4.5
Incorrect; WR 99% quantile= 5.1
B
correct, locked, 30/30% Malta
correct clade; WR 95% quantile= 4
Incorrect; WR 99% quantile= 4.3
correct, locked, 32/30% Malta
correct clade; WR 95% quantile= 4
Incorrect; WR 99% quantile= 4.3
correct, locked, 40/30% Malta
correct clade; WR 95% quantile= 4
Incorrect; WR 99% quantile= 4.3
correct, locked32, 30/30% Malta
correct clade; WR 95% quantile= 4.3
Incorrect; WR 99% quantile= 4.6
correct, locked32, 40/30% Malta
correct clade; WR 95% quantile= 9.3
Incorrect; WR 99% quantile= 9.6
correct, unlocked, 30/30% Malta
correct clade; WR 95% quantile= 3.8
Incorrect; WR 99% quantile= 4.2
correct, unlocked, 32/30% Malta
correct clade; WR 95% quantile= 3.8
Incorrect; WR 99% quantile= 4.2
correct, unlocked, 40/30% Malta
correct clade; WR 95% quantile= 3.8
Incorrect; WR 99% quantile= 4.2
WR

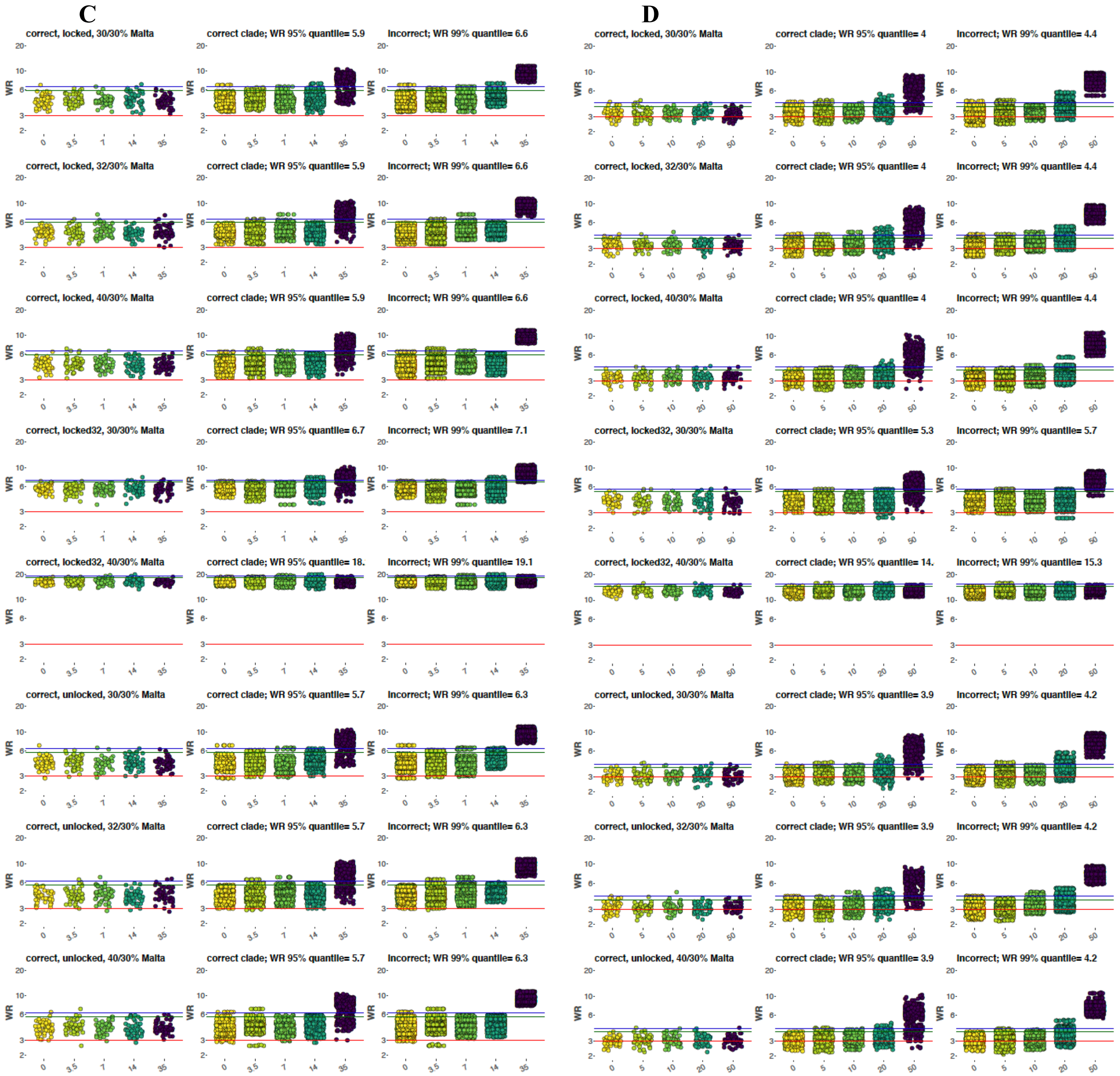


Worst $f_4$-statistic residuals for assessing absolute model fit to the data on the complex history, 3-way admixed models for USR1 ($FAM_{EA}$ + Mal'ta + X), 95,000 sites with (**A**) or without the ascertainment bias (**B**) and 400,000 sites with (**C**) or without the ascertainment bias (**D**). Iterations are grouped by simulated Houtaomuga admixture proportions on the x-axis, and worst residual values are plotted on the y-axis in log10 scale. The traditional threshold of 3 SE is shown with red lines, and 95% and 99% quantiles calculated on the combined dataset with simulated admixture (3.5% to 35% Houtaomuga proportion in USR1, all settings combined) are shown with green and blue lines, respectively. Models are classified into "correct" (X=Houtaomuga), "correct clade" (X=any edge in the Siberian clade), and "incorrect" ones (X=any other edge), and results are plotted separately for these model classes. Results for the "locked", "locked32", and "unlocked" settings are shown (see Table S9).

**Fig. S32.**

**A**

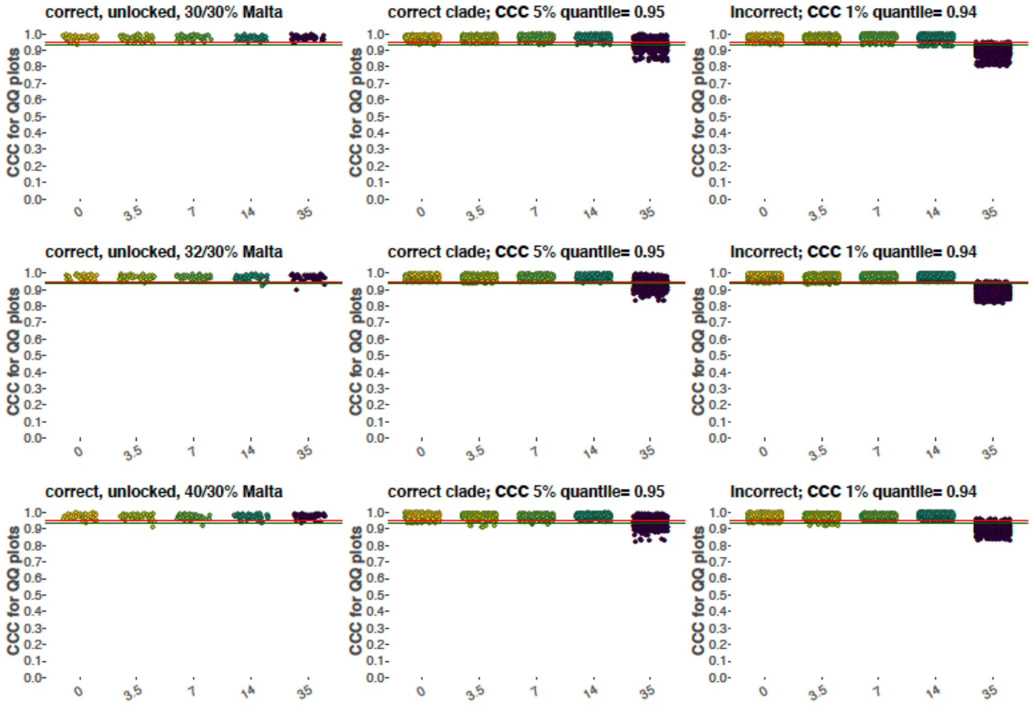
correct, unlocked, 30/30% Malta
correct clade; CCC 5% quantile= 0.95
incorrect; CCC 1% quantile= 0.94
correct, unlocked, 32/30% Malta
correct clade; CCC 5% quantile= 0.95
incorrect; CCC 1% quantile= 0.94
correct, unlocked, 40/30% Malta
correct clade; CCC 5% quantile= 0.95
incorrect; CCC 1% quantile= 0.94
CCC for QQ plots
1.0
0.9
0.8
0.7
0.6
0.5
0.4
0.3
0.2
0.1
0.0
0
3.5
7
14
35

**B**

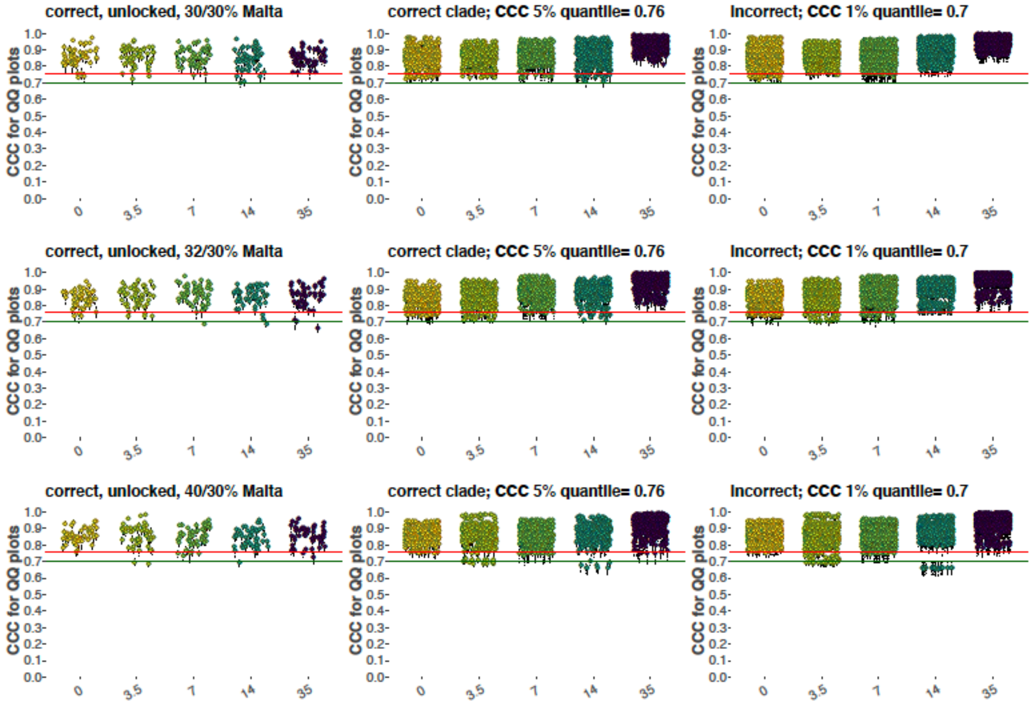
correct, unlocked, 30/30% Malta
correct clade; CCC 5% quantile= 0.76
incorrect; CCC 1% quantile= 0.7
correct, unlocked, 32/30% Malta
correct clade; CCC 5% quantile= 0.76
incorrect; CCC 1% quantile= 0.7
correct, unlocked, 40/30% Malta
correct clade; CCC 5% quantile= 0.76
incorrect; CCC 1% quantile= 0.7
CCC for QQ plots
1.0
0.9
0.8
0.7
0.6
0.5
0.4
0.3
0.2
0.1
0.0
0
3.5
7
14
35

C

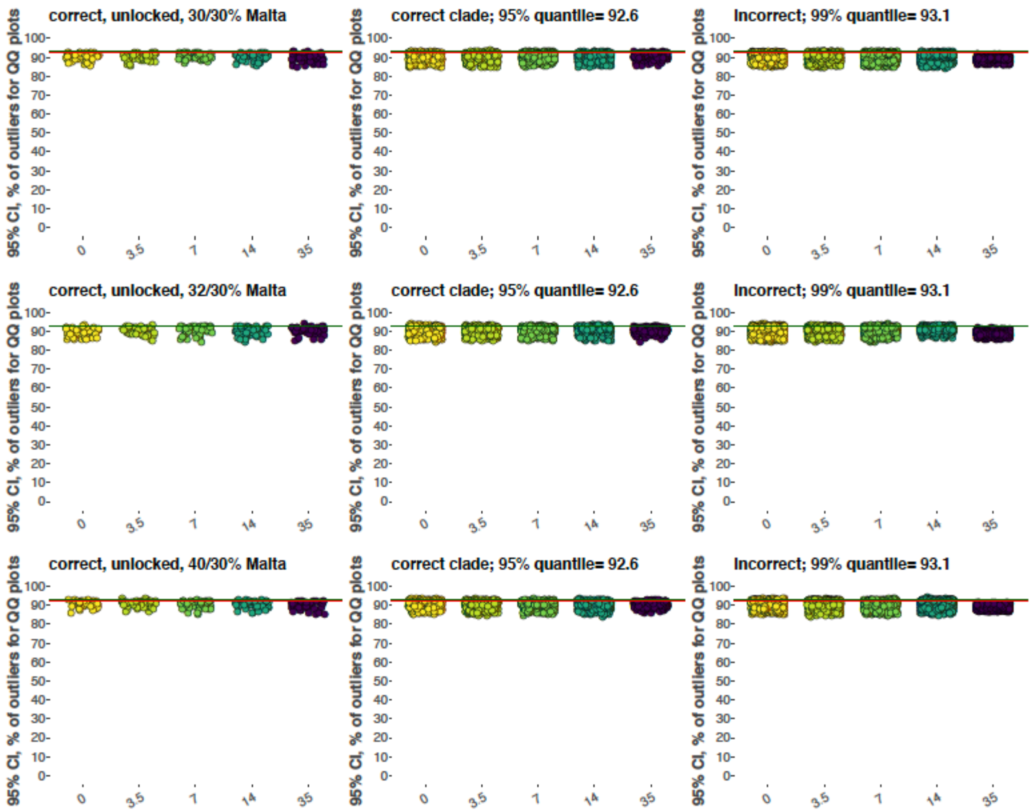


Assessing absolute model fit to the data on the complex graph, 400,000 sites with the ascertainment bias, the "unlocked" setting, and 3-way admixed models for USR1. Three alternative methods were tested. (**A**) Concordance correlation coefficients (and their 95% confidence intervals) for $f_2/f_3/f_4$-statistic residuals vs. the standard normal distribution. 5% and 1% quantiles calculated on the combined dataset with simulated admixture are shown with red and green lines, respectively. The 5% quantile = 0.95, and the 1% quantile = 0.94. (**B**) Concordance correlation coefficients for $f_2$-statistic residuals vs. the standard normal distribution. (**C**) Percentage of $f_2/f_3/f_4$-statistic residuals that lie outside of the 95% confidence interval around the standard normal distribution. 95% and 99% quantiles calculated on the combined dataset with simulated admixture are shown with red and green lines, respectively. The 95% quantile = 92.6%, and the 99% quantile = 93.1%.

**Fig. S40.**

**A**

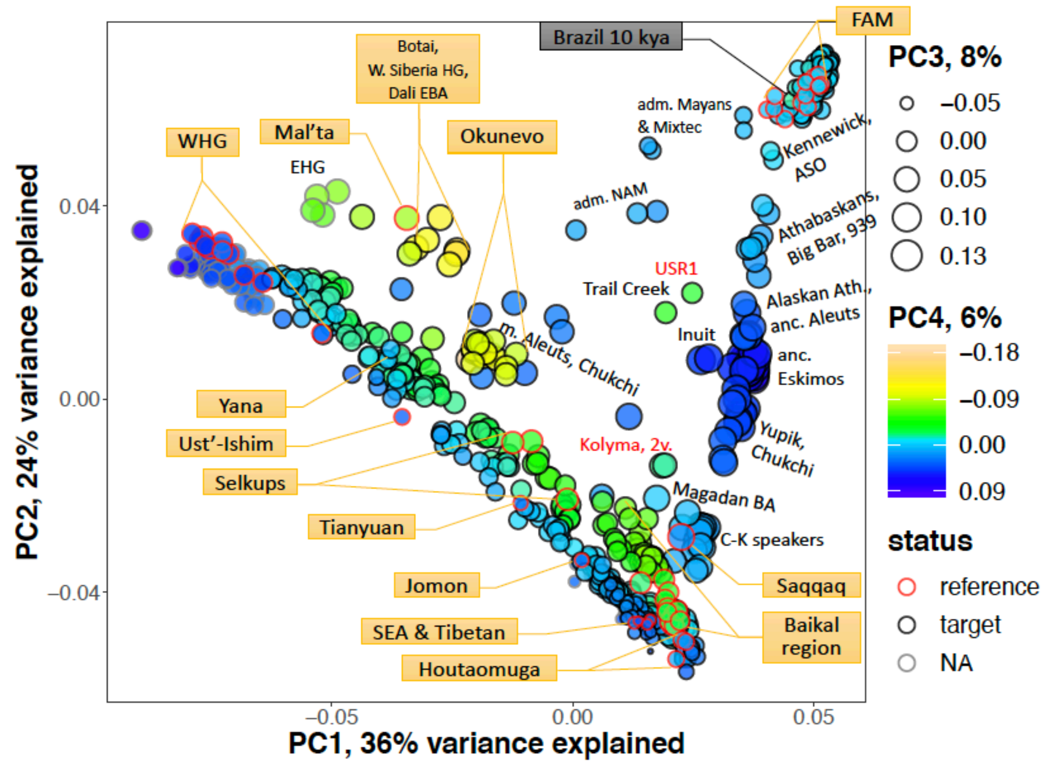

FAM
Brazil 10 kya
Botai, W. Siberia HG, Dali EBA
WHG
Mal'ta
Okunevo
EHG
adm. Mayans & Mixtec
Kennewick, ASO
adm. NAM
Athabaskans, Big Bar, 939
USR1
Trail Creek
Alaskan Ath., anc. Aleuts
Inuit
anc. Eskimos
m. Aleuts, Chukchi
Yupik, Chukchi
Yana
Ust'-Ishim
Kolyma, 2v.
Selkups
Magadan BA
Tianyuan
C-K speakers
Jomon
Saqqaq
SEA & Tibetan
Baikal region
Houtaomuga
PC1, 36% variance explained
PC2, 24% variance explained
PC3, 8%
PC4, 6%
status
reference
target
NA


**B**

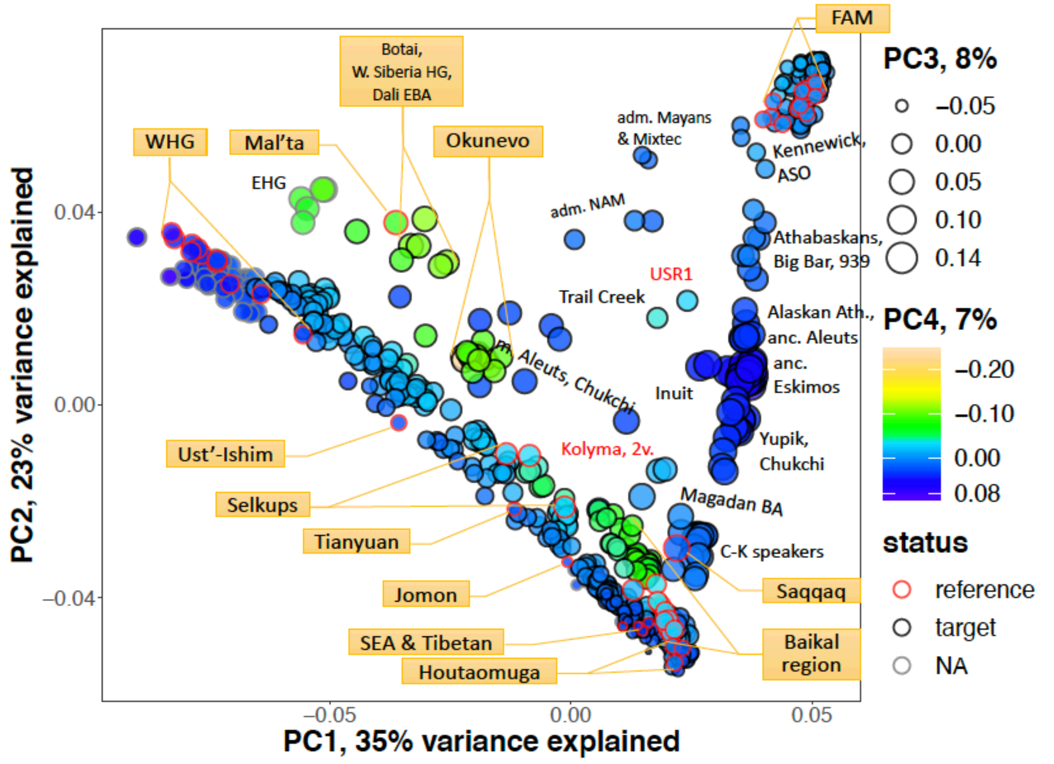

FAM
Botai, W. Siberia HG, Dali EBA
WHG
Mal'ta
Okunevo
EHG
adm. Mayans & Mixtec
Kennewick, ASO
adm. NAM
Athabaskans, Big Bar, 939
USR1
Trail Creek
Alaskan Ath., anc. Aleuts
anc. Eskimos
m. Aleuts, Chukchi
Inuit
Yupik, Chukchi
Ust'-Ishim
Kolyma, 2v.
Selkups
Magadan BA
Tianyuan
C-K speakers
Jomon
Saqqaq
SEA & Tibetan
Baikal region
Houtaomuga
PC1, 35% variance explained
PC2, 23% variance explained
PC3, 8%
PC4, 7%
status
reference
target
NA

C

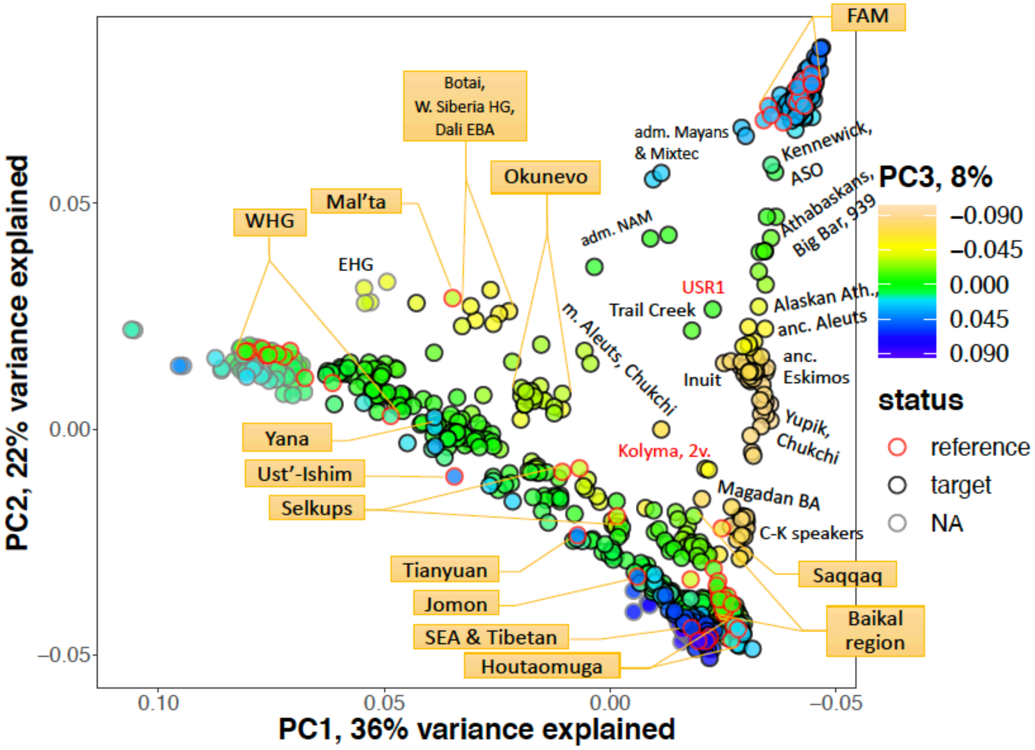
FAM
Botai, W. Siberia HG, Dali EBA
adm. Mayans & Mixtec
Kennewick, ASO
Okunevo
Mal'ta
WHG
EHG
adm. NAM
Athabaskans, Big Bar, 939
USR1
Trail Creek
Alaskan Ath., anc. Aleuts
m. Aleuts, Chukchi
anc. Eskimos
Inuit
Yupik, Chukchi
Yana
Ust'-Ishim
Selkups
Kolyma, 2v.
Magadan BA
C-K speakers
Tianyuan
Jomon
SEA & Tibetan
Houtaomuga
Saqqaq
Baikal region
PC3, 8%
−0.090
−0.045
0.000
0.045
0.090
status
reference
target
NA
PC2, 22% variance explained
PC1, 36% variance explained
0.05
0.00
−0.05
0.10

D

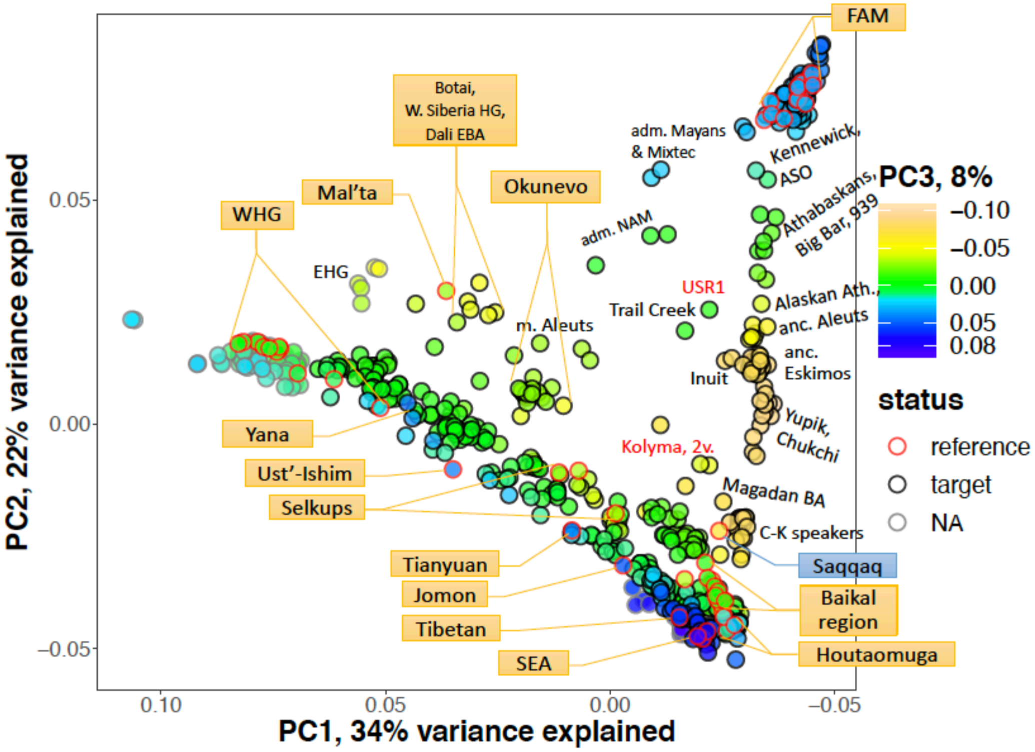
FAM
Botai, W. Siberia HG, Dali EBA
adm. Mayans & Mixtec
Kennewick, ASO
Okunevo
Mal'ta
WHG
EHG
adm. NAM
Athabaskans, Big Bar, 939
USR1
Trail Creek
m. Aleuts
Alaskan Ath., anc. Aleuts
anc. Eskimos
Inuit
Yupik, Chukchi
Yana
Ust'-Ishim
Selkups
Kolyma, 2v.
Magadan BA
C-K speakers
Saqqaq
Tianyuan
Jomon
Tibetan
SEA
Baikal region
Houtaomuga
PC3, 8%
−0.10
−0.05
0.00
0.05
0.08
status
reference
target
NA
PC2, 22% variance explained
PC1, 34% variance explained
0.05
0.00
−0.05
0.10

E

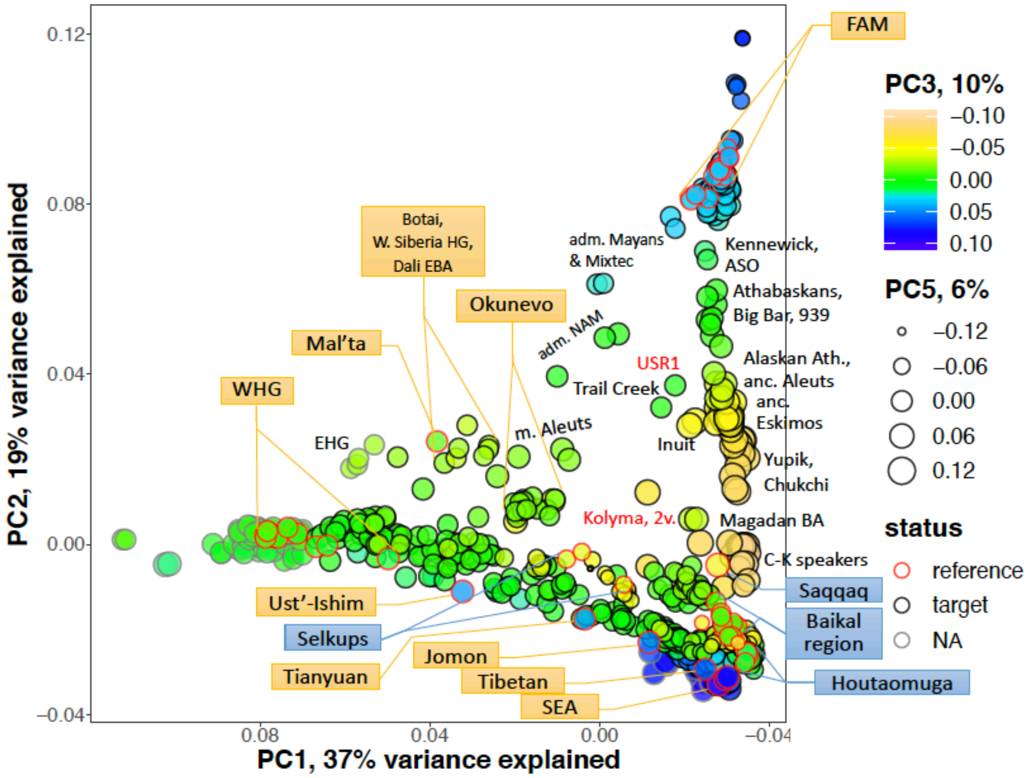

PC2, 19% variance explained
PC1, 37% variance explained
0.12
0.08
0.04
0.00
–0.04
FAM
PC3, 10%
–0.10
–0.05
0.00
0.05
0.10
PC5, 6%
–0.12
–0.06
0.00
0.06
0.12
status
reference
target
NA
Botai, W. Siberia HG, Dali EBA
adm. Mayans & Mixtec
Kennewick, ASO
Okunevo
Athabaskans, Big Bar, 939
Mal'ta
adm. NAM
USR1
Alaskan Ath., anc. Aleuts
anc. Eskimos
WHG
Trail Creek
EHG
m. Aleuts
Inuit
Yupik, Chukchi
Kolyma, 2v.
Magadan BA
C-K speakers
Saqqaq
Ust'-Ishim
Selkups
Baikal region
Jomon
Tianyuan
Tibetan
SEA
Houtaomuga


F

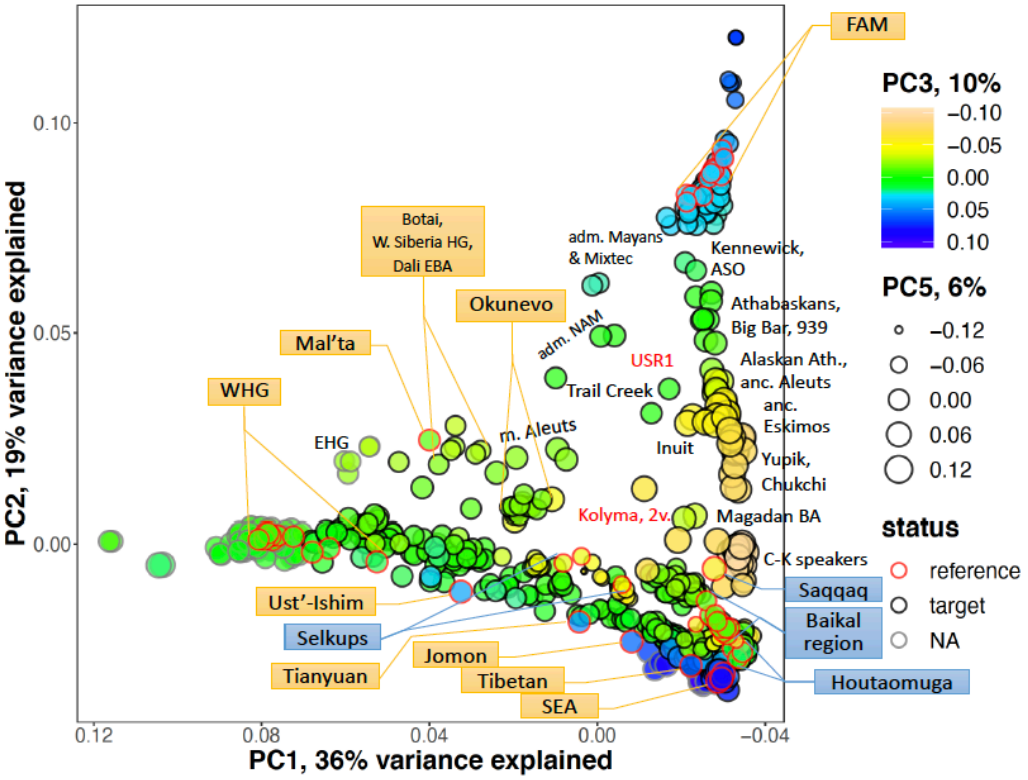

PC2, 19% variance explained
PC1, 36% variance explained
0.10
0.05
0.00
0.12
0.08
0.04
0.00
–0.04
FAM
PC3, 10%
–0.10
–0.05
0.00
0.05
0.10
PC5, 6%
–0.12
–0.06
0.00
0.06
0.12
status
reference
target
NA
Botai, W. Siberia HG, Dali EBA
adm. Mayans & Mixtec
Kennewick, ASO
Okunevo
Athabaskans, Big Bar, 939
Mal'ta
adm. NAM
USR1
Alaskan Ath., anc. Aleuts
anc. Eskimos
WHG
Trail Creek
EHG
m. Aleuts
Inuit
Yupik, Chukchi
Kolyma, 2v.
Magadan BA
C-K speakers
Saqqaq
Ust'-Ishim
Selkups
Baikal region
Jomon
Tianyuan
Tibetan
SEA
Houtaomuga

PCA analyses incorporating ancient and present-day Eurasians and Americans. The analyses were based on the dataset without transitions (**A, C, E**) or on the full dataset (**B, D, F**) used for graph mapping. The following protocols were used: 1) 10 PCs were calculated using PLINK v. 1.90b6.10 based on both ancient and present-day individuals (**A, B**); 2) 10 PCs were calculated using smartPCA v.16000 based on both ancient and present-day individuals, with the "shrinkmode: YES" option (**C, D**); 3) 10 PCs were calculated using smartPCA v.16000 based on present-day individuals, and ancient individuals were projected on the PCs, with the "shrinkmode: YES" option (**E, F**). Individual coordinates on PC3 and PC4 (**A, B**) or PC3 and PC5 (**E, F**) are visualized using marker size and marker color, respectively. In the case of the 2$^{nd}$ setup, PC4 and higher PCs are not visualized (**C, D**) since they separated just few individuals from the others. Individuals that were mapped on the skeleton graphs as targets are circled in black, and individuals belonging to the groups incorporated into the 16-population skeleton graphs are circled in red. Individuals not included into the admixture graph analyses are circled in grey. Key individuals and clusters of individuals are labelled by text and callouts. The Kolyma1 and USR1 individuals are labelled in red. Two versions of the Kolyma1 genotypes are plotted. The following abbreviations are used: 2v., two genotype versions for the same individual; adm., admixed (having colonial European admixture); anc., ancient; ASO, the ancient South Ontario individual (*18*); Ath., Athabaskans; BA, Bronze Age; C-K speakers, peoples speaking languages of the Chukotko-Kamchatkan family; EBA, early Bronze Age; EHG, eastern European hunter-gatherers *sensu* Haak *et al.* (*48*); FAM, First Peoples of the Americas; kya, thousands of years ago; m., modern; NAM, northern North Americans *sensu* Raghavan *et al.* (*15*) and Scheib *et al.* (*18*); SEA, Southeast Asians; W. Siberia HG, West Siberian hunter-gatherers *sensu* Narasimhan *et al*. (*30*); WHG, West European hunter-gatherers *sensu* Lazaridis *et al.* (*140*).

**Fig. S41.**

**A**

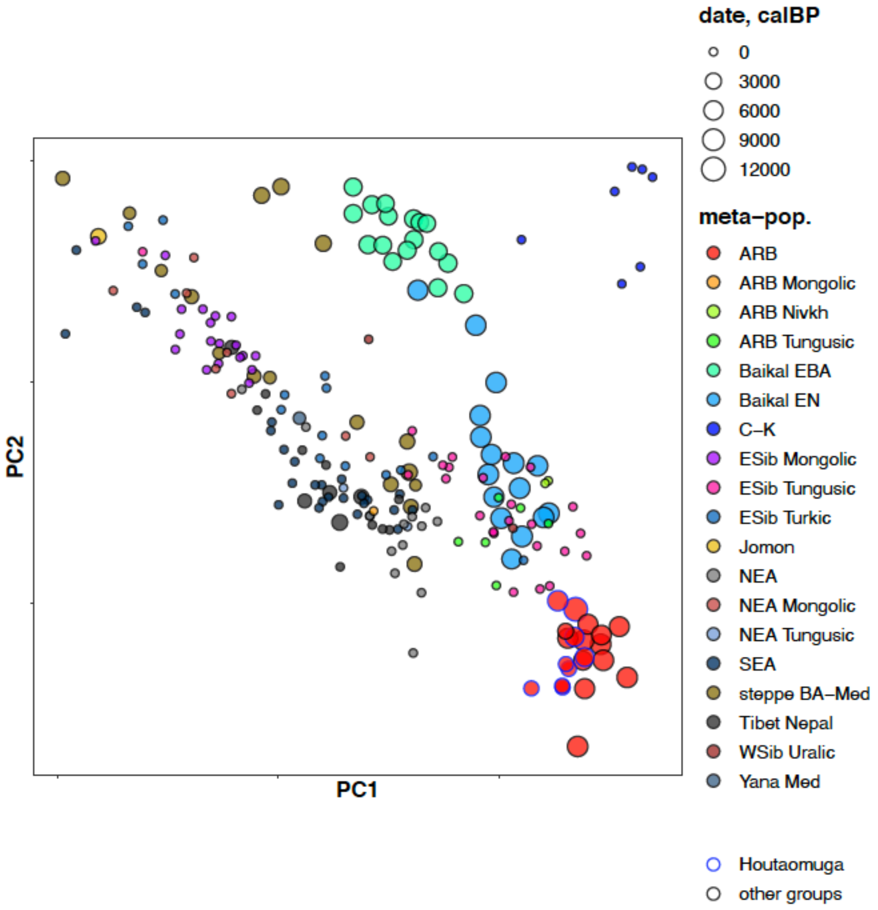

date, calBP
0
3000
6000
9000
12000
meta–pop.
ARB
ARB Mongolic
ARB Nivkh
ARB Tungusic
Baikal EBA
Baikal EN
C–K
ESib Mongolic
ESib Tungusic
ESib Turkic
Jomon
NEA
NEA Mongolic
NEA Tungusic
SEA
steppe BA–Med
Tibet Nepal
WSib Uralic
Yana Med
Houtaomuga
other groups
PC1
PC2

**B**

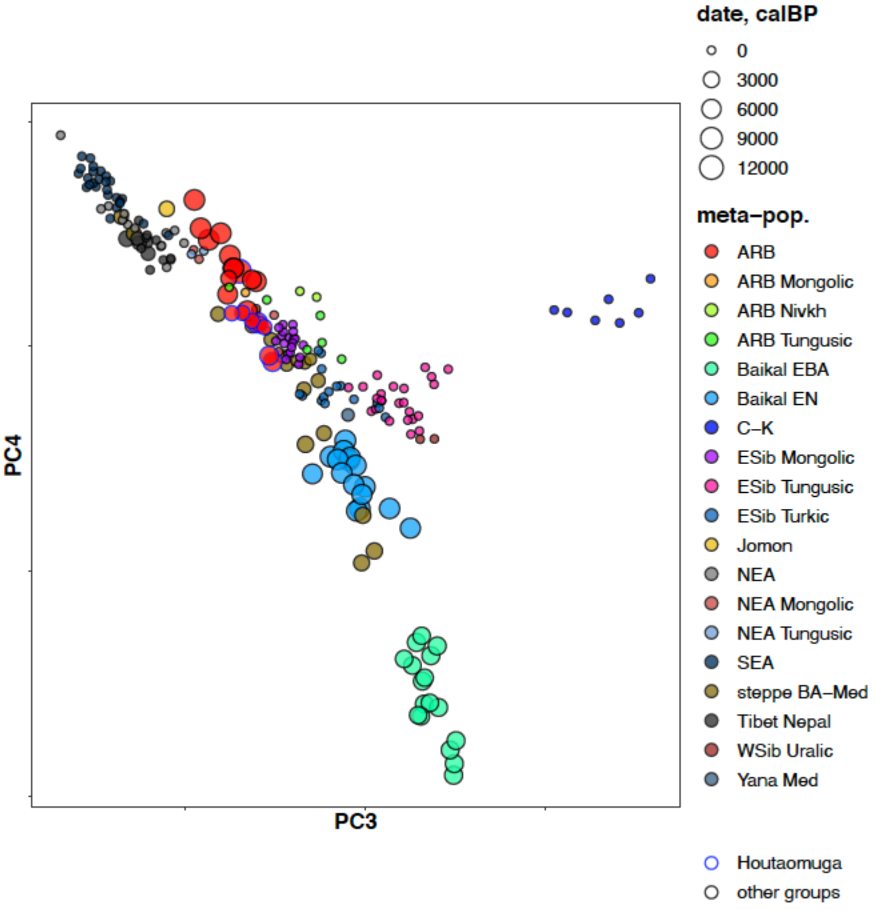


Amur River Basin groups in PC space. This is a zoomed-in version of the PCA plot shown in Fig. S40a. PC1 vs. PC2 (**A**) and PC3 vs. PC4 (**B**) planes are shown. Date in cal yr BP is coded by marker size, geographical and linguistic meta-populations are color-coded according to the legend. Houtaomuga individuals are circled in blue. The following abbreviations are used: ARB, Amur River Basin; EBA, early Bronze Age; EN, early Neolithic; C-K, Chukotko-Kamchatkan speakers; ESib, East Siberian; NEA, northern East Asians; SEA, Southeast Asians; Med, Medieval; WSib, West Siberian.

**Fig. S42.**

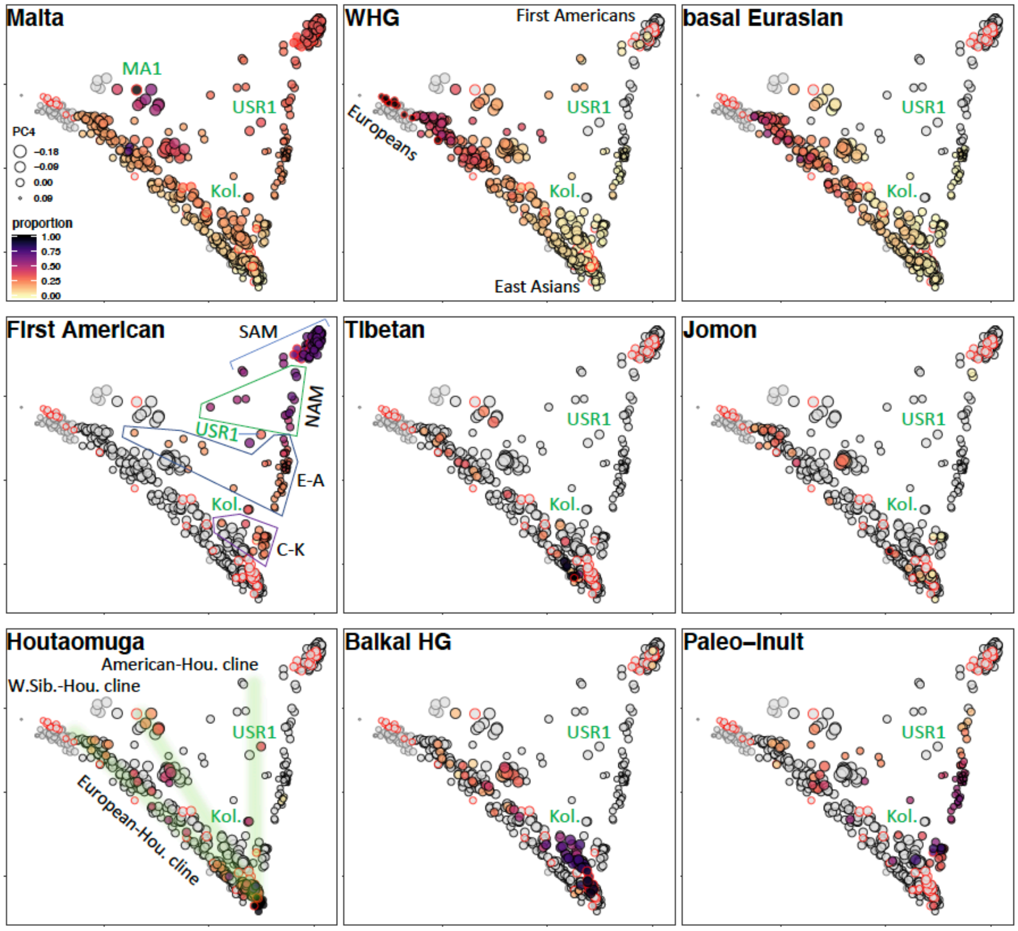


Proportions of selected ancestry sources inferred using the admixture graph approach mapped on the PC1 vs. PC2 plot. Coordinates on PC4 that reflect Mal'ta-related ancestry are visualized using marker size. The ancestry sources are indicated in each panel, and proportions are color-coded according to the legend. Individuals having 0% of a given ancestry source are shown as grey circles with black or red border. Individuals that were mapped on the skeleton graphs as targets are circled in black, and individuals belonging to the groups incorporated into the 16-population skeleton graph are circled in red. Individuals not included into the admixture graph analyses are circled in grey. The USR1 and Kolyma1 (Kol.) individuals are labelled in green. In the Houtaomuga panel, three clines are highlighted: between Houtaomuga and Europeans, West Siberian HG, and First Americans. In the American panel, major population clusters are outlined: C-K, Chukotko-Kamchatkan speakers and Paleo-Inuit; E-A, Eskimo-Aleut Speakers; NAM, northern North Americans including Athabaskans; SAM, the other First Americans. The graph mapping results were taken from Data S6 and S9-1, and the PCA from Fig. S40. If several alternative sources were equally well supported for a given target, all were plotted, and the highest proportion for each source found among the winning models was used.

**Fig. S43.**

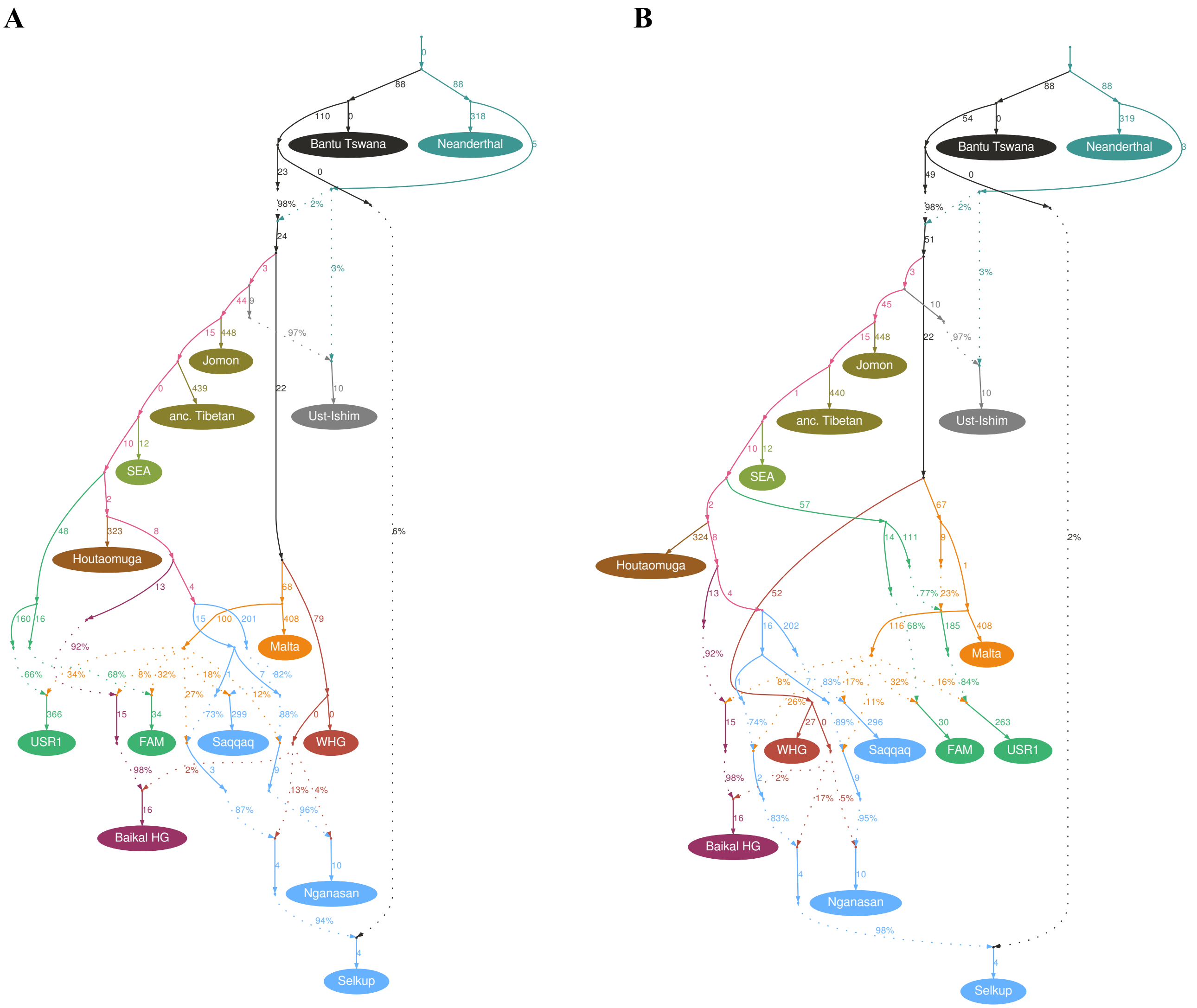
A
B
Bantu Tswana
Neanderthal
Jomon
anc. Tibetan
Ust-Ishim
SEA
Houtaomuga
Malta
USR1
FAM
Saqqaq
WHG
Baikal HG
Nganasan
Selkup

**C**

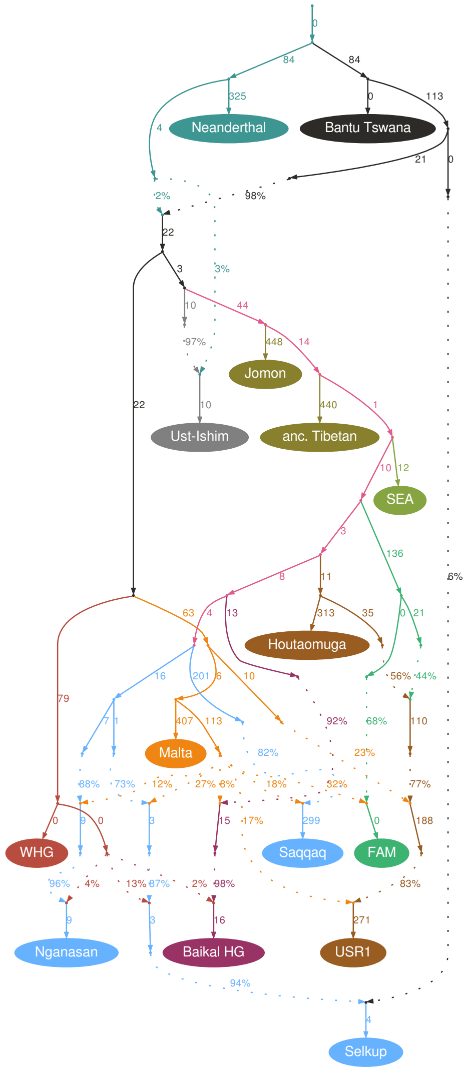


Best two- (**A**), three- (**B**), and four-way (**C**) models for USR1 (the pseudohaploid version, the dataset without transitions) mapped on the 14-population skeleton graph. The USR1 branch is colored according to its largest ancestry component. The following settings were used: West Eurasian gene flows into Siberians were not locked, "useallsnps NO", the Houtaomuga group was composed of two individuals. For the worst residuals and other model statistics see Data S4a.

**Fig. S44.**

**A**

Z-score = 5.86, 17 pops., 12 adm. events, 9 unlocked, 407,407 sites used

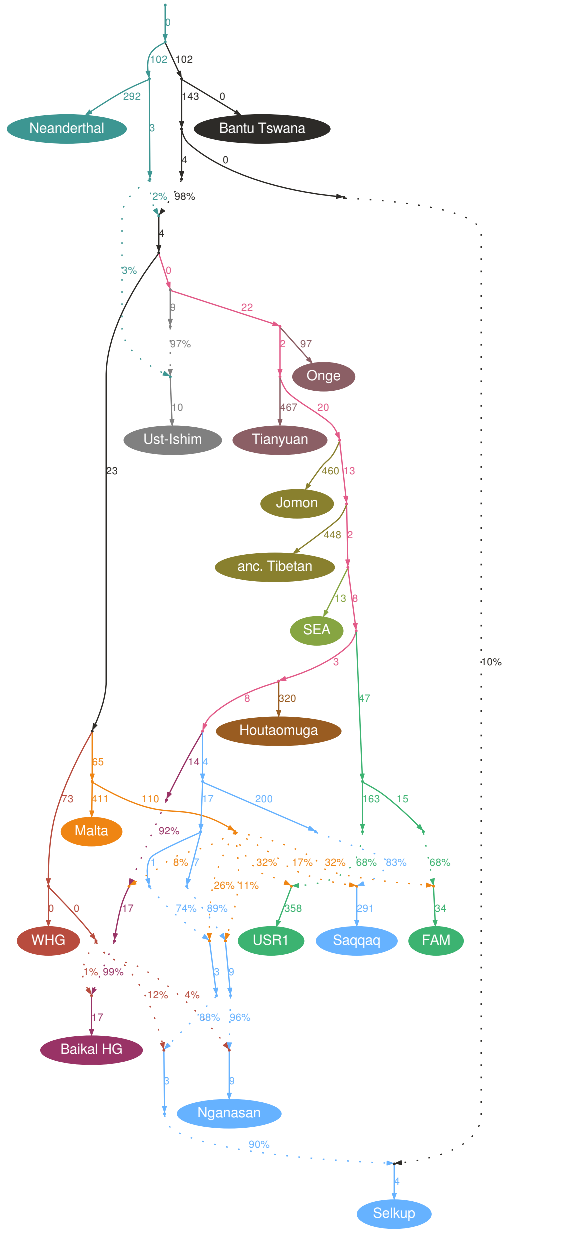


**B**

Z-score = 5.87, 17 pops., 13 adm. events, 10 unlocked, 407,407 sites used

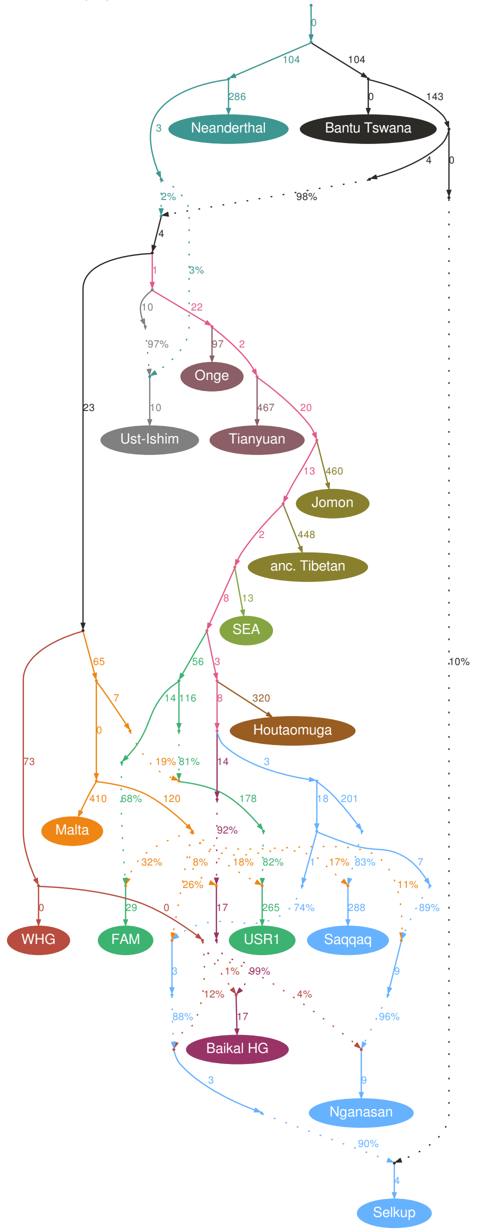

**C**

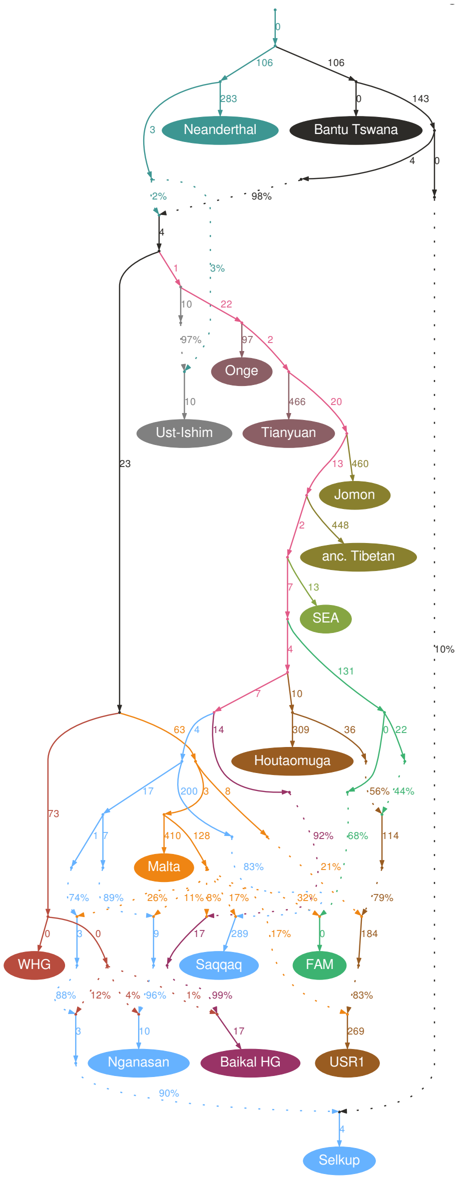


Best two- (**A**), three- (**B**), and four-way (**C**) models for USR1 (the pseudohaploid version, the dataset including transitions) mapped on the 16-population skeleton graph including Tianyuan and Onge. The USR1 branch is colored according to its largest ancestry component. The following settings were used: West Eurasian gene flows into Siberians were not locked, "useallsnps NO", the Houtaomuga group was composed of two individuals. For the worst residuals and other model statistics see Data S4b.

Fig. S45.

A

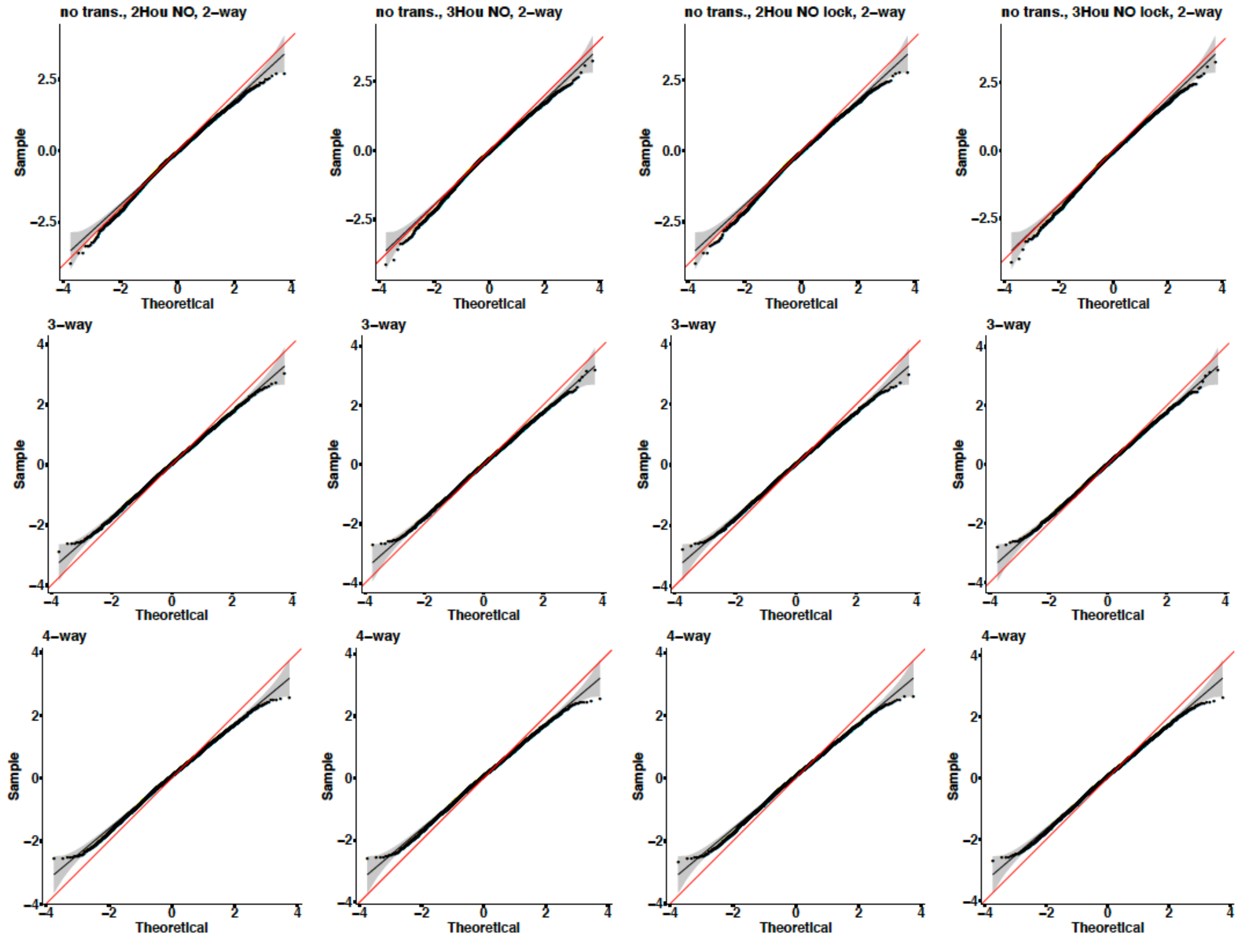
no trans., 2Hou NO, 2–way
no trans., 3Hou NO, 2–way
no trans., 2Hou NO lock, 2–way
no trans., 3Hou NO lock, 2–way
3–way
4–way
Sample
Theoretical

**B**

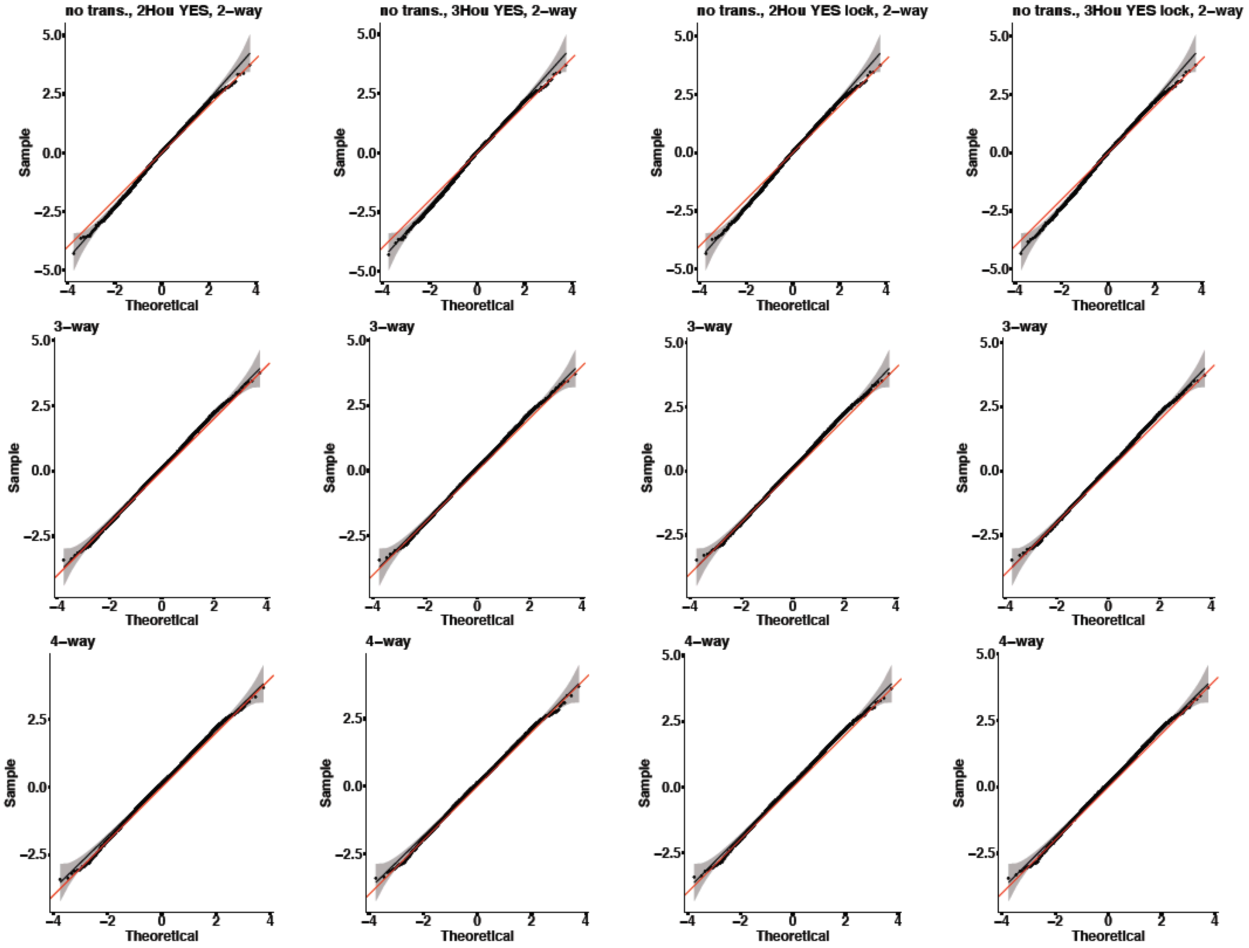
no trans., 2Hou YES, 2−way
no trans., 3Hou YES, 2−way
no trans., 2Hou YES lock, 2−way
no trans., 3Hou YES lock, 2−way
3−way
4−way
Sample
Theoretical

**C**

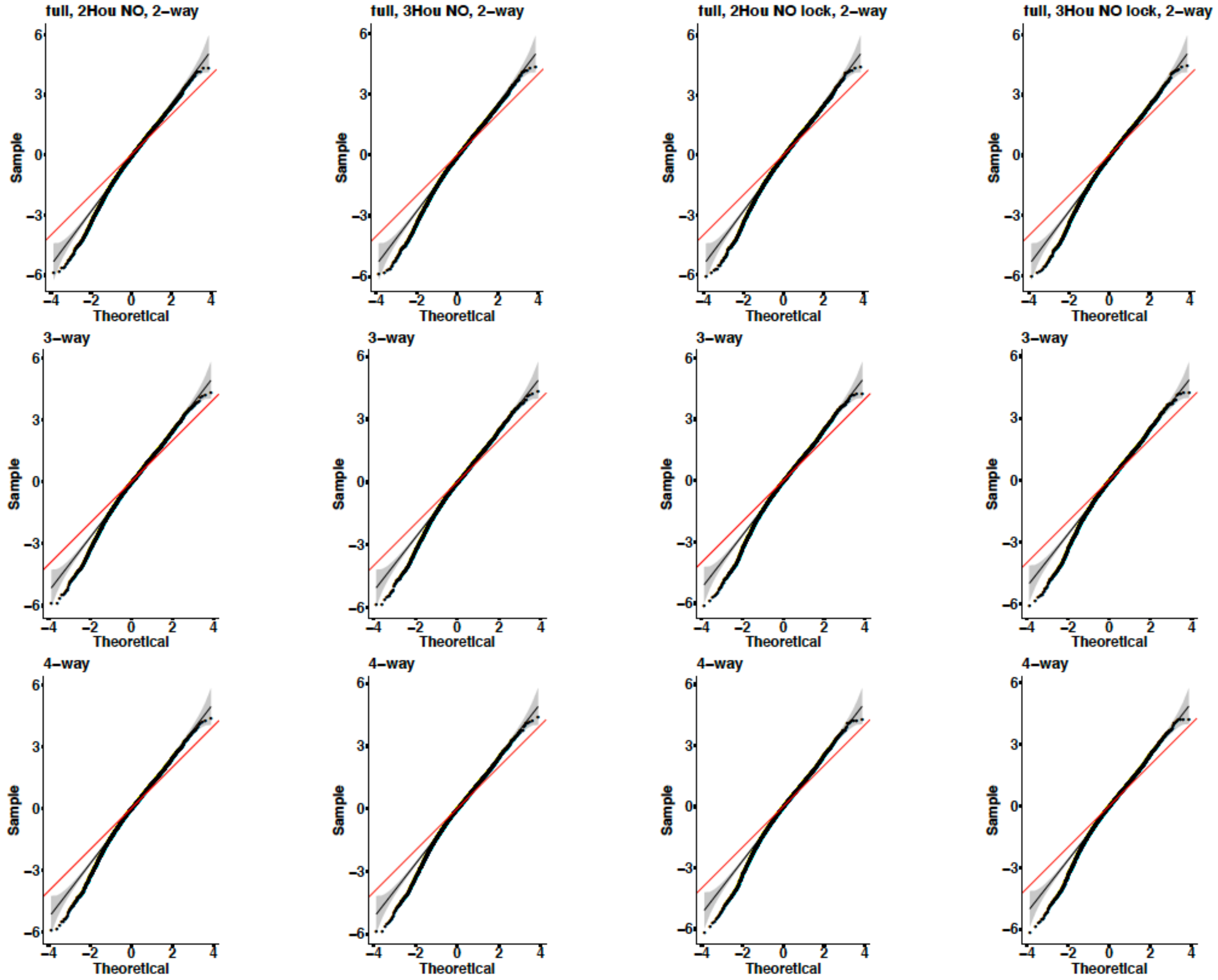


Quantile-quantile plots for all *f*-statistic residuals. The following USR1 models were considered: $FAM_{EA}$ + Mal'ta (labelled as "2-way"); $FAM_{EA}$ + "basal Mal'ta" + Mal'ta (labelled as "3-way"); $FAM_{EA}$ + Houtaomuga + "basal Mal'ta" + Mal'ta (labelled as "4-way"). *qpGraph* settings are indicated in top panels: NT stands for the dataset without transitions (**A**, **B**), "full" for the full dataset (**C**); 2Hou and 3Hou for two and three Houtaomuga individuals, respectively; NO (**A**, **C**) and YES (**B**) for the "useallsnps" setting; "lock" for locked West Eurasian gene flows into Siberian branches. The gray area shows the 95% confidence interval around the general normal distribution (black line). The red line shows the expectation for the standard normal distribution (SD =1).

**Fig. S46.**

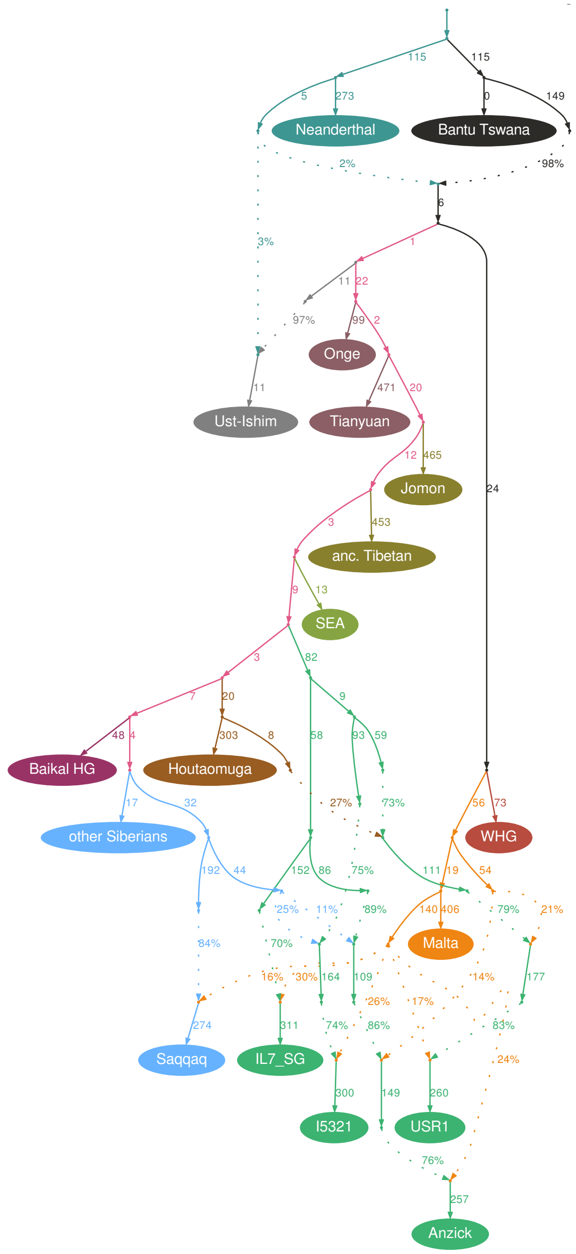


The best topology of the FAM clade ((I5321, USR1), (Clovis, IL7)) selected among 15 alternative topologies tested under 24 setting/dataset combinations. The following settings were used for the graph shown here: 1) the West Eurasian gene flows into Siberians were not locked; 2) external gene flows into the four American individuals were not locked; 3) transitions were kept; 4) "useallsnps: NO"; 5) the Houtaomuga group was composed of two individuals. The Nganasan and Selkup lineages are collapsed under the "other Siberians" label for making the visualization simpler. For the worst residuals and other model statistics see Data S4f.

**Fig. S47.**

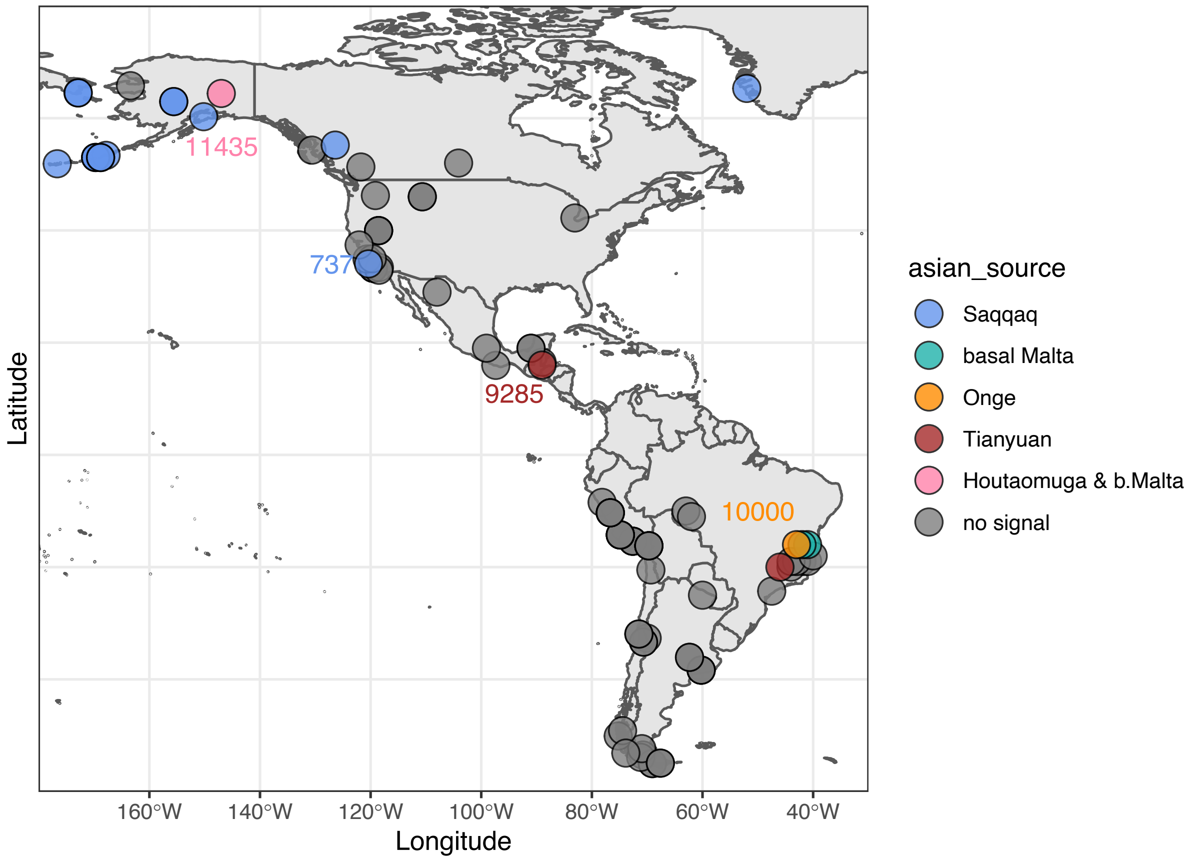


A map of extra-American ancestry sources detected at high confidence in ancient individuals and present-day groups. Individuals/groups showing no signal are labelled in grey. Presence/absence of selected ancestry sources is color-coded according to the legend. Average radiocarbon dates in cal yr BP are plotted next to selected circles.

**Fig. S48.**

Z-score = 4.8, 18 pops., 19 adm. events, 17 unlocked, 349,540 sites used, 1 winning among 105 topol.

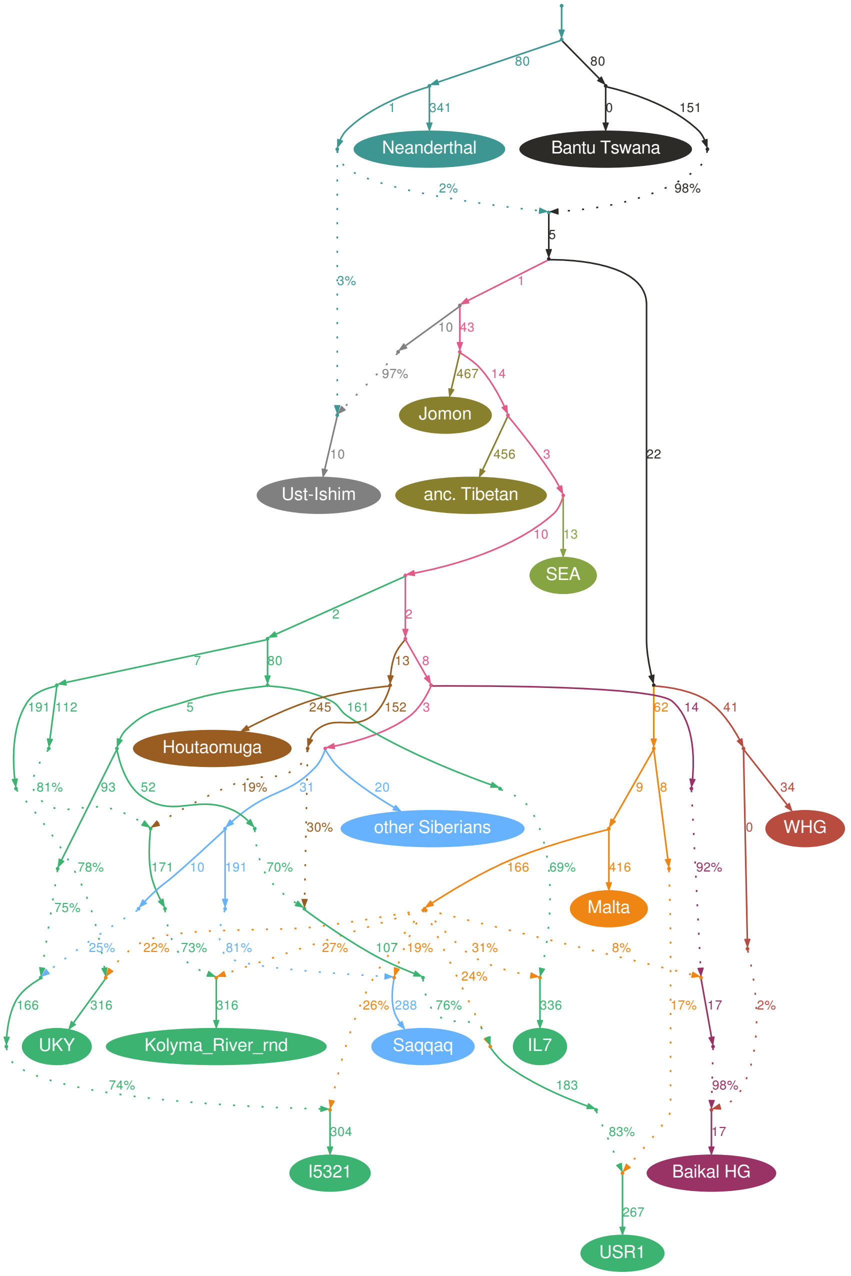


The best topology of the proto-American clade ((Ust'-Kyakhta [unadmixed], Kolyma [admixed]), (SNA, (USR1 [admixed], NNA))) selected among 105 alternative topologies tested under one setting/dataset combination. The following settings were used for the graph shown here: 1) the West Eurasian gene flows into Siberians were not locked; 2) external gene flows into the four American individuals were not locked; 3) transitions were kept; 4) "useallsnps: NO"; 5) the Houtaomuga group was composed of three individuals. The Nganasan and Selkup lineages are collapsed under the "other Siberians" label for making the visualization simpler.

**Fig. S49.**

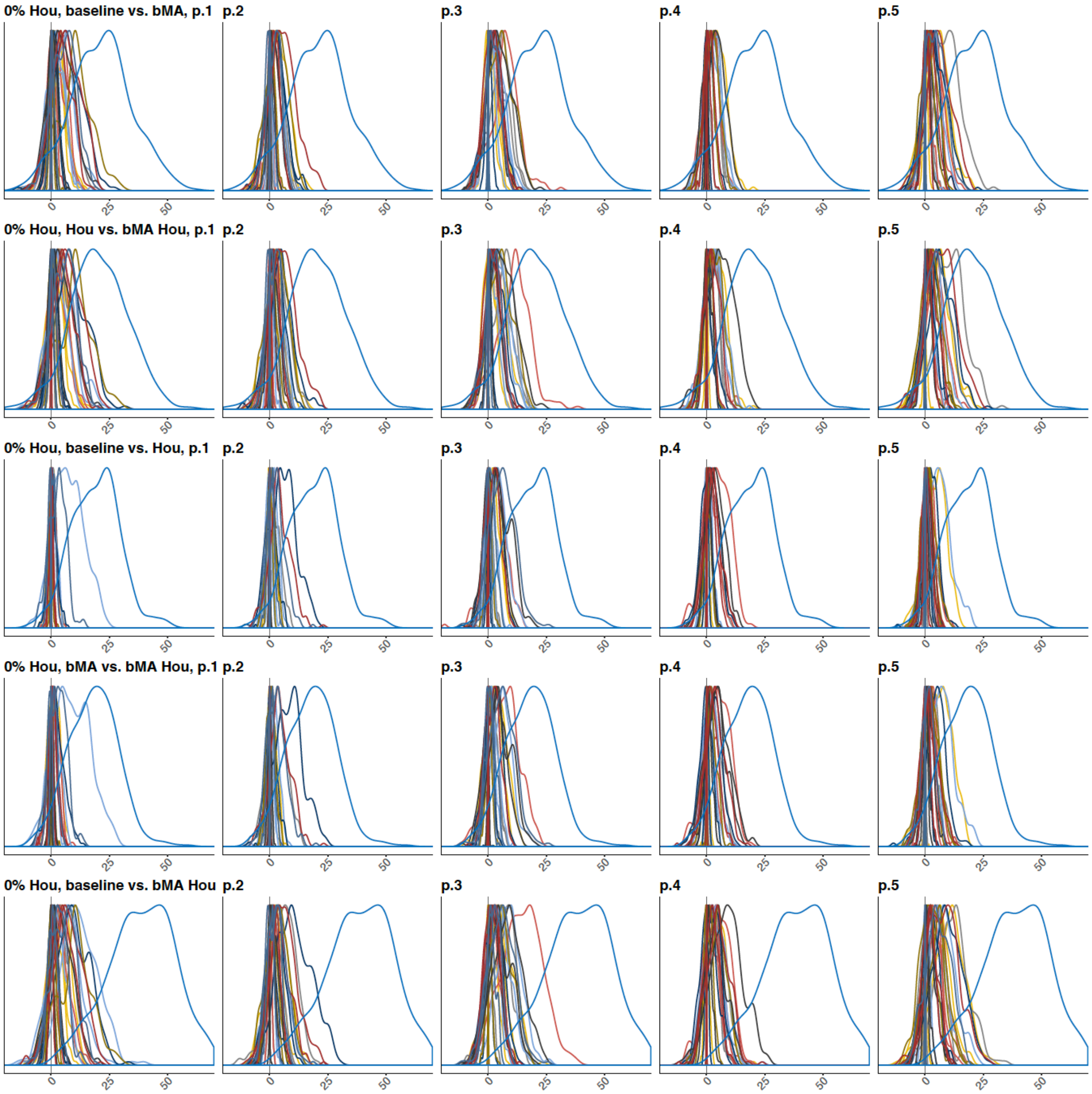


Distributions of out-of-sample LL differences across bootstrap replicates on data simulated under the null hypothesis and on real data. Five model comparisons were performed, see the labels in the leftmost panels. Results for 130 simulated datasets (200 bootstrap replicates each) are shown dataset by dataset, and the curves are shown across five panels and colored to aid visualization. The rightmost "dark-blue" curves in each panel correspond to the standard group composition on real data (based on 500 bootstrap replicates): the composite FAM group, three Houtaomuga individuals. Abbreviations used: bMA, basal Mal'ta; Hou, Houtaomuga.

**Fig. S50.**

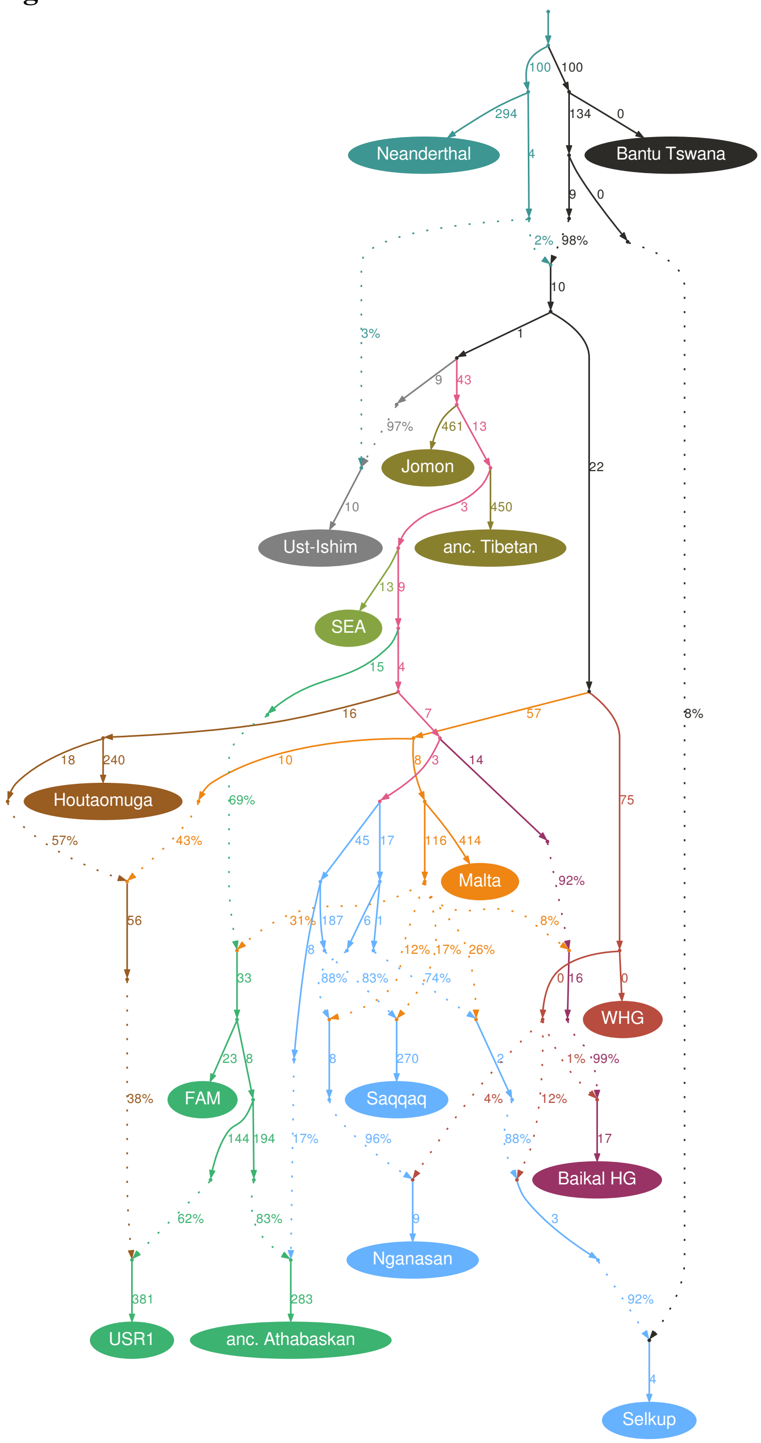


The figure shows a 4-way admixture model for USR1 in the context of the American clade including the FAM group (13 present-day individuals) and an ancient Athabaskan individual I5321. The model shown was constructed on the full 1240K-based dataset and features a "basal" Mal'ta + Houtaomuga admixed group that merges with the USR1 lineage. The model had a WR of 4.8 SE on the full dataset and 3.51 on the dataset without transitions. 450,669 sites were used on the full dataset, and 84,780 on the dataset without transitions.

**Fig. S51.**

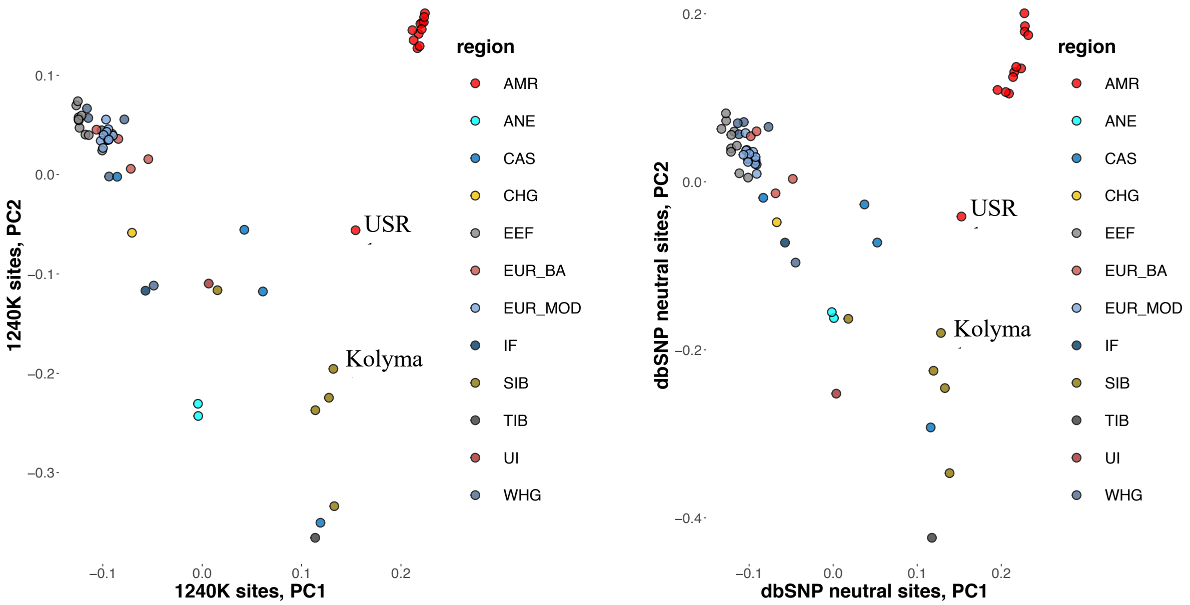


Comparison of PC1 vs. PC2 plots for the 1240K dataset and the neutral dataset. The following abbreviations are used for large regional/genetic groups: AMR, Native Americans; ANE, ancient North Eurasians (Yana); CAS, Central Asians; CHG, Caucasian hunter-gatherers; EEF, early European (and Anatolian) farmers; EUR BA, Bronze Age West Eurasia; EUR MOD, Iron Age and Medieval West Eurasia; IF, Iranian Neolithic; SIB; Siberians; TIB, Tibetans; UI, Ust'-Ishim; WHG, West European hunter-gatherers.

**Fig. S52.**

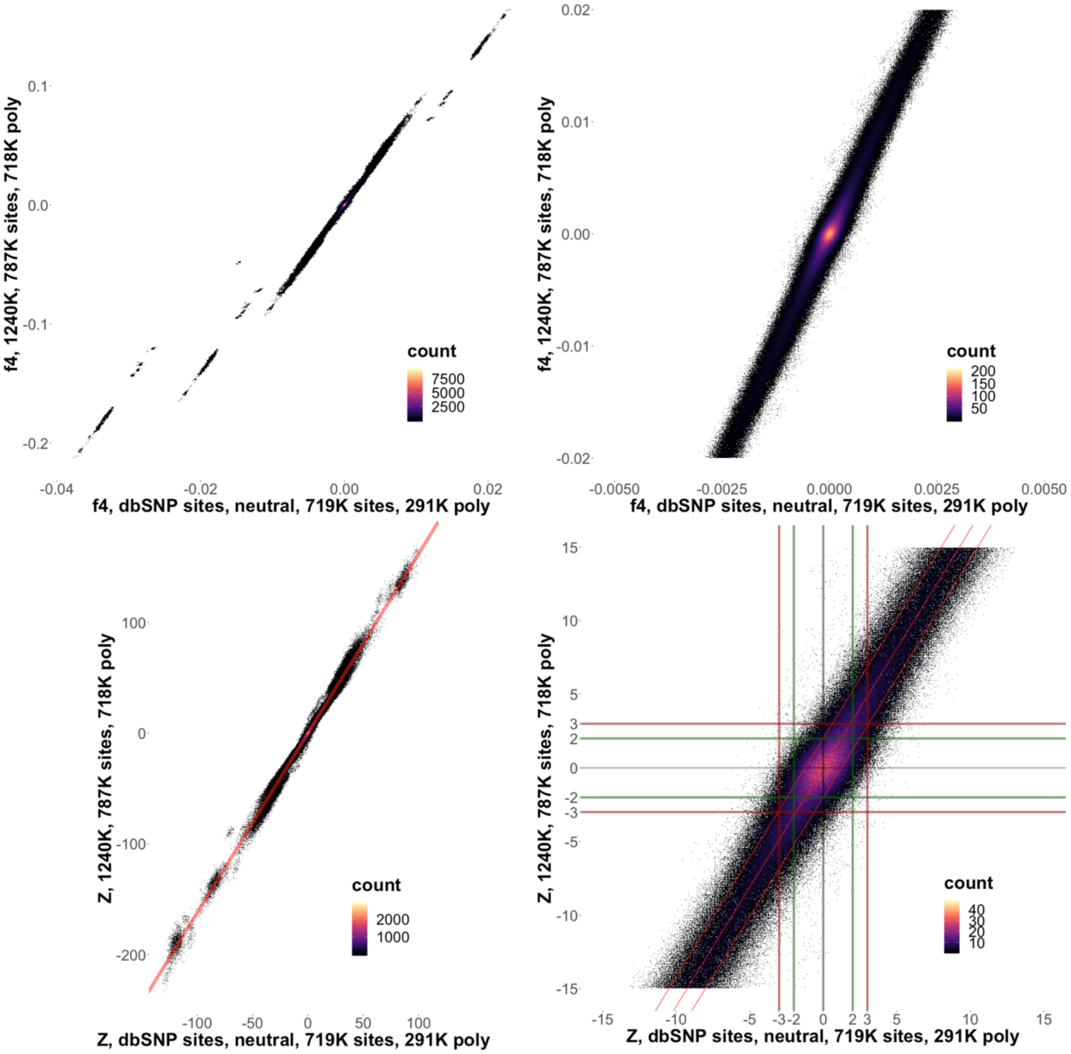


Heatmaps illustrating the distribution of $f_4$-statistics and their Z-scores in the coordinates of statistics on the neutral dataset (x-axis) and on the 1240K dataset (y-axis). Red diagonal lines on the lower right plot show the expected diagonal ± 2 SE. The numbers of sites used for calculating the statistics are shown on the axes. Please pay attention to the distant outliers scattered in the vertical strip between -3 and 3 SE on the x-axis.

**Fig. S53.**

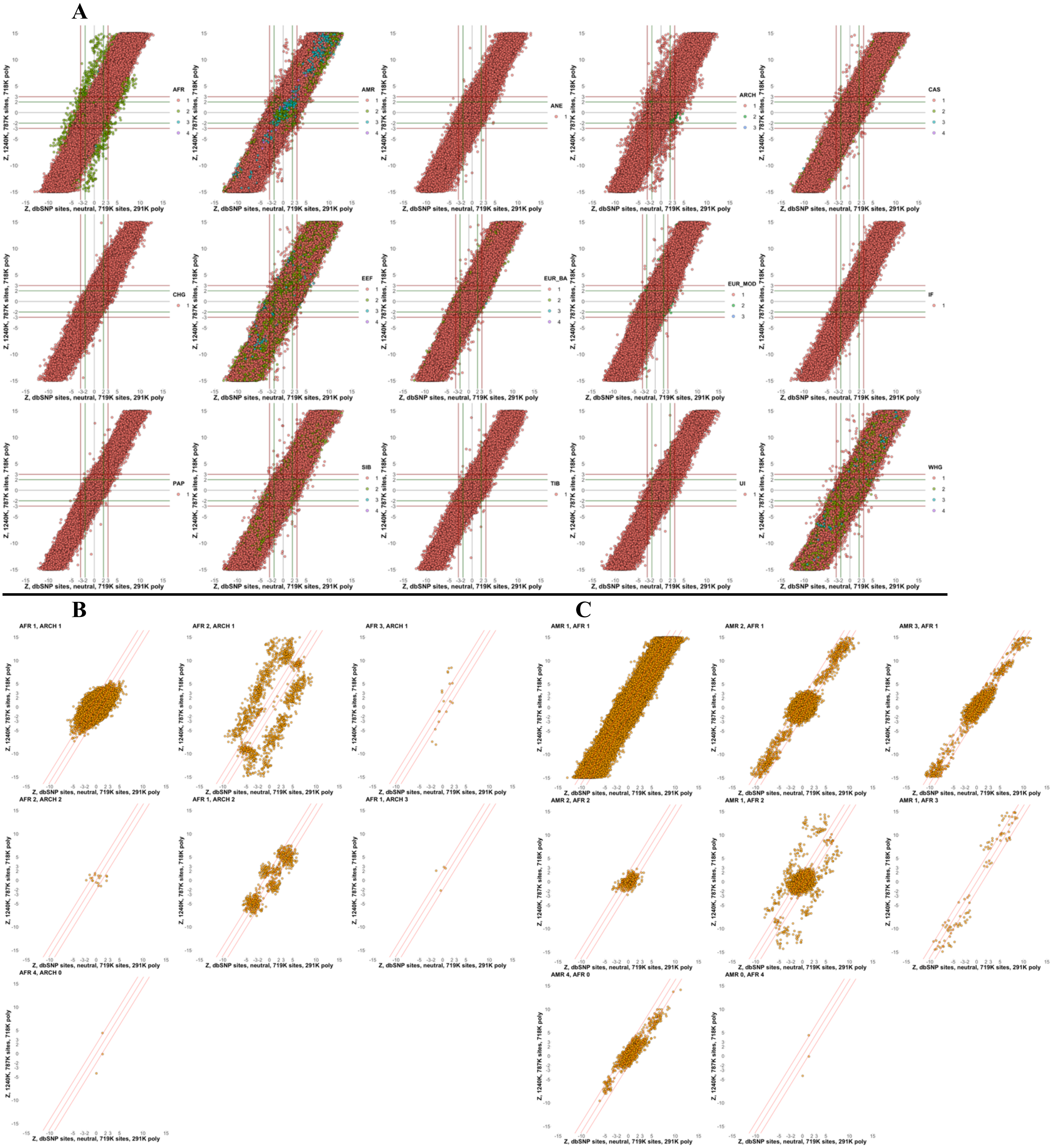
A
B
C
Z, dbSNP sites, neutral, 719K sites, 291K poly
Z, 1240K, 787K sites, 718K poly
AFR
AMR
ANE
ARCH
CAS
CHG
EEF
EUR_BA
EUR_MOD
IF
PAP
SIB
TIB
UI
WHG
AFR 1, ARCH 1
AFR 2, ARCH 1
AFR 3, ARCH 1
AFR 2, ARCH 2
AFR 1, ARCH 2
AFR 1, ARCH 3
AFR 4, ARCH 0
AMR 1, AFR 1
AMR 2, AFR 1
AMR 3, AFR 1
AMR 2, AFR 2
AMR 1, AFR 2
AMR 1, AFR 3
AMR 4, AFR 0
AMR 0, AFR 4

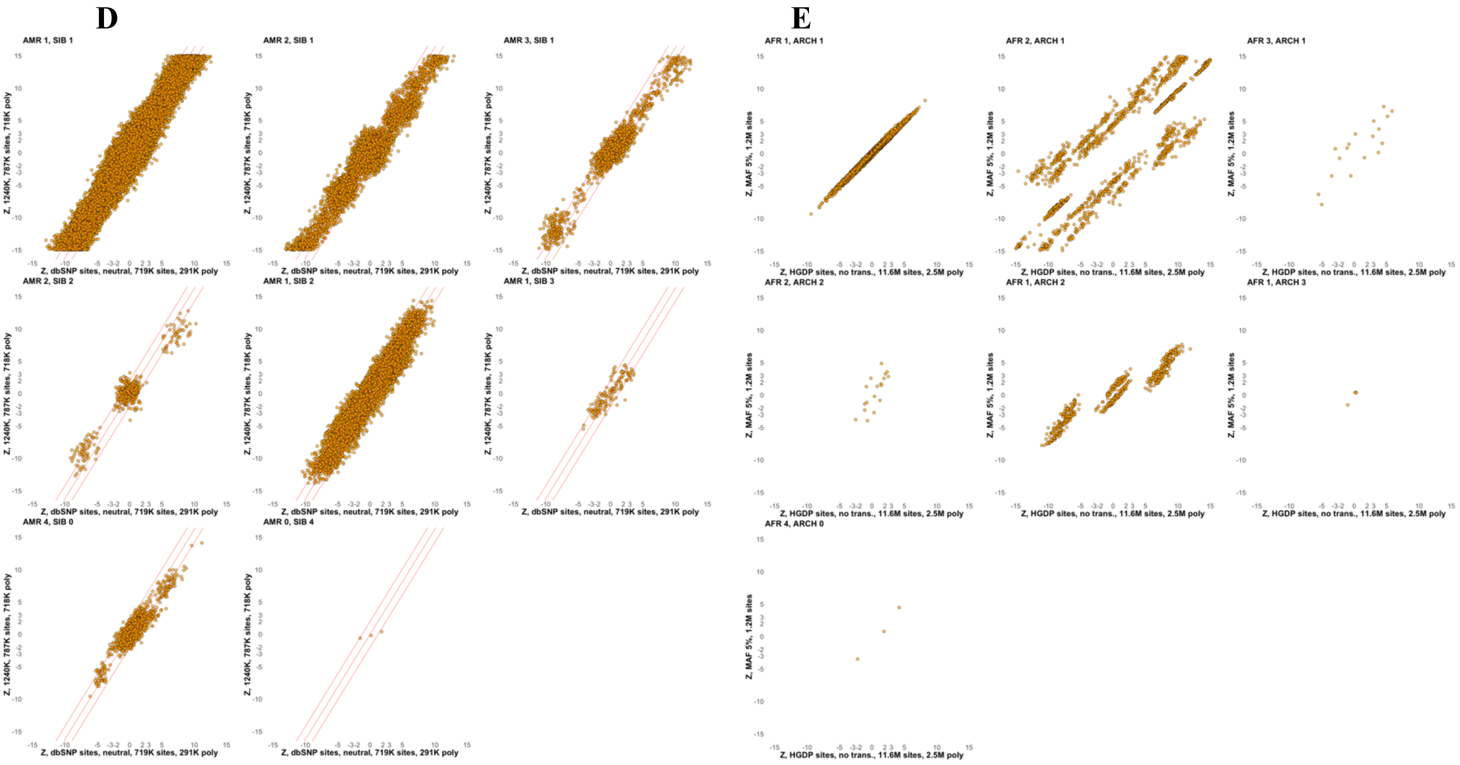


Biplots illustrating the distribution of $f_4$-statistic Z-scores across large regional/genetic groups. (**A**) The number of group members per statistic, which ranges from one to four, is color coded according to the legends. (**B**) Various types of statistics including from 1 to 4 African groups and from 1 to 3 archaic groups are shown. (**C**) Various types of statistics including from 1 to 4 African groups and from 1 to 4 American groups are shown. (**D**) Various types of statistics including from 1 to 4 Siberian groups and from 1 to 4 American groups are shown. (**E**) The worst biases (those affecting the statistics including two African group and an archaic individual) are reproduced by removing rare variants (i.e. by applying a global MAF filter of 5%). The numbers of sites used for calculating the statistics are shown on the axes. The following abbreviations are used: AFR, Africans; AMR, Native Americans; ANE, ancient North Eurasians (Yana); CAS, Central Asians; CHG, Caucasian hunter-gatherers; EEF, early European (and Anatolian) farmers; EUR BA, Bronze Age West Eurasia; EUR MOD, Iron Age and Medieval West Eurasia; IF, Iranian Neolithic; PAP, Papuans and related groups (Andamanese in our case); SIB; Siberians; TIB, Tibetans; UI, Ust'-Ishim; WHG, West European hunter-gatherers.

**Fig. S54.**

**A**

WR = 2.5 SE, LL = 35.8 units, 13 pops., 6 adm. events, 240,506 sites used

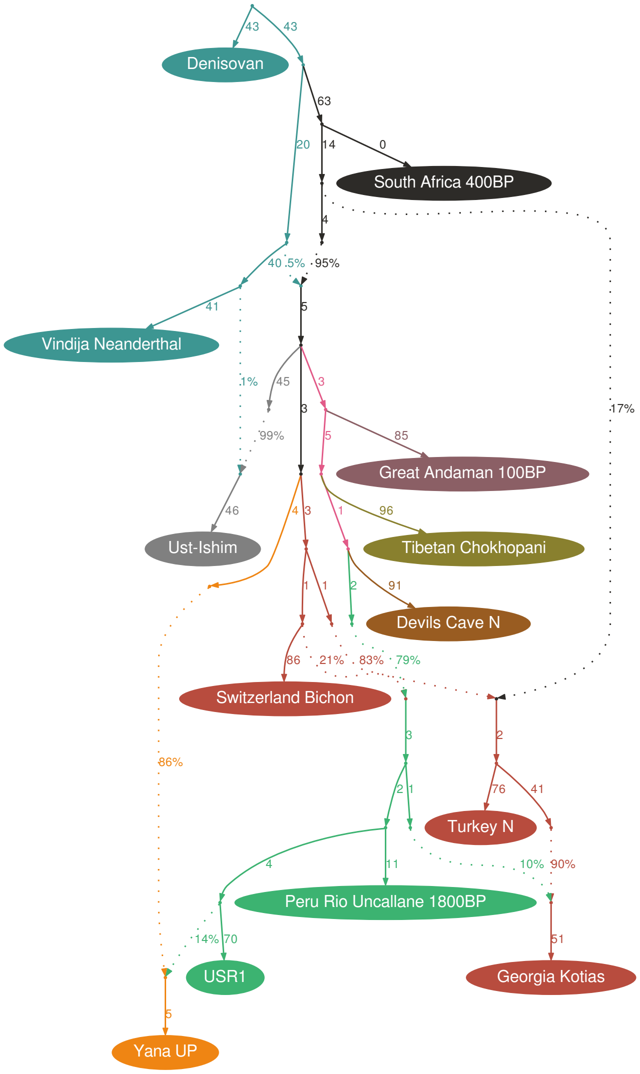


**B**

WR = 2.53 SE, LL = 35.4 units, 13 pops., 6 adm. events, 240,506 sites used

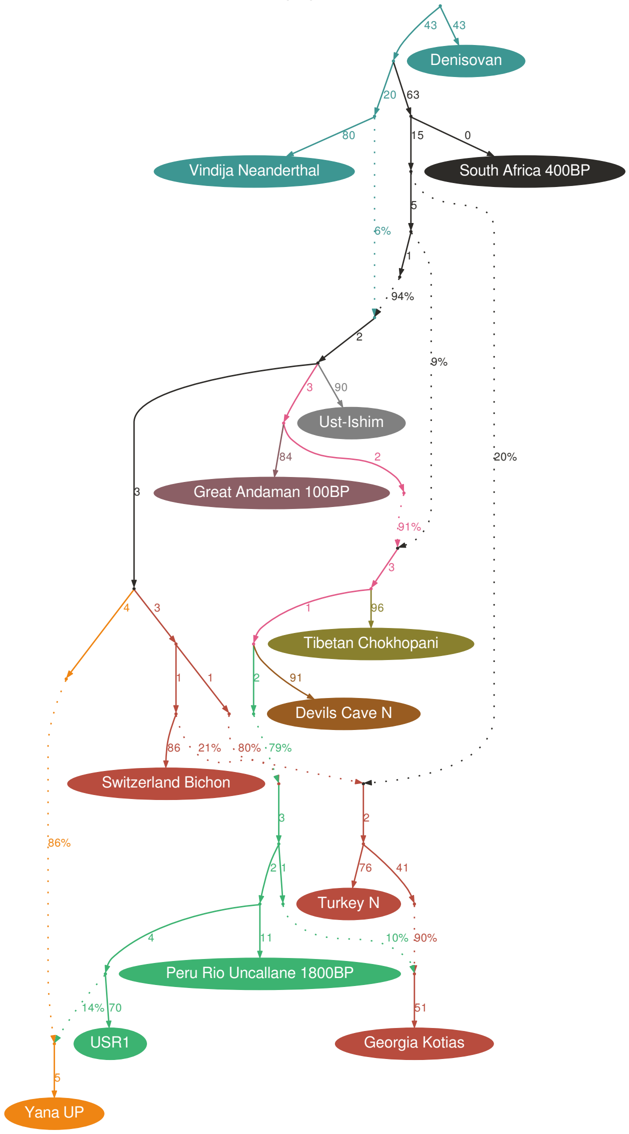

**C** WR = 2.15 SE, LL = 32.9 units, 13 pops., 6 adm. events, 240,506 sites used

**D** WR = 2.13 SE, LL = 33.8 units, 13 pops., 6 adm. events, 240,506 sites used

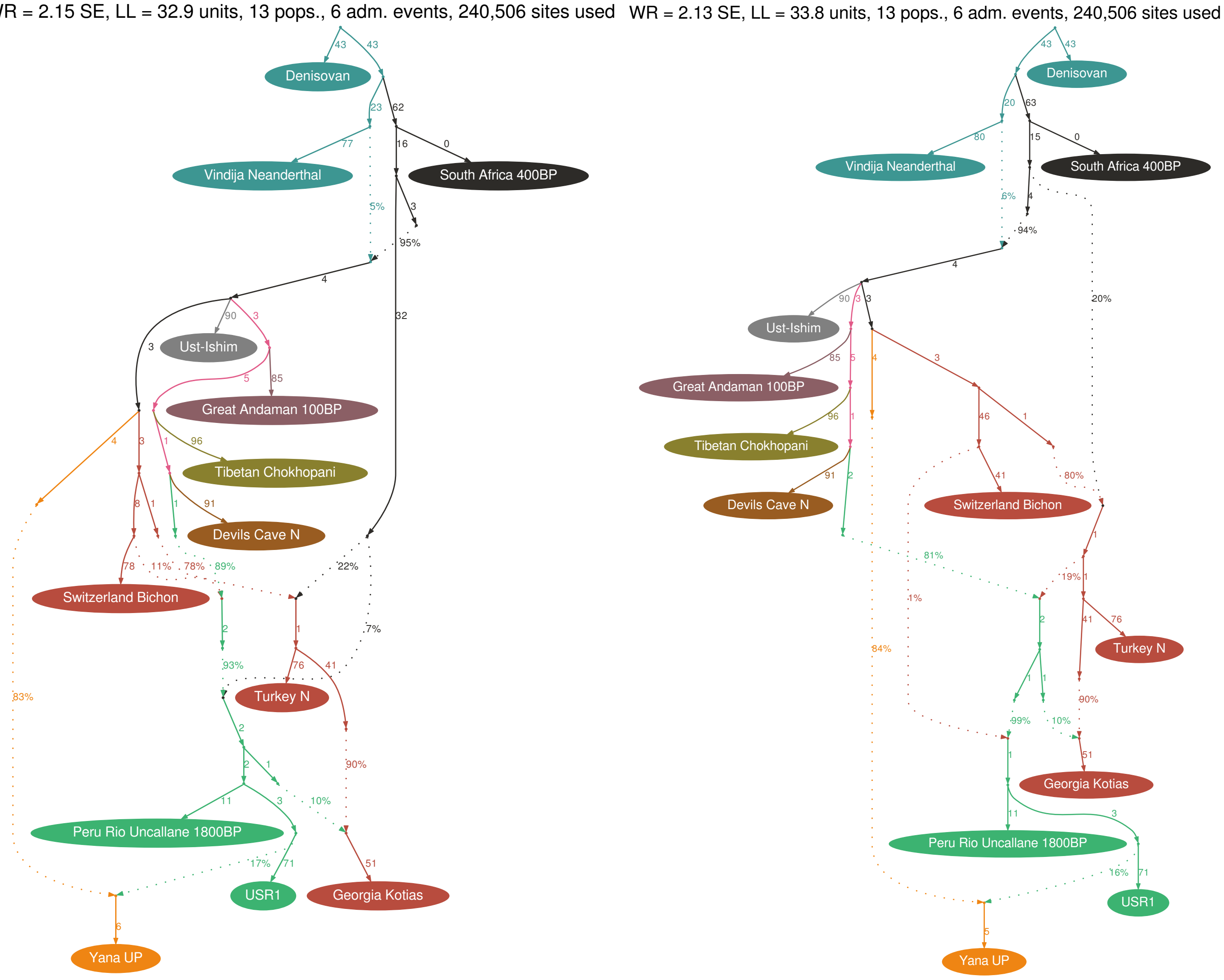


Best-fitting graphs generated by four automated topology exploration runs. Edges having the length of 0 were collapsed for simplicity.

**Fig. S55.**

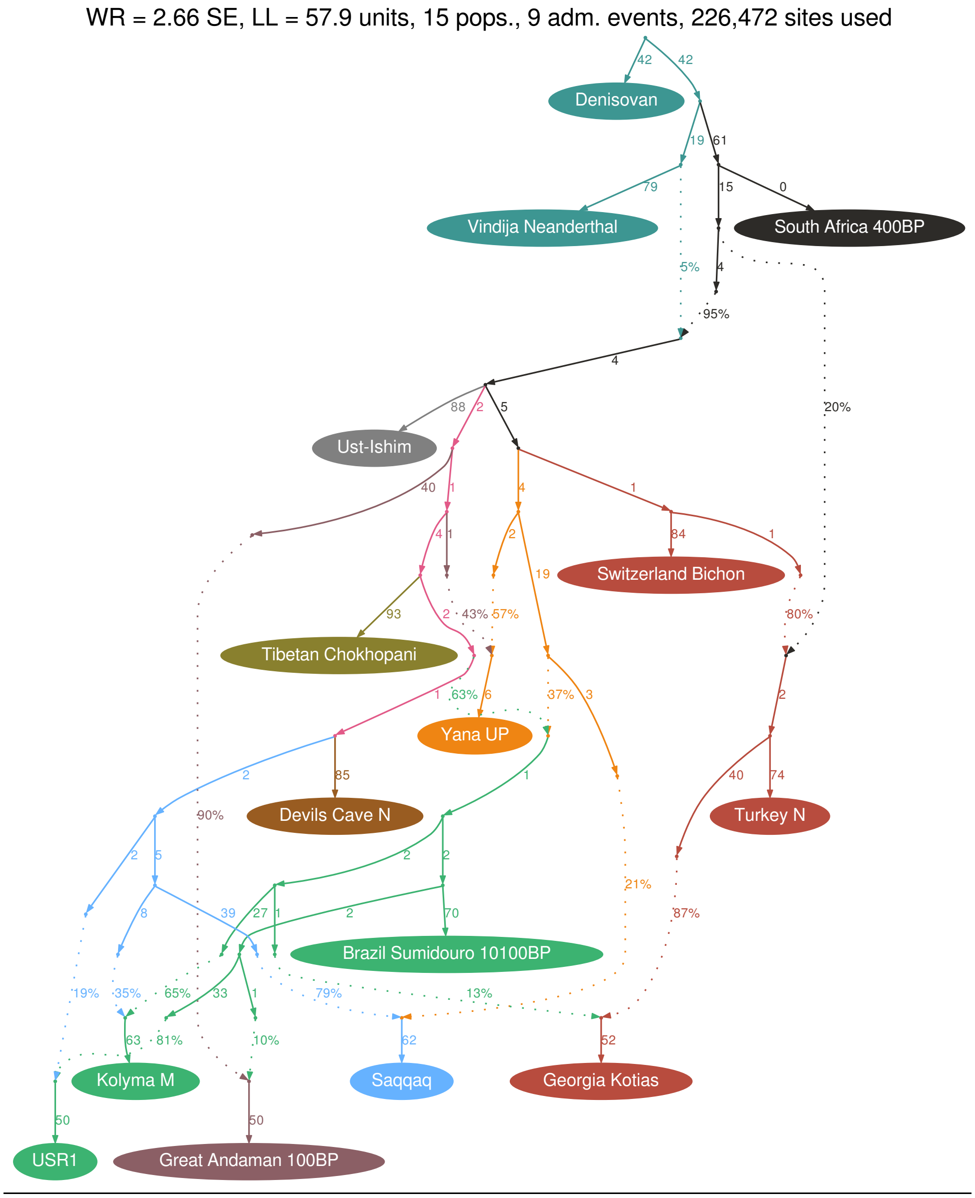


An outcome of manual exploration of more complex graph models incorporating 15 groups and up to 10 admixture events. The model shown incorporates 9 admixture events. Edges having the length of 0 were collapsed for simplicity.

#### Table S1.

| | | | | all models considered, 95K | | | all models considered, 400K | | |
|---|---|---|---|---|---|---|---|---|---|
| Tdiv2 | Tdiv1 | Tadm | classifier | min % | max % | interval width, % | min % | max % | interval width, % |
| 10000 | 8500 | 2500 | LL10 | 20 | 70 | 50 | 10 | 70 | 60 |
| | | | SR20 | 20 | 40 | 20 | 10 | 60 | 50 |
| | | | SSR40 | 20 | 60 | 40 | 10 | 60 | 50 |
| | | | WR08 | 20 | 50 | 30 | 10 | 70 | 60 |
| | | | int. map. | NA | NA | 0 | NA | NA | 0 |
| | | 5000 | LL10 | 20 | 60 | 40 | 20 | 70 | 50 |
| | | | SR20 | 30 | 50 | 20 | 20 | 60 | 40 |
| | | | SSR40 | 30 | 60 | 30 | 20 | 60 | 40 |
| | | | WR08 | 30 | 60 | 30 | 20 | 60 | 40 |
| | | | int. map. | 30 | 30 | 5 | 30 | 30 | 5 |
| | | 7500 | LL10 | 40 | 60 | 20 | 40 | 60 | 20 |
| | | | SR20 | NA | NA | 0 | 40 | 60 | 20 |
| | | | SSR40 | NA | NA | 0 | 40 | 60 | 20 |
| | | | WR08 | NA | NA | 0 | 50 | 50 | 0 |
| | | | int. map. | 40 | 70 | 30 | 50 | 60 | 10 |
| 13000 | 10500 | 2500 | LL10 | 20 | 80 | 60 | 10 | 80 | 70 |
| | | | SR20 | 20 | 70 | 50 | 10 | 70 | 60 |
| | | | SSR40 | 20 | 70 | 50 | 10 | 70 | 60 |
| | | | WR08 | 20 | 70 | 50 | 10 | 80 | 70 |
| | | | int. map. | NA | NA | 0 | NA | NA | 0 |
| | | 5000 | LL10 | 20 | 70 | 50 | 10 | 80 | 70 |
| | | | SR20 | 20 | 70 | 50 | 20 | 70 | 50 |
| | | | SSR40 | 20 | 70 | 50 | 10 | 70 | 60 |
| | | | WR08 | 20 | 70 | 50 | 20 | 70 | 50 |
| | | | int. map. | 30 | 30 | 5 | NA | NA | 0 |
| | | 7500 | LL10 | 30 | 70 | 40 | 20 | 80 | 60 |
| | | | SR20 | 30 | 60 | 30 | 20 | 70 | 50 |
| | | | SSR40 | 30 | 60 | 30 | 20 | 70 | 50 |
| | | | WR08 | 30 | 60 | 30 | 20 | 70 | 50 |
| | | | int. map. | 40 | 40 | 5 | 40 | 40 | 5 |
| 13000 | 8500 | 2500 | LL10 | 10 | 80 | 70 | 10 | 80 | 70 |
| | | | SR20 | 20 | 80 | 60 | 10 | 80 | 70 |
| | | | SSR40 | 20 | 80 | 60 | 10 | 80 | 70 |
| | | | WR08 | 20 | 80 | 60 | 10 | 80 | 70 |
| | | | int. map. | NA | NA | 0 | NA | NA | 0 |
| | | 5000 | LL10 | 20 | 80 | 60 | 10 | 80 | 70 |
| | | | SR20 | 20 | 80 | 60 | 10 | 80 | 70 |
| | | | SSR40 | 20 | 80 | 60 | 10 | 80 | 70 |
| | | | WR08 | 20 | 80 | 60 | 10 | 80 | 70 |
| | | | int. map. | NA | NA | 0 | NA | NA | 0 |
| | | 7500 | LL10 | 20 | 80 | 60 | 20 | 80 | 60 |
| | | | SR20 | 30 | 70 | 40 | 20 | 80 | 60 |
| | | | SSR40 | 20 | 70 | 50 | 20 | 80 | 60 |
| | | | WR08 | 30 | 70 | 40 | 30 | 80 | 50 |
| | | | int. map. | NA | NA | 0 | NA | NA | 0 |
| 15000 | 8500 | 2500 | LL10 | 10 | 90 | 80 | 10 | 95 | 85 |
| | | | SR20 | 20 | 80 | 60 | 10 | 90 | 80 |
| | | | SSR40 | 20 | 80 | 60 | 10 | 90 | 80 |
| | | | WR08 | 20 | 90 | 70 | 10 | 90 | 80 |
| | | | int. map. | 40 | 60 | 20 | 50 | 50 | 5 |
| | | 5000 | LL10 | 10 | 90 | 80 | 10 | 90 | 80 |
| | | | SR20 | 20 | 90 | 70 | 10 | 90 | 80 |
| | | | SSR40 | 20 | 90 | 70 | 10 | 90 | 80 |
| | | | WR08 | 20 | 90 | 70 | 10 | 90 | 80 |
| | | | int. map. | 50 | 70 | 20 | 50 | 70 | 20 |
| | | 7500 | LL10 | 20 | 90 | 70 | 20 | 90 | 70 |
| | | | SR20 | 20 | 80 | 60 | 20 | 90 | 70 |
| | | | SSR40 | 20 | 80 | 60 | 20 | 90 | 70 |
| | | | WR08 | 20 | 80 | 60 | 20 | 90 | 70 |
| | | | int. map. | 50 | 80 | 30 | 50 | 80 | 30 |
| 15000 | 8500 | 2500 | LL10 | 10 | 90 | 80 | 5 | 95 | 90 |
| | | | SR20 | 10 | 90 | 80 | 10 | 90 | 80 |
| | | | SSR40 | 10 | 90 | 80 | 5 | 90 | 85 |
| | | | WR08 | 10 | 90 | 80 | 10 | 90 | 80 |
| | | | int. map. | 30 | 50 | 20 | 40 | 50 | 10 |
| | | 5000 | LL10 | 10 | 90 | 80 | 5 | 90 | 85 |
| | | | SR20 | 10 | 90 | 80 | 5 | 90 | 85 |
| | | | SSR40 | 10 | 90 | 80 | 5 | 90 | 85 |
| | | | WR08 | 10 | 90 | 80 | 10 | 90 | 80 |
| | | | int. map. | 30 | 60 | 30 | 30 | 60 | 30 |
| | | 7500 | LL10 | 10 | 90 | 80 | 10 | 90 | 80 |
| | | | SR20 | 10 | 90 | 80 | 10 | 90 | 80 |
| | | | SSR40 | 10 | 90 | 80 | 10 | 90 | 80 |
| | | | WR08 | 10 | 90 | 80 | 10 | 90 | 80 |
| | | | int. map. | 30 | 80 | 50 | 30 | 80 | 50 |
| 20000 | 8500 | 2500 | LL10 | 5 | 95 | 90 | 5 | 95 | 90 |
| | | | SR20 | 5 | 95 | 90 | 5 | 95 | 90 |
| | | | SSR40 | 5 | 95 | 90 | 5 | 95 | 90 |
| | | | WR08 | 5 | 95 | 90 | 5 | 95 | 90 |
| | | | int. map. | 20 | 60 | 40 | 30 | 50 | 20 |
| | | 5000 | LL10 | 5 | 95 | 90 | 5 | 95 | 90 |
| | | | SR20 | 5 | 95 | 90 | 5 | 95 | 90 |
| | | | SSR40 | 5 | 95 | 90 | 5 | 95 | 90 |
| | | | WR08 | 5 | 95 | 90 | 5 | 95 | 90 |
| | | | int. map. | 20 | 70 | 50 | 20 | 60 | 40 |
| | | 7500 | LL10 | 5 | 95 | 90 | 5 | 95 | 90 |
| | | | SR20 | 5 | 95 | 90 | 5 | 95 | 90 |
| | | | SSR40 | 5 | 95 | 90 | 5 | 95 | 90 |
| | | | WR08 | 10 | 95 | 85 | 5 | 95 | 90 |
| | | | int. map. | 20 | 90 | 70 | 20 | 90 | 70 |

Admixture proportions confidently detectable (0% false negative iterations) by the mapping approach with four alternative classifiers under various setting combinations. Minimal and maximal confidently detectable proportions and their difference (the length of the confidently detectable interval) are shown in percent. Differences among classifiers, settings, and datasets (95,000 or 400,000 sites, "information-rich" and "information-poor") are noticeable. Results for the information-rich graphs are marked in grey shading. We also showed the intervals of simulated admixture proportions which resulted in a non-zero fraction of best one-way modes demonstrating intermediate positions (not on the I or G branches). Those intervals are shown in green if they lie inside the confidently detectable interval, and in red otherwise.

#### Table S2.

| | | | Tadm 2500 | | | | | | Tadm 5000 | | | | | | Tadm 7500 | | | | | |
|---|---|---|---|---|---|---|---|---|---|---|---|---|---|---|---|---|---|---|---|---|
| | | | 95K | | | | 400K | | 95K | | | | 400K | | 95K | | | | 400K | |
| Tdiv2 | Tdiv1 | simulated gene flow % | LL diff. incorr.1-p.corr.1 >3 | WR diff. incorr.1-p.corr.1 >0.1 | LL diff. incorr.2-p.corr.2 >3 | WR diff. incorr.2-p.corr.2 >0.1 | LL diff. incorr.1-p.corr.1 >3 | WR diff. incorr.1-p.corr.1 >0.1 | LL diff. incorr.1-p.corr.1 >3 | WR diff. incorr.1-p.corr.1 >0.1 | LL diff. incorr.2-p.corr.2 >3 | WR diff. incorr.2-p.corr.2 >0.1 | LL diff. incorr.1-p.corr.1 >3 | WR diff. incorr.1-p.corr.1 >0.1 | LL diff. incorr.1-p.corr.1 >3 | WR diff. incorr.1-p.corr.1 >0.1 | LL diff. incorr.2-p.corr.2 >3 | WR diff. incorr.2-p.corr.2 >0.1 | LL diff. incorr.1-p.corr.1 >3 | WR diff. incorr.1-p.corr.1 >0.1 |
| 10000 | 8500 | 0 | 0 | 0 | 0 | 0 | 0 | 0 | 0 | 0 | 0 | 0 | 0 | 0 | 0 | 0 | 0 | 0 | 0 | 0 |
| | | 5 | 6 | 1 | 10 | 0 | 4 | 3 | 3 | 1 | 5 | 0 | 3 | 5 | 2 | 1 | 5 | 0 | 4 | 3 |
| | | 10 | 13 | 3 | 15 | 0 | 16 | 9 | 12 | 8 | 18 | 0 | 15 | 4 | 8 | 6 | 11 | 0 | 11 | 8 |
| | | 20 | 23 | 8 | 27 | 0 | 36 | 17 | 16 | 11 | 17 | 0 | 27 | 9 | 16 | 11 | 20 | 0 | 23 | 7 |
| | | 30 | 43 | 25 | 36 | 0 | 47 | 37 | 30 | 14 | 29 | 0 | 44 | 22 | 23 | 10 | 20 | 0 | 33 | 20 |
| | | 40 | 47 | 35 | 42 | 0 | 46 | 41 | 35 | 25 | 36 | 0 | 45 | 32 | 38 | 17 | 36 | 0 | 44 | 33 |
| | | 50 | 48 | 38 | 43 | 0 | 50 | 47 | 50 | 38 | 47 | 0 | 50 | 46 | 35 | 23 | 35 | 0 | 48 | 35 |
| | | 60 | 48 | 38 | 47 | 0 | 50 | 50 | 48 | 39 | 46 | 0 | 50 | 48 | 42 | 39 | 44 | 0 | 50 | 46 |
| | | 70 | 46 | 43 | 46 | 0 | 50 | 50 | 48 | 38 | 42 | 0 | 50 | 49 | 46 | 39 | 45 | 0 | 50 | 49 |
| | | 80 | 50 | 46 | 49 | 0 | 50 | 49 | 49 | 47 | 48 | 0 | 50 | 49 | 49 | 42 | 47 | 0 | 50 | 50 |
| | | 90 | 50 | 48 | 50 | 0 | 50 | 50 | 49 | 44 | 48 | 0 | 50 | 50 | 50 | 47 | 49 | 0 | 50 | 50 |
| | | 95 | 50 | 46 | 50 | 0 | 50 | 50 | 50 | 50 | 50 | 0 | 50 | 50 | 50 | 49 | 49 | 0 | 50 | 50 |
| | | 100 | 50 | 50 | 50 | 0 | 50 | 50 | 50 | 50 | 50 | 1 | 50 | 50 | 50 | 50 | 49 | 1 | 50 | 50 |
| 13000 | 10500 | 0 | 0 | 0 | 0 | 0 | 0 | 0 | 0 | 0 | 0 | 0 | 0 | 0 | 0 | 0 | 0 | 0 | 0 | 0 |
| | | 5 | 7 | 5 | 8 | 0 | 12 | 6 | 6 | 6 | 8 | 0 | 10 | 5 | 2 | 3 | 7 | 0 | 7 | 8 |
| | | 10 | 15 | 3 | 15 | 0 | 27 | 15 | 13 | 3 | 13 | 0 | 16 | 6 | 11 | 5 | 16 | 0 | 18 | 10 |
| | | 20 | 42 | 31 | 40 | 0 | 47 | 41 | 31 | 14 | 28 | 0 | 41 | 28 | 32 | 17 | 29 | 0 | 42 | 27 |
| | | 30 | 50 | 46 | 49 | 1 | 50 | 49 | 48 | 39 | 46 | 0 | 49 | 47 | 42 | 23 | 37 | 0 | 48 | 38 |
| | | 40 | 50 | 50 | 50 | 0 | 50 | 50 | 48 | 42 | 47 | 0 | 50 | 49 | 47 | 35 | 46 | 0 | 49 | 47 |
| | | 50 | 50 | 50 | 50 | 0 | 50 | 50 | 50 | 49 | 50 | 0 | 50 | 50 | 49 | 45 | 49 | 0 | 50 | 49 |
| | | 60 | 50 | 50 | 50 | 1 | 50 | 50 | 50 | 49 | 50 | 0 | 50 | 50 | 50 | 49 | 50 | 0 | 50 | 50 |
| | | 70 | 50 | 50 | 50 | 0 | 50 | 50 | 50 | 49 | 50 | 1 | 50 | 50 | 50 | 48 | 50 | 1 | 50 | 50 |
| | | 80 | 50 | 50 | 50 | 1 | 50 | 50 | 50 | 50 | 50 | 0 | 50 | 50 | 50 | 50 | 50 | 2 | 50 | 50 |
| | | 90 | 50 | 50 | 50 | 1 | 50 | 50 | 50 | 49 | 50 | 2 | 50 | 50 | 50 | 50 | 50 | 0 | 50 | 50 |
| | | 95 | 50 | 50 | 50 | 1 | 50 | 50 | 50 | 50 | 50 | 1 | 50 | 50 | 50 | 50 | 50 | 3 | 50 | 50 |
| | | 100 | 50 | 50 | 50 | 4 | 50 | 50 | 50 | 50 | 50 | 2 | 50 | 50 | 50 | 50 | 50 | 3 | 50 | 50 |
| 13000 | 8500 | 0 | 0 | 0 | 0 | 0 | 0 | 0 | 0 | 0 | 0 | 0 | 0 | 0 | 0 | 0 | 0 | 0 | 0 | 0 |
| | | 5 | 19 | 10 | 22 | 0 | 22 | 5 | 14 | 5 | 16 | 0 | 16 | 9 | 11 | 5 | 16 | 0 | 9 | 6 |
| | | 10 | 35 | 20 | 31 | 0 | 43 | 29 | 32 | 15 | 35 | 0 | 38 | 25 | 22 | 13 | 21 | 0 | 30 | 16 |
| | | 20 | 50 | 49 | 49 | 0 | 50 | 50 | 47 | 32 | 44 | 0 | 49 | 47 | 41 | 17 | 39 | 0 | 45 | 35 |
| | | 30 | 50 | 50 | 50 | 1 | 50 | 50 | 50 | 49 | 50 | 0 | 50 | 50 | 45 | 37 | 43 | 0 | 49 | 46 |
| | | 40 | 50 | 50 | 50 | 0 | 50 | 50 | 50 | 50 | 50 | 0 | 50 | 50 | 50 | 46 | 50 | 0 | 50 | 50 |
| | | 50 | 50 | 50 | 50 | 1 | 50 | 50 | 50 | 50 | 50 | 3 | 50 | 50 | 49 | 48 | 49 | 0 | 50 | 50 |
| | | 60 | 50 | 50 | 50 | 8 | 50 | 50 | 50 | 50 | 50 | 0 | 50 | 50 | 50 | 50 | 50 | 0 | 50 | 50 |
| | | 70 | 50 | 50 | 50 | 3 | 50 | 50 | 50 | 50 | 50 | 0 | 50 | 50 | 50 | 49 | 50 | 0 | 50 | 50 |
| | | 80 | 50 | 50 | 50 | 3 | 50 | 50 | 50 | 50 | 50 | 2 | 50 | 50 | 50 | 49 | 50 | 1 | 50 | 50 |
| | | 90 | 50 | 50 | 50 | 0 | 50 | 50 | 50 | 50 | 50 | 2 | 50 | 50 | 50 | 50 | 50 | 1 | 50 | 50 |
| | | 95 | 50 | 50 | 50 | 1 | 50 | 50 | 50 | 50 | 50 | 2 | 50 | 50 | 50 | 50 | 50 | 1 | 50 | 50 |
| | | 100 | 50 | 50 | 50 | 7 | 50 | 50 | 50 | 50 | 50 | 2 | 50 | 50 | 50 | 50 | 50 | 1 | 50 | 50 |
| 15000 | 8500 | 0 | 0 | 0 | 0 | 0 | 0 | 0 | 0 | 0 | 0 | 0 | 0 | 0 | 0 | 0 | 0 | 0 | 0 | 0 |
| | | 5 | 33 | 17 | 26 | 0 | 41 | 25 | 23 | 13 | 20 | 0 | 38 | 19 | 27 | 16 | 28 | 0 | 42 | 30 |
| | | 10 | 47 | 44 | 48 | 1 | 50 | 50 | 49 | 40 | 46 | 0 | 50 | 49 | 44 | 35 | 38 | 0 | 50 | 45 |
| | | 20 | 50 | 50 | 50 | 3 | 50 | 50 | 50 | 50 | 50 | 0 | 50 | 50 | 50 | 50 | 50 | 2 | 50 | 50 |
| | | 30 | 50 | 50 | 50 | 15 | 50 | 50 | 50 | 50 | 50 | 2 | 50 | 50 | 50 | 50 | 50 | 1 | 50 | 50 |
| | | 40 | 50 | 50 | 50 | 28 | 50 | 50 | 50 | 50 | 50 | 13 | 50 | 50 | 50 | 50 | 50 | 9 | 50 | 50 |
| | | 50 | 50 | 50 | 50 | 46 | 50 | 50 | 50 | 50 | 50 | 32 | 50 | 50 | 50 | 50 | 50 | 30 | 50 | 50 |
| | | 60 | 50 | 50 | 50 | 49 | 50 | 50 | 50 | 50 | 50 | 47 | 50 | 50 | 50 | 50 | 50 | 46 | 50 | 50 |
| | | 70 | 50 | 50 | 50 | 50 | 50 | 50 | 50 | 50 | 50 | 50 | 50 | 50 | 50 | 50 | 50 | 49 | 50 | 50 |
| | | 80 | 50 | 50 | 50 | 50 | 50 | 50 | 50 | 50 | 50 | 50 | 50 | 50 | 50 | 50 | 50 | 50 | 50 | 50 |
| | | 90 | 50 | 50 | 50 | 50 | 50 | 50 | 50 | 50 | 50 | 50 | 50 | 50 | 50 | 50 | 50 | 50 | 50 | 50 |
| | | 95 | 50 | 50 | 50 | 50 | 50 | 50 | 50 | 50 | 50 | 50 | 50 | 50 | 50 | 50 | 50 | 50 | 50 | 50 |
| | | 100 | 50 | 50 | 50 | 50 | 50 | 50 | 50 | 50 | 50 | 50 | 50 | 50 | 50 | 50 | 50 | 50 | 50 | 50 |

Admixture proportions that allow gene flow direction to be resolved by the LL3 and WR0.1 classifiers under the "mapping" setup, on sets of 95,000 or 400,000 sites. The following "information-poor" graph classes were considered: incorrect 1 and 2, partially correct 1 and 2. Settings that yielded good resolution (>95% iterations yielding a positive result) are highlighted in green.

### Table S3.

| Tdiv2 | Tdiv1 | simulated gene flow, % | qpGraph, 95K SD, Tadm 2500 | qpGraph, 95K SD, Tadm 5000 | qpGraph, 95K SD, Tadm 7500 | qpGraph, 400K SD, Tadm 2500 | qpGraph, 400K SD, Tadm 5000 | qpGraph, 400K SD, Tadm 7500 | qpAdm, 95K SD, Tadm 2500 | qpAdm, 95K SD, Tadm 5000 | qpAdm, 95K SD, Tadm 7500 | qpAdm, 400K SD, Tadm 2500 | qpAdm, 400K SD, Tadm 5000 | qpAdm, 400K SD, Tadm 7500 |
|---|---|---|---|---|---|---|---|---|---|---|---|---|---|---|
| 10000 | 8500 | 0 | 9.4 | 10.8 | 8.2 | 6.7 | 7.7 | 6.7 | 40.5 | 34.3 | 23.3 | 19.3 | 16.6 | 15.0 |
| | | 5 | 12.0 | 12.5 | 10.3 | 8.7 | 8.9 | 8.5 | 51.3 | 58.0 | 24.9 | 21.9 | 15.4 | 19.3 |
| | | 10 | 13.8 | 16.0 | 16.2 | 9.6 | 11.0 | 10.7 | 30.1 | 28.8 | 29.2 | 18.6 | 17.2 | 18.9 |
| | | 20 | 11.9 | 16.0 | 19.3 | 9.9 | 11.8 | 14.2 | 28.8 | 23.4 | 23.3 | 16.2 | 17.0 | 16.3 |
| | | 30 | 12.1 | 15.4 | 15.4 | 9.2 | 11.8 | 12.8 | 21.7 | 27.1 | 24.9 | 13.9 | 19.0 | 15.1 |
| | | 40 | 14.2 | 16.4 | 16.6 | 10.8 | 13.7 | 14.1 | 18.9 | 22.6 | 19.4 | 14.9 | 16.4 | 12.1 |
| | | 50 | 13.7 | 14.3 | 21.5 | 8.3 | 11.9 | 14.6 | 20.5 | 21.7 | 26.1 | 12.2 | 12.7 | 16.8 |
| | | 60 | 15.1 | 16.7 | 17.6 | 11.3 | 13.9 | 14.6 | 30.5 | 19.6 | 20.9 | 16.4 | 14.3 | 16.1 |
| | | 70 | 13.1 | 16.7 | 16.8 | 11.2 | 11.5 | 11.3 | 17.8 | 51.5 | 61.2 | 12.3 | 14.1 | 13.4 |
| | | 80 | 12.8 | 12.8 | 16.2 | 8.8 | 9.1 | 15.3 | 18.0 | 19.2 | 41.0 | 11.1 | 12.1 | 18.9 |
| | | 90 | 10.9 | 14.8 | 14.5 | 9.1 | 10.3 | 12.2 | 21.9 | 27.9 | 23.2 | 14.9 | 14.4 | 15.7 |
| | | 95 | 9.7 | 7.7 | 11.1 | 7.3 | 6.4 | 8.6 | 31.1 | 24.8 | 32.1 | 15.1 | 16.6 | 19.0 |
| | | 100 | 6.0 | 7.5 | 8.7 | 4.5 | 7.6 | 7.0 | 19.0 | 21.2 | 22.6 | 14.4 | 17.0 | 21.9 |
| 13000 | 10500 | 0 | 6.7 | 4.1 | 5.7 | 5.8 | 3.9 | 4.9 | 18.7 | 16.0 | 16.1 | 12.4 | 10.0 | 9.7 |
| | | 5 | 8.8 | 9.1 | 7.5 | 6.9 | 6.9 | 6.7 | 17.3 | 15.2 | 17.8 | 12.0 | 11.2 | 12.4 |
| | | 10 | 6.9 | 8.0 | 9.1 | 6.0 | 6.6 | 8.1 | 15.3 | 16.2 | 17.6 | 9.4 | 11.5 | 11.2 |
| | | 20 | 7.2 | 9.6 | 11.5 | 5.9 | 6.9 | 8.3 | 11.9 | 15.7 | 13.7 | 9.8 | 8.7 | 10.7 |
| | | 30 | 10.5 | 10.3 | 13.0 | 6.0 | 8.7 | 9.7 | 12.0 | 13.0 | 14.1 | 6.5 | 10.6 | 10.1 |
| | | 40 | 9.6 | 11.8 | 12.4 | 6.3 | 9.5 | 10.4 | 13.3 | 13.6 | 14.3 | 8.4 | 9.9 | 10.7 |
| | | 50 | 8.0 | 10.5 | 12.1 | 7.4 | 10.0 | 8.3 | 9.9 | 11.8 | 11.4 | 8.4 | 9.1 | 8.4 |
| | | 60 | 9.1 | 13.1 | 13.1 | 7.9 | 10.6 | 11.0 | 9.7 | 13.9 | 12.8 | 7.9 | 10.9 | 11.1 |
| | | 70 | 8.5 | 12.0 | 12.0 | 6.6 | 9.0 | 10.4 | 9.0 | 13.1 | 15.3 | 6.9 | 9.2 | 11.3 |
| | | 80 | 9.0 | 10.8 | 9.4 | 7.6 | 8.0 | 8.1 | 9.3 | 12.3 | 9.8 | 8.4 | 7.7 | 8.5 |
| | | 90 | 7.8 | 9.7 | 9.9 | 7.8 | 7.3 | 6.7 | 12.9 | 14.2 | 11.9 | 10.0 | 7.8 | 9.4 |
| | | 95 | 5.3 | 7.3 | 6.9 | 4.7 | 5.1 | 5.2 | 10.3 | 14.2 | 13.7 | 7.7 | 8.9 | 9.1 |
| | | 100 | 4.9 | 5.7 | 5.9 | 4.2 | 4.1 | 3.9 | 13.0 | 11.7 | 16.4 | 10.2 | 9.2 | 11.2 |
| 13000 | 8500 | 0 | 6.6 | 5.8 | 5.1 | 4.0 | 4.3 | 5.3 | 11.6 | 15.3 | 16.9 | 8.6 | 11.8 | 12.9 |
| | | 5 | 8.2 | 7.4 | 8.7 | 6.3 | 5.8 | 6.0 | 12.3 | 13.1 | 12.8 | 12.8 | 10.5 | 9.5 |
| | | 10 | 7.2 | 8.6 | 9.1 | 5.9 | 6.8 | 7.1 | 13.1 | 12.4 | 14.6 | 9.7 | 8.0 | 10.5 |
| | | 20 | 6.9 | 10.6 | 11.5 | 5.8 | 7.3 | 10.5 | 14.3 | 13.6 | 15.2 | 9.2 | 10.0 | 11.7 |
| | | 30 | 7.0 | 9.3 | 11.6 | 6.3 | 8.7 | 8.6 | 16.0 | 11.8 | 13.7 | 9.6 | 9.9 | 8.6 |
| | | 40 | 8.5 | 10.4 | 11.9 | 7.0 | 9.0 | 9.8 | 12.0 | 11.2 | 11.6 | 8.6 | 8.5 | 9.8 |
| | | 50 | 8.4 | 9.7 | 10.8 | 5.6 | 7.4 | 9.5 | 10.2 | 10.4 | 11.1 | 6.1 | 8.0 | 9.0 |
| | | 60 | 7.8 | 11.4 | 11.9 | 6.6 | 7.5 | 9.6 | 9.0 | 11.4 | 12.3 | 7.3 | 7.5 | 10.7 |
| | | 70 | 8.3 | 10.4 | 11.2 | 7.3 | 9.2 | 9.9 | 9.7 | 12.0 | 11.3 | 7.4 | 9.4 | 9.5 |
| | | 80 | 8.3 | 8.3 | 10.3 | 7.2 | 7.7 | 8.9 | 11.1 | 8.8 | 12.8 | 8.1 | 7.9 | 9.8 |
| | | 90 | 8.4 | 7.6 | 9.9 | 7.0 | 6.3 | 7.2 | 13.3 | 11.0 | 12.9 | 8.2 | 8.5 | 10.1 |
| | | 95 | 6.1 | 7.6 | 8.9 | 5.0 | 6.0 | 7.2 | 11.7 | 12.9 | 16.1 | 7.4 | 9.1 | 10.4 |
| | | 100 | 4.7 | 4.4 | 6.3 | 3.7 | 3.5 | 4.4 | 10.7 | 15.2 | 12.5 | 8.2 | 10.9 | 11.9 |
| 15000 | 8500 | 0 | 1.3 | 1.7 | 1.4 | 1.0 | 1.2 | 1.7 | 2.9 | 3.2 | 3.5 | 2.4 | 2.4 | 2.6 |
| | | 5 | 2.4 | 2.6 | 2.6 | 1.8 | 2.2 | 2.4 | 3.1 | 3.1 | 3.8 | 2.3 | 2.2 | 2.3 |
| | | 10 | 2.9 | 3.1 | 3.4 | 1.9 | 2.4 | 2.7 | 3.3 | 3.6 | 3.5 | 2.1 | 2.5 | 2.6 |
| | | 20 | 3.1 | 2.8 | 3.2 | 2.2 | 1.9 | 2.5 | 3.4 | 2.8 | 2.9 | 2.2 | 1.9 | 2.5 |
| | | 30 | 3.1 | 2.8 | 3.2 | 2.0 | 2.2 | 2.1 | 3.1 | 2.6 | 3.1 | 1.9 | 2.2 | 2.0 |
| | | 40 | 2.6 | 3.0 | 3.5 | 2.4 | 2.3 | 2.2 | 2.6 | 2.7 | 3.4 | 2.3 | 2.3 | 2.2 |
| | | 50 | 2.9 | 3.1 | 2.8 | 2.2 | 2.4 | 2.4 | 2.8 | 3.0 | 2.8 | 2.3 | 2.3 | 2.2 |
| | | 60 | 2.9 | 3.0 | 3.1 | 2.4 | 2.0 | 2.0 | 2.7 | 3.0 | 3.1 | 2.4 | 1.9 | 1.9 |
| | | 70 | 2.7 | 2.7 | 2.8 | 2.4 | 2.1 | 2.3 | 2.6 | 2.7 | 2.8 | 2.3 | 2.2 | 2.2 |
| | | 80 | 2.7 | 2.4 | 2.9 | 1.6 | 2.3 | 2.1 | 2.6 | 2.3 | 2.9 | 1.6 | 2.4 | 2.0 |
| | | 90 | 2.8 | 2.9 | 2.6 | 1.7 | 2.2 | 2.4 | 3.2 | 2.9 | 3.2 | 1.6 | 2.1 | 2.5 |
| | | 95 | 2.2 | 2.6 | 2.3 | 1.8 | 2.0 | 1.9 | 2.3 | 3.2 | 2.8 | 1.8 | 2.4 | 2.3 |
| | | 100 | 1.4 | 1.7 | 1.4 | 0.8 | 1.0 | 1.0 | 2.6 | 3.1 | 2.5 | 2.1 | 2.2 | 2.2 |
| 15000 | 8500 | 0 | 0.9 | 0.8 | 0.9 | 0.6 | 0.7 | 0.6 | 2.0 | 1.7 | 1.8 | | | |
| | | 5 | 1.5 | 1.2 | 1.4 | 1.0 | 1.1 | 1.3 | 1.9 | 1.4 | 1.6 | | | |
| | | 10 | 1.3 | 1.7 | 1.6 | 1.1 | 1.3 | 1.2 | 1.5 | 1.9 | 1.8 | | | |
| | | 20 | 1.5 | 1.5 | 1.5 | 1.3 | 1.4 | 1.3 | 1.5 | 1.5 | 1.6 | | | |
| | | 30 | 1.7 | 1.5 | 1.5 | 1.3 | 1.3 | 1.3 | 1.9 | 1.5 | 1.5 | | | |
| | | 40 | 1.7 | 1.6 | 1.5 | 1.2 | 1.3 | 1.2 | 1.5 | 1.7 | 1.5 | | | |
| | | 50 | 1.5 | 1.3 | 1.6 | 1.4 | 1.1 | 1.4 | 1.6 | 1.4 | 1.6 | | | |
| | | 60 | 1.2 | 1.3 | 1.7 | 1.1 | 1.1 | 1.6 | 1.3 | 1.4 | 1.6 | | | |
| | | 70 | 1.4 | 1.3 | 1.8 | 1.0 | 1.2 | 1.4 | 1.5 | 1.4 | 1.8 | | | |
| | | 80 | 1.4 | 1.2 | 1.5 | 1.3 | 1.1 | 1.3 | 1.6 | 1.4 | 1.8 | | | |
| | | 90 | 1.6 | 1.6 | 1.4 | 1.2 | 1.3 | 1.4 | 1.5 | 1.5 | 1.6 | | | |
| | | 95 | 1.6 | 1.0 | 1.6 | 1.2 | 1.1 | 1.5 | 1.6 | 1.2 | 1.9 | | | |
| | | 100 | 0.6 | 0.7 | 0.5 | 0.5 | 0.6 | 0.7 | 1.3 | 1.7 | 1.6 | | | |
| 20000 | 8500 | 0 | 0.4 | 0.5 | 0.6 | 0.3 | 0.4 | 0.4 | 0.8 | 0.8 | 1.0 | | | |
| | | 5 | 0.8 | 0.9 | 0.8 | 0.6 | 0.7 | 0.6 | 0.9 | 0.9 | 0.7 | | | |
| | | 10 | 0.9 | 0.8 | 0.9 | 0.7 | 0.6 | 0.8 | 1.0 | 0.9 | 0.9 | | | |
| | | 20 | 0.9 | 1.1 | 0.9 | 0.7 | 0.8 | 0.8 | 0.9 | 1.1 | 1.0 | | | |
| | | 30 | 0.8 | 1.0 | 0.9 | 0.8 | 0.8 | 0.9 | 0.8 | 1.1 | 0.9 | | | |
| | | 40 | 0.8 | 0.8 | 1.0 | 0.8 | 0.8 | 0.9 | 0.8 | 0.8 | 1.0 | | | |
| | | 50 | 0.9 | 0.9 | 1.0 | 0.8 | 0.8 | 0.8 | 0.9 | 1.0 | 1.0 | | | |
| | | 60 | 0.9 | 1.0 | 1.0 | 0.9 | 0.8 | 0.8 | 0.9 | 0.9 | 1.1 | | | |
| | | 70 | 0.9 | 0.6 | 0.9 | 0.7 | 0.7 | 0.9 | 0.9 | 0.7 | 0.9 | | | |
| | | 80 | 0.8 | 0.8 | 0.9 | 0.7 | 0.7 | 0.7 | 0.8 | 0.8 | 1.0 | | | |
| | | 90 | 0.7 | 0.8 | 0.7 | 0.7 | 0.8 | 0.6 | 0.7 | 1.0 | 0.7 | | | |
| | | 95 | 0.8 | 0.7 | 0.9 | 0.6 | 0.5 | 0.7 | 0.8 | 0.7 | 0.8 | | | |
| | | 100 | 0.3 | 0.3 | 0.4 | 0.2 | 0.3 | 0.3 | 0.7 | 0.7 | 0.8 | | | |

Standard deviations for admixture proportions (in percent) estimated by *qpGraph* under the "mapping" setup and by *qpAdm*. The "information-poor" or "information-rich" graphs and 95,000 or 400,000 sites were used. Results for the "information-rich" graphs are marked in grey shading.

### Table S4.

| classifier, 95K, information-poor | G ancestry in H, % | av. MCC, 0%-based | av. MCC, 100%-based | classifier, 400K, information-poor | av. MCC, 0%-based | av. MCC, 100%-based | classifier, 95K, information-rich | av. MCC, 0%-based | av. MCC, 100%-based |
|---|---|---|---|---|---|---|---|---|---|
| D-statistics Z2 | 5 | 0.39 | 0.39 | D-statistics Z2 | 0.61 | 0.60 | qpGraph LL10 | 0.93 | 0.95 |
| qpGraph LL10 | 5 | 0.30 | 0.31 | qpGraph LL10 | 0.51 | 0.51 | D-statistics Z2 | 0.93 | 0.94 |
| qpAdm/qpWave GI 0.05 | 5 | 0.27 | 0.28 | qpAdm/qpWave GI 0.05 | 0.41 | 0.41 | qpGraph WR08 | 0.84 | 0.85 |
| D-statistics Z3 | 5 | 0.24 | 0.21 | qpGraph WR08 | 0.38 | 0.39 | qpAdm/qpWave GI 0.05 | 0.84 | 0.84 |
| qpGraph WR08 | 5 | 0.23 | 0.26 | D-statistics Z3 | 0.37 | 0.37 | D-statistics Z3 | 0.83 | 0.83 |
| qpAdm/qpWave GI 0.01 | 5 | 0.23 | 0.23 | qpAdm/qpWave GI 0.01 | 0.31 | 0.31 | qpAdm/qpWave GI 0.01 | 0.81 | 0.81 |
| qpAdm/qpWave 0.05 | 5 | 0.14 | 0.15 | | | | | | |
| qpAdm/qpWave 0.01 | 5 | 0.09 | 0.09 | | | | | | |
| D-statistics Z2 | 10 | 0.77 | 0.77 | D-statistics Z2 | 0.90 | 0.90 | D-statistics Z3 | 1.00 | 1.00 |
| qpGraph LL10 | 10 | 0.75 | 0.75 | qpGraph LL10 | 0.87 | 0.87 | qpGraph WR08 | 0.99 | 1.00 |
| qpGraph WR08 | 10 | 0.66 | 0.67 | qpGraph WR08 | 0.81 | 0.82 | qpAdm/qpWave GI 0.01 | 0.99 | 0.99 |
| qpAdm/qpWave GI 0.05 | 10 | 0.60 | 0.61 | D-statistics Z3 | 0.80 | 0.80 | qpGraph LL10 | 0.97 | 1.00 |
| D-statistics Z3 | 10 | 0.60 | 0.59 | qpAdm/qpWave GI 0.01 | 0.78 | 0.77 | D-statistics Z2 | 0.97 | 0.98 |
| qpAdm/qpWave GI 0.01 | 10 | 0.58 | 0.58 | qpAdm/qpWave GI 0.05 | 0.74 | 0.74 | qpAdm/qpWave GI 0.05 | 0.96 | 0.97 |
| qpAdm/qpWave 0.05 | 10 | 0.26 | 0.28 | | | | | | |
| qpAdm/qpWave 0.01 | 10 | 0.23 | 0.28 | | | | | | |
| qpGraph LL10 | 20 | 0.94 | 0.94 | D-statistics Z2 | 0.97 | 0.96 | D-statistics Z3 | 1.00 | 1.00 |
| D-statistics Z2 | 20 | 0.94 | 0.94 | D-statistics Z3 | 0.96 | 0.96 | qpAdm/qpWave GI 0.01 | 1.00 | 1.00 |
| qpGraph WR08 | 20 | 0.92 | 0.92 | qpGraph LL10 | 0.96 | 0.96 | qpGraph WR08 | 0.99 | 1.00 |
| D-statistics Z3 | 20 | 0.92 | 0.91 | qpGraph WR08 | 0.94 | 0.95 | qpGraph LL10 | 0.97 | 1.00 |
| qpAdm/qpWave GI 0.01 | 20 | 0.83 | 0.83 | qpAdm/qpWave GI 0.01 | 0.91 | 0.90 | D-statistics Z2 | 0.97 | 0.98 |
| qpAdm/qpWave GI 0.05 | 20 | 0.82 | 0.82 | qpAdm/qpWave GI 0.05 | 0.88 | 0.88 | qpAdm/qpWave GI 0.05 | 0.95 | 0.96 |
| qpAdm/qpWave 0.05 | 20 | 0.38 | 0.39 | | | | | | |
| qpAdm/qpWave 0.01 | 20 | 0.34 | 0.38 | | | | | | |
| qpGraph LL10 | 30 | 0.97 | 0.97 | D-statistics Z3 | 0.99 | 0.99 | D-statistics Z3 | 1.00 | 1.00 |
| D-statistics Z3 | 30 | 0.97 | 0.96 | D-statistics Z2 | 0.98 | 0.97 | qpGraph WR08 | 0.99 | 1.00 |
| D-statistics Z2 | 30 | 0.97 | 0.97 | qpGraph LL10 | 0.98 | 0.98 | qpAdm/qpWave GI 0.01 | 0.99 | 0.99 |
| qpGraph WR08 | 30 | 0.96 | 0.96 | qpAdm/qpWave GI 0.01 | 0.97 | 0.96 | qpGraph LL10 | 0.97 | 1.00 |
| qpAdm/qpWave GI 0.01 | 30 | 0.93 | 0.93 | qpGraph WR08 | 0.96 | 0.97 | D-statistics Z2 | 0.97 | 0.98 |
| qpAdm/qpWave GI 0.05 | 30 | 0.89 | 0.89 | qpAdm/qpWave GI 0.05 | 0.94 | 0.94 | qpAdm/qpWave GI 0.05 | 0.93 | 0.94 |
| qpAdm/qpWave 0.05 | 30 | 0.48 | 0.48 | | | | | | |
| qpAdm/qpWave 0.01 | 30 | 0.42 | 0.47 | | | | | | |
| D-statistics Z3 | 40 | 0.99 | 0.99 | D-statistics Z3 | 1.00 | 1.00 | D-statistics Z3 | 1.00 | 1.00 |
| qpGraph LL10 | 40 | 0.98 | 0.98 | qpAdm/qpWave GI 0.01 | 0.99 | 0.98 | qpAdm/qpWave GI 0.01 | 0.99 | 0.99 |
| qpAdm/qpWave GI 0.01 | 40 | 0.97 | 0.97 | D-statistics Z2 | 0.98 | 0.97 | qpGraph WR08 | 0.99 | 1.00 |
| D-statistics Z2 | 40 | 0.97 | 0.97 | qpGraph LL10 | 0.98 | 0.98 | qpGraph LL10 | 0.97 | 1.00 |
| qpGraph WR08 | 40 | 0.96 | 0.96 | qpGraph WR08 | 0.97 | 0.98 | D-statistics Z2 | 0.97 | 0.98 |
| qpAdm/qpWave GI 0.05 | 40 | 0.93 | 0.94 | qpAdm/qpWave GI 0.05 | 0.94 | 0.94 | qpAdm/qpWave GI 0.05 | 0.91 | 0.92 |
| qpAdm/qpWave 0.05 | 40 | 0.55 | 0.56 | | | | | | |
| qpAdm/qpWave 0.01 | 40 | 0.47 | 0.48 | | | | | | |
| D-statistics Z3 | 50 | 1.00 | 0.99 | D-statistics Z3 | 1.00 | 1.00 | D-statistics Z3 | 1.00 | 1.00 |
| qpGraph LL10 | 50 | 0.98 | 0.98 | qpAdm/qpWave GI 0.01 | 0.99 | 0.99 | qpAdm/qpWave GI 0.01 | 0.99 | 0.99 |
| qpAdm/qpWave GI 0.01 | 50 | 0.98 | 0.98 | D-statistics Z2 | 0.98 | 0.97 | qpGraph WR08 | 0.99 | 1.00 |
| qpGraph WR08 | 50 | 0.97 | 0.97 | qpGraph LL10 | 0.98 | 0.98 | qpGraph LL10 | 0.97 | 1.00 |
| D-statistics Z2 | 50 | 0.97 | 0.97 | qpGraph WR08 | 0.97 | 0.98 | D-statistics Z2 | 0.97 | 0.98 |
| qpAdm/qpWave GI 0.05 | 50 | 0.93 | 0.94 | qpAdm/qpWave GI 0.05 | 0.93 | 0.93 | qpAdm/qpWave GI 0.05 | 0.93 | 0.94 |
| qpAdm/qpWave 0.05 | 50 | 0.63 | 0.63 | | | | | | |
| qpAdm/qpWave 0.01 | 50 | 0.49 | 0.55 | | | | | | |
| D-statistics Z3 | 60 | 1.00 | 0.99 | D-statistics Z3 | 1.00 | 1.00 | D-statistics Z3 | 1.00 | 1.00 |
| qpGraph LL10 | 60 | 0.98 | 0.98 | qpAdm/qpWave GI 0.01 | 0.98 | 0.98 | qpAdm/qpWave GI 0.01 | 1.00 | 1.00 |
| qpGraph WR08 | 60 | 0.98 | 0.98 | D-statistics Z2 | 0.98 | 0.97 | qpGraph WR08 | 0.99 | 1.00 |
| D-statistics Z2 | 60 | 0.97 | 0.97 | qpGraph LL10 | 0.98 | 0.98 | qpGraph LL10 | 0.97 | 1.00 |
| qpAdm/qpWave GI 0.01 | 60 | 0.96 | 0.96 | qpGraph WR08 | 0.97 | 0.98 | D-statistics Z2 | 0.97 | 0.98 |
| qpAdm/qpWave GI 0.05 | 60 | 0.92 | 0.92 | qpAdm/qpWave GI 0.05 | 0.93 | 0.93 | qpAdm/qpWave GI 0.05 | 0.96 | 0.96 |
| qpAdm/qpWave 0.05 | 60 | 0.59 | 0.60 | | | | | | |
| qpAdm/qpWave 0.01 | 60 | 0.54 | 0.61 | | | | | | |
| D-statistics Z3 | 70 | 0.97 | 0.97 | D-statistics Z3 | 0.99 | 1.00 | D-statistics Z3 | 1.00 | 1.00 |
| D-statistics Z2 | 70 | 0.96 | 0.97 | D-statistics Z2 | 0.98 | 0.97 | qpGraph WR08 | 0.99 | 1.00 |
| qpGraph LL10 | 70 | 0.96 | 0.96 | qpGraph LL10 | 0.98 | 0.98 | qpAdm/qpWave GI 0.01 | 0.98 | 0.98 |
| qpGraph WR08 | 70 | 0.93 | 0.93 | qpAdm/qpWave GI 0.01 | 0.97 | 0.97 | qpGraph LL10 | 0.97 | 1.00 |
| qpAdm/qpWave GI 0.01 | 70 | 0.92 | 0.92 | qpGraph WR08 | 0.95 | 0.96 | D-statistics Z2 | 0.97 | 0.98 |
| qpAdm/qpWave GI 0.05 | 70 | 0.91 | 0.91 | qpAdm/qpWave GI 0.05 | 0.94 | 0.94 | qpAdm/qpWave GI 0.05 | 0.95 | 0.95 |
| qpAdm/qpWave 0.05 | 70 | 0.53 | 0.53 | | | | | | |
| qpAdm/qpWave 0.01 | 70 | 0.45 | 0.46 | | | | | | |
| D-statistics Z2 | 80 | 0.92 | 0.92 | D-statistics Z2 | 0.96 | 0.96 | D-statistics Z3 | 1.00 | 1.00 |
| qpGraph LL10 | 80 | 0.88 | 0.88 | qpGraph LL10 | 0.94 | 0.94 | qpGraph WR08 | 0.99 | 1.00 |
| D-statistics Z3 | 80 | 0.87 | 0.86 | D-statistics Z3 | 0.93 | 0.93 | qpAdm/qpWave GI 0.01 | 0.98 | 0.98 |
| qpGraph WR08 | 80 | 0.80 | 0.81 | qpAdm/qpWave GI 0.01 | 0.88 | 0.87 | qpGraph LL10 | 0.97 | 1.00 |
| qpAdm/qpWave GI 0.05 | 80 | 0.80 | 0.80 | qpAdm/qpWave GI 0.05 | 0.87 | 0.87 | D-statistics Z2 | 0.97 | 0.98 |
| qpAdm/qpWave GI 0.01 | 80 | 0.79 | 0.79 | qpGraph WR08 | 0.87 | 0.88 | qpAdm/qpWave GI 0.05 | 0.94 | 0.94 |
| qpAdm/qpWave 0.05 | 80 | 0.40 | 0.41 | | | | | | |
| qpAdm/qpWave 0.01 | 80 | 0.39 | 0.45 | | | | | | |
| D-statistics Z2 | 90 | 0.72 | 0.73 | D-statistics Z2 | 0.83 | 0.82 | D-statistics Z3 | 1.00 | 1.00 |
| qpGraph LL10 | 90 | 0.60 | 0.60 | qpGraph LL10 | 0.72 | 0.72 | qpGraph WR08 | 0.99 | 1.00 |
| D-statistics Z3 | 90 | 0.58 | 0.57 | D-statistics Z3 | 0.69 | 0.70 | qpAdm/qpWave GI 0.01 | 0.99 | 0.99 |
| qpAdm/qpWave GI 0.05 | 90 | 0.53 | 0.54 | qpAdm/qpWave GI 0.05 | 0.65 | 0.65 | qpGraph LL10 | 0.97 | 1.00 |
| qpAdm/qpWave GI 0.01 | 90 | 0.49 | 0.49 | qpAdm/qpWave GI 0.01 | 0.62 | 0.61 | D-statistics Z2 | 0.97 | 0.98 |
| qpGraph WR08 | 90 | 0.49 | 0.50 | qpGraph WR08 | 0.61 | 0.62 | qpAdm/qpWave GI 0.05 | 0.93 | 0.94 |
| qpAdm/qpWave 0.05 | 90 | 0.29 | 0.31 | | | | | | |
| qpAdm/qpWave 0.01 | 90 | 0.22 | 0.24 | | | | | | |
| D-statistics Z2 | 95 | 0.42 | 0.42 | D-statistics Z2 | 0.55 | 0.54 | qpGraph LL10 | 0.94 | 0.96 |
| D-statistics Z3 | 95 | 0.33 | 0.24 | D-statistics Z3 | 0.40 | 0.41 | D-statistics Z2 | 0.93 | 0.94 |
| qpGraph LL10 | 95 | 0.31 | 0.28 | qpGraph LL10 | 0.39 | 0.39 | qpGraph WR08 | 0.85 | 0.86 |
| qpAdm/qpWave GI 0.01 | 95 | 0.27 | 0.32 | qpAdm/qpWave GI 0.05 | 0.37 | 0.37 | qpAdm/qpWave GI 0.01 | 0.85 | 0.85 |
| qpAdm/qpWave GI 0.05 | 95 | 0.22 | 0.25 | qpAdm/qpWave GI 0.01 | 0.35 | 0.30 | qpAdm/qpWave GI 0.05 | 0.84 | 0.85 |
| qpGraph WR08 | 95 | 0.19 | 0.25 | qpGraph WR08 | 0.26 | 0.28 | D-statistics Z3 | 0.83 | 0.83 |
| qpAdm/qpWave 0.05 | 95 | 0.12 | 0.15 | | | | | | |
| qpAdm/qpWave 0.01 | 95 | 0.10 | 0.13 | | | | | | |

MCC averaged across all setting combinations for each classifier and ancestry proportion. The results are shown for three graph/dataset combinations: 1/ "information-poor", 95,000 sites sampled; 2/ "information-poor", 400,000 sites sampled; and 3/ "information-rich", 95,000 sites sampled. The classifiers are sorted in the order of decreasing average MCC.

#### Table S5.

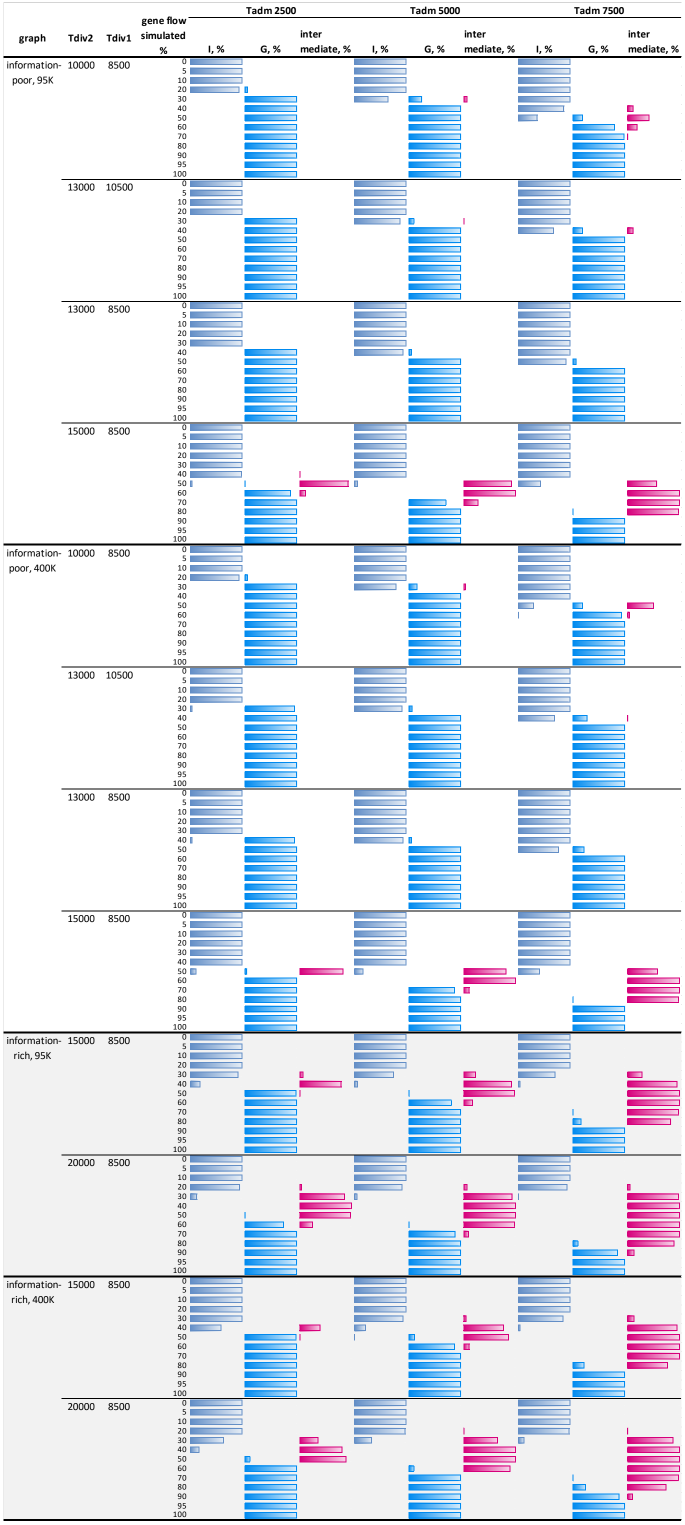


Fraction of best one-way admixture graph models with ancestry sources on the I, G, or intermediate branches. The results are shown for four graph/dataset combinations tested: "information-poor" and "information-rich" graphs, 95,000 or 400,000 sites sampled. Results for the "information-rich" graphs are marked in grey shading.

**Table S6.**

| | | information-rich | | information-poor | | average FP |
|---|---|---|---|---|---|---|
| metric | threshold | 95K | 400K | 95K | 400K | rate |
| LL | 3 | 38.2% | 37.8% | 32.2% | 32.8% | **35.3%** |
| LL | 4 | 23.0% | 26.0% | 21.0% | 22.3% | **23.1%** |
| LL | 5 | 13.7% | 15.7% | 13.5% | 14.1% | **14.2%** |
| LL | 6 | 7.5% | 10.2% | 9.2% | 9.1% | **9.0%** |
| LL | 7 | 4.7% | 7.0% | 6.1% | 5.8% | **5.9%** |
| **LL** | **8** | 3.7% | 3.5% | 3.9% | 3.8% | **3.7%** |
| LL | 9 | 2.7% | 2.3% | 2.6% | 3.2% | **2.7%** |
| **LL** | **10** | 1.5% | 1.2% | 1.7% | 2.0% | **1.6%** |
| LL | 11 | 0.7% | 0.2% | 1.1% | 1.4% | **0.8%** |
| LL | 12 | 0.5% | 0.2% | 0.8% | 0.8% | **0.6%** |
| SR | 5% | 23.5% | 28.8% | 34.0% | 34.6% | **30.2%** |
| SR | 10% | 5.3% | 6.2% | 11.2% | 10.8% | **8.4%** |
| SR | 15% | 1.5% | 1.5% | 3.4% | 4.5% | **2.7%** |
| SR | 20% | 0.3% | 0.5% | 1.8% | 2.2% | **1.2%** |
| SSR | 5% | 55.0% | 54.0% | 57.2% | 58.5% | **56.2%** |
| SSR | 15% | 12.7% | 13.2% | 18.2% | 19.3% | **15.8%** |
| SSR | 30% | 1.5% | 1.7% | 4.6% | 4.8% | **3.1%** |
| SSR | 40% | 0.2% | 0.5% | 2.0% | 2.6% | **1.3%** |
| WR | 0.1 | 22.8% | 27.8% | 27.8% | 27.4% | **26.5%** |
| WR | 0.2 | 15.0% | 17.3% | 19.8% | 19.8% | **18.0%** |
| WR | 0.3 | 9.7% | 12.0% | 13.1% | 13.9% | **12.2%** |
| WR | 0.4 | 6.0% | 7.2% | 9.3% | 10.6% | **8.3%** |
| WR | 0.5 | 4.2% | 4.5% | 6.3% | 7.6% | **5.6%** |
| WR | 0.6 | 2.7% | 3.0% | 4.3% | 5.3% | **3.8%** |
| **WR** | **0.7** | 1.3% | 1.3% | 2.7% | 3.8% | **2.3%** |
| **WR** | **0.8** | 0.5% | 0.8% | 1.7% | 2.8% | **1.5%** |
| WR | 0.9 | 0.3% | 0.5% | 1.3% | 1.8% | **1.0%** |
| WR | 1 | 0.2% | 0.5% | 1.1% | 0.9% | **0.7%** |

False positive rates for various *qpGraph*-based classifiers averaged across four graph/dataset combinations tested: "information-poor" and "information-rich" graphs, 95,000 or 400,000 sites sampled.

#### Table S7

| gene flow simulated, % | percentile | information-poor 95K | 400K | information-rich 95K | 400K |
|---|---|---|---|---|---|
| 0 | 95% | 7.8 | 7.2 | 7.6 | 7.6 |
| 5 | 5% | 1.0 | 1.3 | 10.5 | 16.9 |
| 10 | 5% | 2.5 | 5.0 | 44.2 | 72.5 |
| 20 | 5% | 10.9 | 17.0 | 183.3 | 282.4 |
| 30 | 5% | 19.5 | 33.5 | 398.6 | 633.3 |
| 40 | 5% | 31.6 | 51.4 | 639.8 | 1002.8 |
| 50 | 5% | 35.8 | 62.4 | 729.2 | 1198.7 |
| 60 | 5% | 26.0 | 43.9 | 705.2 | 1099.2 |
| 70 | 5% | 14.0 | 23.7 | 458.8 | 721.7 |
| 80 | 5% | 5.5 | 10.3 | 222.7 | 350.0 |
| 90 | 5% | 1.3 | 1.6 | 52.8 | 88.7 |
| 95 | 5% | 0.5 | 0.8 | 10.2 | 18.6 |
| 100 | 95% | 7.4 | 7.4 | 6.1 | 7.3 |

95% and 5% percentiles for the LL difference calculated across four graph/dataset combinations tested: "information-poor" and "information-rich" graphs, 95,000 or 400,000 sites sampled. Values above 8 are highlighted in green.

**Table S8**

| Group name | Effective population size, ind. | Sample size, diploid individuals | Date, cal yr BP |
|---|---|---|---|
| Neanderthal | 3000 | 1 | 50000 BP |
| Bantu | 22500 | 2 | Present |
| Ust'-Ishim | 1500 | 1 | 45000 BP |
| Jomon | 2000 | 1 | 2500 BP |
| Tibetan Chokhopani | 2000 | 1 | 2700 BP |
| SEA | 4900 | 3 | Present |
| Houtaomuga | 1500 | 3 | 12000 BP |
| Mal'ta | 2700 | 1 | 24000 BP |
| USR1 | 1100 | 1 | 11500 BP |
| FAM | 3000 | 13 | Present |
| Saqqaq | 670 | 1 | 3900 BP |
| WHG | 2700 | 11 | 13102 BP |
| Shamanka EN | 2000 | 10 | 7541 BP |
| Nganasan | 2000 | 2 | Present |
| Selkup | 2000 | 3 | Present |
| Ghost 1 | 3800 | - | - |
| Ghost 2 | 2000 | - | - |
| Ghost 3 | 1100 | - | - |
| Ghost 4 | 2000 | - | - |
| Ghost 5 | 2000 | - | - |
| Ghost 6 | 2000 | - | - |
| Ghost 7 | 2000 | - | - |
| Ghost 8 | 1100 | - | - |

Parameters of the simulated populations for the complex simulated history (Fig. S21).

### Table S9

| setting | MA adm. in FAM,% | Hou. adm. in USR1,% | 400,000 sites, 3 vs. 2-way | | | | 400,000 sites, 4 vs. 3-way | | | | 95,000 sites, 3 vs. 2-way | | | | 95,000 sites, 4 vs. 3-way | | | | |
|---|---|---|---|---|---|---|---|---|---|---|---|---|---|---|---|---|---|---|---|
| | | | LL diff. percentile | WR diff. percentile | LL diff. percentile | WR diff. percentile | LL diff. percentile | WR diff. percentile | LL diff. percentile | WR diff. percentile | LL diff. percentile | WR diff. percentile | LL diff. percentile | WR diff. percentile | LL diff. percentile | WR diff. percentile | LL diff. percentile | WR diff. percentile | |
| USR1 model(s) compared | | | FAM+MA<br>FAM+MA+X | | FAM+MA<br>FAM+MA+Hou. | | FAM+MA+X<br>FAM+MA+X+Y | | FAM+MA+Hou.<br>FAM+MA+Hou.+Y | | FAM+MA<br>FAM+MA+X | | FAM+MA<br>FAM+MA+Hou. | | FAM+MA+X<br>FAM+MA+X+Y | | FAM+MA+Hou.<br>FAM+MA+Hou.+Y | | |
| locked | 30 | 0 | 11.3 | 0.09 | 5.4 | 0.01 | 7.5 | 0.02 | 11.5 | 0.08 | 10.1 | 0.17 | 5.5 | 0.07 | 5.6 | 0.04 | 10.7 | 0.15 | 95th percentile |
| | 32 | 0 | 9.7 | 0.05 | 2.8 | 0.01 | 6.1 | 0.02 | 10.2 | 0.05 | 8.4 | 0.05 | 4.4 | 0.03 | 6.6 | 0.02 | 9.1 | 0.04 | |
| | 40 | 0 | 8.8 | 0.05 | 5.5 | 0.01 | 6.5 | 0.04 | 7.7 | 0.05 | 8.9 | 0.10 | 4.4 | 0.02 | 7.4 | 0.06 | 11.5 | 0.19 | |
| locked32 | 30 | 0 | 15.8 | 0.19 | 1.6 | 0.02 | 8.0 | 0.05 | 16.0 | 0.17 | 10.0 | 0.24 | 3.4 | 0.02 | 6.0 | 0.04 | 12.5 | 0.25 | |
| | 40 | 0 | 45.1 | 0.10 | 45.1 | 0.08 | 16.5 | 0.08 | 19.6 | 0.08 | 27.0 | 0.09 | 26.2 | 0.08 | 15.2 | 0.05 | 17.1 | 0.08 | |
| unlocked | 30 | 0 | 13.0 | 0.21 | 7.3 | 0.04 | 7.3 | 0.08 | 14.0 | 0.20 | 9.1 | 0.22 | 7.0 | 0.08 | 5.4 | 0.09 | 11.2 | 0.27 | |
| | 32 | 0 | 10.3 | 0.08 | 3.2 | 0.03 | 5.3 | 0.08 | 11.5 | 0.08 | 9.4 | 0.20 | 4.1 | 0.07 | 6.5 | 0.04 | 10.6 | 0.14 | |
| | 40 | 0 | 9.5 | 0.14 | 5.5 | 0.05 | 6.1 | 0.06 | 8.8 | 0.13 | 8.5 | 0.19 | 4.8 | 0.06 | 6.7 | 0.08 | 10.6 | 0.21 | |
| locked | 30 | 3.5 | 1.6 | 0.00 | 0.0 | -0.04 | 9.2 | 0.03 | 14.9 | 0.12 | 1.4 | 0.00 | 0.0 | -0.05 | 7.3 | 0.04 | 9.1 | 0.24 | 5th percentile |
| | | 7 | 2.0 | 0.00 | 0.3 | -0.03 | 10.0 | 0.03 | 11.6 | 0.06 | 1.6 | 0.00 | 0.4 | -0.03 | 7.3 | 0.05 | 7.3 | 0.06 | |
| | | 14 | 11.8 | 0.00 | 11.8 | -0.03 | 11.9 | 0.07 | 11.9 | 0.08 | 3.4 | 0.00 | 3.4 | -0.03 | 10.8 | 0.09 | 11.0 | 0.09 | |
| | | 35 | 121.3 | 3.39 | 121.3 | 3.39 | 11.3 | 0.04 | 11.3 | 0.04 | 46.6 | 1.60 | 46.6 | 1.60 | 10.2 | 0.14 | 10.2 | 0.14 | |
| | 32 | 3.5 | 1.5 | 0.00 | 0.0 | -0.02 | 7.5 | 0.03 | 8.7 | 0.03 | 0.9 | 0.00 | 0.0 | -0.02 | 6.6 | 0.04 | 9.9 | 0.09 | |
| | | 7 | 3.6 | 0.00 | 3.6 | -0.01 | 8.5 | 0.02 | 9.3 | 0.03 | 1.4 | 0.00 | 0.0 | -0.04 | 10.3 | 0.06 | 10.3 | 0.11 | |
| | | 14 | 10.5 | 0.00 | 10.5 | -0.02 | 11.7 | 0.09 | 11.7 | 0.14 | 5.7 | 0.00 | 3.9 | -0.04 | 11.2 | 0.07 | 13.0 | 0.15 | |
| | | 35 | 108.3 | 3.09 | 108.3 | 3.09 | 11.9 | 0.03 | 11.9 | 0.03 | 48.4 | 1.84 | 48.4 | 1.82 | 10.0 | 0.06 | 10.0 | 0.08 | |
| | 40 | 3.5 | 1.5 | 0.00 | 0.0 | -0.02 | 8.8 | 0.04 | 10.7 | 0.05 | 1.4 | 0.00 | 0.0 | -0.05 | 5.9 | 0.09 | 8.5 | 0.17 | |
| | | 7 | 2.5 | 0.00 | 1.3 | -0.02 | 7.6 | 0.02 | 8.4 | 0.02 | 2.0 | 0.00 | 0.2 | -0.02 | 8.7 | 0.05 | 8.9 | 0.11 | |
| | | 14 | 15.2 | 0.00 | 15.2 | -0.01 | 11.5 | 0.06 | 11.5 | 0.07 | 4.8 | 0.00 | -0.2 | -0.01 | 5.0 | 0.04 | 33.9 | 1.01 | |
| | | 35 | 118.5 | 2.96 | 118.5 | 2.96 | 10.3 | 0.10 | 10.3 | 0.11 | 42.2 | 1.58 | 42.2 | 1.58 | 8.9 | 0.08 | 8.9 | 0.08 | |
| locked32 | 30 | 3.5 | 1.4 | 0.00 | 0.0 | -0.11 | 8.2 | 0.05 | 15.3 | 0.08 | 1.0 | 0.00 | 0.0 | -0.06 | 6.4 | 0.05 | 9.2 | 0.11 | |
| | | 7 | 1.2 | 0.00 | 0.1 | -0.14 | 9.5 | 0.04 | 11.3 | 0.12 | 1.5 | 0.00 | 0.0 | -0.13 | 6.6 | 0.05 | 7.6 | 0.06 | |
| | | 14 | 6.2 | 0.00 | 6.2 | -0.15 | 10.3 | 0.05 | 10.3 | 0.10 | 2.1 | 0.00 | 1.4 | -0.12 | 9.4 | 0.07 | 9.4 | 0.14 | |
| | | 35 | 103.1 | 1.71 | 103.1 | 1.71 | 9.5 | 0.09 | 9.5 | 0.10 | 35.6 | 0.63 | 35.6 | 0.63 | 7.8 | 0.07 | 7.8 | 0.08 | |
| | 40 | 3.5 | 8.9 | 0.00 | 4.1 | -0.15 | 27.4 | 0.07 | 28.5 | 0.07 | 6.8 | 0.00 | 3.7 | -0.08 | 16.5 | 0.06 | 16.6 | 0.09 | |
| | | 7 | 14.3 | 0.00 | 14.3 | -0.19 | 20.3 | 0.06 | 20.2 | 0.07 | 9.7 | 0.00 | 7.5 | -0.13 | 17.3 | 0.08 | 20.9 | 0.08 | |
| | | 14 | 42.2 | 0.00 | 42.2 | -0.23 | 24.7 | 0.07 | 24.7 | 0.11 | 26.8 | 0.00 | 26.8 | -0.14 | 16.1 | 0.07 | 16.1 | 0.11 | |
| | | 35 | 196.9 | 0.00 | 196.9 | -0.33 | 27.4 | 0.11 | 27.4 | 0.12 | 101.2 | 0.00 | 101.2 | -0.29 | 15.1 | 0.06 | 15.1 | 0.10 | |
| unlocked | 30 | 3.5 | 2.2 | 0.00 | 0.1 | -0.03 | 8.4 | 0.04 | 15.2 | 0.08 | 1.4 | 0.00 | 0.0 | -0.02 | 8.3 | 0.08 | 9.5 | 0.28 | |
| | | 7 | 2.1 | 0.00 | 0.9 | -0.05 | 9.6 | 0.08 | 12.1 | 0.09 | 1.7 | 0.00 | 0.1 | -0.04 | 6.6 | 0.11 | 7.8 | 0.12 | |
| | | 14 | 12.3 | 0.00 | 12.3 | -0.01 | 12.4 | 0.14 | 12.4 | 0.14 | 4.3 | 0.00 | 4.1 | -0.01 | 12.0 | 0.17 | 12.0 | 0.24 | |
| | | 35 | 115.7 | 3.08 | 115.7 | 3.08 | 14.4 | 0.10 | 14.4 | 0.10 | 47.0 | 1.77 | 47.0 | 1.77 | 9.5 | 0.30 | 9.5 | 0.30 | |
| | 32 | 3.5 | 1.7 | 0.00 | 0.0 | -0.02 | 8.1 | 0.05 | 8.9 | 0.09 | 1.1 | 0.00 | -1.1 | -0.04 | 6.3 | 0.11 | 10.6 | 0.22 | |
| | | 7 | 3.7 | 0.00 | 3.6 | -0.01 | 7.7 | 0.05 | 8.1 | 0.07 | 1.8 | 0.00 | 0.2 | -0.10 | 9.5 | 0.07 | 11.4 | 0.14 | |
| | | 14 | 10.4 | 0.00 | 10.4 | -0.01 | 9.7 | 0.22 | 10.3 | 0.28 | 4.3 | 0.00 | 4.2 | -0.03 | 9.3 | 0.13 | 9.3 | 0.24 | |
| | | 35 | 111.0 | 3.21 | 111.0 | 3.20 | 10.1 | 0.16 | 10.1 | 0.16 | 45.2 | 1.79 | 45.2 | 1.79 | 9.1 | 0.12 | 9.1 | 0.11 | |
| | 40 | 3.5 | 1.8 | 0.00 | -0.1 | -0.01 | 8.4 | 0.05 | 9.7 | 0.05 | 1.5 | 0.00 | -0.1 | -0.01 | 6.1 | 0.09 | 10.3 | 0.25 | |
| | | 7 | 3.3 | 0.00 | 1.5 | -0.01 | 7.2 | 0.05 | 7.2 | 0.06 | 1.8 | 0.00 | -0.1 | -0.08 | 8.7 | 0.19 | 8.9 | 0.30 | |
| | | 14 | 15.1 | 0.00 | 15.1 | 0.00 | 12.1 | 0.12 | 12.1 | 0.10 | 5.3 | 0.00 | 4.7 | -0.01 | 7.2 | 0.17 | 6.7 | 0.17 | |
| | | 35 | 114.9 | 3.26 | 114.9 | 3.26 | 10.9 | 0.09 | 10.9 | 0.08 | 39.4 | 1.86 | 39.4 | 1.86 | 10.0 | 0.19 | 10.0 | 0.19 | |
| locked | 30 | 0 | 5.3 | 0.21 | 3.1 | 0.03 | | | | | 8.4 | 0.32 | 4.8 | 0.03 | | | | | 95th percentile |
| | 32 | 0 | 7.9 | 0.20 | 2.5 | 0.05 | | | | | 7.5 | 0.14 | 3.0 | 0.05 | | | | | |
| | 40 | 0 | 13.0 | 0.53 | 3.4 | 0.05 | | | | | 9.7 | 0.31 | 4.6 | 0.06 | | | | | |
| locked32 | 30 | 0 | 6.8 | 0.15 | 1.3 | 0.02 | | | | | 9.9 | 0.16 | 3.0 | 0.01 | | | | | |
| | 40 | 0 | 30.4 | 0.13 | 30.0 | 0.10 | | | | | 19.2 | 0.11 | 18.1 | 0.09 | | | | | |
| unlocked | 30 | 0 | 5.9 | 0.30 | 3.9 | 0.04 | | | | | 8.0 | 0.39 | 5.2 | 0.13 | | | | | |
| | 32 | 0 | 6.4 | 0.23 | 1.8 | 0.06 | | | | | 7.4 | 0.31 | 2.9 | 0.08 | | | | | |
| | 40 | 0 | 11.7 | 0.54 | 2.9 | 0.14 | | | | | 10.2 | 0.67 | 6.4 | 0.12 | | | | | |
| locked | 30 | 3.5 | 1.3 | 0.00 | 0.0 | -0.02 | | | | | 0.7 | 0.00 | 0.0 | -0.04 | | | | | 5th percentile |
| | | 7 | 1.9 | 0.00 | 0.1 | -0.07 | | | | | 1.4 | 0.00 | 0.0 | -0.01 | | | | | |
| | | 14 | 8.9 | 0.00 | 7.7 | 0.00 | | | | | 2.3 | 0.00 | 1.0 | -0.06 | | | | | |
| | | 35 | 64.3 | 2.87 | 64.3 | 2.87 | | | | | 17.8 | 0.48 | 17.8 | 0.48 | | | | | |
| | 32 | 3.5 | 1.5 | 0.00 | 0.1 | -0.07 | | | | | 1.1 | 0.00 | 0.0 | -0.03 | | | | | |
| | | 7 | 2.4 | 0.00 | 1.3 | -0.05 | | | | | 1.3 | 0.00 | 0.1 | -0.04 | | | | | |
| | | 14 | 6.9 | 0.01 | 6.9 | -0.02 | | | | | 1.6 | 0.00 | 0.8 | -0.03 | | | | | |
| | | 35 | 68.2 | 2.94 | 68.2 | 2.94 | | | | | 17.5 | 0.30 | 17.5 | 0.28 | | | | | |
| | 40 | 3.5 | 1.5 | 0.00 | 0.0 | -0.05 | | | | | 0.9 | 0.00 | 0.0 | -0.09 | | | | | |
| | | 7 | 1.8 | 0.00 | 0.6 | -0.09 | <= no global MAF filtering (no ascertainment bias) | | | | 1.2 | 0.00 | 0.0 | -0.04 | <= no global MAF filtering (no ascertainment bias) | | | | |
| | | 14 | 4.5 | 0.01 | 4.3 | -0.08 | | | | | 2.1 | 0.00 | 0.8 | -0.07 | | | | | |
| | | 35 | 63.9 | 2.85 | 63.9 | 2.85 | | | | | 19.7 | 0.31 | 19.7 | 0.31 | | | | | |
| locked32 | 30 | 3.5 | 1.2 | 0.00 | 0.0 | -0.10 | | | | | 0.7 | 0.00 | 0.0 | -0.03 | | | | | |
| | | 7 | 1.7 | 0.00 | 0.0 | -0.10 | | | | | 1.0 | 0.00 | 0.0 | -0.04 | | | | | |
| | | 14 | 5.1 | 0.00 | 4.2 | -0.07 | | | | | 1.0 | 0.00 | 0.2 | -0.04 | | | | | |
| | | 35 | 50.6 | 1.35 | 50.6 | 1.35 | | | | | 12.8 | 0.14 | 12.8 | 0.13 | | | | | |
| | 40 | 3.5 | 8.2 | 0.00 | 6.0 | -0.07 | | | | | 3.0 | 0.00 | 1.4 | -0.05 | | | | | |
| | | 7 | 14.0 | 0.00 | 12.6 | -0.11 | | | | | 5.4 | 0.00 | 4.4 | -0.13 | | | | | |
| | | 14 | 31.9 | 0.00 | 31.9 | -0.16 | | | | | 9.8 | 0.00 | 9.8 | -0.10 | | | | | |
| | | 35 | 126.8 | 0.01 | 126.8 | -0.29 | | | | | 44.2 | 0.00 | 44.2 | -0.13 | | | | | |
| unlocked | 30 | 3.5 | 1.2 | 0.00 | -0.3 | -0.06 | | | | | 0.5 | 0.00 | -0.3 | -0.06 | | | | | |
| | | 7 | 1.8 | 0.00 | 0.1 | -0.06 | | | | | 1.0 | 0.00 | -0.4 | -0.07 | | | | | |
| | | 14 | 8.3 | 0.00 | 7.2 | 0.00 | | | | | 2.3 | 0.00 | 0.8 | -0.04 | | | | | |
| | | 35 | 59.1 | 2.64 | 59.1 | 2.64 | | | | | 16.5 | 0.30 | 16.5 | 0.30 | | | | | |
| | 32 | 3.5 | 1.3 | 0.00 | -0.1 | -0.08 | | | | | 0.9 | 0.00 | -0.6 | -0.05 | | | | | |
| | | 7 | 2.3 | 0.00 | 1.3 | -0.06 | | | | | 1.4 | 0.00 | -0.1 | -0.07 | | | | | |
| | | 14 | 6.5 | 0.00 | 6.5 | 0.00 | | | | | 1.7 | 0.00 | 0.5 | -0.06 | | | | | |
| | | 35 | 66.6 | 3.07 | 66.6 | 3.07 | | | | | 16.8 | 0.43 | 16.8 | 0.43 | | | | | |
| | 40 | 3.5 | 1.1 | 0.00 | -0.2 | -0.07 | | | | | 1.2 | 0.00 | -1.0 | -0.09 | | | | | |
| | | 7 | 2.2 | 0.00 | 0.7 | -0.11 | | | | | 1.0 | 0.00 | 0.0 | -0.03 | | | | | |
| | | 14 | 3.7 | 0.02 | 3.6 | -0.06 | | | | | 1.7 | 0.00 | 0.6 | -0.09 | | | | | |
| | | 35 | 56.7 | 2.85 | 56.7 | 2.85 | | | | | 16.7 | 0.06 | 16.7 | 0.01 | | | | | |

Percentiles for the distributions of key admixture graph model ranking metrics obtained on simulated data: complex history, 95,000 and 400,000 sites with the ascertainment bias (the top half of the table), 95,000 and 400,000 sites without the ascertainment bias (the bottom half of the table). LL difference was calculated between the 2-way model USR1 = $FAM_{EA}$ + Mal'ta and either the best-ranking 3-way model USR1 = $FAM_{EA}$ + Mal'ta + X or the correct 3-way model, where X = Houtaomuga. We also compared 3- and 4-way models: the best-ranking 3-way model USR1 = $FAM_{EA}$ + Mal'ta + X and the best-ranking 4-way model USR1 = $FAM_{EA}$ + Mal'ta + X + Y; USR1 = $FAM_{EA}$ + Mal'ta + Houtaomuga and USR1 = $FAM_{EA}$ + Mal'ta + Houtaomuga + Y. The same was done for WR difference. $95^{th}$ percentiles of LL and WR difference were calculated across 50 iterations for measuring specificity, and $5^{th}$ percentiles for measuring sensitivity. LL difference is shown in ln-units, and WR difference in SE units. Values of $5^{th}$ LL difference percentiles above 13 ln-units are highlighted in green (multiple model comparison). Values of $5^{th}$ LL difference percentiles above 7 ln-units are highlighted in yellow (single model comparison). Values of $5^{th}$ WR difference percentiles above 0.25 SE are highlighted in green. Values of $95^{th}$ LL difference percentiles above 13 ln-units are highlighted in red. Values of $95^{th}$ WR difference percentiles above 0.25 SE are also highlighted in red.

**Table S17.**

| *f4* (FAM, USR1; X, Y) | | X, Y | Z-score vs. 0 | additional sources: none | basal Mal'ta | Houtaomuga + b. Mal'ta |
|---|---|---|---|---|---|---|
| | | | | Z-score model/data | | |
| **Houtaomuga** | Selkups | Sib, Sib | -3.6 | 4.2 | 2.6 | 0.4 |
| **Houtaomuga** | Nganasans | Sib, Sib | -3.1 | 3.4 | 2.5 | 0.5 |
| **Shamanka EN** | Selkups | Sib, Sib | -2.7 | 3.3 | 1.8 | 1.5 |
| **Houtaomuga** | Shamanka EN | Sib, Sib | -2.0 | 2.2 | 1.6 | 0.6 |
| **Bantu Tswana** | Selkups | X, Sib | -4.0 | 4.0 | 1.6 | 1.6 |
| **Neanderthal** | Selkups | X, Sib | -3.2 | 3.2 | 1.3 | 1.3 |
| **Mal'ta** | Selkups | X, Sib | -3.2 | 2.6 | 0.5 | 0.4 |
| **Ust'-Ishim** | Selkups | X, Sib | -3.1 | 3.1 | 1.2 | 1.2 |
| **WHG** | Selkups | X, Sib | -3.1 | 2.8 | 0.0 | 0.2 |
| **Bantu Tswana** | Nganasans | X, Sib | -2.9 | 2.6 | 1.3 | 1.4 |
| **anc. Tibetan** | Selkups | X, Sib | -2.7 | 3.2 | 1.7 | 1.9 |
| **Neanderthal** | Nganasans | X, Sib | -2.5 | 2.2 | 1.1 | 1.2 |
| **Mal'ta** | Nganasans | X, Sib | -2.4 | 1.7 | 0.4 | 0.4 |
| **Ust'-Ishim** | Nganasans | X, Sib | -2.4 | 2.1 | 1.0 | 1.1 |
| **anc. Tibetan** | Nganasans | X, Sib | -2.2 | 2.4 | 1.6 | 2.0 |
| SEA | **Houtaomuga** | X, Sib | 3.0 | 3.1 | 3.1 | 0.6 |
| **Bantu Tswana** | SEA | X, Y | -2.9 | 2.3 | 2.0 | 1.6 |
| **Neanderthal** | SEA | X, Y | -2.3 | 1.9 | 1.6 | 1.3 |
| **Ust'-Ishim** | SEA | X, Y | -2.3 | 1.8 | 1.5 | 1.2 |
| SEA | **anc. Tibetan** | X, Y | 2.2 | 2.2 | 2.1 | 2.1 |
| SEA | **Mal'ta** | X, Y | 2.3 | 1.3 | 0.8 | 0.4 |

$f_4$-statistics measuring asymmetry between USR1 and First Americans (First Americans, USR1; X, Y). Only statistics deviating from 0 by more than 2 SE intervals are shown here. For each statistic deviation between fitted and observed values is shown for the admixture graph models shown in Fig. S43. Residuals above 3 SE are highlighted in violet, and those between 2 and 3 SE in green. Groups attracted to USR1 vs. First Americans are marked in bold, and Siberian groups attracted to USR1 are highlighted in yellow.

**Table S18.**

| haploid USR1, 3 Houtaomuga ind.<br>model class | skeleton graph type | sites | worst *f*-statistic residual, SE | WR difference, SE | LL difference, ln-units |
|---|---|---|---|---|---|
| 2-way baseline | 14-pop., locked adm. % | 95K | 4.1 | | |
| 3-way | | | 3.2 | 0.9 | 14.6 |
| 4-way | | | 2.7 | 0.5 | 8.8 |
| 2-way baseline | 14-pop., unlocked adm. % | | 4.1 | | |
| 3-way | | | 3.1 | 1.0 | 14.1 |
| 4-way | | | 2.6 | 0.6 | 8.4 |
| 2-way baseline | 16-pop., locked adm. % | 407K | 6.1 | | |
| 3-way | | | 6.1 | 0.0 | 21.4 |
| 4-way | | | 6.1 | 0.0 | 16.3 |
| 2-way baseline | 16-pop., unlocked adm. % | | 5.9 | | |
| 3-way | | | 5.9 | 0.0 | 22.6 |
| 4-way | | | 5.9 | 0.0 | 17.7 |

Key fit statistics for various admixture models for the USR1 individual (pseudohaploid version). The following models were compared: the best 2-way model (the baseline model "$FAM_{EA}$ + Mal'ta") (*4*), the best 3-way model "$FAM_{EA}$ + Mal'ta + X" (X=basal Mal'ta in all cases), and the best 4-way model "$FAM_{EA}$ + Mal'ta + X + Y" (X=basal Mal'ta and Y=Houtaomuga in all cases). Worst *f*-statistic residuals (WR) for the best model and the differences in log-likelihood (LL) and WR between the best models in each class are shown. The difference values that were observed for less that 5% of datasets simulated under the baseline 14-population skeleton model (under any setting combination tested) are highlighted in green. Skeleton graph type and the number of sites are also shown.

**Table S19.**

| no | topology | LL | WR | LL difference vs. the best model | WR difference vs. the best model | trifurcations in the American clade |
|---|---|---|---|---|---|---|
| 1 | ((Kolyma, Ust'-Kyakhta), (USR1, (NNA, SNA))) | 234.6 | 4.78 | 0.0 | 0.0 | 2 |
| 2 | ((Kolyma, Ust'-Kyakhta), (NNA, (SNA, USR1))) | 234.7 | 4.60 | 0.1 | -0.2 | 2 |
| **3** | **((Kolyma, Ust'-Kyakhta), (SNA, (NNA, USR1)))** | **235.3** | **4.82** | **0.7** | **0.0** | **0** |
| 4 | (Ust'-Kyakhta, (Kolyma, (SNA, (NNA, USR1)))) | 238.1 | 4.75 | 3.5 | 0.0 | 0 |
| 5 | (Kolyma, (Ust'-Kyakhta, (SNA, (NNA, USR1)))) | 238.4 | 4.75 | 3.8 | 0.0 | 1 |
| 6 | (Ust'-Kyakhta, (Kolyma, (NNA, (SNA, USR1)))) | 238.4 | 4.77 | 3.8 | 0.0 | 2 |
| 7 | (Ust'-Kyakhta, (Kolyma, (USR1, (NNA, SNA)))) | 238.7 | 4.72 | 4.1 | -0.1 | 2 |
| 8 | (Kolyma, (Ust'-Kyakhta, (USR1, (NNA, SNA)))) | 238.7 | 4.79 | 4.1 | 0.0 | 2 |
| 9 | (Kolyma, (Ust'-Kyakhta, (NNA, (SNA, USR1)))) | 238.8 | 4.80 | 4.2 | 0.0 | 2 |
| 10 | (Ust'-Kyakhta, (SNA, (USR1, (Kolyma, NNA)))) | 239.4 | 4.82 | 4.8 | 0.0 | 1 |
| 11 | (Ust'-Kyakhta, (USR1, (SNA, (Kolyma, NNA)))) | 239.7 | 4.85 | 5.2 | 0.1 | 2 |
| 12 | (Ust'-Kyakhta, ((Kolyma, NNA), (SNA, USR1))) | 239.7 | 4.84 | 5.2 | 0.1 | 2 |
| 13 | (Ust'-Kyakhta, (SNA, (NNA, (Kolyma, USR1)))) | 243.1 | 4.81 | 8.6 | 0.0 | 1 |
| 14 | (Ust'-Kyakhta, (SNA, (Kolyma, (NNA, USR1)))) | 243.2 | 4.86 | 8.6 | 0.1 | 2 |
| 15 | (Ust'-Kyakhta, ((Kolyma, SNA), (NNA, USR1))) | 243.4 | 4.85 | 8.8 | 0.1 | 2 |
| 16 | (Ust'-Kyakhta, ((Kolyma, USR1), (NNA, SNA))) | 243.5 | 4.85 | 8.9 | 0.1 | 2 |
| 17 | (Ust'-Kyakhta, (USR1, (NNA, (Kolyma, SNA)))) | 243.5 | 4.87 | 8.9 | 0.1 | 3 |
| 18 | (Ust'-Kyakhta, (NNA, (USR1, (Kolyma, SNA)))) | 243.5 | 4.84 | 8.9 | 0.1 | 3 |
| 19 | (Ust'-Kyakhta, (NNA, (SNA, (Kolyma, USR1)))) | 243.5 | 4.88 | 8.9 | 0.1 | 2 |
| 20 | (Ust'-Kyakhta, (NNA, (Kolyma, (SNA, USR1)))) | 243.5 | 4.85 | 9.0 | 0.1 | 3 |
| 21 | (Ust'-Kyakhta, (USR1, (Kolyma, (NNA, SNA)))) | 243.7 | 4.85 | 9.1 | 0.1 | 2 |
| 22 | ((NNA, SNA), (USR1, (Kolyma, Ust'-Kyakhta))) | 364.4 | 5.71 | 129.8 | 0.9 | 2 |
| 23 | (USR1, ((Kolyma, Ust'-Kyakhta), (NNA, SNA))) | 366.7 | 6.05 | 132.1 | 1.3 | 2 |
| 24 | (Kolyma, (USR1, (Ust'-Kyakhta, (NNA, SNA)))) | 373.4 | 5.72 | 138.8 | 0.9 | 3 |
| 25 | ((NNA, SNA), (Ust'-Kyakhta, (Kolyma, USR1))) | 374.4 | 5.87 | 139.8 | 1.1 | 3 |

Twenty-five best topologies for the American clade on the full dataset.

### Table S20.

| ARB source | First American source | group type | clade | baseline vs. Hou | bMA vs. bMA Hou | pos. result recorded | baseline vs. bMA | Hou vs. bMA Hou | baseline vs. bMA Hou | SNPs |
|---|---|---|---|---|---|---|---|---|---|---|
| | | | | one-tailed empirical *p*-value | | | | | | |
| 0% | | | min. | 0.110 | 0.095 | | 0.100 | 0.070 | 0.050 | 400,000 |
| | | | median | 0.360 | 0.335 | | 0.300 | 0.275 | 0.255 | 400,000 |
| | | | max. | 0.500 | 0.500 | | 0.500 | 0.495 | 0.495 | 400,000 |
| 3.5% | | | min. | 0.060 | 0.020 | | 0.105 | 0.050 | 0.020 | 400,000 |
| | | | median | 0.238 | 0.223 | | 0.340 | 0.303 | 0.195 | 400,000 |
| | | | max. | 0.500 | 0.455 | | 0.500 | 0.500 | 0.395 | 400,000 |
| 7% | | | min. | 0.010 | 0.010 | | 0.100 | 0.080 | 0.010 | 400,000 |
| | | | median | 0.120 | 0.115 | | 0.335 | 0.330 | 0.100 | 400,000 |
| | | | max. | 0.495 | 0.475 | | 0.495 | 0.495 | 0.405 | 400,000 |
| 14% | | | min. | 0.003 | 0.003 | | 0.140 | 0.080 | 0.003 | 400,000 |
| | | | median | 0.025 | 0.020 | | 0.350 | 0.335 | 0.018 | 400,000 |
| | | | max. | 0.235 | 0.215 | | 0.495 | 0.490 | 0.145 | 400,000 |
| 35% | | | min. | 0.003 | 0.003 | | 0.140 | 0.040 | 0.003 | 400,000 |
| | | | median | 0.003 | 0.003 | | 0.338 | 0.305 | 0.003 | 400,000 |
| | | | max. | 0.005 | 0.005 | | 0.500 | 0.490 | 0.005 | 400,000 |
| USR1 pseudohaploid 3 Houtaomuga ind. | Argentina Aconcagua Inca 500BP | ancient | SNA | 0.176 | 0.102 | | 0.114 | 0.058 | 0.034 | 360,220 |
| | Argentina ArroyoSeco2 7700BP | ancient | SNA | 0.138 | 0.208 | | 0.062 | 0.066 | 0.046 | 374,409 |
| | Argentina Fuego Patagonian | ancient | SNA | 0.058 | 0.116 | yes | 0.012 | 0.020 | 0.008 | 384,335 |
| | Argentina LagunaChica 6800BP | ancient | SNA | 0.238 | 0.226 | | 0.110 | 0.100 | 0.082 | 243,410 |
| | ASO | ancient | NNA | 0.114 | 0.096 | yes | 0.220 | 0.194 | 0.060 | 295,334 |
| | Aymara | present-day | SNA | 0.132 | 0.158 | | 0.048 | 0.042 | 0.034 | 377,706 |
| | Belize MayahakCabPek 9300BP | ancient | SNA | 0.104 | 0.146 | | 0.062 | 0.068 | 0.022 | 192,777 |
| | Belize SakiTzul 7400BP | ancient | SNA | 0.268 | 0.204 | | 0.300 | 0.230 | 0.180 | 203,967 |
| | Brazil LapaDoSanto 9600BP | ancient | SNA | 0.126 | 0.194 | | 0.056 | 0.072 | 0.042 | 376,490 |
| | Brazil LapaDoSanto 9600BP publ. | ancient | SNA | 0.216 | 0.228 | | 0.274 | 0.314 | 0.176 | 245,468 |
| | Brazil Laranjal 6700BP | ancient | SNA | 0.240 | 0.196 | | 0.122 | 0.092 | 0.088 | 192,898 |
| | Brazil Sumidouro 10100BP | ancient | SNA | 0.086 | 0.076 | yes | 0.234 | 0.240 | 0.070 | 407,195 |
| | Canada 6000BP | ancient | NNA | 0.214 | 0.212 | | 0.036 | 0.036 | 0.038 | 133,364 |
| | Canada BigBar 5700BP | ancient | NNA | 0.236 | 0.272 | | 0.062 | 0.080 | 0.044 | 280,059 |
| | Chane | present-day | SNA | 0.084 | 0.090 | yes | 0.098 | 0.110 | 0.030 | 405,728 |
| | Chile Ayayema 5100BP | ancient | SNA | 0.138 | 0.204 | | 0.040 | 0.066 | 0.028 | 406,086 |
| | Chile Conchali 700BP | ancient | SNA | 0.138 | 0.192 | | 0.052 | 0.068 | 0.034 | 374,201 |
| | Chile Fuego Patagonian | ancient | SNA | 0.092 | 0.134 | yes | 0.154 | 0.210 | 0.052 | 176,433 |
| | Chile Kaweskar 1000BP | ancient | SNA | 0.098 | 0.112 | yes | 0.058 | 0.062 | 0.040 | 406,225 |
| | Chile LosRieles 12000BP | ancient | SNA | 0.142 | 0.168 | | 0.128 | 0.128 | 0.080 | 336,790 |
| | Chile LosRieles 5100BP | ancient | SNA | 0.200 | 0.198 | | 0.210 | 0.204 | 0.108 | 332,940 |
| | Chile PicaOcho 700BP | ancient | SNA | 0.212 | 0.304 | | 0.022 | 0.034 | 0.016 | 226,632 |
| | Chile PuntaSantaAna 7300BP | ancient | SNA | 0.290 | 0.246 | | 0.398 | 0.338 | 0.240 | 317,431 |
| | Chile Yamana 1000BP | ancient | SNA | 0.098 | 0.122 | yes | 0.088 | 0.094 | 0.032 | 376,880 |
| | E San Nicolas | ancient | SNA | 0.106 | 0.114 | | 0.124 | 0.112 | 0.046 | 391,128 |
| | FAM 13 ind. | present-day | SNA | 0.032 | 0.044 | yes | 0.062 | 0.048 | 0.002 | 407,556 |
| | Huichol | present-day | SNA | 0.198 | 0.246 | | 0.068 | 0.078 | 0.068 | 379,530 |
| | Island Chumash SanCruz | ancient | SNA | 0.132 | 0.156 | | 0.092 | 0.096 | 0.052 | 384,011 |
| | Island Chumash SanMiguel | ancient | SNA | 0.084 | 0.158 | yes | 0.016 | 0.020 | 0.004 | 350,176 |
| | Karitiana | present-day | SNA | 0.038 | 0.038 | yes | 0.066 | 0.044 | 0.010 | 406,482 |
| | L San Nicolas | ancient | SNA | 0.062 | 0.050 | yes | 0.060 | 0.040 | 0.010 | 404,827 |
| | LSCI | ancient | SNA | 0.076 | 0.092 | yes | 0.074 | 0.060 | 0.016 | 379,284 |
| | Mainland Chumash | ancient | SNA | 0.208 | 0.204 | | 0.188 | 0.182 | 0.124 | 279,141 |
| | Mainland Chumash NewCuyama | ancient | SNA | 0.266 | 0.342 | | 0.104 | 0.108 | 0.090 | 161,149 |
| | Mixe | present-day | SNA | 0.050 | 0.074 | yes | 0.030 | 0.038 | 0.010 | 406,571 |
| | Nahua | present-day | SNA | 0.112 | 0.132 | | 0.190 | 0.198 | 0.088 | 405,851 |
| | Peru Cuncaicha 3300BP | ancient | SNA | 0.188 | 0.192 | | 0.226 | 0.202 | 0.124 | 332,149 |
| | Peru Cuncaicha 4200BP | ancient | SNA | 0.112 | 0.086 | yes | 0.158 | 0.134 | 0.050 | 315,276 |
| | Peru Cuncaicha 9000BP | ancient | SNA | 0.216 | 0.248 | | 0.106 | 0.122 | 0.080 | 311,203 |
| | Peru LaGalgada 4100BP | ancient | SNA | 0.132 | 0.156 | | 0.074 | 0.072 | 0.038 | 367,804 |
| | Peru Laramate 900BP | ancient | SNA | 0.120 | 0.054 | yes | 0.034 | 0.022 | 0.008 | 390,835 |
| | Peru Lauricocha 3500BP | ancient | SNA | 0.092 | 0.082 | yes | 0.176 | 0.148 | 0.044 | 259,382 |
| | Peru Lauricocha 5800BP | ancient | SNA | 0.180 | 0.198 | | 0.100 | 0.090 | 0.040 | 264,629 |
| | Peru Lauricocha 8600BP | ancient | SNA | 0.074 | 0.050 | yes | 0.100 | 0.072 | 0.024 | 388,829 |
| | Peru RioUncallane 1800BP | ancient | SNA | 0.082 | 0.058 | yes | 0.024 | 0.018 | 0.004 | 407,296 |
| | Piapoco | present-day | SNA | 0.088 | 0.068 | yes | 0.054 | 0.034 | 0.012 | 406,325 |
| | Pima | present-day | SNA | 0.072 | 0.068 | yes | 0.046 | 0.046 | 0.008 | 406,390 |
| | Quechua | present-day | SNA | 0.062 | 0.060 | yes | 0.088 | 0.084 | 0.018 | 406,630 |
| | Surui | present-day | SNA | 0.132 | 0.180 | | 0.066 | 0.064 | 0.054 | 406,258 |
| | USA Anzick | ancient | SNA | 0.182 | 0.242 | | 0.174 | 0.210 | 0.122 | 405,244 |
| | USA Anzick realigned | ancient | SNA | 0.166 | 0.206 | | 0.192 | 0.226 | 0.120 | 405,833 |
| | USA Nevada LovelockCave 1850BP | ancient | SNA | 0.048 | 0.072 | yes | 0.018 | 0.032 | 0.004 | 406,299 |
| | USA Nevada LovelockCave 600BP | ancient | SNA | 0.174 | 0.156 | | 0.088 | 0.084 | 0.064 | 406,077 |
| | USA Nevada SpiritCave 11000BP | ancient | SNA | 0.084 | 0.116 | yes | 0.104 | 0.110 | 0.038 | 406,092 |
| | USA WA Kennewick | ancient | NNA | 0.208 | 0.230 | | 0.272 | 0.270 | 0.154 | 213,998 |
| | Yukpa | present-day | SNA | 0.112 | 0.130 | | 0.042 | 0.044 | 0.030 | 371,426 |
| | Zapotec | present-day | SNA | 0.076 | 0.120 | yes | 0.034 | 0.042 | 0.016 | 406,497 |
| **USR1 hapl. 2 Hou** | FAM 13 ind. | present-day | SNA | 0.068 | 0.060 | yes | 0.048 | 0.040 | 0.010 | 407,407 |
| **USR1 dipl. 2 Hou** | FAM 13 ind. | present-day | SNA | 0.090 | 0.136 | yes | 0.042 | 0.056 | 0.028 | 377,481 |
| **USR1 dipl. 3 Hou** | FAM 13 ind. | present-day | SNA | 0.060 | 0.094 | yes | 0.046 | 0.060 | 0.018 | 377,622 |

Results of out-of-sample testing of admixture graph models on 200 resampled replicates of simulated (400,000 sites with ascertainment bias) and 500 resampled replicates of real data. Distributions of model-comparison empirical *p*-values (proportion of bootstrap replicates with a negative LL difference $LL_{simple\ model} - LL_{complex\ model}$) are summarized across 130 simulated datasets for each simulated ARB ancestry proportion (minimal, median, and maximal values are shown). For each combination of the USR1 target (pseudohaploid and diploid versions), the ARB source proxy (3 or 2 Houtaomuga individuals) and an American source proxy (57 alternative groups) one test on 500 bootstrap replicates was performed. Empirical *p*-values below 0.10 (the lowest value observed on data simulated under the baseline model when that model was compared to the "$FAM_{EA}$ + Mal'ta + Houtaomuga" or "$FAM_{EA}$ + Mal'ta + basal Mal'ta" models) and above 0.05 are highlighted in yellow. *P*-values below 0.05 are highlighted in green. The following abbreviations are used: bMA, basal Mal'ta; Hou, Houtaomuga.

#### Table S21.

| American clade topology | gene flows in the USR1 history | WR NT | WR | WR diff. NT | WR diff. | LL NT | LL | LL diff. NT | LL diff. | trifurcations in the FAM clade, NT | trifurcations in the FAM clade |
|---|---|---|---|---|---|---|---|---|---|---|---|
| (IL7, (proto-Athabaskan, USR1)) | Mal'ta | 5.07 | 7.32 | | | 124.5 | 270.9 | | | 1 | 1 |
| (proto-Athabaskan, (IL7, USR1)) | Mal'ta | 5.07 | 7.33 | | | 124.5 | 270.8 | | | 1 | 1 |
| (USR1, (proto-Athabaskan, IL7)) | Mal'ta | 4.09 | 5.30 | | | 109.6 | 228.3 | | | 0 | 0 |
| (IL7, (proto-Athabaskan, proto-USR1)) | Mal'ta, basal Mal'ta | 4.05 | 5.11 | | | 107.7 | 219.4 | | | 1 | 1 |
| (proto-Athabaskan, (IL7, proto-USR1)) | Mal'ta, basal Mal'ta | 4.05 | 5.11 | | | 107.7 | 219.4 | | | 0 | 1 |
| (proto-USR1, (proto-Athabaskan, IL7)) | Mal'ta, basal Mal'ta | 3.78 | 4.81 | 0.31 | 0.49 | 101.6 | 206.8 | 8.0 | 21.5 | 0 | 0 |
| (IL7, (proto-Athabaskan, proto-USR1)) | Mal'ta, Houtaomuga | 4.91 | 6.99 | | | 119.8 | 255.9 | | | 1 | 1 |
| (proto-Athabaskan, (IL7, proto-USR1)) | Mal'ta, Houtaomuga | 4.92 | 6.99 | | | 119.9 | 255.9 | | | 1 | 1 |
| (proto-USR1, (proto-Athabaskan, IL7)) | Mal'ta, Houtaomuga | 4.12 | 5.39 | -0.03 | -0.08 | 107.1 | 220.8 | 2.5 | 7.5 | 0 | 0 |
| (IL7, (proto-Athabaskan, proto-USR1)) | Mal'ta, Houtaomuga, basal Mal'ta | 3.58 | 4.81 | 0.20 | 0.00 | 92.8 | 184.3 | 8.8 | 22.5 | 1 | 0 |
| (proto-Athabaskan, (IL7, proto-USR1)) | Mal'ta, Houtaomuga, basal Mal'ta | 3.51 | 4.81 | 0.27 | 0.00 | 91.2 | 184.8 | 10.4 | 22.0 | 0 | 0 |
| (proto-USR1, (proto-Athabaskan, IL7)) | Mal'ta, Houtaomuga, basal Mal'ta | 3.58 | 4.81 | 0.20 | 0.00 | 92.8 | 184.9 | 8.8 | 21.9 | 1 | 1 |
| (IL7, (proto-Athabaskan, proto-USR1)) | Mal'ta, Houtaomuga+basal Mal'ta | 3.58 | 4.81 | | | 92.8 | 184.3 | | | 1 | 0 |
| (proto-Athabaskan, (IL7, proto-USR1)) | Mal'ta, Houtaomuga+basal Mal'ta | 3.51 | 4.81 | | | 91.2 | 184.8 | | | 0 | 0 |
| (proto-USR1, (proto-Athabaskan, IL7)) | Mal'ta, Houtaomuga+basal Mal'ta | 3.58 | 4.81 | | | 92.8 | 184.8 | | | 1 | 1 |
| (IL7, (proto-Athabaskan, proto-USR1)) | Mal'ta, Houtaomuga rev, basal Mal'ta | 3.95 | 4.81 | -0.37 | 0.00 | 103.5 | 204.9 | -10.8 | -20.6 | 1 | 1 |
| (proto-Athabaskan, (IL7, proto-USR1)) | Mal'ta, Houtaomuga rev, basal Mal'ta | 3.96 | 4.81 | -0.44 | 0.00 | 103.0 | 204.9 | -11.9 | -20.1 | 0 | 1 |
| (proto-USR1, (proto-Athabaskan, IL7)) | Mal'ta, Houtaomuga rev, basal Mal'ta | 3.81 | 4.81 | -0.23 | 0.00 | 97.8 | 195.3 | -5.1 | -10.4 | 0 | 0 |
| (FAM, (proto-Athabaskan, USR1)) | Mal'ta | 6.59 | 10.02 | | | 161.3 | 332.2 | | | 1 | 1 |
| (proto-Athabaskan, (FAM, USR1)) | Mal'ta | 6.59 | 10.03 | | | 161.3 | 332.2 | | | 1 | 1 |
| (USR1, (proto-Athabaskan, FAM)) | Mal'ta | 4.15 | 5.55 | | | 113.2 | 217.1 | | | 0 | 0 |
| (FAM, (proto-Athabaskan, proto-USR1)) | Mal'ta, basal Mal'ta | 4.69 | 6.34 | | | 130.2 | 245.5 | | | 1 | 1 |
| (proto-Athabaskan, (FAM, proto-USR1)) | Mal'ta, basal Mal'ta | 4.69 | 6.34 | | | 130.2 | 245.6 | | | 1 | 1 |
| (proto-USR1, (proto-Athabaskan, FAM)) | Mal'ta, basal Mal'ta | 3.77 | 4.80 | 0.38 | 0.75 | 106.6 | 201.0 | 6.6 | 16.1 | 0 | 0 |
| (FAM, (proto-Athabaskan, proto-USR1)) | Mal'ta, Houtaomuga | 5.69 | 7.64 | | | 142.7 | 277.2 | | | 1 | 1 |
| (proto-Athabaskan, (FAM, proto-USR1)) | Mal'ta, Houtaomuga | 5.69 | 7.64 | | | 142.7 | 277.2 | | | 1 | 1 |
| (proto-USR1, (proto-Athabaskan, FAM)) | Mal'ta, Houtaomuga | 4.16 | 5.61 | -0.01 | -0.05 | 109.3 | 206.7 | 3.9 | 10.4 | 0 | 0 |
| (FAM, (proto-Athabaskan, proto-USR1)) | Mal'ta, Houtaomuga, basal Mal'ta | 3.51 | 4.80 | 0.26 | 0.00 | 95.1 | 171.4 | 11.5 | 29.7 | 1 | 0 |
| (proto-Athabaskan, (FAM, proto-USR1)) | Mal'ta, Houtaomuga, basal Mal'ta | 3.50 | 4.80 | 0.28 | 0.00 | 95.0 | 175.4 | 11.6 | 25.7 | 0 | 1 |
| (proto-USR1, (proto-Athabaskan, FAM)) | Mal'ta, Houtaomuga, basal Mal'ta | 3.51 | 4.80 | 0.26 | 0.00 | 95.1 | 175.4 | 11.5 | 25.7 | 1 | 1 |
| (FAM, (proto-Athabaskan, proto-USR1)) | Mal'ta, Houtaomuga+basal Mal'ta | 3.51 | 4.80 | | | 95.1 | 171.4 | | | 1 | 0 |
| (proto-Athabaskan, (FAM, proto-USR1)) | Mal'ta, Houtaomuga+basal Mal'ta | 3.50 | 4.80 | | | 95.1 | 175.4 | | | 0 | 1 |
| (proto-USR1, (proto-Athabaskan, FAM)) | Mal'ta, Houtaomuga+basal Mal'ta | 3.51 | 4.80 | | | 95.1 | 175.4 | | | 1 | 1 |
| (FAM, (proto-Athabaskan, proto-USR1)) | Mal'ta, Houtaomuga rev, basal Mal'ta | 4.39 | 6.23 | -0.88 | -1.43 | 123.2 | 225.9 | -28.1 | -54.5 | 1 | 1 |
| (proto-Athabaskan, (FAM, proto-USR1)) | Mal'ta, Houtaomuga rev, basal Mal'ta | 4.39 | 6.26 | -0.90 | -1.46 | 123.3 | 225.9 | -28.2 | -50.5 | 1 | 1 |
| (proto-USR1, (proto-Athabaskan, FAM)) | Mal'ta, Houtaomuga rev, basal Mal'ta | 3.96 | 4.80 | -0.45 | 0.00 | 101.7 | 186.3 | -6.6 | -10.9 | 0 | 0 |

A summary of admixture graph models incorporating the Mal'ta-related gene flow shared by all Native Americans. Three possible topologies of the American clade are shown on the left, and gene flows in the USR1 history in the next column. WR and LL differences were calculated between the "$\text{FAM}_{\text{EA}}$ + Mal'ta" and "$\text{FAM}_{\text{EA}}$ + Mal'ta + basal Mal'ta", "$\text{FAM}_{\text{EA}}$ + Mal'ta" and "$\text{FAM}_{\text{EA}}$ + Mal'ta + Houtaomuga" models, and also between "$\text{FAM}_{\text{EA}}$ + Mal'ta + basal Mal'ta" and various 4-way models. NT, the 1240K-based dataset without transitions. LL and WR differences exceeding the cut-off values optimized on simulated data in Supplementary Text 2 are highlighted in green.

**Table S25.**

| method | data type | *p*-value | *p*-value, comment | reference |
|---|---|---|---|---|
| "mapping" USR1 on skeleton admixture graphs as derived from various numbers of all possible sources | a 1240K-based dataset, pseudohaploid and diploid, 407K variable sites | <0.05 | a significance cut-off for log-likelihood difference was set according to a simulation study, exact *p*-value is unavailable | this study |
| comparing admixture graph models using a bootstrapping algorithm and out-of-sample likelihood (Supplementary Text 11) | a 1240K-based dataset, pseudohaploid and diploid, 407K variable sites | 0.032 | a one-tailed empirical *p*-value is shown; see results for various FAM reference groups in Table S20 | this study |
| comparing admixture graph models using a bootstrapping algorithm and out-of-sample likelihood (Supplementary Text 11) | shotgun, pseudohaploid, neutrally evolving transversion sites selected according to Pouyet *et al*. (*32*), 226K variable sites | 0.05 | a one-tailed empirical *p*-value is shown | this study |
| exploration of alternative topologies for the proto-American clade | a 1240K-based dataset, pseudohaploid and diploid, 95K or 407K variable sites | N/A | according to three topology explorations (Supplementary Texts 7 section g, 9, and 12), the $FAM_{EA}$ or FAM source for USR1 was placed on the NNA branch. model selection relied on likelihood and the number of trifurcations in the American clade | this study |
| testing for cladality of USR1 and First Americans using *qpWave* | a 1240K-based dataset, pseudohaploid and diploid, 407K variable sites | $1.3\times10^{-5}$ – $7\times10^{-5}$ | a two-tailed *p*-value is shown; see results for various FAM reference groups in Data S8a | this study |

| | | | | |
|---|---|---|---|---|
| *f4*-statistic (First Americans, USR1; Southeast Asians, Houtaomuga) | a 1240K-based dataset, pseudohaploid and diploid, 407K variable sites | 0.0027 | a two-tailed *p*-value is shown | this study |
| *D*-statistics (a First American group, USR1; a Siberian group, Han) | shotgun | ~ 0.0027 | a two-tailed *p*-value is shown | Moreno-Mayar *et al*. (*4*) |
| testing simple demographic models using *diCal*, a haplotype-based coalescent method (*39*) | shotgun, diploid, first 10 chromosomes | <0.05 | a significance cut-off for log-likelihood difference was set according to a simulation study, exact *p*-value is unavailable | Moreno-Mayar *et al*. (*4*) |

Various lines of evidence suggesting the existence of Siberian admixture in the USR1 lineage in this study and those reported by Moreno-Mayar *et al*. (*4*).

**Data S1. (separate file)**

SSR vs. WR and SSR vs. LL biplots summarized for "partially correct" graphs, 2-way admixed models, all settings, and all gene flow intensities from 5 to 95% ("information-poor" graphs, 95,000 sites). Results for unadmixed models are shown for simulated gene flows of 0% and 100%. We did not pay attention to admixture proportions inferred by *qpGraph*, only to source combinations. Correct models are coded as 2, partially correct models as 1, and incorrect models as 0. Thus, a notation "2 & 1 & 0" means that correct, partially correct, and incorrect ancestry source pairs fell below the chosen WR and LL thresholds. The curve shape is described in separate columns for the WR and LL metrics, and the overall conclusion is shown in the rightmost column.

**Data S2. (separate file)**

An interactive three-dimensional plot of PC1, PC2, and PC3 based on simulated genetic data (the complex simulated history, 95,000 variable sites sampled with the 5% MAF filter). Results for 150 simulated datasets with 0% Houtaomuga admixture in USR1 (red marker border) are plotted along with those for 35% Houtaomuga admixture in USR1 (150 simulated datasets, black marker border). Each dot is labelled with its population and individual name. The dots are colored according to PC4.

**Data S3. (separate file)**

Sample information and sequencing statistics for 22 samples screened int his study. The samples selected for downstream analyses based on contamination estimates are shown in bold.

**Data S4. (separate file)**

Results of admixture graph mapping for the USR1 individual. (**A**) Mapping the diploid and pseudohaploid USR1 versions on the 14-population skeleton graph under 8 setting combinations, relying on the 1240K-based dataset without transitions. The default Mal'ta source is not shown. 2-, 3-, and 4-way admixed models are shown. (**B**) Mapping the diploid and pseudohaploid USR1 versions on the 16-population skeleton graph under 4 setting combinations, relying on the 1240K-based dataset including transitions. The default Mal'ta source is not shown. 2-, 3-, and 4-way admixed models are shown. (**C**) Mapping the pseudohaploid USR1 version on the 14-population skeleton graph under 8 setting combinations, relying on the 1240K-based dataset without transitions. Only the $FAM_{EA}$ source, but not the Mal'ta source, was set by default, and the default source is not shown. 2-, 3-, 4-, and 5-way admixed models are shown, but most low-ranking 4- and 5-way models are omitted to meet the file size limit. (**D**) Mapping the pseudohaploid USR1 version on the 16-population skeleton graph under 4 setting combinations, relying on the 1240K-based dataset including transitions. Only the $FAM_{EA}$ source, but not the Mal'ta source, was set by default, and the default source is not shown. 2-, 3-, and 4-way admixed models are shown, but most low-ranking 4-way models are omitted to meet the file size limit. (**E**) Exploring the likelihood space of admixture proportions under the final 4-way admixture model selected for USR1. Results are shown for 4 dataset/skeleton graph/setting combinations, and most low-ranking models are omitted to meet the file size limit. (**F**) Exploring all possible topologies of $FAM_{EA}$ sources for USR1, ancient Athabaskan I5321, and two SNA individuals (Anzick and IL7), with non-American gene flows added according to the best models for these individuals. Anzick was modelled as having Siberian and "basal Mal'ta" admixture. Results for

48 dataset/skeleton graph/setting combinations are shown. (**G**) Exploring all possible topologies of $FAM_{EA}$ sources for USR1, ancient Athabaskan I5321, and two SNA individuals (Anzick and IL7), with non-American gene flows added according to the best models for these individuals. Anzick was modelled as having no gene flows from extra-American sources. Results for 48 dataset/skeleton graph/setting combinations are shown.

**Data S5. (separate file)**

An interactive three-dimensional plot visualizing model ranking in the likelihood space of admixture proportions under the final 4-way admixture model selected for USR1. Best-ranking models are colored according to the "final rank" column in Data S4e.

**Data S6. (separate file)**

Summary of graph mapping results for other ancient American individuals and present-day groups. Only highest-ranking models are shown for each target. Results for two setups and shown: the 1240K-based dataset without transitions and the 14-population skeleton graph; the 1240K-based dataset including transitions and the 16-population skeleton graph

**Data S7. (separate file)**

Results of admixture graph mapping for the Kolyma1 and Ust'-Kyakhta individuals. (**A**) Mapping the diploid and pseudohaploid Kolyma1 versions and the pseudohaploid Ust'-Kyakhta version on the 14-population skeleton graph under 8 setting combinations, relying on the 1240K-based dataset without transitions. The default Mal'ta source is not shown. 2-, 3-, and 4-way admixed models are shown, but most low-ranking 4-way models are omitted to meet the file size limit. (**B**) Mapping the diploid and pseudohaploid Kolyma1 versions and the pseudohaploid Ust'-Kyakhta version on the 16-population skeleton graph under 4 setting combinations, relying on the 1240K-based dataset including transitions. The default Mal'ta source is not shown. 2-, 3-, and 4-way admixed models are shown, but most low-ranking 4-way models are omitted to meet the file size limit.

**Data S8. (separate file)**

*qpWave* and *qpAdm* results. Results for the diploid and pseudo-haploid USR1 versions and for the Houtaomuga group composed of two or three individuals are shown side by side. (**A**) Testing the cladality of USR1 and various FAM ancient individuals and present-day groups using *qpWave*. The 1240K-based dataset including transitions was used for this analysis. (**B**) Testing the cladality of all possible group pairs on the 14-population skeleton graph using *qpWave*. The 1240K-based dataset including transitions was used for this analysis. (**C**) Testing all possible 2-way models for the USR1, FAM, and Houtaomuga target groups in the context of the 14-population skeleton graph. The 1240K-based dataset including transitions was used for this analysis. (**D**) Testing all possible 3-way models for the USR1, FAM, and Houtaomuga target groups in the context of the 14-population skeleton graph. The 1240K-based dataset including transitions was used for this analysis. (**E**) Testing the cladality of all possible group pairs on the 14-population skeleton graph using *qpWave*. The 1240K-based dataset without transitions was used for this analysis. (**F**) Testing all possible 2-way models for the USR1, FAM, and Houtaomuga target groups in the context of the 14-population skeleton graph. The 1240K-based dataset without transitions was used for this analysis. (**G**) Testing all possible 3-way models for the USR1, FAM, and Houtaomuga target groups in the context of the 14-population skeleton graph. The 1240K-based dataset without transitions was used for this analysis.

**Data S9. (separate file)**

A list of ancient individuals (with published whole-genome data) included into the analyses relying on the set neutrally evolving sites.

**Data S10. (separate file)**

Graph mapping results for Siberian, Central and East Asian ancient individuals and present-day groups mapped on the 14-population skeleton graph. The results are based on the 1240K dataset without transitions. (**A**) Metadata for target groups and model fit statistics. Only highest-ranking models are shown for each target. (**B**) Breakdown of target genomes into ancestry components according to the best-ranking models.
